# Unveiling Crucivirus Diversity by Mining Metagenomic Data

**DOI:** 10.1101/2020.02.27.967703

**Authors:** Ignacio de la Higuera, George W. Kasun, Ellis L. Torrance, Alyssa A. Pratt, Amberlee Maluenda, Jonathan Colombet, Maxime Bisseux, Viviane Ravet, Anisha Dayaram, Daisy Stainton, Simona Kraberger, Peyman Zawar-Reza, Sharyn Goldstien, James V. Briskie, Robyn White, Helen Taylor, Christopher Gomez, David G. Ainley, Jon S. Harding, Rafaela S. Fontenele, Joshua Schreck, Simone G. Ribeiro, Stephen A. Oswald, Jennifer M. Arnold, François Enault, Arvind Varsani, Kenneth M. Stedman

## Abstract

The discovery of cruciviruses revealed the most explicit example of a common protein homologue between DNA and RNA viruses to date. Cruciviruses are a novel group of circular Rep-encoding ssDNA (CRESS-DNA) viruses that encode capsid proteins (CPs) that are most closely related to those encoded by RNA viruses in the family *Tombusviridae*. The apparent chimeric nature of the two core proteins encoded by crucivirus genomes suggests horizontal gene transfer of CP genes between DNA and RNA viruses. Here, we identified and characterized 451 new crucivirus genomes and ten CP-encoding circular genetic elements through *de novo* assembly and mining of metagenomic data. These genomes are highly diverse, as demonstrated by sequence comparisons and phylogenetic analysis of subsets of the protein sequences they encode. Most of the variation is reflected in the replication associated protein (Rep) sequences, and much of the sequence diversity appears to be due to recombination. Our results suggest that recombination tends to occur more frequently among groups of cruciviruses with relatively similar capsid proteins, and that the exchange of Rep protein domains between cruciviruses is rarer than gene exchange. Altogether, we provide a comprehensive and descriptive characterization of cruciviruses.

**IMPORTANCE:** Viruses are the most abundant biological entities on Earth. In addition to their impact on animal and plant health, viruses have important roles in ecosystem dynamics as well as in the evolution of the biosphere. Circular Rep-encoding single-stranded (CRESS) DNA viruses are ubiquitous in nature, many are agriculturally important, and are viruses that appear to have multiple origins from prokaryotic plasmids. CRESS-DNA viruses such as the cruciviruses, have homologues of capsid proteins (CPs) encoded by RNA viruses. The genetic structure of cruciviruses attests to the transfer of capsid genes between disparate groups of viruses. However, the evolutionary history of cruciviruses is still unclear. By collecting and analyzing cruciviral sequence data, we provide a deeper insight into the evolutionary intricacies of cruciviruses. Our results reveal an unexpected diversity of this virus group, with frequent recombination as an important determinant of variability.

## INTRODUCTION

In the last decade, metagenomics has allowed for the study of viruses from a new angle; viruses are not merely agents of disease, but abundant and diverse members of ecosystems (1, 2). Viruses have been shaping the biosphere probably since the origin of life, as they are important drivers of the evolution of the organisms they infect (3–5). However, the origin of viruses is not entirely clear. Viruses, as replicons and mobile elements, are also subject to evolution. Virus variability is driven by various mutation rates, recombination and reassortment of genetic components (6). These attributes, coupled with types of genomes (RNA or DNA, single or double stranded and circular or linear), lead to a large genetic diversity in the ‘viral world’.

Viruses are generally classified based on the nature of their transmitted genetic material (7). Viral genetic information is coded in either RNA or DNA. Moreover, these genomes can be single (positive or negative sense) or double stranded, linear or circular, and can be comprised of a single or multiple molecules of nucleic acid (monopartite or multipartite, respectively). These different groups of viruses have different replication strategies, and they harbor distinct taxa based on their genome arrangement and composition (1). The striking differences between viral groups with disparate genome types suggest polyphyletic virus origins (8).

For example, the highly abundant circular Rep-encoding single-stranded DNA (CRESS-DNA) viruses may have been derived from plasmids on multiple occasions by acquiring capsid genes from RNA viruses (9–11). Eukaryotic CRESS-DNA viruses constitute a diverse and widespread group of viruses with circular genomes -some of them multipartite– that contains the families *Geminiviridae, Circoviridae*, *Nanoviridae*, *Alphasatellitidae*, *Genomoviridae*, *Bacilladnaviridae*, *Smacoviridae* and *Redondoviridae* (ICTV classification for some groups is pending at this time), in addition to vast numbers of unclassified viruses (12, 13). Universal to all CRESS-DNA viruses is the Rep, which is involved in the initiation of the virus’ rolling-circle replication. Rep homologues are also encoded in plasmids (13, 14). Some pathogenic CRESS-DNA viruses are agriculturally important, such as porcine circoviruses, and nanoviruses and geminiviruses that infect a wide range of plant hosts (12). However, many CRESS-DNA viruses have been identified in apparently healthy organisms, and metagenomics have revealed their presence in most environments (12).

In 2012, a metagenomic survey of a hot and acidic lake in the volcanic Cascade Range of the western USA uncovered a new type of circular DNA virus (15). The genome of this virus is a CRESS-DNA virus based on the circularity of its sequence, the presence of a *rep* gene, and a predicted stem-loop structure with a conserved nucleotide sequence (*ori*) that serves as an origin for CRESS-DNA virus rolling-circle replication (RCR; reviewed in (16, 17)). Interestingly, the sequence of CP encoded by this genome resembles those encoded by the RNA viruses in the family *Tombusviridae* (15). It was hypothesized that this virus originated by the acquisition of a capsid gene from an RNA virus through a yet to be demonstrated RNA-DNA recombination event (15, 18). Since the discovery of this putatively “chimeric virus”, 80 circular sequences encoding a Rep and a CP that share homology to tombusvirus CPs have been found in different environments around the globe (19, 20, 29–31, 21–28). This growing group of viruses have been branded “cruciviruses”, as they imply the *crossing* between CRESS-DNA viruses and RNA tombusviruses (27). Cruciviruses have been found associated with forams (20), alveolates hosted by isopods (26), arthropods (19, 22), and in peatland ecosystems (27), but no host for cruciviruses have been elucidated to date.

The circular genome of known cruciviruses is variable in size, ranging from 2.7 to 5.7 kb and often contains ORFs in addition to the Rep and CP, which have been found in either a unisense or an ambisense orientation (20, 27). The function of additional crucivirus ORFs is unclear due to the lack of sequence similarity with any characterized protein. The genome replication of CRESS-DNA viruses is initiated by the Rep protein, that binds to direct repeats present just downstream of the stem of the *ori*-containing stem-loop structure and nicks the ssDNA (32, 33). The exposed 3’OH serves as a primer for cellular enzymes to replicate the viral genome via RCR (33–35). The exact terminating events of CRESS-DNA virus replication are poorly understood for most CRESS-DNA viruses, but Rep is known to be involved in the sealing of newly replicated genomes (33, 35–37).

Rep has a domain in the N-terminus that belongs to the HUH endonuclease superfamily (38). This family of proteins is characterized by a HUH motif (Motif II), in which two histidine residues are separated by a bulky hydrophobic amino acid, and a Tyr-containing motif (Motif III) that catalyzes the nicking of the ssDNA (38–41). CRESS-DNA virus Reps also contain a third conserved motif in the N-terminal portion of the protein (Motif I), likely responsible for dsDNA binding specificity (42). In many CRESS-DNA viruses, the HUH motif has been substituted for a similar motif that lacks the second histidine residue (e.g. circoviruses have replaced HUH with HLQ) (10, 38). The C-terminal portion of eukaryotic CRESS-DNA virus Reps contain a Superfamily 3 helicase domain (S3H) that may be responsible for unwinding dsDNA replicative intermediates (43, 44). This helicase domain is characterized by Walker A and B motifs, Motif C and an Arg finger. Previous studies have identified evidence of recombination in the endonuclease and helicase domains of Rep, which contributes to the potential ambiguity of Rep phylogenies (45). Interestingly, the Rep proteins of different cruciviruses have been shown to be similar to CRESS-DNA viruses in different families, including circoviruses, nanoviruses, and geminiviruses (20, 27). In some cruciviruses, these differences in phylogeny have been observed between the individual domains of a single Rep protein (21, 27). The apparent polyphyly of crucivirus Reps suggests recombination events involving cruciviruses and other CRESS-DNA viruses, even within Reps (20, 21).

All characterized CRESS-DNA viruses package their DNA into small capsids with icosahedral symmetry or their geminate variants, built from multiple copies of the CP encoded in their genome (12). The CP of these CRESS-DNA viruses appears to fold into an eight-strand ß-barrel that conforms to the single jelly-roll (SJR) architecture, which is also commonly found in eukaryotic RNA viruses (46). The CP of cruciviruses has no detectable sequence similarity with the capsid of other CRESS-DNA viruses, and is predicted to adopt the SJR conformation found in the CP of tombusviruses (15, 20, 21). Three domains can be distinguished in tombusviral CPs (47, 48). From the N- to the C-terminus: i) The RNA-interacting or R-domain, a disordered region that faces the interior of the viral particle to interact with the nucleic acid through abundant basic residues (49, 50), ii) the shell or S-domain containing the single jelly-roll fold and the architectural base of the capsid (48) and iii) the protruding or P-domain, that decorates the surface of the virion and is involved in host transmission (51). In tombusviruses, the S-domain of 180 CP subunits interact with each other to assemble around the viral RNA in a T=3 fashion, forming a Ø~35 nm virion (48, 52).

The study of cruciviruses suggests evidence for the transfer of capsid genes between disparate viral groups, which can shed light on virus origins and the phenotypic plasticity of virus capsids. Here, we document the discovery of 461 new crucivirus genomes (CruV) and cruci-like circular genetic elements (CruCGE) identified in metagenomic data obtained from different environments and organisms. This study provides a comprehensive analysis of this greatly expanded dataset and explores the extent of cruciviral diversity -mostly due to Rep heterogeneity– impacted by rampant recombination.

## MATERIALS AND METHODS

### Recovery of viral genomes from assembled viromes

A total of 461 crucivirus-related sequences were identified from 1168 metagenomic surveys (Supp. Tables 1 and 2). 1167 viromes from 57 published datasets and one unpublished virome were obtained from different types of environments: i) aquatic systems (freshwater, seawater, hypersaline ponds, thermal springs and hydrothermal vents), ii) engineered systems (bioreactor, food production), and iii) eukaryote-associated flora (human, insect and other animal feces, human saliva and fluids, cnidarians and plants). New cruciviral sequences were identified in these viromes by screening circular contigs for the presence of CPs from previously known cruciviruses (20) and tombusviruses, using a BLASTx bit-score threshold of 50.

Additionally, sequences CruV-240, CruV-300, CruV-331, CruV-338 and CruV-367 were retrieved from Joint Genome Institute (JGI)’s IMG/VR repository (53), by searching scaffolds with a function set including the protein family pfam00729, corresponding to the S-domain of tombusvirus capsids. The sequences with an RdRP coding region were excluded, and the circularity of the sequences, as well as the presence of an ORF encoding a tombusvirus-like capsid, were confirmed with Geneious 11.0.4 (Biomatters, Ltd).

### Annotation of crucivirus putative genes

The 461 cruciviral sequences were annotated and analyzed in Geneious 11.0.4. Coding sequences (CDSs) were semi-automatically annotated from a custom database (Supp. Table 3) of protein sequences of published cruciviruses and close homologs obtained from GenBank, using Geneious 11.0.4’s annotation function with a 25% nucleotide similarity threshold. Annotated CDS were re-checked with GenBank database using BLASTx to identify sequences similar to previously described cruciviruses and putative relatives. Sequences containing in-frame stop codons were checked for putative splicing sites (54), or translated using a ciliate genetic code only when usage rendered a complete ORF with similarity to other putative crucivirus CDSs. Predicted ORFs longer than 300 bases with no obvious homologs and no overlap with CP or Rep-like ORFs were annotated as “putative ORFs”.

### Putative Stem-loop annotation

Stem-loop structures that could serve as an origin of replication for circular ssDNA viruses were identified and annotated using StemLoop-Finder (Pratt *et* al., unpublished, (55, 56)). The 461 cruciviral sequences were scanned for the presence of conserved nonanucleotide motifs described for other CRESS-DNA viruses (NANTANTAN, NAKWRTTAC, TAWWDHWAN, & TRAKATTRC) (12). The integrated ViennaRNA 2.0 library was used to predict secondary structures of DNA around the detected motif, including the surrounding 15-20 nucleotides on either side (57, 58). Predicted structures with a stem longer than four base pairs and a loop including seven or more bases were subjected to the default scoring system, which increases the score by one point for each deviation from ideal stem lengths of 11 base pairs and loop lengths of 11 nucleotides. A set of annotations for stem-loops and nonanucleotides was created with StemLoop-Finder for those with a score of 15 or below. Putative stem-loops were excluded from annotation when a separate stem-loop was found with the same first base, but attained a greater score, as well as those that appeared to have a nonanucleotide within four bases of its stem-loop structure’s first or last nucleotide.

### Conservation analysis and visualization

#### Pairwise identity matrices

The pairwise identity (PI) between the protein sequence from translated cruciviral genes was calculated with SDTv1.2 (59), with MAFFT alignment option for CPs and S-domains, and MUSCLE alignment options for Reps.

#### Sequence conservation annotation

CP sequence conservation represented in Fig. 2A was generated with Jalview v2.11.0 (60), and reflects the conservation of the physicochemical properties for each column of the alignment (61).

#### Sequence logos

Sequence logos showing frequency of bases in nonanucleotides at the origin of replication or residue in conserved Rep motifs were made using the weblogo server (http://weblogo.threeplusone.com/; (62)).

#### Structural representation of capsid conservation

The 3D structure of CP was modeled with Phyre^2^ (63). The generated graphic was colored by sequence conservation with Chimera v.1.13 (64), from the alignment of the 47 capsid sequences from each of the CP-based clusters (Fig. 3B).

### Phylogenetic analyses

#### Multiple sequence alignments

CP sequences were aligned using MAFFT (65) in Geneious 11.0.4, with a G-INS-i algorithm and BLOSUM 45 as exchange matrix, with an open gap penalty of 1.53 and an offset value of 0.123, and manually curated. Rep protein sequences were aligned using PSI-Coffee [http://tcoffee.crg.cat/; (66)]. Rep alignments were manually inspected and corrected in Geneious 11.0.4, and trimmed using TrimAI v1.3 with a *strict plus* setting (67). To produce individual alignments of the endonuclease and helicase domains the full length trimmed alignments were split at the Walker A motif (45).

#### Phylogenetic trees

Phylogenetic trees containing the entire dataset of cruciviral sequences were built on Geneious using the FastTree plugin (68). For the analysis of sequence subsets, trees were inferred with PhyML 3.0 web server [http://www.atgc-montpellier.fr/phyml/; (69)], using an aLRT SH-like support (70). The substitution model for each analysis was automatically selected by the program.

#### Intergene and interdomain comparison

Tanglegrams were made using Dendroscope v3.5.10 (71) to compare the phylogenies between different genes or domains within the same set of crucivirus genomes.

#### Sequence similarity networks

A total of 540 CP amino acid sequences, and 600 Rep amino acid sequences were uploaded to EFI–EST web server for the calculation of PIs [https://efi.igb.illinois.edu/efi-est/; (72)]. A specific alignment score cutoff was established for each dataset analyzed, and xgmml files generated by EFI-EST were visualized and edited in Cytoscape v3.7.2 (73).

### Accession numbers

Provided in Supp. Table 1.

## RESULTS & DISCUSSION

### Expansion of the crucivirus group

To broaden our understanding of the diversity and relationships of cruciviruses, 461 uncharacterized circular DNA sequences containing predicted CDSs with sequence similarity to the CP of tombusviruses were compiled from metagenomic sequencing data. The data came from published and unpublished metagenomic studies, carried out in a wide variety of environments, from permafrost to temperate lakes, and on various organisms from red algae to invertebrates (Metagenomes and their metadata are provided in Supp. Table 2). The selected genomes are assumed to be complete and circular based on the terminal redundancy identified in *de novo* assembled genomes.

The cruciviral sequences were named sequentially, beginning with the smallest genome, which was named CruV-81 to account for the 80 crucivirus genomes reported in prior literature (15, 19, 28–31, 20–27). The average GC content of the newly described cruciviral sequences is 42.9 ± 4.9 % (Fig. 1B) with genome lengths spanning from 2,474 to 7,947 bases (Fig. 1A), some exceeding the size of described bacilladnaviruses (≤6,000 nt (74)), the largest CRESS-DNA viruses known (12).

**Figure 1.**
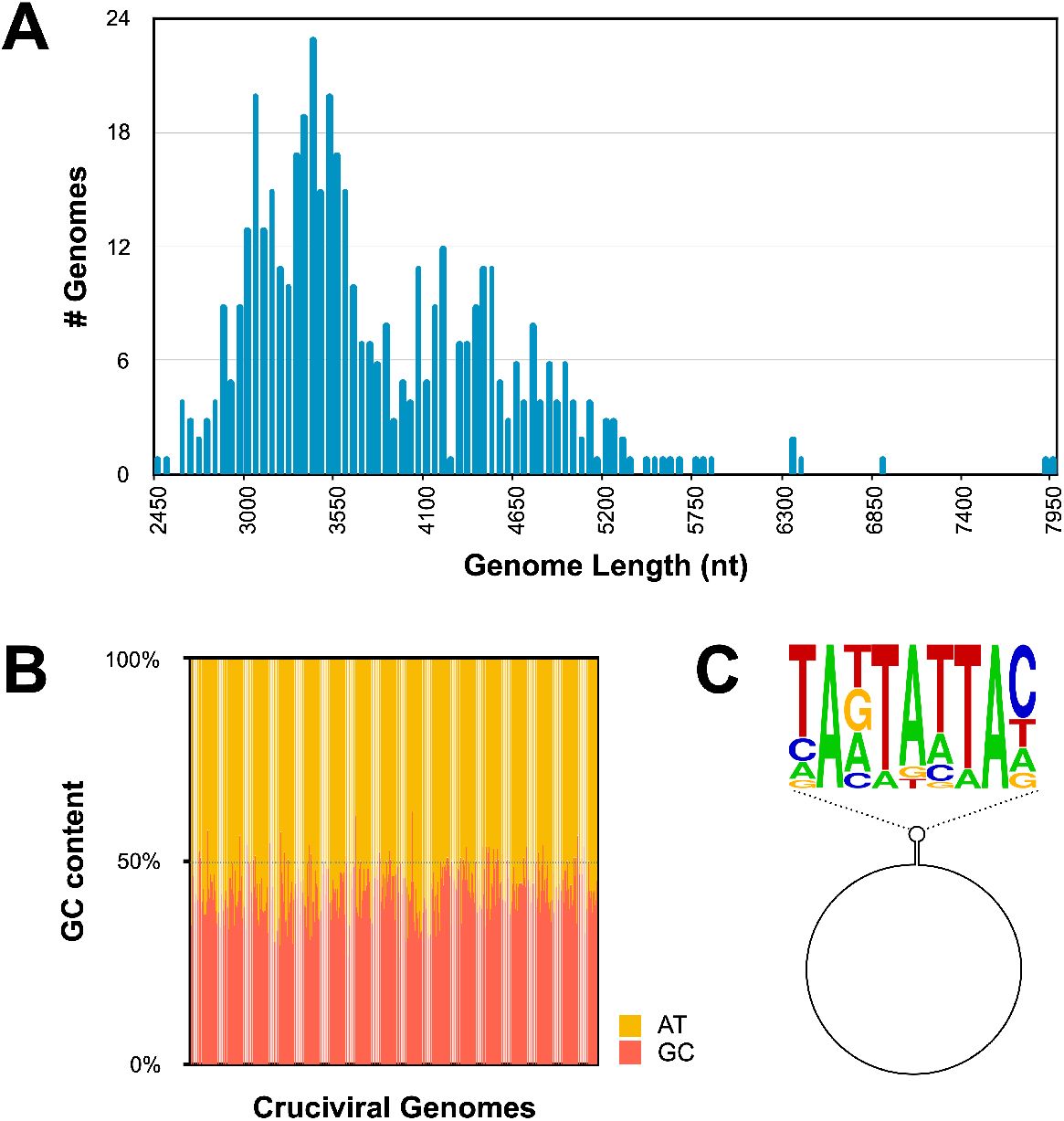
Genome properties of 461 new cruciviral circular sequences. **(A)** Histogram of cruciviral genome lengths categorized in 50 nt bins. **(B)** Percentage of G + C content versus A + T in each of the sequences described in this study **(C)** Relative abundance of nucleotides in the conserved nonanucleotide sequence of the 211 stem-loops and putative origins of replication represented predicted with StemLoop-Finder (Pratt *et* al., unpublished) in Sequence Logo format.

Of the 461 sequences that contain a CP ORF, 451 have putative coding regions with sequence similarity to Rep of CRESS-DNA viruses (10). The CP and Rep ORFs are encoded in a unisense orientation in 40% of the genomes and an ambisense orientation in 58% of the genomes. The remaining ~2% correspond to ten CruCGEs with no clear Rep CDS. Five of these CruCGEs contain a predicted origin of rolling-circle replication (RCR) (Supp. Table 1), indicating that they are circular genomes that undergo RCR characteristic of other CRESS-DNA virus genomes (16, 17).

One possible reason for the lack of a Rep ORF in certain sequences is that some of these may be sub-genomic molecules or possible components of multipartite viruses (75). Some CRESS-DNA viruses, such as geminiviruses and nanoviruses, have multipartite genomes (76). Moreover, some ssRNA tombunodaviruses; including *Plasmopara halstedii* virus A and *Schlerophthora macrospora* virus A -viruses that contain the most similar capsid sequences to cruciviral capsids (15, 27)– also have multipartite genomes (77). Unfortunately, no reliable method yet exists to match different sequences belonging to the same multisegmented virus in metagenomes, making identification of multipartite or segmented viruses from metagenomic data challenging (76).

Stem-loop structures with conserved nonanucleotide motifs as putative origins of replication were predicted and annotated in 277 cruciviral sequences with StemLoop-Finder (Pratt *et* al., unpublished). In some cases, more than one nonanucleotide motif with similar scores were found for a single genome, resulting in more than one stem-loop annotation. Of the annotated genomes, 223 contain a stem-loop with a nonanucleotide with a NANTANTAN pattern, with the most common sequence being the canonical circovirus motif TAGTATTAC, found in 64 of the genomes (Supp. Table 1; (78)). The majority of the 54 sequences that do not correspond to NANTANTAN contain a TAWWDHWAN nonanucleotide motif, typical of genomoviruses (79). The frequency of bases at each position in the nonanucleotide sequence is given in Fig. 1C, and reflects similarity to motifs found in other CRESS-DNA viruses (10).

### Crucivirus capsid protein (CP)

The CP of cruciviruses is predicted to have a single jelly-roll (SJR) architecture, based on its homology to tombusvirus CPs for which 3D structures have been determined [Fig. 2A; (80–82)]. The SJR conformation is found in CPs of both RNA and DNA viruses (46). The SJR CP of tombusviruses and cruciviruses contains three distinct domains: the RNA-binding or R-domain, the shell or S-domain, and the protruding or P-domain (Fig. 2A). All 461 crucivirus CPs analyzed in this study contain a complete S-domain. This domain contains a distinct jelly-roll fold and interacts with the S-domain of other capsid subunits in the virion of related tombusviruses (48). The S-domain has greater sequence conservation than the remaining regions of the CP (Fig. 2A), likely due to its functional importance in capsid structure. In tombusviruses, the S-domain contains a calcium binding motif (DxDxxD), which was not identified in previously described cruciviruses (83). However, we detected this Ca-binding motif in 68 CPs of the newly identified cruciviral sequences. These crucivirus sequences form a distinct cluster, shown in red in Fig. 3B. The S-domain is flanked on the N-terminus by the R-domain, which in cruciviruses appears variable in size (up to 320 amino acids long), and appears to be truncated in some of the CP sequences (e.g. CruV-386 and CruV-493). The R-domain is characterized by an abundance of basic residues at the N-terminus, followed by a Gly-rich tract (Fig. 2A). The P-domain, on the C-terminal end of the CP sequence, is generally the largest domain, with the exception of CruV-385, where it appears to be truncated. The conservation of CP suggests a similar structure for all cruciviruses. However, those cruciviruses with larger genomes may assemble their capsids in a different arrangement to accommodate their genome. While the capsids of tombusviruses have been shown to adopt a T=1 icosahedral conformation, rather than the usual T=3, when the R-domain is partially or totally removed (82), we have not seen a correlation between the length of CP domains and genome size in our dataset that could be indicative of alternative capsid arrangements. Furthermore, no packaging dynamics relating genome size and virion T-number arrangement have been determined in CRESS-DNA viruses, although sub-genomic elements of geminiviruses can be packaged in non-geminate capsids (84, 85).

**Figure 2.**
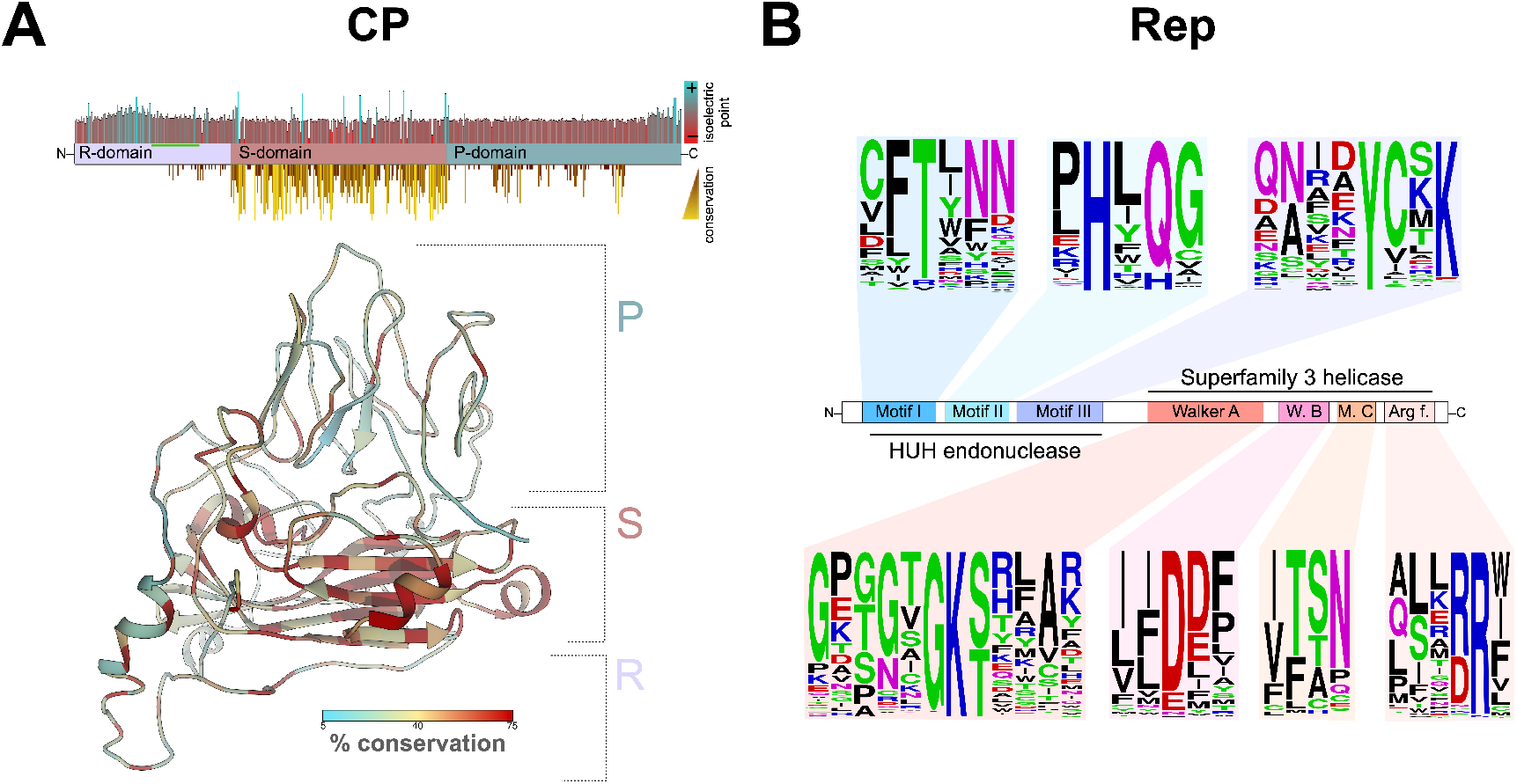
Protein conservation in cruciviruses. **(A) Top:** distribution of domains, isoelectric point and conservation in a consensus capsid protein (CP). 461 CP protein sequences were aligned in Geneious 11.0.4 with MAFFT (G-INS-i, BLOSUM 45, open gap penalty 1.53, offset 0.123) and trimmed manually. The conservation of the physico-chemical properties at each position was obtained with Jalview v2.11.0, and the isoelectric point was estimated in Geneious 11.0.4. The region of CP rich in glycine is highlighted with a green bar. **Bottom:** Structure of a cruciviral CP (CruV-359) as predicted by Phyre2 showing sequence conservation based on an alignment of the 47 CP protein sequences from the CP-based clusters. **(B)** Conserved motifs found in cruciviral Reps after aligning all the extracted Rep protein sequences using PSI-Coffee. Sequence logos were generated at http://weblogo.threeplusone.com to indicate the frequency of residues at each position.

**Figure 3.**
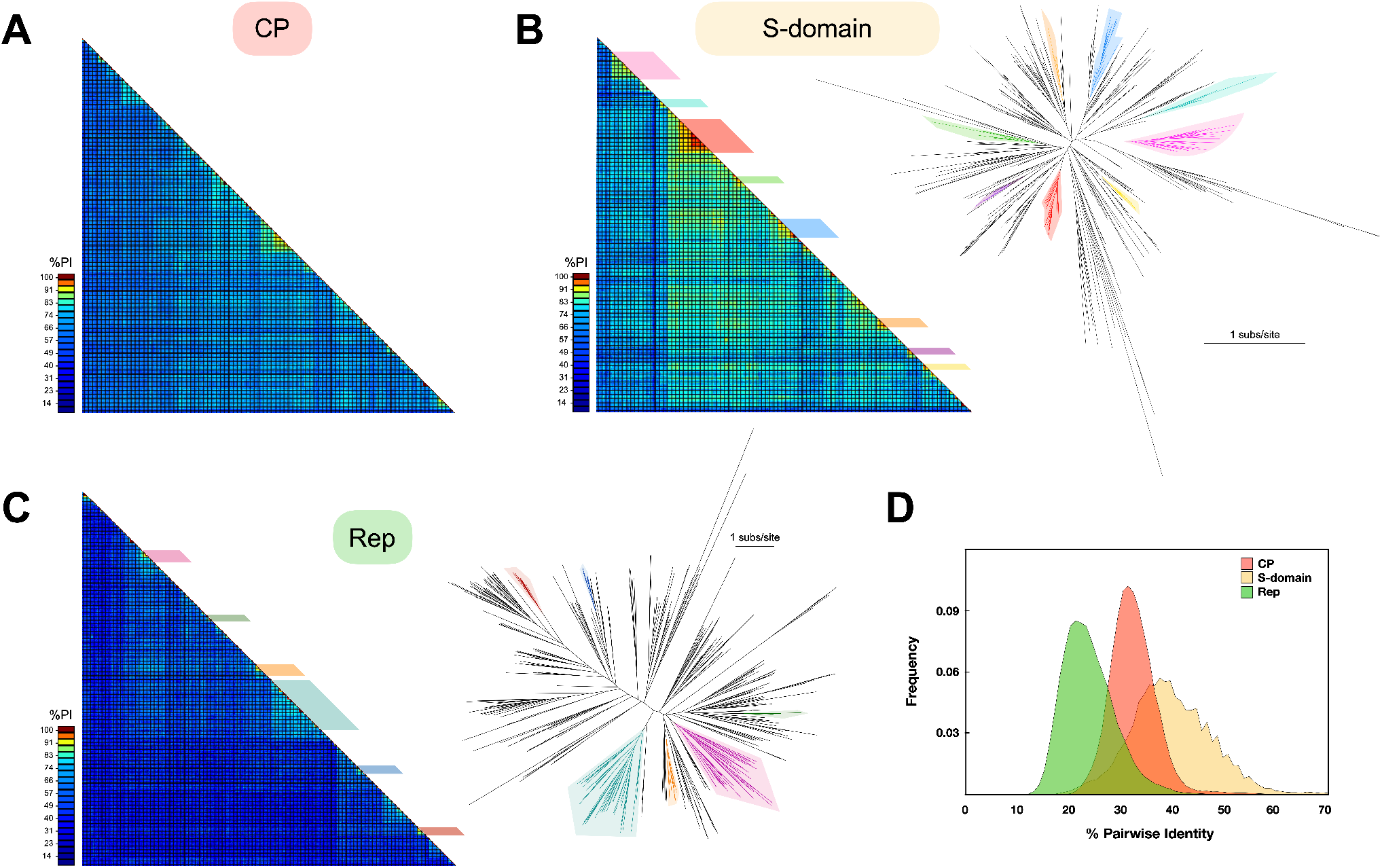
Diversity of cruciviral proteins. **(A) CP diversity.** Pairwise amino acid identity (PI) between the CPs predicted for 461 cruciviral sequences. The alignment and analysis were carried out with SDT, using the integrated MAFFT algorithm. **(B) S-domain diversity. Left:** PI matrix between the capsid protein (CP) predicted S-domain of the 461 sequences described in this study. The alignment and analysis were carried out with SDT, using the integrated MAFFT algorithm. The colored boxes indicate the different clusters of sequences used to create the CP-based clusters sequence subset. **Right:** Unrooted phylogenetic tree obtained with FastTree from a manually curated MAFFT alignment of the translated sequences of the S-domain (G-INS-i, BLOSUM 45, open gap penalty 1.53, offset 0.123). The colored branches represent the different clusters observed in the matrix. Scale bar indicates substitutions per site. **(C) Rep diversity. Left:** Pairwise identity (PI) matrix between all Reps found in cruciviral genomes in this study. The alignment and analysis were carried out with SDT, using the integrated MUSCLE algorithm. **Right:** Unrooted phylogenetic tree obtained with FastTree from an PSI-Coffee alignment of the translated sequences of Rep trimmed with TrimAl v1.3. The colored branches represent the different clusters that contain *Rep-based clusters* sequence subset. Scale bar indicates substitutions per site. **(D) PI frequency distribution.** The frequency of PI values for each of the putative proteins or domains analyzed is shown.

Interestingly, CruV-420 contains not one tombusvirus-related CP, but two. A recent compilation of CRESS-DNA viruses from animal metagenomes also contains four genomes with two different CPs in their capsid (31). Whether these viruses use two different CPs in their capsid (as some RNA viruses do), or whether these are intermediates in the exchange of CP genes, as predicted from the gene capture mechanism proposed by Stedman (2013) (18), is unclear. If the latter is true, CP gene acquisition by CRESS-DNA viruses may be much more common than previously thought.

### Crucivirus Rep

The Reps of CRESS-DNA viruses typically contain an endonuclease domain characterized by conserved motifs I, II and III, and a helicase domain with Walker A and B motifs, motif C, and an Arg-finger [Fig. 2B; (12)]. The majority (85.9%) of the crucivirus genomes described in this dataset contain all of the expected Rep motifs (Supp. Table 4). However, five genomes (CruCGE-110, CruCGE-296, CruCGE-436, CruCGE-471 and CruCGE-533) with overall sequence homology to other Reps (35.8, 32.7, 49.7, 60.2 and 57.2 % PI with other putative Reps in the databases, respectively), lack any detectable conserved motifs within their sequence. Thus, these sequences are considered CP-encoding cruci-like circular genetic elements (CruCGEs).

The endonuclease catalytic domain of Rep (motif II), including HUH, was identified in 441 of the genomes of which 95.2% had an alternative HUH, with the most common arrangement being HUQ (70.0%), also found in circoviruses and nanoviruses [(10, 25, 39); Fig. 2B]. 26.2% of the crucivirus motif II deviate from the HUH motif by additionally replacing the second hydrophobic residue (U) with a polar amino acid (Fig. 2B; supp. Table 4), with 53 of Reps with the sequence HYQ (12.0%), also found in smacoviruses (10, 23, 45).

We identified thirteen putative Reps in these crucivirus genomes that lack all four motifs typically found in S3H helicases (e.g. CruV-166, CruV-202, CruV-499; Supp. Table 4). Recent work has shown that the deletion of individual conserved motifs in the helicase domain of the Rep protein of beak and feather disease virus does not abolish ATPase and GTPase activity (86). The absence of all four motifs may prevent these putative Reps from performing helicase and ATPase activity using previously characterized mechanisms. However, it is possible that crucivirus Reps that lack these motifs are still capable of ATP hydrolysis and associated helicase activity. Alternatively, these activities may be provided by host factors (87), or by a viral replication-enhancer protein – as is the case with the AC3 protein of begomoviruses (88).

We identified 36 crucivirus genomes whose putative *rep* genes contain in-frame stop codons or the HUH and SF3 helicase are in different frames, suggesting that their transcripts may require intron splicing prior to translation. Acceptor and donor splicing sites identical to those found in maize streak virus (54) were found in all these sequences, and the putatively spliced Reps annotated accordingly. In five of the 36 spliced Reps, we were unable to detect any of the four conserved motifs associated with helicase/ATPase activity, which are encoded in the predicted second exon in most cases. CruV-513 and CruV-518 also contain predicted splicing sites in their CP gene.

No GRS motifs -which have been identified as necessary for geminivirus replication (89), and have also been found in genomoviruses (90)– were detected in Reps in our dataset. We were unable to detect any conserved Rep motifs present in cruciviruses that are absent in other CRESS-DNA viruses. Given the conservation of Rep motifs in these newly-described cruciviruses, we expect most to be active in RCR.

### Crucivirus CPs share higher genetic identity than their Rep proteins

To assess the diversity in the proteins of cruciviruses, the percentage pairwise identity (% PI) between the protein sequences was calculated for CP and Rep using SDTv1.2 (Fig. 3). The average % PI for CP was found to be 33.1± 4.9 % PI (Figs. 3A and 3D), likely due to the high levels of conservation found in the S-domain (40.5 ± 8.4 % PI; Figs. 3B and 3D), while the average % PI for Rep is quite low at 24.7% (± 5.6 SD; Figs. 3C and 3D). The high variation of the Rep protein sequence relative to CP in cruciviruses correlates with a previous observation on a smaller dataset (20).

To compare cruciviruses to other viral groups with homologous proteins, sequence similarity networks were built for CP and Rep (Fig. 4). For the CP, related protein sequences from tombusviruses and unclassified RNA viruses were included. The virus sequences were connected when the similarity between their protein sequence had an e-value < 1e–20, sufficient to connect all cruciviruses and tombusviruses, with the exception of CruV-523 (Fig. 4A). However, using BLASTp, CruV-523 showed similarity to other RNA viruses with an e-value < 1e–9, which were not included in the analysis. The CP sequence similarity network analysis demonstrates the apparent homology of the CPs in our dataset with the CP of RNA viruses; specifically to unclassified RNA viruses that have RdRPs similar to either tombusviruses -also described as tombus-like viruses (77, 91, 92)- or to nodaviruses. The latter RNA viruses are proposed to belong to a chimeric group of viruses named tombunodaviruses (93).

**Figure 4.**
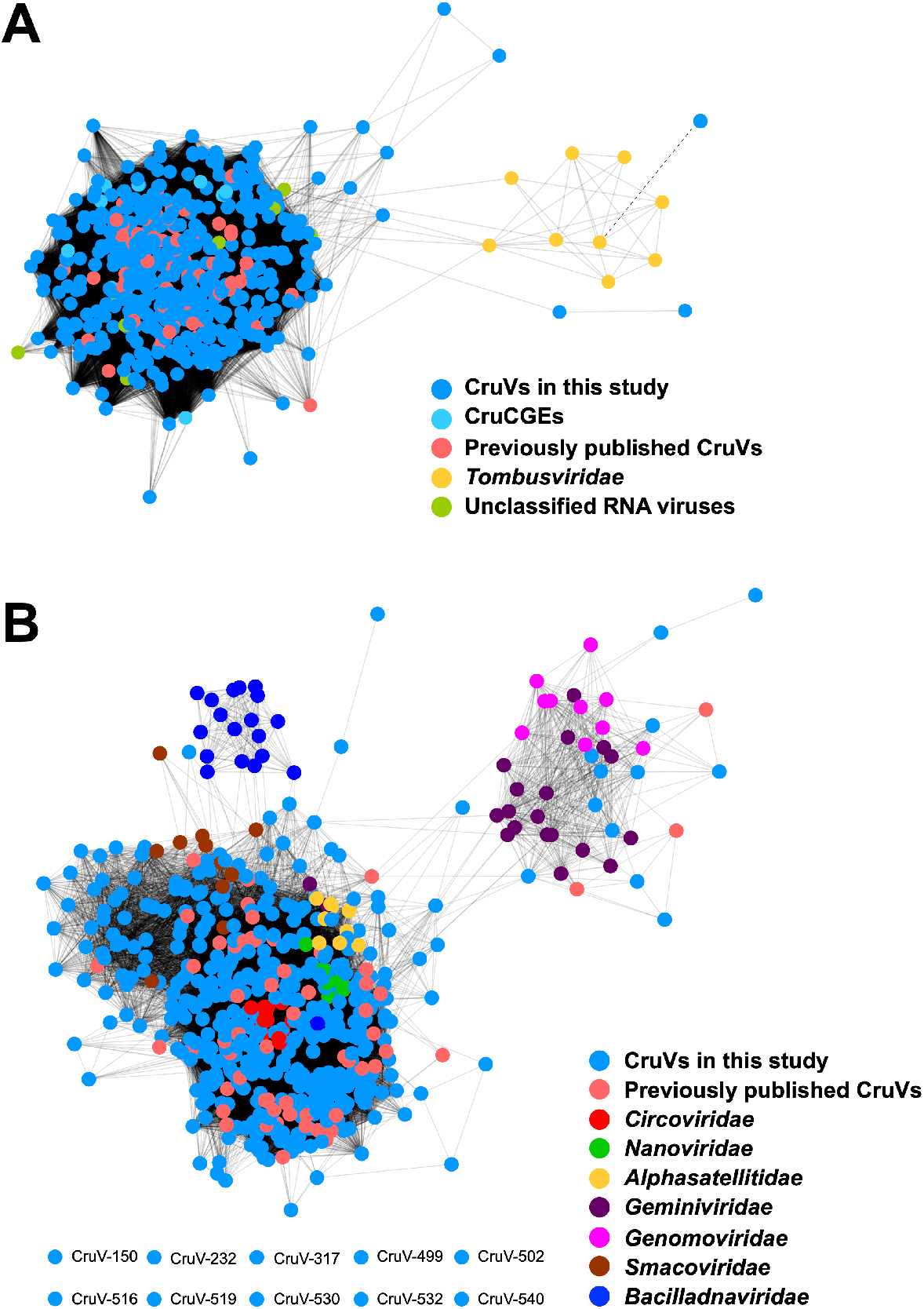
Similarity networks of cruciviral proteins with related viruses. **(A)** Capsid proteins (CP) represented by colored dots are connected with a solid line when the similarity between them is greater than e-value=1e^−20^. The dashed line represents an e-value = 6e^-7^ between the nodes corresponding to the CP of CruV-523 and turnip crinkle virus, as given by BLASTp. **(B)** Replication-associated protein (Rep) translations, represented by colored dots, are connected with a solid line when the similarity between them greater than e-value=1e^−10^. The eight nodes at the bottom left did not connect to any other node. All networks were carried out with pairwise identities calculated in the EFI–EST web server and visualized in Cytoscape v3.7.2.

For sequence similarity network analysis of Rep, sequences from CRESS-DNA viruses belonging to the families *Circoviridae, Nanoviridae, Alphasatellitidae, Geminiviridae, Genomoviridae, Smacoviridae* and *Bacilladnaviridae* were used (Fig. 4B). Due to the heterogeneity of Rep (Fig. 3C), the score cutoff for the network was relaxed to an e-value < 1e–10; nonetheless, ten divergent sequences lacked sufficient similarity to form connections within the network. While the Rep of the different viral families clustered in specific regions of the network, the similarity of cruciviral Reps spans the diversity of all CRESS-DNA viruses, and blurs the borders between them. Though there are cruciviruses that appear to be closely related to geminiviruses and genomoviruses, these connections are less common than with other classified CRESS-DNA families (Fig. 4B). While still highly divergent from each other, the conserved motifs in the Rep still share the most sequence similarity with CRESS-DNA viruses (Fig. 2B).

The broad sequence space distribution of cruciviral Rep sequences has been proposed to reflect multiple Rep acquisition events through recombination with viruses from different CRESS-DNA viral families (20). However, the apparent larger diversity of cruciviral Reps relative to classified CRESS-DNA viruses can be due to the method of study, as most classified CRESS-DNA viruses have been discovered from infected organisms and are grouped mainly based on Rep similarity (1). By contrast, here crucivirus sequences are selected according to the presence of a tombusvirus-like CP. Moreover, the Rep of cruciviruses could be subject to higher substitution rates than CP (26). It is possible that sequence divergence in CP is more limited than in the Rep due to structural constraints.

### Horizontal gene transfer among cruciviruses

To gain insight into the evolutionary history of cruciviruses, we carried out phylogenetic analyses of their CPs and Reps. Due to the high sequence diversity in the dataset, two smaller subsets of sequences were analyzed:

i. *CP-based clusters*. Clusters with more than six non-identical CP sequences whose S-domains share a % PI greater than 70% were identified from Fig. 3B. This resulted in the identification of seven clusters, and a more divergent, yet clearly distinct, cluster was included (pink in Fig. 3B). A total of 47 genomes from the eight different clusters were selected for sequence comparison. The protein sequences of CP and Rep were extracted, aligned, and their phylogenies inferred and analyzed using tanglegrams (Fig. 5A). The CP phylogeny shows that the eight CP-based clusters form separate clades (Fig. 5A). On the other hand, the phylogeny of Rep shows a different pattern of relatedness between those genomes (Fig. 5A). This suggests different evolutionary histories for the CP and Rep proteins, which could be due to recombination events between cruciviruses, as previously proposed with smaller datasets (20, 21).
ii. *Rep-based clusters*. To account for the possible bias introduced by selecting genomes from CP cluster groups and to increase the resolution in the phylogeny of the Rep sequences, clusters with more than six Rep sequences sharing PI > 45 and < 98% were identified. A total of 53 genomes from six clusters (Fig. 3C) were selected, and their CP and Rep protein sequences analyzed. The phylogeny of Reps shows distinct clades between the sequences from different clusters (Fig. 5B). When the phylogeny of Rep was compared to that of their corresponding CPs, we observed the presence of groups of cruciviruses that clade with each other in both the Rep and the CP. Discrepancies in topology between Rep and CP were observed as well, particularly in the CP clade marked with an asterisk in Fig. 5B. This clade corresponds to the highly-homogeneous red CP-based cluster shown in Fig. 3B, and suggests that gene transfer is more common in cruciviruses with a more similar CP, likely infecting the same type of organism. On the other hand, the presence of cruciviral groups with no trace of genetic exchange may indicate that lineages within the cruciviral group may have undergone speciation in the course of evolution.

**Figure 5.**
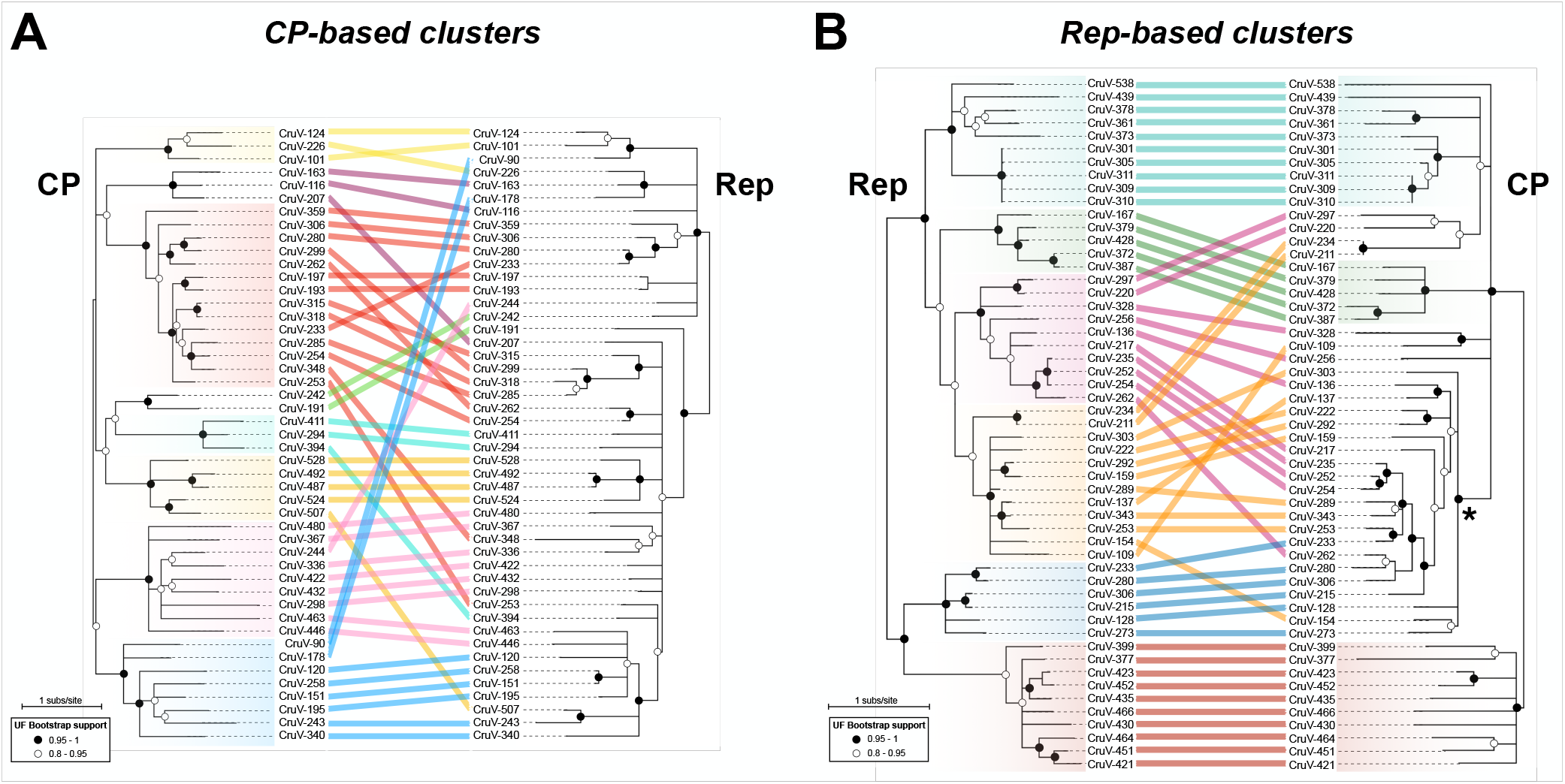
Comparison of phylogenies of CP and Rep proteins of representative cruciviruses. **(A)** Tanglegram calculated with Dendroscope v3.5.10 from phylogenetic trees generated with PhyML from Cp (PhyML automatic model selection LG+G+I+F) and Rep (PhyML automatic model selection RtREV+G+I) alignments. The tips corresponding to the same viral genome are linked by lines that are color-coded according to the clusters obtained from Fig. 3A (CP-based clusters). **(B)** Tanglegram calculated with Dendroscope v3.5.10 from phylogenetic trees generated with PhyML from Cp (PhyML automatic model selection LG+G+I+F) and Rep (PhyML automatic model selection RtREV+G+I) alignments. The tips corresponding to the same viral sequence are linked by lines that are color-coded according to the clusters obtained from Fig. 3B (Rep-based clusters). The clade marked with an asterisk is formed by members of the red cluster of subset A. Branch support is given according to aLRT SH-like (Anisimova & Gascuel, 2006). All nodes with an aLRT SH-like branch support inferior to 0.8 were collapsed with Dendroscope prior to constructing the tanglegram.

To investigate possible exchanges of individual Rep domains among cruciviruses, the Rep alignments of the analyses of the CP-based and Rep-based clusters were split at the beginning of the Walker A motif to separate endonuclease and helicase domains. From the analysis of the CP-based clusters, we observed incongruence in the phylogenies between endonuclease and helicase domains (Fig. 6A), suggesting recombination within crucivirus Reps, as has been previously hypothesized with a much smaller dataset (21). This incongruency is not observed in the analyzed Rep-based clusters (Fig. 6B). This is likely due to the higher similarity between Reps in this subset of sequences, biased by the clustering based on Rep. We do observe different topologies between the trees, which may be a consequence of different evolutionary constraints to which the endonuclease and helicase domains are subject. The detection of CP/Rep exchange and not of individual Rep domains in Rep-based clusters suggests that the rate of intergenic recombination is higher than intragenic recombination in cruciviruses.

**Figure 6.**
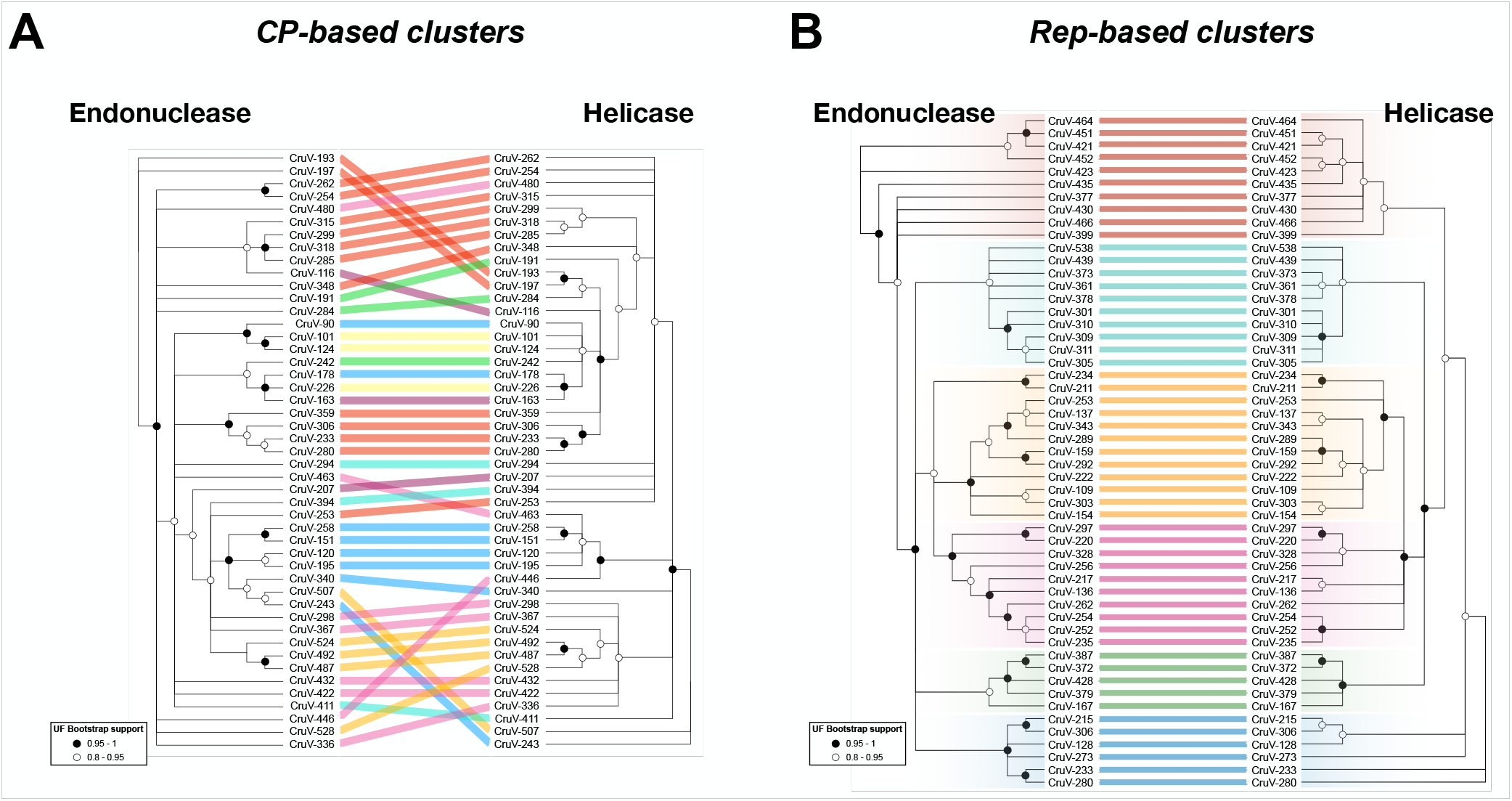
Comparison of phylogenies between the endonuclease and helicase domains of Reps from representative cruciviruses. **(A)** Tanglegram calculated with Dendroscope v3.5.10 from phylogenetic trees generated with PhyML from separate alignments of Rep endonuclease and helicase domains. The tips corresponding to the same viral genome are linked by lines that are color-coded according to the clusters obtained from Fig. 3A (CP-based clusters). **(B)** Same as A but with sequences from the clusters obtained from Fig. 3B (Rep-based clusters). All nodes with an aLRT SH-like branch support inferior to 0.8 were collapsed with Dendroscope v3.5.10 prior to constructing the tanglegram.

### Members of the SAR supergroup are potential crucivirus hosts

While no crucivirus host has been identified to date, the architecture of the Rep protein found in most cruciviruses, as well as the presence of introns in some of the genomes, suggests a eukaryotic host. The fusion of an endonuclease domain to a S3H helicase domain is observed in other CRESS-DNA viruses which are known to infect eukaryotes (38). This is distinct from Reps found in prokaryote-infecting CRESS-DNA viruses -which lack a fused S3H helicase domains (94)– and other related HUH endonucleases involved in plasmid RCR and HUH transposases (38). Additionally, the CP of cruciviruses, a suggested determinant of tropism (95, 96), is homologous to the capsid of RNA viruses known to infect eukaryotes. The RNA viruses with a known host with capsids most similar to cruciviral capsids (tombunodaviruses) infect oomycetes, a group of filamentous eukaryotic stramenopiles (77).

Cruciviruses have been found as contaminants of spin columns made of diatomaceous silica (21), in aquatic metagenomes enriched with unicellular algae (20), in the metagenome of *Astrammina rara* -a foraminiferan protist part of the rhizaria- (20), and associated with epibionts of isopods, mainly comprised of apicomplexans and ciliates, both belonging to the alveolates (26). These pieces of evidence point toward the stramenopiles/alveolates/rhizaria (SAR) supergroup as a candidate taxon to contain potential crucivirus hosts (97). No host prediction can be articulated from our sequence data. However, at least five of the crucivirus genomes only render complete translated CP and Rep sequences when using a relaxed genetic code. Such alternative genetic codes have been detected in ciliates, in which the hypothetical termination codons UAA and UAG encode for a glutamine (98). The usage of an alternative genetic code seems evident in CruV-502 -found in the metagenome from seawater collected above diseased coral colonies (99)– that uses a UAA codon for a glutamine of the S-domain conserved in 33.5% of the sequences. While the data accumulated suggest unicellular eukaryotes and SAR members as crucivirus-associated organisms, the host of cruciviruses remains elusive, and further investigations are necessary.

#### Classification of cruciviruses

Cruciviruses have circular genomes that encode a Rep protein probably involved in RCR. The single-stranded nature of packaged crucivirus genomes has not been demonstrated experimentally; however, the overall genomic structure and sequence similarity underpins the placement of cruciviruses within the CRESS-DNA viruses.

The classification of the CRESS-DNA viruses is primarily based upon the phylogeny of the Rep proteins, although commonalities in CP and genome organization are also considered (13). This taxonomic criteria is challenging in cruciviruses, whose Rep proteins are highly diverse and apparently paralogous. Whether the use of proteins involved in replication for virus classification should be preferred over structural proteins has been previously questioned (100).

The capsid of cruciviruses, as well as the capsid of other CRESS-DNA virus families like circoviruses, geminiviruses and bacilladnaviruses, possess the single-jelly roll architecture (46). However, there is no obvious sequence similarity between the CP of cruciviruses and that of classified CRESS-DNA viruses. The crucivirus CP – homologous to the capsid of tombusviruses– is an orthologous trait within the CRESS-DNA viruses. Hence, CP constitutes a synapomorphic character that demarcate this group of viruses from the rest of the CRESS-DNA viral families. Thus, the classification of cruciviruses is challenging.

## CONCLUDING REMARKS

Cruciviruses are a growing group of CRESS-DNA viruses that encode CPs that are homologous to those encoded by tombusviruses. Over 500 crucivirus genomes have been recovered from various environments across the globe. These genomes vary in size, sequence and genome organization. While crucivirus CPs are relatively homogeneous, the Reps are relatively diverse amongst the cruciviruses, spanning the diversity of all classified CRESS-DNA viruses. It has been hypothesized that cruciviruses emerged from the recombination between a CRESS-DNA virus and a tombus-like RNA virus (15, 18). Furthermore, cruciviruses seem to have recombined with each other to exchange functional modules between them, and probably with other viral groups, which blurs their evolutionary history. Cruciviruses show evidence of genetic transfer, not just between viruses with similar genomic properties, but also between disparate groups of viruses such as CRESS-DNA and RNA viruses.

## ABBREVIATIONS

CruV: Crucivirus
CRESS: Circular Rep-Encoding Single Stranded
CP: Capsid
RdRP: RNA-dependent RNA Polymerase
ORF: Open Reading Frame
SJR: Single Jelly Roll
PI: Pairwise Identity
MDA: Multiple Displacement Amplification
S3H: Superfamily 3 helicase
RCR: Rolling Circle Replication
CDS: Coding Sequence
CruCGE: Cruci-like Circular Genetic Element

## ACKNOWLEDGEMENTS

This work was supported by the NASA Exobiology Program, grant 80NSSC17K0301 (I. dl H., G. K., E. T., A. P. and K. S.), the NIH BUILD EXITO Program (A. M.), BUILD EXITO was supported by grants from the National Institutes of Health: UL1GM118964; RL5GM118963; TL4GM118965, and the Portland State University Ronald E. McNair Scholars Program (E. T.), supported by grants from the U.S. Department of Education and Portland State University. The Antarctic field work was supported by the US National Science Foundation (NSF) under grant ANT-0944411, with logistics supplied by the US Antarctic Program. The fresh water work in New Zealand was supported by a grant (UC-E6007) from the American New Zealand Association (USA) awarded to P. Z-R., C. G., J. S. H. and A. V. The green lipped mussel work was supported by a grant from the Brian Mason Scientific & Technical Trust of New Zealand awarded to S.G. and A.V. EU-s Horizon 2020 Framework Program for Research and Innovation (‘Virus-X’, project no. 685778) supported F. E.

