## Supplementary material for "Unveiling Crucivirus Diversity by Mining Metagenomic Data": Supp. Table 1

Supplementary Table 1: Properties of cruviruses (CruV) and cruci-like circular genetic elements (CruCGE)

| Name | Length | %GC | Genetic Code | Genome organization | ori (nona) | Sequences Subset | Location | Publication | Source | Sequencing | Assembly software | Notes | Accession number* |
|---|---|---|---|---|---|---|---|---|---|---|---|---|---|
| CruV-081 | 2472 | 34.1% | Standard | Ambisense | TAATATTAC | | River (NZ) | | vDNA, MDA | Illumina HiSeq | metaSPAdes | | MT263540 |
| CruCGE-082 | 2535 | 50.1% | Standard | no rep | CATTATTAC | | unnamed Arctic pond (78°02.935′N; 13°41.973′E) | Aguirre de Cárcer et al., 2015 | vDNA, MDA | Illumina HiSeq | IDBA-UD | | PRJEB5265 |
| CruV-083 | 2612 | 46.3% | Standard | Ambisense | #N/A | | unnamed Arctic pond (78°02.935′N; 13°41.973′E) | Aguirre de Cárcer et al., 2015 | vDNA, MDA | Illumina HiSeq | IDBA-UD | Spliced Rep | PRJEB5265 |
| CruV-084 | 2612 | 46.3% | Standard | Ambisense | #N/A | | Lake Tunsjøen (78°03.375′N; 13°40.313′E) | Aguirre de Cárcer et al., 2015 | vDNA, MDA | Illumina HiSeq | IDBA-UD | Spliced Rep | PRJEB5265 |
| CruCGE-085 | 2618 | 39.6% | Standard | no rep | #N/A | | Chirominidae (NZ) | | vDNA, MDA | Illumina HiSeq | metaSPAdes | | MT263538 |
| CruV-086 | 2621 | 35.8% | Standard | Unisense | #N/A | | Lake Tunsjøen (78°03.375′N; 13°40.313′E) | Aguirre de Cárcer et al., 2015 | vDNA, MDA | Illumina HiSeq | IDBA-UD | | PRJEB5265 |
| CruV-087 | 2667 | 42.2% | Standard | Ambisense | #N/A | | Lake Nordammen (78°38.279′N; 16°44.025′E) | Aguirre de Cárcer et al., 2015 | vDNA, MDA | Illumina HiSeq | IDBA-UD | | PRJEB5265 |
| CruV-088 | 2671 | 31.1% | Standard | Ambisense | TAGTATTAC | | Green lipped muscles (NZ) | | vDNA, MDA | Illumina HiSeq | metaSPAdes | | MT263541 |
| CruV-089 | 2690 | 40.4% | Standard | Unisense | #N/A | | Lake Tunsjøen (78°03.375′N; 13°40.313′E) | Aguirre de Cárcer et al., 2015 | vDNA, MDA | Illumina HiSeq | IDBA-UD | | PRJEB5265 |
| CruV-090 | 2702 | 52.5% | Standard | Ambisense | #N/A | Cp-based clusters | Lake Tunsjøen (78°03.375′N; 13°40.313′E) | Aguirre de Cárcer et al., 2015 | vDNA, MDA | Illumina HiSeq | IDBA-UD | | PRJEB5265 |
| CruV-091 | 2702 | 52.6% | Standard | Ambisense | #N/A | | unnamed Arctic pond (78°02.935′N; 13°41.973′E) | Aguirre de Cárcer et al., 2015 | vDNA, MDA | Illumina HiSeq | IDBA-UD | | PRJEB5265 |
| CruV-092 | 2789 | 50.8% | Standard | Unisense | #N/A | | Lake Aydat (45°39'52.859''N; 2°59'11.943''E) surface water | | vDNA, MDA | Illumina HiSeq | IDBA-UD | | MT478561 |
| CruV-093 | 2795 | 38.4% | Standard | Ambisense | TAATACTAA | | River (NZ) | | vDNA, MDA | Illumina HiSeq | metaSPAdes | | MT263542 |
| CruV-094 | 2797 | 40.2% | Standard | Ambisense | AAATATTAT | | Soil (NZ) | | vDNA, MDA | Illumina HiSeq | metaSPAdes | | MT263543 |
| CruV-095 | 2810 | 43.5% | Standard | Unisense | GATTAATAT | | Lake Aydat (45°39'52.859''N; 2°59'11.943''E) surface water | | vDNA, MDA | Illumina HiSeq | IDBA-UD | | MT478560 |
| CruV-096 | 2839 | 37.0% | Standard | Ambisense | TATTATAAT | | Lake Tenndammen (78°06.118′N; 15°02.024′E) | Aguirre de Cárcer et al., 2015 | vDNA, MDA | Illumina HiSeq | IDBA-UD | | PRJEB5265 |
| CruV-097 | 2839 | 36.8% | Standard | Ambisense | CAGTATTAC | | River (NZ) | | vDNA, MDA | Illumina HiSeq | metaSPAdes | | MT263544 |
| CruV-098 | 2849 | 37.2% | Standard | Ambisense | GATTATTAC / TACTATTAA / TAATAGTAA / TATTACTAT | | Lake Tenndammen (78°06.118′N; 15°02.024′E) | Aguirre de Cárcer et al., 2015 | vDNA, MDA | Illumina HiSeq | IDBA-UD | | PRJEB5265 |
| CruV-099 | 2860 | 40.1% | Standard | Ambisense | TATAGCTAC | | River bank soil (NZ) | | vDNA, MDA | Illumina HiSeq | metaSPAdes | | MT263545 |
| CruV-100 | 2867 | 57.7% | Standard | Ambisense | #N/A | | Lake Tunsjøen (78°03.375′N; 13°40.313′E) | Aguirre de Cárcer et al., 2015 | vDNA, MDA | Illumina HiSeq | IDBA-UD | | PRJEB5265 |
| CruV-101 | 2871 | 43.5% | Standard | Ambisense | TAATGTTAA | Cp-based clusters | Lake Nordammen (78°38.279′N; 16°44.025′E) | Aguirre de Cárcer et al., 2015 | vDNA, MDA | Illumina HiSeq | IDBA-UD | | PRJEB5265 |
| CruV-102 | 2874 | 40.9% | Standard | Ambisense | TAGTATTAC | | River bank soil (NZ) | | vDNA, MDA | Illumina HiSeq | metaSPAdes | | MT263546 |
| CruV-103 | 2881 | 46.9% | Standard | Ambisense | TAATATTAC | | Lake Nordammen (78°38.279′N; 16°44.025′E) | Aguirre de Cárcer et al., 2015 | vDNA, MDA | Illumina HiSeq | IDBA-UD | Spliced Rep | PRJEB5265 |
| CruV-104 | 2883 | 45.2% | Standard | Ambisense | TATTATAAT | | Lake Tunsjøen (78°03.375′N; 13°40.313′E) | Aguirre de Cárcer et al., 2015 | vDNA, MDA | Illumina HiSeq | IDBA-UD | Spliced Rep | PRJEB5265 |
| CruV-105 | 2883 | 45.1% | Standard | Ambisense | TAATATTAC | | unnamed Arctic pond (78°02.935′N; 13°41.973′E) | Aguirre de Cárcer et al., 2015 | vDNA, MDA | Illumina HiSeq | IDBA-UD | Spliced Rep | PRJEB5265 |
| CruV-106 | 2886 | 40.7% | Standard | Ambisense | #N/A | | Lake Aydat (45°39'52.859''N; 2°59'11.943''E) surface water | | vDNA, MDA | Illumina HiSeq | IDBA-UD | | MT478559 |
| CruV-107 | 2899 | 46.8% | Standard | Ambisense | #N/A | | unnamed Arctic pond (78°02.935′N; 13°41.973′E) | Aguirre de Cárcer et al., 2015 | vDNA, MDA | Illumina HiSeq | IDBA-UD | Spliced Rep | PRJEB5265 |
| CruV-108 | 2905 | 42.8% | Standard | Ambisense | CAATAATAT | | Lake Tenndammen (78°06.118′N; 15°02.024′E) | Aguirre de Cárcer et al., 2015 | vDNA, MDA | Illumina HiSeq | IDBA-UD | | PRJEB5265 |
| CruV-109 | 2911 | 38.1% | Standard | Ambisense | CATTAATAA / CATTACTAA | Rep-based clusters | unnamed Arctic pond (78°02.935′N; 13°41.973′E) | Aguirre de Cárcer et al., 2015 | vDNA, MDA | Illumina HiSeq | IDBA-UD | | PRJEB5265 |
| CruV-110 | 2919 | 34.8% | Standard | Ambisense | TACTATTAC | | Lake Tenndammen (78°06.118′N; 15°02.024′E) | Aguirre de Cárcer et al., 2015 | | | IDBA-UD | | PRJEB5265 |
| CruV-111 | 2929 | 36.5% | Standard | Ambisense | AAGTAATAA | | River (NZ) | | vDNA, MDA | Illumina HiSeq | metaSPAdes | | MT263547 |
| CruV-112 | 2942 | 37.4% | Standard | Ambisense | AATTACTAT | | River (NZ) | | vDNA, MDA | Illumina HiSeq | metaSPAdes | | MT263548 |
| CruV-113 | 2952 | 29.7% | Standard | Ambisense | GACTATTAC | | River (NZ) | | vDNA, MDA | Illumina HiSeq | metaSPAdes | | MT263549 |
| CruCGE-114 | 2953 | 33.9% | Standard | no rep | TAATATTAC | | unnamed Arctic pond (78°02.935′N; 13°41.973′E) | Aguirre de Cárcer et al., 2015 | vDNA, MDA | Illumina HiSeq | IDBA-UD | | PRJEB5265 |
| CruV-115 | 2954 | 37.0% | Standard | Ambisense | TATTTCAAG | | unnamed Arctic pond (78°02.935′N; 13°41.973′E) | Aguirre de Cárcer et al., 2015 | vDNA, MDA | Illumina HiSeq | IDBA-UD | | PRJEB5265 |
| CruV-116 | 2963 | 41.8% | Standard | Ambisense | #N/A | Cp-based clusters | Lake Nordammen (78°38.279′N; 16°44.025′E) | Aguirre de Cárcer et al., 2015 | vDNA, MDA | Illumina HiSeq | IDBA-UD | | PRJEB5265 |
| CruV-117 | 2965 | 37.9% | Standard | Unisense | TAAAATTAC / CATTATTAA | | Borgdammane pond (78°04.254′N; 13°47.652′E) | Aguirre de Cárcer et al., 2015 | vDNA, MDA | Roche 454 | IDBA-UD | | PRJEB5265 |
| CruCGE-118 | 2965 | 32.5% | Standard | Unisense | #N/A | | unnamed Arctic pond (78°02.935′N; 13°41.973′E) | Aguirre de Cárcer et al., 2015 | vDNA, MDA | Illumina HiSeq | IDBA-UD | | PRJEB5265 |
| CruV-119 | 2966 | 34.5% | Standard | Unisense | #N/A | | River (NZ) | | vDNA, MDA | Illumina HiSeq | metaSPAdes | | MT263550 |
| CruV-120 | 2983 | 40.6% | Standard | Ambisense | TATATAAAA | Cp-based clusters | Lake Tenndammen (78°06.118′N; 15°02.024′E) | Aguirre de Cárcer et al., 2015 | vDNA, MDA | Illumina HiSeq | IDBA-UD | | PRJEB5265 |
| CruV-121 | 2991 | 43.1% | Standard | Unisense | AAATACTAC | | unnamed Arctic pond (78°02.935′N; 13°41.973′E) | Aguirre de Cárcer et al., 2015 | vDNA, MDA | Illumina HiSeq | IDBA-UD | | PRJEB5265 |

| Name | Length | %GC | Genetic Code | Genome organization | ori (nona) | Sequences Subset | Location | Publication | Source | Sequencing | Assembly software | Notes | Accession number* |
|---|---|---|---|---|---|---|---|---|---|---|---|---|---|
| CruV-122 | 3003 | 38.4% | Standard | Unisense | TAATGTTAA | | River sediments (NZ) | | vDNA, MDA | Illumina HiSeq | metaSPAdes | | MT263551 |
| CruV-123 | 3009 | 38.6% | Standard | Ambisense | CAGTATTAC | | unnamed Arctic pond (78°02.935′N; 13°41.973′E) | Aguirre de Cárcer et al., 2015 | vDNA, MDA | Illumina HiSeq | IDBA-UD | | PRJEB5265 |
| CruV-124 | 3012 | 33.6% | Standard | Ambisense | #N/A | Cp-based clusters | Sewage Oxydation Pond (NZ) | | vDNA, MDA | Illumina HiSeq | metaSPAdes | | MT263552 |
| CruV-125 | 3012 | 46.1% | Standard | Ambisense | #N/A | | Lake Aydat (45°39'52.859''N; 2°59'11.943''E) surface water | | vDNA, MDA | Illumina HiSeq | IDBA-UD | | MT478558 |
| CruV-126 | 3019 | 46.0% | Standard | Unisense | #N/A | | Lake Tunsjøen (78°03.375′N; 13°40.313′E) | Aguirre de Cárcer et al., 2015 | vDNA, MDA | Illumina HiSeq | IDBA-UD | | PRJEB5265 |
| CruV-127 | 3020 | 43.2% | Standard | Ambisense | #N/A | | Lake Tenndammen (78°06.118′N; 15°02.024′E) | Aguirre de Cárcer et al., 2015 | vDNA, MDA | Illumina HiSeq | IDBA-UD | | PRJEB5265 |
| CruV-128 | 3023 | 49.7% | Standard | Unisense | TAGTATTAC | Rep-based clusters | Lake Aydat (45°39'52.859''N; 2°59'11.943''E) surface water | | vDNA, MDA | Illumina HiSeq | IDBA-UD | | MT478557 |
| CruV-129 | 3026 | 41.8% | Standard | Unisense | #N/A | | Lake Aydat (45°39'52.859''N; 2°59'11.943''E) surface water | | vDNA, MDA | Illumina HiSeq | IDBA-UD | | MT478556 |
| CruV-130 | 3026 | 34.4% | Standard | Ambisense | #N/A | | Lake Aydat (45°39'52.859''N; 2°59'11.943''E) surface water | | vDNA, MDA | Illumina HiSeq | IDBA-UD | | MT478555 |
| CruV-131 | 3030 | 48.0% | Standard | Ambisense | #N/A | | Lake Tenndammen (78°06.118′N; 15°02.024′E) | Aguirre de Cárcer et al., 2015 | vDNA, MDA | Illumina HiSeq | IDBA-UD | | PRJEB5265 |
| CruV-132 | 3030 | 39.2% | Standard | Ambisense | TAGTATTAC | | Lake Nordammen (78°38.279′N; 16°44.025′E) | Aguirre de Cárcer et al., 2015 | vDNA, MDA | Illumina HiSeq | IDBA-UD | | PRJEB5265 |
| CruV-133 | 3040 | 38.7% | Standard | Unisense | TACTATTAC | | Sewage Oxydation Pond (NZ) | | vDNA, MDA | Illumina HiSeq | metaSPAdes | | MT263553 |
| CruV-134 | 3049 | 40.1% | Standard | Unisense | #N/A | | Lake Aydat (45°39'52.859''N; 2°59'11.943''E) surface water | | vDNA, MDA | Illumina HiSeq | IDBA-UD | | MT478554 |
| CruV-135 | 3052 | 42.8% | Standard | Ambisense | AATTACTAT | | Lake Aydat (45°39'52.859''N; 2°59'11.943''E) surface water | | vDNA, MDA | Illumina HiSeq | IDBA-UD | | MT478553 |
| CruV-136 | 3054 | 56.2% | Standard | Ambisense | TAGTAATAG TATTACTAC / AACTATTAG | Rep-based clusters | Lake Aydat (45°39'52.859''N; 2°59'11.943''E) surface water | | vDNA, MDA | Illumina HiSeq | IDBA-UD | | MT478552 |
| CruV-137 | 3056 | 43.0% | Standard | Ambisense | #N/A | Rep-based clusters | Lake Aydat (45°39'52.859''N; 2°59'11.943''E) surface water | | vDNA, MDA | Illumina HiSeq | IDBA-UD | | MT478551 |
| CruV-138 | 3060 | 46.3% | Standard | Ambisense | TAATACTAC | | Lake Aydat (45°39'52.859''N; 2°59'11.943''E) surface water | | vDNA, MDA | Illumina HiSeq | IDBA-UD | | MT478550 |
| CruV-139 | 3065 | 36.2% | Standard | Ambisense | #N/A | | unnamed Arctic pond (78°02.935′N; 13°41.973′E) | Aguirre de Cárcer et al., 2015 | vDNA, MDA | Illumina HiSeq | IDBA-UD | | PRJEB5265 |
| CruV-140 | 3066 | 40.2% | Standard | Ambisense | #N/A | | Lake Aydat (45°39'52.859''N; 2°59'11.943''E) surface water | | vDNA, MDA | Illumina HiSeq | IDBA-UD | | MT478549 |
| CruV-141 | 3068 | 39.0% | Standard | Ambisense | TATTTCTAC | | Soil (NZ) | | vDNA, MDA | Illumina HiSeq | metaSPAdes | | MT263554 |
| CruV-142 | 3068 | 35.8% | Standard | Ambisense | TAGTATTAC | | Lake Tenndammen (78°06.118′N; 15°02.024′E) | Aguirre de Cárcer et al., 2015 | vDNA, MDA | Illumina HiSeq | IDBA-UD | | PRJEB5265 |
| CruV-143 | 3069 | 33.3% | Standard | Ambisense | #N/A | | Lake Nordammen (78°38.279′N; 16°44.025′E) | Aguirre de Cárcer et al., 2015 | vDNA, MDA | Illumina HiSeq | IDBA-UD | | PRJEB5265 |
| CruV-144 | 3070 | 47.9% | Standard | Unisense | TAGTATTAC | | Lake Tenndammen (78°06.118′N; 15°02.024′E) | Aguirre de Cárcer et al., 2015 | vDNA, MDA | Illumina HiSeq | IDBA-UD | | PRJEB5265 |
| CruV-145 | 3076 | 54.1% | Standard | Ambisense | TATAGTAAG / TATAAAAAC / TATTATAAG | | Bat (China) | Wu et al. 2015 | vDNA, MDA | Illumina | IDBA-UD | | SRR2063921 |
| CruV-146 | 3084 | 49.6% | Standard | Unisense | TACTACTAA / TAGTAGTAA | | River (NZ) | | vDNA, MDA | Illumina HiSeq | metaSPAdes | | MT263555 |
| CruV-147 | 3084 | 49.7% | Standard | Unisense | AATTATTAA | | River (NZ) | | vDNA, MDA | Illumina HiSeq | metaSPAdes | | MT263556 |
| CruV-148 | 3084 | 49.6% | Standard | Unisense | #N/A | | River (NZ) | | vDNA, MDA | Illumina HiSeq | metaSPAdes | | MT263557 |
| CruV-149 | 3089 | 43.2% | Standard | Unisense | #N/A | | Lake Tunsjøen (78°03.375′N; 13°40.313′E) | Aguirre de Cárcer et al., 2015 | vDNA, MDA | Illumina HiSeq | IDBA-UD | | PRJEB5265 |
| CruV-150 | 3094 | 42.6% | Standard | Unisense | TAGTATTAC | | Lake Pavin (45°29'45.11''N; 2°53'14.60''E), sampling depth = 22 meters | | vDNA, MDA | Illumina HiSeq | IDBA-UD | | MT478548 |
| CruV-151 | 3096 | 37.6% | Standard | Unisense | TAGTATTAC | Cp-based clusters | Lake Tenndammen (78°06.118′N; 15°02.024′E) | Aguirre de Cárcer et al., 2015 | vDNA, MDA | Illumina HiSeq | IDBA-UD | | PRJEB5265 |
| CruV-152 | 3096 | 50.5% | Standard | Unisense | TAGTATTAC | | River (NZ) | | vDNA, MDA | Illumina HiSeq | metaSPAdes | | MT263558 |
| CruV-153 | 3097 | 43.9% | Standard | Ambisense | TATTGTTAC | | Lake Aydat (45°39'52.859''N; 2°59'11.943''E) surface water | | vDNA, MDA | Illumina HiSeq | IDBA-UD | | MT478547 |
| CruV-154 | 3099 | 51.4% | Standard | Unisense | #N/A | Rep-based clusters | Lake Aydat (45°39'52.859''N; 2°59'11.943''E) surface water | | vDNA, MDA | Illumina HiSeq | IDBA-UD | | MT478546 |
| CruCGE-155 | 3102 | 41.2% | Standard | no rep | TAGTATTAC | | Lake Aydat (45°39'52.859''N; 2°59'11.943''E) surface water | | vDNA, MDA | Illumina HiSeq | IDBA-UD | | MT478562 |
| CruV-156 | 3105 | 44.7% | Standard | Unisense | #N/A | | Lake Nordammen (78°38.279′N; 16°44.025′E) | Aguirre de Cárcer et al., 2015 | vDNA, MDA | Illumina HiSeq | IDBA-UD | | PRJEB5265 |
| CruV-157 | 3113 | 35.8% | Standard | Unisense | TAGTATTAC | | Lake Aydat (45°39'52.859''N; 2°59'11.943''E) surface water | | vDNA, MDA | Illumina HiSeq | IDBA-UD | | MT478545 |
| CruV-158 | 3121 | 34.2% | Standard | Unisense | #N/A | | River (NZ) | | vDNA, MDA | Illumina HiSeq | metaSPAdes | | MT263559 |
| CruV-159 | 3122 | 44.6% | Standard | Ambisense | #N/A | Rep-based clusters | Lake Aydat (45°39'52.859''N; 2°59'11.943''E) surface water | | vDNA, MDA | Illumina HiSeq | IDBA-UD | | MT478544 |
| CruV-160 | 3129 | 38.2% | Standard | Unisense | TAGTATTAC | | Lake Aydat (45°39'52.859''N; 2°59'11.943''E) surface water | | vDNA, MDA | Illumina HiSeq | IDBA-UD | | MT478543 |

| Name | Length | %GC | Genetic Code | Genome organization | ori (nona) | Sequences Subset | Location | Publication | Source | Sequencing | Assembly software | Notes | Accession number* |
|---|---|---|---|---|---|---|---|---|---|---|---|---|---|
| CruV-161 | 3137 | 46.1% | Standard | Unisense | GAATATTAT / GACTATTAT / GAATAATAG | | unnamed Arctic pond (78°02.935′N; 13°41.973′E) | Aguirre de Cárcer et al., 2015 | vDNA, MDA | Illumina HiSeq | IDBA-UD | | PRJEB5265 |
| CruV-162 | 3139 | 37.8% | Standard | Ambisense | #N/A | | Lake Tunsjøen (78°03.375′N; 13°40.313′E) | Aguirre de Cárcer et al., 2015 | vDNA, MDA | Illumina HiSeq | IDBA-UD | | PRJEB5265 |
| CruV-163 | 3142 | 37.6% | Standard | Unisense | #N/A | Cp-based clusters | River bank soil (NZ) | | vDNA, MDA | Illumina HiSeq | metaSPAdes | | MT263560 |
| CruV-164 | 3142 | 37.7% | Standard | Ambisense | TAGTATTAC | | unnamed Arctic pond (78°02.935′N; 13°41.973′E) | Aguirre de Cárcer et al., 2015 | vDNA, MDA | Illumina HiSeq | IDBA-UD | | PRJEB5265 |
| CruV-165 | 3142 | 37.8% | Standard | Unisense | #N/A | | unnamed Arctic pond (78°02.935′N; 13°41.973′E) | Aguirre de Cárcer et al., 2015 | vDNA, MDA | Illumina HiSeq | IDBA-UD | | PRJEB5265 |
| CruV-166 | 3144 | 39.5% | Standard | Unisense | AACTAATAA | | unnamed Arctic pond (78°02.935′N; 13°41.973′E) | Aguirre de Cárcer et al., 2015 | vDNA, MDA | Illumina HiSeq | IDBA-UD | | PRJEB5265 |
| CruV-167 | 3145 | 44.0% | Standard | Unisense | CAATACTAG | Rep-based clusters | unnamed Arctic pond (78°02.935′N; 13°41.973′E) | Aguirre de Cárcer et al., 2015 | vDNA, MDA | Illumina HiSeq | IDBA-UD | | PRJEB5265 |
| CruV-168 | 3150 | 32.4% | Standard | Unisense | #N/A | | River (NZ) | | vDNA, MDA | Illumina HiSeq | metaSPAdes | | MT263561 |
| CruV-169 | 3150 | 38.5% | Standard | Unisense | TATTGCTAC / TAATTATAG | | Lake Nordammen (78°38.279′N; 16°44.025′E) | Aguirre de Cárcer et al., 2015 | vDNA, MDA | Illumina HiSeq | IDBA-UD | | PRJEB5265 |
| CruV-170 | 3154 | 54.5% | Standard | Unisense | TATTTAAAA / AAATAATAA | | Soil (NZ) | | vDNA, MDA | Illumina HiSeq | metaSPAdes | Spliced Rep | MT263562 |
| CruV-171 | 3157 | 45.7% | Standard | Ambisense | TAGTATTAC | | Lake Aydat (45°39'52.859''N; 2°59'11.943''E) surface water | | vDNA, MDA | Illumina HiSeq | IDBA-UD | | MT478542 |
| CruV-172 | 3168 | 40.4% | Standard | Ambisense | TATTGCTAG | | Lake Nordammen (78°38.279′N; 16°44.025′E) | Aguirre de Cárcer et al., 2015 | vDNA, MDA | Illumina HiSeq | IDBA-UD | | PRJEB5265 |
| CruV-173 | 3171 | 47.5% | Standard | Ambisense | TATAATAAT / TATTATTAT / TAATAATAA / GATTATTAT / CACTATTAA | | Lake Aydat (45°39'52.859''N; 2°59'11.943''E) surface water | | vDNA, MDA | Illumina HiSeq | IDBA-UD | | MT478541 |
| CruV-174 | 3172 | 43.9% | Standard | Ambisense | #N/A | | River (NZ) | | vDNA, MDA | Illumina HiSeq | metaSPAdes | Spliced Rep | MT263563 |
| CruV-175 | 3172 | 43.8% | Standard | Ambisense | TAGTATTAC | | Chiromindae (NZ) | | vDNA, MDA | Illumina HiSeq | metaSPAdes | | MT263564 |
| CruV-176 | 3176 | 30.3% | Standard | Unisense | TATTGCTAG | | Lake Aydat (45°39'52.859''N; 2°59'11.943''E) surface water | | vDNA, MDA | Illumina HiSeq | IDBA-UD | | MT478540 |
| CruV-177 | 3176 | 30.1% | Standard | Unisense | #N/A | | unnamed Arctic pond (78°02.935′N; 13°41.973′E) | Aguirre de Cárcer et al., 2015 | vDNA, MDA | Illumina HiSeq | IDBA-UD | | PRJEB5265 |
| CruV-178 | 3177 | 39.1% | Standard | Ambisense | TAGTATTAC | Cp-based clusters | Lake Aydat (45°39'52.859''N; 2°59'11.943''E) surface water | | vDNA, MDA | Illumina HiSeq | IDBA-UD | | MT478539 |
| CruV-179 | 3180 | 33.1% | Standard | Unisense | #N/A | | Lake Tunsjøen (78°03.375′N; 13°40.313′E) | Aguirre de Cárcer et al., 2015 | vDNA, MDA | Illumina HiSeq | IDBA-UD | | PRJEB5265 |
| CruV-180 | 3180 | 33.1% | Standard | Unisense | TAGTATTAC | | unnamed Arctic pond (78°02.935′N; 13°41.973′E) | Aguirre de Cárcer et al., 2015 | vDNA, MDA | Illumina HiSeq | IDBA-UD | | PRJEB5265 |
| CruV-181 | 3181 | 36.0% | Standard | Unisense | AACTATTAC | | Lake Tunsjøen (78°03.375′N; 13°40.313′E) | Aguirre de Cárcer et al., 2015 | vDNA, MDA | Illumina HiSeq | IDBA-UD | | PRJEB5265 |
| CruV-182 | 3199 | 29.5% | Standard | Unisense | AACTATTAC | | Estuary benthic sediments (NZ) | | vDNA, MDA | Illumina HiSeq | metaSPAdes | | MT263565 |
| CruV-183 | 3202 | 57.2% | Standard | Ambisense | GATTAATAT | | Lake Mary (AZ) | | vDNA, MDA | Illumina HiSeq | metaSPAdes | | MT263566 |
| CruV-184 | 3206 | 47.6% | Standard | Ambisense | CATTAATAT | | | | vDNA, MDA | Illumina HiSeq | metaSPAdes | | MT263533 |
| CruV-185 | 3212 | 44.5% | Standard | Ambisense | CAATAATAT | | unnamed Arctic pond (78°02.935′N; 13°41.973′E) | Aguirre de Cárcer et al., 2015 | vDNA, MDA | Illumina HiSeq | IDBA-UD | | PRJEB5265 |
| CruV-186 | 3222 | 40.2% | Standard | Ambisense | CAGTATTAC | | Lake Tenndammen (78°06.118′N; 15°02.024′E) | Aguirre de Cárcer et al., 2015 | vDNA, MDA | Illumina HiSeq | IDBA-UD | | PRJEB5265 |
| CruV-187 | 3227 | 52.3% | Standard | Ambisense | CAGTATTAC | | Lake Aydat (45°39'52.859''N; 2°59'11.943''E) surface water | | vDNA, MDA | Illumina HiSeq | IDBA-UD | | MT478538 |
| CruV-188 | 3230 | 35.3% | Standard | Unisense | #N/A | | Lake Aydat (45°39'52.859''N; 2°59'11.943''E) surface water | | vDNA, MDA | Illumina HiSeq | IDBA-UD | | MT478537 |
| CruV-189 | 3230 | 35.7% | Standard | Unisense | TAGTATTAC | | unnamed Arctic pond (78°02.935′N; 13°41.973′E) | Aguirre de Cárcer et al., 2015 | vDNA, MDA | Illumina HiSeq | IDBA-UD | | PRJEB5265 |
| CruV-190 | 3241 | 31.7% | Standard | Unisense | #N/A | | River (NZ) | | vDNA, MDA | Illumina HiSeq | metaSPAdes | | MT263567 |
| CruV-191 | 3243 | 50.4% | Standard | Ambisense | TAATACTAC | Cp-based clusters | Lake Tunsjøen (78°03.375′N; 13°40.313′E) | Aguirre de Cárcer et al., 2015 | vDNA, MDA | Illumina HiSeq | IDBA-UD | | PRJEB5265 |
| CruV-192 | 3244 | 37.3% | Standard | Ambisense | #N/A | | Lake Aydat (45°39'52.859''N; 2°59'11.943''E) surface water | | vDNA, MDA | Illumina HiSeq | IDBA-UD | | MT478536 |
| CruV-193 | 3248 | 39.9% | Standard | Ambisense | CAATAATAA | Cp-based clusters | Lake Aydat (45°39'52.859''N; 2°59'11.943''E) surface water | | vDNA, MDA | Illumina HiSeq | IDBA-UD | | MT478535 |
| CruV-194 | 3250 | 42.6% | Standard | Unisense | #N/A | | Lake Aydat (45°39'52.859''N; 2°59'11.943''E) surface water | | vDNA, MDA | Illumina HiSeq | IDBA-UD | | MT478534 |
| CruV-195 | 3256 | 45.1% | Standard | Unisense | #N/A | Cp-based clusters | Chiromindae (NZ) | | vDNA, MDA | Illumina HiSeq | metaSPAdes | | MT263568 |
| CruV-196 | 3267 | 29.9% | Standard | Ambisense | #N/A | | Lake Tenndammen (78°06.118′N; 15°02.024′E) | Aguirre de Cárcer et al., 2015 | vDNA, MDA | Illumina HiSeq | IDBA-UD | | PRJEB5265 |
| CruV-197 | 3273 | 40.5% | Standard | Ambisense | TATTATTAC | Cp-based clusters | Lake Aydat (45°39'52.859''N; 2°59'11.943''E) surface water | | vDNA, MDA | Illumina HiSeq | IDBA-UD | | MT478533 |
| CruV-198 | 3279 | 43.5% | Standard | Ambisense | #N/A | | unnamed Arctic pond (78°02.935′N; 13°41.973′E) | Aguirre de Cárcer et al., 2015 | vDNA, MDA | Illumina HiSeq | IDBA-UD | | PRJEB5265 |
| CruV-199 | 3279 | 44.9% | Standard | Unisense | #N/A | | Lake Aydat (45°39'52.859''N; 2°59'11.943''E) surface water | | vDNA, MDA | Illumina HiSeq | IDBA-UD | | MT478532 |

| Name | Length | %GC | Genetic Code | Genome organization | ori (nona) | Sequences Subset | Location | Publication | Source | Sequencing | Assembly software | Notes | Accession number* |
|---|---|---|---|---|---|---|---|---|---|---|---|---|---|
| CruV-200 | 3282 | 34.0% | Standard | Unisense | #N/A | | Lake Aydat (45°39'52.859''N; 2°59'11.943''E) surface water | | vDNA, MDA | Illumina HiSeq | IDBA-UD | | MT478531 |
| CruV-201 | 3283 | 40.9% | Standard | Ambisense | TAATAATAG / TATTATTAC | | unnamed Arctic pond (78°02.935′N; 13°41.973′E) | Aguirre de Cárcer et al., 2015 | vDNA, MDA | Illumina HiSeq | IDBA-UD | | PRJEB5265 |
| CruV-202 | 3285 | 40.3% | Standard | Unisense | #N/A | | Lake Aydat (45°39'52.859''N; 2°59'11.943''E) surface water | | vDNA, MDA | Illumina HiSeq | IDBA-UD | | MT478530 |
| CruV-203 | 3299 | 48.3% | Standard | Ambisense | TAGTATTAC | | unnamed Arctic pond (78°02.935′N; 13°41.973′E) | Aguirre de Cárcer et al., 2015 | vDNA, MDA | Illumina HiSeq | IDBA-UD | | PRJEB5265 |
| CruV-204 | 3303 | 37.1% | Standard | Ambisense | #N/A | | River (NZ) | | vDNA, MDA | Illumina HiSeq | metaSPAdes | | MT263569 |
| CruV-205 | 3303 | 37.1% | Standard | Ambisense | TATAACTAG / CAGTATTAG | | River (NZ) | | vDNA, MDA | Illumina HiSeq | metaSPAdes | | MT263570 |
| CruV-206 | 3304 | 45.1% | Standard | Ambisense | #N/A | | Lake Aydat (45°39'52.859''N; 2°59'11.943''E) surface water | | vDNA, MDA | Illumina HiSeq | IDBA-UD | | MT478529 |
| CruV-207 | 3304 | 35.8% | Standard | Unisense | #N/A | Cp-based clusters | River sediments (NZ) | | vDNA, MDA | Illumina HiSeq | metaSPAdes | | MT263571 |
| CruV-208 | 3306 | 42.4% | Standard | Unisense | #N/A | | Lake Tunsjøen (78°03.375′N; 13°40.313′E) | Aguirre de Cárcer et al., 2015 | vDNA, MDA | Illumina HiSeq | IDBA-UD | | PRJEB5265 |
| CruV-209 | 3309 | 45.4% | Standard | Ambisense | #N/A | | Lake Tenndammen (78°06.118′N; 15°02.024′E) | Aguirre de Cárcer et al., 2015 | vDNA, MDA | Illumina HiSeq | IDBA-UD | | PRJEB5265 |
| CruV-210 | 3310 | 39.1% | Standard | Ambisense | TATTATTAC | | Lake Aydat (45°39'52.859''N; 2°59'11.943''E) surface water | | vDNA, MDA | Illumina HiSeq | IDBA-UD | | MT478528 |
| CruV-211 | 3310 | 38.8% | Standard | Ambisense | #N/A | Rep-based clusters | River (NZ) | | vDNA, MDA | Illumina HiSeq | metaSPAdes | | MT263572 |
| CruV-212 | 3311 | 34.0% | Standard | Unisense | TATTATTAC | | River (NZ) | | vDNA, MDA | Illumina HiSeq | metaSPAdes | | MT263573 |
| CruV-213 | 3314 | 49.7% | Standard | Ambisense | TATTATTAC | | unnamed Arctic pond (78°02.935′N; 13°41.973′E) | Aguirre de Cárcer et al., 2015 | vDNA, MDA | Illumina HiSeq | IDBA-UD | | PRJEB5265 |
| CruV-214 | 3316 | 41.5% | Standard | Ambisense | TAGTATTAC | | Lake Nordammen (78°38.279′N; 16°44.025′E) | Aguirre de Cárcer et al., 2015 | vDNA, MDA | Illumina HiSeq | IDBA-UD | | PRJEB5265 |
| CruV-215 | 3326 | 54.0% | Standard | Unisense | TAATATTAC | Rep-based clusters | Lake Tenndammen (78°06.118′N; 15°02.024′E) | Aguirre de Cárcer et al., 2015 | vDNA, MDA | Illumina HiSeq | IDBA-UD | | PRJEB5265 |
| CruV-216 | 3330 | 30.6% | Standard | Unisense | CAGTATTAC | | unnamed Arctic pond (78°02.935′N; 13°41.973′E) | Aguirre de Cárcer et al., 2015 | vDNA, MDA | Illumina HiSeq | IDBA-UD | | PRJEB5265 |
| CruV-217 | 3333 | 51.8% | Standard | Ambisense | ...G | Rep-based clusters | Lake Aydat (45°39'52.859''N; 2°59'11.943''E) surface water | | vDNA, MDA | Illumina HiSeq | IDBA-UD | | MT478527 |
| CruV-218 | 3344 | 31.8% | Standard | Ambisense | TATTTAAAT | | Lake Aydat (45°39'52.859''N; 2°59'11.943''E) surface water | | vDNA, MDA | Illumina HiSeq | IDBA-UD | | MT478526 |
| CruV-219 | 3344 | 40.1% | Standard | Unisense | #N/A | | Lake Aydat (45°39'52.859''N; 2°59'11.943''E) surface water | | vDNA, MDA | Illumina HiSeq | IDBA-UD | | MT478525 |
| CruV-220 | 3344 | 40.2% | Standard | Ambisense | #N/A | Rep-based clusters | Lake Nordammen (78°38.279′N; 16°44.025′E) | Aguirre de Cárcer et al., 2015 | vDNA, MDA | Illumina HiSeq | IDBA-UD | | PRJEB5265 |
| CruV-221 | 3350 | 35.0% | Standard | Unisense | #N/A | | unnamed Arctic pond (78°02.935′N; 13°41.973′E) | Aguirre de Cárcer et al., 2015 | vDNA, MDA | Illumina HiSeq | IDBA-UD | | PRJEB5265 |
| CruV-222 | 3352 | 47.7% | Standard | Ambisense | TAAAGATAT | Rep-based clusters | Lake Aydat (45°39'52.859''N; 2°59'11.943''E) surface water | | vDNA, MDA | Illumina HiSeq | IDBA-UD | | MT478524 |
| CruV-223 | 3354 | 44.9% | Standard | Ambisense | TAAAGTTAT | | unnamed Arctic pond (78°02.935′N; 13°41.973′E) | Aguirre de Cárcer et al., 2015 | vDNA, MDA | Illumina HiSeq | IDBA-UD | | PRJEB5265 |
| CruV-224 | 3359 | 43.8% | Standard | Ambisense | TAGTATTAC | | River bank soil (NZ) | | vDNA, MDA | Illumina HiSeq | metaSPAdes | | MT263574 |
| CruV-225 | 3362 | 36.2% | Standard | Unisense | #N/A | | Lake Pavin (45°29'45.11''N; 2°53'14.60''E), sampling depth = 22 meters | | vDNA, MDA | Illumina HiSeq | IDBA-UD | | MT478523 |
| CruV-226 | 3362 | 35.8% | Standard | Ambisense | #N/A | Cp-based clusters | unnamed Arctic pond (78°02.935′N; 13°41.973′E) | Aguirre de Cárcer et al., 2015 | vDNA, MDA | Illumina HiSeq | IDBA-UD | | PRJEB5265 |
| CruV-227 | 3365 | 42.5% | Standard | Unisense | #N/A | | unnamed Arctic pond (78°02.935′N; 13°41.973′E) | Aguirre de Cárcer et al., 2015 | vDNA, MDA | Illumina HiSeq | IDBA-UD | | PRJEB5265 |
| CruV-228 | 3369 | 37.8% | Standard | Ambisense | #N/A | | Lake Tunsjøen (78°03.375′N; 13°40.313′E) | Aguirre de Cárcer et al., 2015 | vDNA, MDA | Illumina HiSeq | IDBA-UD | | PRJEB5265 |
| CruV-229 | 3372 | 34.4% | Standard | Unisense | #N/A | | Lake Aydat (45°39'52.859''N; 2°59'11.943''E) surface water | | vDNA, MDA | Illumina HiSeq | IDBA-UD | | MT478522 |
| CruV-230 | 3373 | 46.8% | Standard | Ambisense | CAATATTAC | | Lake Aydat (45°39'52.859''N; 2°59'11.943''E) surface water | | vDNA, MDA | Illumina HiSeq | IDBA-UD | | MT478521 |
| CruV-231 | 3375 | 48.8% | Standard | Ambisense | TATTATTAA | | Lake Aydat (45°39'52.859''N; 2°59'11.943''E) surface water | | vDNA, MDA | Illumina HiSeq | IDBA-UD | | MT478520 |
| CruV-232 | 3383 | 49.6% | Standard | Ambisense | TAGTATTAC | | Lake Tenndammen (78°06.118′N; 15°02.024′E) | Aguirre de Cárcer et al., 2015 | vDNA, MDA | Illumina HiSeq | IDBA-UD | | PRJEB5265 |
| CruV-233 | 3384 | 42.2% | Standard | Ambisense | TATTATTAC | Cp- and Rep-based clusters | Lake Aydat (45°39'52.859''N; 2°59'11.943''E) surface water | | vDNA, MDA | Illumina HiSeq | IDBA-UD | | MT478519 |
| CruV-234 | 3385 | 37.6% | Standard | Ambisense | AAATAATAT | Rep-based clusters | River (NZ) | | vDNA, MDA | Illumina HiSeq | metaSPAdes | | MT263575 |
| CruV-235 | 3386 | 48.3% | Standard | Ambisense | TACTATTAC | Rep-based clusters | Lake Aydat (45°39'52.859''N; 2°59'11.943''E) surface water | | vDNA, MDA | Illumina HiSeq | IDBA-UD | | MT478518 |
| CruV-236 | 3391 | 37.0% | Standard | Ambisense | GATTATTAT | | unnamed Arctic pond (78°02.935′N; 13°41.973′E) | Aguirre de Cárcer et al., 2015 | vDNA, MDA | Illumina HiSeq | IDBA-UD | | PRJEB5265 |
| CruV-237 | 3394 | 48.1% | Standard | Ambisense | #N/A | | Lake Tenndammen (78°06.118′N; 15°02.024′E) | Aguirre de Cárcer et al., 2015 | vDNA, MDA | Illumina HiSeq | IDBA-UD | | PRJEB5265 |

| Name | Length | %GC | Genetic Code | Genome organization | ori (nona) | Sequences Subset | Location | Publication | Source | Sequencing | Assembly software | Notes | Accession number* |
|---|---|---|---|---|---|---|---|---|---|---|---|---|---|
| CruV-238 | 3396 | 39.5% | Standard | Unisense | #N/A | | unnamed Arctic pond (78°02.935′N; 13°41.973′E) | Aguirre de Cárcer et al., 2015 | vDNA, MDA | Illumina HiSeq | IDBA-UD | | PRJEB5265 |
| CruV-239 | 3398 | 39.6% | Standard | Unisense | CAGTATTAC | | Lake Tunsjøen (78°03.375′N; 13°40.313′E) | Aguirre de Cárcer et al., 2015 | vDNA, MDA | Illumina HiSeq | IDBA-UD | | PRJEB5265 |
| CruV-240 | 3400 | 35.8% | Standard | Ambisense | #N/A | | Arabidopsis rhizosphere microbial communities from the University of North Carolina | Lundberg et al. 2012 | eDNA | Illumina HiSeq | MEGAHIT v1.0.6 | | PRJNA336851 |
| CruV-241 | 3402 | 40.6% | Standard | Ambisense | TATTAGTAA / TACTAATAA | | River (NZ) | | vDNA, MDA | Illumina HiSeq | metaSPAdes | | MT263576 |
| CruV-242 | 3403 | 45.5% | Standard | Ambisense | AATTATTAC | Cp-based clusters | Lake Nordammen (78°38.279′N; 16°44.025′E) | Aguirre de Cárcer et al., 2015 | vDNA, MDA | Illumina HiSeq | IDBA-UD | | PRJEB5265 |
| CruV-243 | 3405 | 43.5% | Standard | Ambisense | TAGTATTAC | Cp-based clusters | River (NZ) | | vDNA, MDA | Illumina HiSeq | metaSPAdes | | MT263577 |
| CruV-244 | 3405 | 42.4% | Standard | Unisense | CACTAATAT | Cp-based clusters | unnamed Arctic pond (78°02.935′N; 13°41.973′E) | Aguirre de Cárcer et al., 2015 | vDNA, MDA | Illumina HiSeq | IDBA-UD | | PRJEB5265 |
| CruV-245 | 3406 | 43.5% | Standard | Ambisense | #N/A | | River (NZ) | | vDNA, MDA | Illumina HiSeq | metaSPAdes | | MT263578 |
| CruV-246 | 3408 | 37.8% | Standard | Ambisense | | | Lake Tenndammen (78°06.118′N; 15°02.024′E) | Aguirre de Cárcer et al., 2015 | vDNA, MDA | Illumina HiSeq | IDBA-UD | | PRJEB5265 |
| CruV-247 | 3413 | 43.2% | Standard | Ambisense | TAGTATTAC | | Lake Aydat (45°39'52.859''N; 2°59'11.943''E) surface water | | vDNA, MDA | Illumina HiSeq | IDBA-UD | | MT478517 |
| CruV-248 | 3413 | 36.0% | Standard | Ambisense | #N/A | | Lake Linnevatnet (78°03.864′N; 13°46.308′E) | Aguirre de Cárcer et al., 2015 | vDNA, MDA | Illumina HiSeq | IDBA-UD | | PRJEB5265 |
| CruV-249 | 3419 | 38.6% | Standard | Unisense | #N/A | | Lake Tenndammen (78°06.118′N; 15°02.024′E) | Aguirre de Cárcer et al., 2015 | vDNA, MDA | Illumina HiSeq | IDBA-UD | | PRJEB5265 |
| CruV-250 | 3420 | 47.2% | Standard | Ambisense | #N/A | | River (NZ) | | vDNA, MDA | Illumina HiSeq | metaSPAdes | | MT263579 |
| CruV-251 | 3420 | 47.2% | Standard | Ambisense | #N/A | | River bank soil (NZ) | | vDNA, MDA | Illumina HiSeq | metaSPAdes | | MT263580 |
| CruV-252 | 3423 | 47.0% | Standard | Ambisense | #N/A | Rep-based clusters | unnamed Arctic pond (78°02.935′N; 13°41.973′E) | Aguirre de Cárcer et al., 2015 | vDNA, MDA | Illumina HiSeq | IDBA-UD | | PRJEB5265 |
| CruV-253 | 3424 | 45.3% | Standard | Ambisense | #N/A | Cp- and Rep-based clusters | Lake Aydat (45°39'52.859''N; 2°59'11.943''E) surface water | | vDNA, MDA | Illumina HiSeq | IDBA-UD | | MT478516 |
| CruV-254 | 3425 | 47.4% | Standard | Ambisense | #N/A | Cp- and Rep-based clusters | Lake Tenndammen (78°06.118′N; 15°02.024′E) | Aguirre de Cárcer et al., 2015 | vDNA, MDA | Illumina HiSeq | IDBA-UD | | PRJEB5265 |
| CruV-255 | 3426 | 37.2% | Standard | Ambisense | GATTATTAT | | Sewage Oxydation Pond (NZ) | | vDNA, MDA | Illumina HiSeq | metaSPAdes | | MT263581 |
| CruV-256 | 3426 | 51.3% | Standard | Unisense | GAATAATAA | Rep-based clusters | Lake Aydat (45°39'52.859''N; 2°59'11.943''E) surface water | | vDNA, MDA | Illumina HiSeq | IDBA-UD | | MT478515 |
| CruV-257 | 3434 | 49.8% | Standard | Ambisense | TACTATTAA / TAATAGTAT | | Lake Nordammen (78°38.279′N; 16°44.025′E) | Aguirre de Cárcer et al., 2015 | vDNA, MDA | Illumina HiSeq | IDBA-UD | | PRJEB5265 |
| CruV-258 | 3435 | 37.4% | Standard | Unisense | TACTATTAA / TAATAGTAT | Cp-based clusters | unnamed Arctic pond (78°02.935′N; 13°41.973′E) | Aguirre de Cárcer et al., 2015 | vDNA, MDA | Illumina HiSeq | IDBA-UD | | PRJEB5265 |
| CruV-259 | 3437 | 26.6% | Ciliate | Unisense | TAATAATAT | | Lake Aydat (45°39'52.859''N; 2°59'11.943''E) surface water | | vDNA, MDA | Illumina HiSeq | IDBA-UD | | MT478514 |
| CruV-260 | 3439 | 34.2% | Standard | Ambisense | AATTATTAT | | Lake Aydat (45°39'52.859''N; 2°59'11.943''E) surface water | | vDNA, MDA | Illumina HiSeq | IDBA-UD | | MT478513 |
| CruV-261 | 3444 | 38.1% | Standard | Unisense | #N/A | | Lake Aydat (45°39'52.859''N; 2°59'11.943''E) surface water | | vDNA, MDA | Illumina HiSeq | IDBA-UD | | MT478512 |
| CruV-262 | 3447 | 47.5% | Standard | Ambisense | GAATAATAA | Cp- and Rep-based clusters | unnamed Arctic pond (78°02.935′N; 13°41.973′E) | Aguirre de Cárcer et al., 2015 | vDNA, MDA | Illumina HiSeq | IDBA-UD | | PRJEB5265 |
| CruV-263 | 3451 | 33.7% | Standard | Ambisense | TATAGTAAC | | Lake Aydat (45°39'52.859''N; 2°59'11.943''E) surface water | | vDNA, MDA | Illumina HiSeq | IDBA-UD | | MT478511 |
| CruV-264 | 3454 | 43.2% | Standard | Ambisense | #N/A | | River (NZ) | | vDNA, MDA | Illumina HiSeq | metaSPAdes | | MT263582 |
| CruV-265 | 3457 | 48.7% | Standard | Ambisense | TATAGTAAC | | River sediments (NZ) | | vDNA, MDA | Illumina HiSeq | metaSPAdes | | MT263583 |
| CruV-266 | 3458 | 48.6% | Standard | Ambisense | #N/A | | River sediments (NZ) | | vDNA, MDA | Illumina HiSeq | metaSPAdes | | MT263584 |
| CruV-267 | 3465 | 50.3% | Standard | Ambisense | CAGTATTAC | | Lake Aydat (45°39'52.859''N; 2°59'11.943''E) surface water | | vDNA, MDA | Illumina HiSeq | IDBA-UD | | MT478510 |
| CruV-268 | 3470 | 61.5% | Standard | Ambisense | #N/A | | Lake Nordammen (78°38.279′N; 16°44.025′E) | Aguirre de Cárcer et al., 2015 | vDNA, MDA | Illumina HiSeq | IDBA-UD | | PRJEB5265 |
| CruV-269 | 3475 | 38.7% | Standard | Unisense | TAGTATTAC | | unnamed Arctic pond (78°02.935′N; 13°41.973′E) | Aguirre de Cárcer et al., 2015 | vDNA, MDA | Illumina HiSeq | IDBA-UD | | PRJEB5265 |
| CruV-270 | 3477 | 37.5% | Standard | Ambisense | TAGTATTAC | | Lake Tenndammen (78°06.118′N; 15°02.024′E) | Aguirre de Cárcer et al., 2015 | vDNA, MDA | Illumina HiSeq | IDBA-UD | | PRJEB5265 |
| CruV-271 | 3479 | 36.4% | Standard | Unisense | TAGTATTAC | | unnamed Arctic pond (78°02.935′N; 13°41.973′E) | Aguirre de Cárcer et al., 2015 | vDNA, MDA | Illumina HiSeq | IDBA-UD | | PRJEB5265 |
| CruV-272 | 3479 | 48.3% | Standard | Ambisense | #N/A | | Dragonfly larvae (NZ) | | vDNA, MDA | Illumina HiSeq | metaSPAdes | | MT263585 |
| CruV-273 | 3485 | 50.2% | Standard | Unisense | #N/A | Rep-based clusters | Lake Aydat (45°39'52.859''N; 2°59'11.943''E) surface water | | vDNA, MDA | Illumina HiSeq | IDBA-UD | | MT478509 |
| CruV-274 | 3485 | 36.2% | Standard | Unisense | #N/A | | unnamed Arctic pond (78°02.935′N; 13°41.973′E) | Aguirre de Cárcer et al., 2015 | vDNA, MDA | Illumina HiSeq | IDBA-UD | | PRJEB5265 |
| CruV-275 | 3497 | 40.6% | Standard | Ambisense | #N/A | | South Island Robin feces (NZ) | | vDNA, MDA | Illumina HiSeq | metaSPAdes | | MT263586 |
| CruV-276 | 3498 | 43.0% | Standard | Ambisense | TAGTATTAC | | River (NZ) | | vDNA, MDA | Illumina HiSeq | metaSPAdes | | MT263587 |

| Name | Length | %GC | Genetic Code | Genome organization | ori (nona) | Sequences Subset | Location | Publication | Source | Sequencing | Assembly software | Notes | Accession number* |
|---|---|---|---|---|---|---|---|---|---|---|---|---|---|
| CruV-277 | 3499 | 48.2% | Standard | Ambisense | AAATATTAA / AATTAATAT | | River (NZ) | | vDNA, MDA | Illumina HiSeq | metaSPAdes | | MT263588 |
| CruV-278 | 3500 | 42.9% | Standard | Ambisense | #N/A | | River (NZ) | | vDNA, MDA | Illumina HiSeq | metaSPAdes | | MT263589 |
| CruV-279 | 3501 | 42.9% | Standard | Unisense | #N/A | | unnamed Arctic pond (78°02.935′N; 13°41.973′E) | Aguirre de Cárcer et al., 2015 | vDNA, MDA | Illumina HiSeq | IDBA-UD | Spliced Rep | PRJEB5265 |
| CruV-280 | 3507 | 44.1% | Standard | Ambisense | TAGTATTAC / TAATACTAG | Cp- and Rep-based clusters | Lake Aydat (45°39'52.859''N; 2°59'11.943''E) surface water | | vDNA, MDA | Illumina HiSeq | IDBA-UD | | MT478508 |
| CruV-281 | 3508 | 48.2% | Standard | Ambisense | TAGTATTAC | | Blood worms (NZ) | | vDNA, MDA | Illumina HiSeq | metaSPAdes | | MT263590 |
| CruV-282 | 3509 | 48.2% | Standard | Ambisense | TATTTCTAA | | River (NZ) | | vDNA, MDA | Illumina HiSeq | metaSPAdes | | MT263591 |
| CruV-283 | 3511 | 45.1% | Standard | Ambisense | TAATGTTAA | | Lake Aydat (45°39'52.859''N; 2°59'11.943''E) surface water | | vDNA, MDA | Illumina HiSeq | IDBA-UD | | MT478507 |
| CruV-284 | 3515 | 36.1% | Standard | Unisense | #N/A | | unnamed Arctic pond (78°02.935′N; 13°41.973′E) | Aguirre de Cárcer et al., 2015 | vDNA, MDA | Illumina HiSeq | IDBA-UD | | PRJEB5265 |
| CruV-285 | 3517 | 47.8% | Standard | Ambisense | CAATATTAC | Cp-based clusters | Lake Aydat (45°39'52.859''N; 2°59'11.943''E) surface water | | vDNA, MDA | Illumina HiSeq | IDBA-UD | | MT478506 |
| CruV-286 | 3518 | 34.8% | Standard | Unisense | TATAGTAAC | | Lake Pavin (45°29'45.11''N; 2°53'14.60''E), sampling depth = 22 meters | | vDNA, MDA | Illumina HiSeq | IDBA-UD | | MT478505 |
| CruV-287 | 3521 | 38.9% | Standard | Unisense | TAGTATTAC | | Lake Tunsjøen (78°03.375′N; 13°40.313′E) | Aguirre de Cárcer et al., 2015 | vDNA, MDA | Illumina HiSeq | IDBA-UD | | PRJEB5265 |
| CruV-288 | 3526 | 41.7% | Standard | Unisense | TAGTATTAC | | Lake Aydat (45°39'52.859''N; 2°59'11.943''E) surface water | | vDNA, MDA | Illumina HiSeq | IDBA-UD | | MT478504 |
| CruV-289 | 3536 | 46.1% | Standard | Ambisense | TACTATTAC | Rep-based clusters | Lake Aydat (45°39'52.859''N; 2°59'11.943''E) surface water | | vDNA, MDA | Illumina HiSeq | IDBA-UD | | MT478503 |
| CruV-290 | 3538 | 50.9% | Standard | Ambisense | #N/A | | Lake Aydat (45°39'52.859''N; 2°59'11.943''E) surface water | | vDNA, MDA | Illumina HiSeq | IDBA-UD | | MT478502 |
| CruV-291 | 3538 | 48.6% | Standard | Ambisense | #N/A | | River (NZ) | | vDNA, MDA | Illumina HiSeq | metaSPAdes | | MT263592 |
| CruV-292 | 3538 | 45.1% | Standard | Ambisense | #N/A | Rep-based clusters | Lake Aydat (45°39'52.859''N; 2°59'11.943''E) surface water | | vDNA, MDA | Illumina HiSeq | IDBA-UD | | MT478501 |
| CruV-293 | 3545 | 45.8% | Standard | Ambisense | TATAACTAG | | Odonatan larvae (NZ) | | vDNA, MDA | Illumina HiSeq | metaSPAdes | | MT263593 |
| CruV-294 | 3546 | 39.8% | Standard | Unisense | #N/A | Cp-based clusters | Lake Aydat (45°39'52.859''N; 2°59'11.943''E) surface water | | vDNA, MDA | Illumina HiSeq | IDBA-UD | | MT478500 |
| CruV-295 | 3548 | 52.3% | Standard | Ambisense | #N/A | | Sewage Oxydation Pond (NZ) | | vDNA, MDA | Illumina HiSeq | metaSPAdes | | MT263594 |
| CruV-296 | 3549 | 51.2% | Standard | Unisense | TATAAATAC | | Lake Tunsjøen (78°03.375′N; 13°40.313′E) | Aguirre de Cárcer et al., 2015 | vDNA, MDA | Illumina HiSeq | IDBA-UD | | PRJEB5265 |
| CruV-297 | 3549 | 40.1% | Standard | Ambisense | AAGTATTAT | Rep-based clusters | unnamed Arctic pond (78°02.935′N; 13°41.973′E) | Aguirre de Cárcer et al., 2015 | vDNA, MDA | Illumina HiSeq | IDBA-UD | | PRJEB5265 |
| CruV-298 | 3551 | 43.6% | Standard | Ambisense | TAGTATTAC | Cp-based clusters | Lake Mary (AZ) | | vDNA, MDA | Illumina HiSeq | metaSPAdes | | MT263595 |
| CruV-299 | 3552 | 47.1% | Standard | Ambisense | AACTAGTAT | Cp-based clusters | unnamed Arctic pond (78°02.935′N; 13°41.973′E) | Aguirre de Cárcer et al., 2015 | vDNA, MDA | Illumina HiSeq | IDBA-UD | | PRJEB5265 |
| CruV-300 | 3558 | 38.2% | Standard | Ambisense | TATTGTTAC | | Arabidopsis rhizosphere microbial communities from the University of North Carolina | Lundberg et al. 2012 | eDNA | Illumina GAIIx | MEGAHIT v1.0.6 | | PRJNA336798 |
| CruV-301 | 3560 | 46.0% | Standard | Unisense | AACTACTAT | Rep-based clusters | Lake Aydat (45°39'52.859''N; 2°59'11.943''E) surface water | | vDNA, MDA | Illumina HiSeq | IDBA-UD | | MT478499 |
| CruV-302 | 3563 | 41.4% | Standard | Unisense | TATAGTAAC | | Lake Nordammen (78°38.279′N; 16°44.025′E) | Aguirre de Cárcer et al., 2015 | vDNA, MDA | Illumina HiSeq | IDBA-UD | | PRJEB5265 |
| CruV-303 | 3564 | 50.2% | Standard | Unisense | #N/A | Rep-based clusters | Lake ALake Aydat (45°39'52.859''N; 2°59'11.943''E) surface waterydat | | vDNA, MDA | Illumina HiSeq | IDBA-UD | | MT478498 |
| CruV-304 | 3567 | 45.0% | Standard | Ambisense | TATAAATAC | | River (NZ) | | vDNA, MDA | Illumina HiSeq | metaSPAdes | | MT263596 |
| CruV-305 | 3569 | 47.1% | Standard | Unisense | AAGTATTAG | Rep-based clusters | River (NZ) | | vDNA, MDA | Illumina HiSeq | metaSPAdes | | MT263597 |
| CruV-306 | 3576 | 53.0% | Standard | Unisense | #N/A | Cp- and Rep-based clusters | River sediments (NZ) | | vDNA, MDA | Illumina HiSeq | metaSPAdes | | MT263598 |
| CruV-307 | 3577 | 50.6% | Standard | Ambisense | TATAGTAAC | | unnamed Arctic pond (78°02.935′N; 13°41.973′E) | Aguirre de Cárcer et al., 2015 | vDNA, MDA | Illumina HiSeq | IDBA-UD | | PRJEB5265 |
| CruV-308 | 3578 | 40.8% | Standard | Ambisense | #N/A | | Lake Aydat (45°39'52.859''N; 2°59'11.943''E) surface water | | vDNA, MDA | Illumina HiSeq | IDBA-UD | | MT478497 |
| CruV-309 | 3584 | 46.6% | Standard | Unisense | #N/A | Rep-based clusters | River (NZ) | | vDNA, MDA | Illumina HiSeq | metaSPAdes | | MT263599 |
| CruV-310 | 3584 | 46.6% | Standard | Unisense | #N/A | Rep-based clusters | River (NZ) | | vDNA, MDA | Illumina HiSeq | metaSPAdes | | MT263600 |
| CruV-311 | 3584 | 46.6% | Standard | Unisense | #N/A | Rep-based clusters | River (NZ) | | vDNA, MDA | Illumina HiSeq | metaSPAdes | | MT263601 |
| CruV-312 | 3593 | 35.8% | Standard | Ambisense | TAGTATTAC | | River (NZ) | | vDNA, MDA | Illumina HiSeq | metaSPAdes | | MT263602 |
| CruV-313 | 3594 | 46.8% | Standard | Unisense | TATAAATAC | | Lake Tenndammen (78°06.118′N; 15°02.024′E) | Aguirre de Cárcer et al., 2015 | vDNA, MDA | Illumina HiSeq | IDBA-UD | | PRJEB5265 |
| CruV-314 | 3597 | 49.0% | Standard | Unisense | TATAAATAC | | Lake Nordammen (78°38.279′N; 16°44.025′E) | Aguirre de Cárcer et al., 2015 | vDNA, MDA | Illumina HiSeq | IDBA-UD | | PRJEB5265 |

| Name | Length | %GC | Genetic Code | Genome organization | ori (nona) | Sequences Subset | Location | Publication | Source | Sequencing | Assembly software | Notes | Accession number* |
|---|---|---|---|---|---|---|---|---|---|---|---|---|---|
| CruV-315 | 3600 | 48.7% | Standard | Ambisense | #N/A | Cp-based clusters | unnamed Arctic pond (78°02.935′N; 13°41.973′E) | Aguirre de Cárcer et al., 2015 | vDNA, MDA | Illumina HiSeq | IDBA-UD | | PRJEB5265 |
| CruV-316 | 3603 | 42.4% | Standard | Unisense | GAATATTAT | | Lake Aydat (45°39'52.859''N; 2°59'11.943''E) surface water | | vDNA, MDA | Illumina HiSeq | IDBA-UD | | MT478496 |
| CruV-317 | 3611 | 26.1% | Standard | Unisense | #N/A | | Lake Pavin (45°29'45.11''N; 2°53'14.60''E), sampling depth = 22 meters | | vDNA, MDA | Illumina HiSeq | IDBA-UD | | MT478495 |
| CruV-318 | 3611 | 49.6% | Standard | Ambisense | #N/A | Cp-based clusters | Lake Aydat (45°39'52.859''N; 2°59'11.943''E) surface water | | vDNA, MDA | Illumina HiSeq | IDBA-UD | | MT478494 |
| CruV-319 | 3613 | 41.8% | Standard | Unisense | TAAAACAAA | | Lake Tunsjøen (78°03.375′N; 13°40.313′E) | Aguirre de Cárcer et al., 2015 | vDNA, MDA | Illumina HiSeq | IDBA-UD | | PRJEB5265 |
| CruV-320 | 3613 | 41.8% | Standard | Unisense | AACTATTAC | | unnamed Arctic pond (78°02.935′N; 13°41.973′E) | Aguirre de Cárcer et al., 2015 | vDNA, MDA | Illumina HiSeq | IDBA-UD | | PRJEB5265 |
| CruV-321 | 3614 | 47.6% | Standard | Ambisense | #N/A | | Lake Aydat (45°39'52.859''N; 2°59'11.943''E) surface water | | vDNA, MDA | Illumina HiSeq | IDBA-UD | | MT478493 |
| CruV-322 | 3616 | 46.1% | Standard | Unisense | TATTTAAAT | | River (NZ) | | vDNA, MDA | Illumina HiSeq | metaSPAdes | | MT263603 |
| CruV-323 | 3616 | 46.1% | Standard | Unisense | TATAAATAC | | River (NZ) | | vDNA, MDA | Illumina HiSeq | metaSPAdes | | MT263604 |
| CruV-324 | 3616 | 46.1% | Standard | Unisense | #N/A | | River bank soil (NZ) | | vDNA, MDA | Illumina HiSeq | metaSPAdes | | MT263605 |
| CruV-325 | 3619 | 30.3% | Standard | Unisense | #N/A | | unnamed Arctic pond (78°02.935′N; 13°41.973′E) | Aguirre de Cárcer et al., 2015 | vDNA, MDA | Illumina HiSeq | IDBA-UD | | PRJEB5265 |
| CruV-326 | 3620 | 44.0% | Standard | Ambisense | AATTATTAA | | River bank soil (NZ) | | vDNA, MDA | Illumina HiSeq | metaSPAdes | | MT263606 |
| CruV-327 | 3630 | 31.1% | Standard | Ambisense | TAATACTAC | | Lake Nordammen (78°38.279′N; 16°44.025′E) | Aguirre de Cárcer et al., 2015 | vDNA, MDA | Illumina HiSeq | IDBA-UD | | PRJEB5265 |
| CruV-328 | 3630 | 42.1% | Standard | Ambisense | #N/A | Rep-based clusters | unnamed Arctic pond (78°02.935′N; 13°41.973′E) | Aguirre de Cárcer et al., 2015 | vDNA, MDA | Illumina HiSeq | IDBA-UD | Capsid ORF has smaller ORF in opposite orientation that has homology to Circo Cap | PRJEB5265 |
| CruV-329 | 3635 | 40.2% | Standard | Ambisense | #N/A | | unnamed Arctic pond (78°02.935′N; 13°41.973′E) | Aguirre de Cárcer et al., 2015 | vDNA, MDA | Illumina HiSeq | IDBA-UD | | PRJEB5265 |
| CruV-330 | 3656 | 40.4% | Standard | Unisense | TAGTATTAC | | unnamed Arctic pond (78°02.935′N; 13°41.973′E) | Aguirre de Cárcer et al., 2015 | vDNA, MDA | Illumina HiSeq | IDBA-UD | | PRJEB5265 |
| CruV-331 | 3657 | 45.2% | Standard | Unisense | #N/A | | Marine microbial communities from Delaware Coast | | eDNA | Illumina HiSeq | SOAPdenovo, Newbler, Minimus2 | | PRJNA336828 |
| CruV-332 | 3658 | 62.1% | Standard | Unisense | #N/A | | Lake Nordammen (78°38.279′N; 16°44.025′E) | Aguirre de Cárcer et al., 2015 | vDNA, MDA | Illumina HiSeq | IDBA-UD | | PRJEB5265 |
| CruV-333 | 3665 | 45.3% | Standard | Ambisense | TAATAATAT / TATTATTAG | | Lake Aydat (45°39'52.859''N; 2°59'11.943''E) surface water | | vDNA, MDA | Illumina HiSeq | IDBA-UD | | MT478492 |
| CruV-334 | 3666 | 35.4% | Standard | Ambisense | TAGTATTAC | | Lake Aydat (45°39'52.859''N; 2°59'11.943''E) surface water | | vDNA, MDA | Illumina HiSeq | IDBA-UD | | MT478491 |
| CruV-335 | 3667 | 33.0% | Standard | Ambisense | TATAGATAA | | unnamed Arctic pond (78°02.935′N; 13°41.973′E) | Aguirre de Cárcer et al., 2015 | vDNA, MDA | Illumina HiSeq | IDBA-UD | | PRJEB5265 |
| CruV-336 | 3676 | 43.5% | Standard | Unisense | #N/A | Cp-based clusters | unnamed Arctic pond (78°02.935′N; 13°41.973′E) | Aguirre de Cárcer et al., 2015 | vDNA, MDA | Illumina HiSeq | IDBA-UD | | PRJEB5265 |
| CruV-337 | 3682 | 37.5% | Standard | Unisense | #N/A | | Lake Tunsjøen (78°03.375′N; 13°40.313′E) | Aguirre de Cárcer et al., 2015 | vDNA, MDA | Illumina HiSeq | IDBA-UD | | PRJEB5265 |
| CruV-338 | 3686 | 36.1% | Standard | Ambisense | TAGTATTAC | | Host-associated microbial communities from peat moss Sphagnum species from Minnesota, USA | | eDNA | Illumina HiSeq | MEGAHIT v1.0.3 | | PRJNA364930 |
| CruV-339 | 3691 | 31.3% | Standard | Ambisense | CAATATTAG | | River (NZ) | | vDNA, MDA | Illumina HiSeq | metaSPAdes | | MT263607 |
| CruV-340 | 3708 | 38.0% | Standard | Ambisense | TAATAAAAT | Cp-based clusters | River (NZ) | | vDNA, MDA | Illumina HiSeq | metaSPAdes | Spliced Rep | MT263608 |
| CruV-341 | 3708 | 32.6% | Standard | Ambisense | TAATATTAC | | unnamed Arctic pond (78°02.935′N; 13°41.973′E) | Aguirre de Cárcer et al., 2015 | vDNA, MDA | Illumina HiSeq | IDBA-UD | | PRJEB5265 |
| CruV-342 | 3709 | 32.9% | Standard | Ambisense | TAGTATTAC | | unnamed Arctic pond (78°02.935′N; 13°41.973′E) | Aguirre de Cárcer et al., 2015 | vDNA, MDA | Illumina HiSeq | IDBA-UD | | PRJEB5265 |
| CruV-343 | 3719 | 49.7% | Standard | Ambisense | AATTATTAA | Rep-based clusters | Lake Aydat (45°39'52.859''N; 2°59'11.943''E) surface water | | vDNA, MDA | Illumina HiSeq | IDBA-UD | | MT478490 |
| CruV-344 | 3726 | 37.5% | Standard | Unisense | TAGTATTAC | | unnamed Arctic pond (78°02.935′N; 13°41.973′E) | Aguirre de Cárcer et al., 2015 | vDNA, MDA | Illumina HiSeq | IDBA-UD | | PRJEB5265 |
| CruV-345 | 3728 | 41.0% | Standard | Ambisense | TAGTATTAC | | River biofilm (NZ) | | vDNA, MDA | Illumina HiSeq | metaSPAdes | | MT263609 |
| CruV-346 | 3749 | 40.3% | Standard | Unisense | #N/A | | unnamed Arctic pond (78°02.935′N; 13°41.973′E) | Aguirre de Cárcer et al., 2015 | vDNA, MDA | Illumina HiSeq | IDBA-UD | Spliced Rep | PRJEB5265 |
| CruV-347 | 3766 | 38.4% | Standard | Unisense | TAATAGTAG / TACTATTAC | | unnamed Arctic pond (78°02.935′N; 13°41.973′E) | Aguirre de Cárcer et al., 2015 | vDNA, MDA | Illumina HiSeq | IDBA-UD | Spliced Rep | PRJEB5265 |
| CruV-348 | 3769 | 41.2% | Standard | Ambisense | #N/A | Cp-based clusters | Lake Tenndammen (78°06.118′N; 15°02.024′E) | Aguirre de Cárcer et al., 2015 | vDNA, MDA | Illumina HiSeq | IDBA-UD | | PRJEB5265 |
| CruV-349 | 3770 | 32.3% | Standard | Ambisense | #N/A | | River (NZ) | | vDNA, MDA | Illumina HiSeq | metaSPAdes | | MT263610 |
| CruV-350 | 3777 | 47.8% | Standard | Ambisense | TATT... | | River (NZ) | | vDNA, MDA | Illumina HiSeq | metaSPAdes | | MT263611 |
| CruV-351 | 3786 | 31.3% | Standard | Ambisense | #N/A | | Lake Nordammen (78°38.279′N; 16°44.025′E) | Aguirre de Cárcer et al., 2015 | vDNA, MDA | Illumina HiSeq | IDBA-UD | | PRJEB5265 |

| Name | Length | %GC | Genetic Code | Genome organization | ori (nona) | Sequences Subset | Location | Publication | Source | Sequencing | Assembly software | Notes | Accession number* |
|---|---|---|---|---|---|---|---|---|---|---|---|---|---|
| CruV-352 | 3790 | 34.9% | Standard | Ambisense | #N/A | | Lake Tunsjøen (78°03.375′N; 13°40.313′E) | Aguirre de Cárcer et al., 2015 | vDNA, MDA | Illumina HiSeq | IDBA-UD | | PRJEB5265 |
| CruV-353 | 3791 | 32.1% | Standard | Ambisense | TACTATTAC | | River (NZ) | | vDNA, MDA | Illumina HiSeq | metaSPAdes | | MT263612 |
| CruV-354 | 3800 | 44.8% | Standard | Ambisense | TACTATTAC | | unnamed Arctic pond (78°02.935′N; 13°41.973′E) | Aguirre de Cárcer et al., 2015 | vDNA, MDA | Illumina HiSeq | IDBA-UD | | PRJEB5265 |
| CruV-355 | 3807 | 43.2% | Standard | Unisense | TATTATTAC | | unnamed Arctic pond (78°02.935′N; 13°41.973′E) | Aguirre de Cárcer et al., 2015 | vDNA, MDA | Illumina HiSeq | IDBA-UD | | PRJEB5265 |
| CruV-356 | 3823 | 47.4% | Standard | Unisense | CAGTATTAC | | unnamed Arctic pond (78°02.935′N; 13°41.973′E) | Aguirre de Cárcer et al., 2015 | vDNA, MDA | Illumina HiSeq | IDBA-UD | | PRJEB5265 |
| CruV-357 | 3828 | 38.1% | Standard | Ambisense | CAGTATTAC | | Lake Aydat (45°39'52.859''N; 2°59'11.943''E) surface water | | vDNA, MDA | Illumina HiSeq | IDBA-UD | | MT478489 |
| CruV-358 | 3835 | 41.7% | Standard | Ambisense | CAGTATTAC | | Lake Aydat (45°39'52.859''N; 2°59'11.943''E) surface water | | vDNA, MDA | Illumina HiSeq | IDBA-UD | | MT478488 |
| CruV-359 | 3848 | 31.5% | Standard | Ambisense | TAATGATAA | Cp-based clusters | River (NZ) | | vDNA, MDA | Illumina HiSeq | metaSPAdes | | MT263613 |
| CruV-360 | 3852 | 41.8% | Standard | Ambisense | #N/A | | Lake Tunsjøen (78°03.375′N; 13°40.313′E) | Aguirre de Cárcer et al., 2015 | vDNA, MDA | Illumina HiSeq | IDBA-UD | | PRJEB5265 |
| CruV-361 | 3864 | 45.3% | Standard | Ambisense | #N/A | Rep-based clusters | Lake Tunsjøen (78°03.375′N; 13°40.313′E) | Aguirre de Cárcer et al., 2015 | vDNA, MDA | Illumina HiSeq | IDBA-UD | | PRJEB5265 |
| CruV-362 | 3866 | 37.3% | Standard | Ambisense | TAAAATAAA | | River (NZ) | | vDNA, MDA | Illumina HiSeq | metaSPAdes | | MT263614 |
| CruV-363 | 3875 | 33.0% | Standard | Ambisense | TAGTATTAC | | unnamed Arctic pond (78°02.935′N; 13°41.973′E) | Aguirre de Cárcer et al., 2015 | vDNA, MDA | Illumina HiSeq | IDBA-UD | | PRJEB5265 |
| CruV-364 | 3876 | 49.6% | Standard | Ambisense | #N/A | | River (NZ) | | vDNA, MDA | Illumina HiSeq | metaSPAdes | | MT263615 |
| CruV-365 | 3879 | 45.9% | Standard | Ambisense | TATTTATAC / TATAAATAG / CAGTGTTAC | | unnamed Arctic pond (78°02.935′N; 13°41.973′E) | Aguirre de Cárcer et al., 2015 | vDNA, MDA | Illumina HiSeq | IDBA-UD | | PRJEB5265 |
| CruV-366 | 3882 | 48.8% | Standard | Ambisense | #N/A | | Lake Tunsjøen (78°03.375′N; 13°40.313′E) | Aguirre de Cárcer et al., 2015 | vDNA, MDA | Illumina HiSeq | IDBA-UD | | PRJEB5265 |
| CruV-367 | 3899 | 44.1% | Standard | Ambisense | #N/A | Cp-based clusters | Delisea pulchra microbial communities from Sydney, Australia, affected by bleaching disease | Zozaya-Valdés et al., 2017 | eDNA | Illumina HiSeq | IDBA-UD + SOAPdenovo + GAA | | Gp0060493 (GOLD) |
| CruV-368 | 3908 | 48.9% | Standard | Ambisense | TATTATTAC | | Lake Aydat (45°39'52.859''N; 2°59'11.943''E) surface water | | vDNA, MDA | Illumina HiSeq | IDBA-UD | | MT478487 |
| CruV-369 | 3929 | 49.2% | Standard | Ambisense | CATTATTAC | | Lake Nordammen (78°38.279′N; 16°44.025′E) | Aguirre de Cárcer et al., 2015 | vDNA, MDA | Illumina HiSeq | IDBA-UD | | PRJEB5265 |
| CruV-370 | 3948 | 47.9% | Standard | Ambisense | AACTATTAC | | unnamed Arctic pond (78°02.935′N; 13°41.973′E) | Aguirre de Cárcer et al., 2015 | vDNA, MDA | Illumina HiSeq | IDBA-UD | Spliced Rep | PRJEB5265 |
| CruV-371 | 3970 | 51.1% | Standard | Unisense | TAATACTAA | | River (NZ) | | vDNA, MDA | Illumina HiSeq | metaSPAdes | | MT263616 |
| CruV-372 | 3974 | 51.0% | Standard | Unisense | #N/A | Rep-based clusters | River (NZ) | | vDNA, MDA | Illumina HiSeq | metaSPAdes | | MT263617 |
| CruV-373 | 3981 | 49.2% | Standard | Ambisense | TAAAGATAC / TATTTCAAG | Rep-based clusters | River (NZ) | | vDNA, MDA | Illumina HiSeq | metaSPAdes | | MT263618 |
| CruV-374 | 3994 | 46.6% | Standard | Unisense | #N/A | | unnamed Arctic pond (78°02.935′N; 13°41.973′E) | Aguirre de Cárcer et al., 2015 | vDNA, MDA | Illumina HiSeq | IDBA-UD | | PRJEB5265 |
| CruV-375 | 3998 | 34.1% | Standard | Ambisense | TATTATAAC | | Lake Nordammen (78°38.279′N; 16°44.025′E) | Aguirre de Cárcer et al., 2015 | vDNA, MDA | Illumina HiSeq | IDBA-UD | | PRJEB5265 |
| CruV-376 | 4026 | 50.1% | Standard | Ambisense | TATAGATAG / TATTAAAAT | | Lake Aydat (45°39'52.859''N; 2°59'11.943''E) surface water | | vDNA, MDA | Illumina HiSeq | IDBA-UD | | MT478486 |
| CruV-377 | 4037 | 40.2% | Standard | Ambisense | TAATATTAC | Rep-based clusters | unnamed Arctic pond (78°02.935′N; 13°41.973′E) | Aguirre de Cárcer et al., 2015 | vDNA, MDA | Illumina HiSeq | IDBA-UD | | PRJEB5265 |
| CruV-378 | 4042 | 45.1% | Standard | Unisense | #N/A | Rep-based clusters | Lake Tunsjøen (78°03.375′N; 13°40.313′E) | Aguirre de Cárcer et al., 2015 | vDNA, MDA | Illumina HiSeq | IDBA-UD | | PRJEB5265 |
| CruV-379 | 4049 | 54.0% | Standard | Unisense | #N/A | Rep-based clusters | unnamed Arctic pond (78°02.935′N; 13°41.973′E) | Aguirre de Cárcer et al., 2015 | vDNA, MDA | Illumina HiSeq | IDBA-UD | | PRJEB5265 |
| CruV-380 | 4056 | 45.6% | Standard | Ambisense | CAGTATTAC | | Tern feces (Canada) | | vDNA, MDA | Illumina HiSeq | metaSPAdes | | MT263619 |
| CruV-381 | 4057 | 35.2% | Standard | Unisense | TAGTATTAC | | Lake Tunsjøen (78°03.375′N; 13°40.313′E) | Aguirre de Cárcer et al., 2015 | vDNA, MDA | Illumina HiSeq | IDBA-UD | | PRJEB5265 |
| CruV-382 | 4058 | 43.5% | Standard | Ambisense | #N/A | | Lake Tenndammen (78°06.118′N; 15°02.024′E) | Aguirre de Cárcer et al., 2015 | vDNA, MDA | Illumina HiSeq | IDBA-UD | | PRJEB5265 |
| CruV-383 | 4064 | 42.8% | Standard | Ambisense | #N/A | | unnamed Arctic pond (78°02.935′N; 13°41.973′E) | Aguirre de Cárcer et al., 2015 | vDNA, MDA | Illumina HiSeq | IDBA-UD | | PRJEB5265 |
| CruV-384 | 4065 | 48.1% | Standard | Unisense | TATATAAAA | | Sewage Oxydation Pond (NZ) | | vDNA, MDA | Illumina HiSeq | metaSPAdes | | MT263620 |
| CruV-385 | 4067 | 47.9% | Standard | Unisense | #N/A | | Lake Pavin (45°29'45.11''N; 2°53'14.60''E), sampling depth = 22 meters | | vDNA, MDA | Illumina HiSeq | IDBA-UD | | MT478485 |
| CruV-386 | 4072 | 49.2% | Standard | Unisense | TATAACAAC | | Lake Nordammen (78°38.279′N; 16°44.025′E) | Aguirre de Cárcer et al., 2015 | vDNA, MDA | Illumina HiSeq | IDBA-UD | | PRJEB5265 |
| CruV-387 | 4076 | 50.8% | Standard | Unisense | CAGTATTAC | Rep-based clusters | Chiromindae (NZ) | | vDNA, MDA | Illumina HiSeq | metaSPAdes | | MT263621 |
| CruV-388 | 4077 | 34.9% | Standard | Ambisense | TAGTATTAC | | Lake Nordammen (78°38.279′N; 16°44.025′E) | Aguirre de Cárcer et al., 2015 | vDNA, MDA | Illumina HiSeq | IDBA-UD | | PRJEB5265 |
| CruV-389 | 4087 | 45.4% | Standard | Ambisense | TAGTATTAC | | River (NZ) | | vDNA, MDA | Illumina HiSeq | metaSPAdes | | MT263622 |
| CruV-390 | 4094 | 50.5% | Standard | Ambisense | TAGTATTAC | | Lake Nordammen (78°38.279′N; 16°44.025′E) | Aguirre de Cárcer et al., 2015 | vDNA, MDA | Illumina HiSeq | IDBA-UD | | PRJEB5265 |

| Name | Length | %GC | Genetic Code | Genome organization | ori (nona) | Sequences Subset | Location | Publication | Source | Sequencing | Assembly software | Notes | Accession number* |
|---|---|---|---|---|---|---|---|---|---|---|---|---|---|
| CruV-391 | 4115 | 49.1% | Standard | Ambisense | #N/A | | Lake Aydat (45°39'52.859''N; 2°59'11.943''E) surface water | | vDNA, MDA | Illumina HiSeq | IDBA-UD | | MT478484 |
| CruV-392 | 4132 | 45.8% | Standard | Ambisense | CATTAATAT | | River (NZ) | | vDNA, MDA | Illumina HiSeq | metaSPAdes | | MT263623 |
| CruV-393 | 4134 | 53.9% | Standard | Unisense | #N/A | | Lake Aydat (45°39'52.859''N; 2°59'11.943''E) surface water | | vDNA, MDA | Illumina HiSeq | IDBA-UD | | MT478483 |
| CruV-394 | 4137 | 45.6% | Standard | Ambisense | TAGTATTAC | Cp-based clusters | unnamed Arctic pond (78°02.935′N; 13°41.973′E) | Aguirre de Cárcer et al., 2015 | vDNA, MDA | Illumina HiSeq | IDBA-UD | | PRJEB5265 |
| CruV-395 | 4141 | 42.8% | Standard | Ambisense | #N/A | | Lake Nordammen (78°38.279′N; 16°44.025′E) | Aguirre de Cárcer et al., 2015 | vDNA, MDA | Illumina HiSeq | IDBA-UD | | PRJEB5265 |
| CruV-396 | 4153 | 49.7% | Standard | Unisense | #N/A | | unnamed Arctic pond (78°02.935′N; 13°41.973′E) | Aguirre de Cárcer et al., 2015 | vDNA, MDA | Illumina HiSeq | IDBA-UD | | PRJEB5265 |
| CruV-397 | 4158 | 42.5% | Standard | Unisense | TAGTATTAC | | Lake Tunsjøen (78°03.375′N; 13°40.313′E) | Aguirre de Cárcer et al., 2015 | vDNA, MDA | Illumina HiSeq | IDBA-UD | | PRJEB5265 |
| CruV-398 | 4169 | 40.9% | Standard | Ambisense | TATTATTAG / TAATAATAG | | Lake Aydat (45°39'52.859''N; 2°59'11.943''E) surface water | | vDNA, MDA | Illumina HiSeq | IDBA-UD | | MT478482 |
| CruV-399 | 4174 | 47.3% | Standard | Ambisense | CAGTATTAC | Rep-based clusters | unnamed Arctic pond (78°02.935′N; 13°41.973′E) | Aguirre de Cárcer et al., 2015 | vDNA, MDA | Illumina HiSeq | IDBA-UD | | PRJEB5265 |
| CruV-400 | 4184 | 46.9% | Standard | Unisense | #N/A | | Lake Aydat (45°39'52.859''N; 2°59'11.943''E) surface water | | vDNA, MDA | Illumina HiSeq | IDBA-UD | | MT478481 |
| CruV-401 | 4185 | 42.3% | Standard | Ambisense | TAATAGTAA / CAATACTAA / AATTAATAT | | unnamed Arctic pond (78°02.935′N; 13°41.973′E) | Aguirre de Cárcer et al., 2015 | vDNA, MDA | Illumina HiSeq | IDBA-UD | | PRJEB5265 |
| CruV-402 | 4188 | 39.5% | Standard | Ambisense | TATATTTAT | | River (NZ) | | vDNA, MDA | Illumina HiSeq | metaSPAdes | | MT263624 |
| CruV-403 | 4191 | 50.1% | Standard | Ambisense | TAGTATTAC | | unnamed Arctic pond (78°02.935′N; 13°41.973′E) | Aguirre de Cárcer et al., 2015 | vDNA, MDA | Illumina HiSeq | IDBA-UD | | PRJEB5265 |
| CruV-404 | 4197 | 42.4% | Standard | Unisense | AAATAATAC | | Lake Nordammen (78°38.279′N; 16°44.025′E) | Aguirre de Cárcer et al., 2015 | vDNA, MDA | Illumina HiSeq | IDBA-UD | | PRJEB5265 |
| CruV-405 | 4209 | 37.6% | Standard | Unisense | AATTAGTAA | | Lake Tunsjøen (78°03.375′N; 13°40.313′E) | Aguirre de Cárcer et al., 2015 | vDNA, MDA | Illumina HiSeq | IDBA-UD | | PRJEB5265 |
| CruV-406 | 4212 | 45.8% | Standard | Ambisense | #N/A | | River (NZ) | | vDNA, MDA | Illumina HiSeq | metaSPAdes | | MT263625 |
| CruV-407 | 4212 | 45.8% | Standard | Ambisense | TAGTATTAC | | River (NZ) | | vDNA, MDA | Illumina HiSeq | metaSPAdes | | MT263626 |
| CruV-408 | 4213 | 35.9% | Standard | Ambisense | #N/A | | Lake Aydat (45°39'52.859''N; 2°59'11.943''E) surface water | | vDNA, MDA | Illumina HiSeq | IDBA-UD | | MT478480 |
| CruV-409 | 4223 | 39.4% | Standard | Ambisense | CAATACTAT | | River sediments (NZ) | | vDNA, MDA | Illumina HiSeq | metaSPAdes | | MT263627 |
| CruV-410 | 4227 | 39.4% | Standard | Ambisense | #N/A | | River sediments (NZ) | | vDNA, MDA | Illumina HiSeq | metaSPAdes | | MT263628 |
| CruV-411 | 4227 | 38.3% | Standard | Ambisense | TAATATTAC | Cp-based clusters | River (NZ) | | vDNA, MDA | Illumina HiSeq | metaSPAdes | | MT263629 |
| CruV-412 | 4231 | 53.8% | Standard | Unisense | TAATATTAC | | Lake Pavin (45°29'45.11''N; 2°53'14.60''E), sampling depth = 22 meters | | vDNA, MDA | Illumina HiSeq | IDBA-UD | | MT478479 |
| CruV-413 | 4234 | 41.9% | Standard | Unisense | #N/A | | unnamed Arctic pond (78°02.935′N; 13°41973′E) | Aguirre de Cárcer et al., 2015 | vDNA, MDA | Illumina HiSeq | IDBA-UD | Spliced Rep | PRJEB5265 |
| CruV-414 | 4237 | 48.4% | Standard | Unisense | #N/A | | Lake Tunsjøen (78°03.375′N; 13°40.313′E) | Aguirre de Cárcer et al., 2015 | vDNA, MDA | Illumina HiSeq | IDBA-UD | | PRJEB5265 |
| CruV-415 | 4241 | 34.0% | Standard | Ambisense | TAATAGTAA / TACTATTAT | | unnamed Arctic pond (78°02.935′N; 13°41.973′E) | Aguirre de Cárcer et al., 2015 | vDNA, MDA | Illumina HiSeq | IDBA-UD | | PRJEB5265 |
| CruV-416 | 4249 | 53.9% | Standard | Unisense | #N/A | | River (NZ) | | vDNA, MDA | Illumina HiSeq | metaSPAdes | | MT263630 |
| CruV-417 | 4268 | 38.4% | Standard | Unisense | TAATGTTAC | | unnamed Arctic pond (78°02.935′N; 13°41.973′E) | Aguirre de Cárcer et al., 2015 | vDNA, MDA | Illumina HiSeq | IDBA-UD | | PRJEB5265 |
| CruV-418 | 4317 | 52.4% | Standard | Ambisense | #N/A | | Lake Aydat (45°39'52.859''N; 2°59'11.943''E) surface water | | vDNA, MDA | Illumina HiSeq | IDBA-UD | | MT478478 |
| CruV-419 | 4319 | 53.9% | Standard | Unisense | #N/A | | Sewage Oxydation Pond (NZ) | | vDNA, MDA | Illumina HiSeq | metaSPAdes | | MT263631 |
| CruV-420 | 4320 | 35.5% | Standard | Unisense | #N/A | | unnamed Arctic pond (78°02.935′N; 13°41.973′E) | Aguirre de Cárcer et al., 2015 | vDNA, MDA | Illumina HiSeq | IDBA-UD | Double CP | PRJEB5265 |
| CruV-421 | 4329 | 38.4% | Standard | Unisense | AAATAGTAT | Rep-based clusters | unnamed Arctic pond (78°02.935′N; 13°41.973′E) | Aguirre de Cárcer et al., 2015 | vDNA, MDA | Illumina HiSeq | IDBA-UD | | PRJEB5265 |
| CruV-422 | 4336 | 44.8% | Standard | Ambisense | CAGTATTAC | Cp-based clusters | unnamed Arctic pond (78°02.935′N; 13°41.973′E) | Aguirre de Cárcer et al., 2015 | vDNA, MDA | Illumina HiSeq | IDBA-UD | | PRJEB5265 |
| CruV-423 | 4344 | 43.9% | Standard | Ambisense | GATTATTAG | Rep-based clusters | unnamed Arctic pond (78°02.935′N; 13°41.973′E) | Aguirre de Cárcer et al., 2015 | vDNA, MDA | Illumina HiSeq | IDBA-UD | | PRJEB5265 |
| CruV-424 | 4348 | 53.2% | Standard | Unisense | #N/A | | unnamed Arctic pond (78°02.935′N; 13°41.973′E) | Aguirre de Cárcer et al., 2015 | vDNA, MDA | Illumina HiSeq | IDBA-UD | | PRJEB5265 |
| CruV-425 | 4361 | 48.4% | Standard | Unisense | TAGTATTAC | | Lake Nordammen (78°38.279′N; 16°44.025′E) | Aguirre de Cárcer et al., 2015 | vDNA, MDA | Illumina HiSeq | IDBA-UD | | PRJEB5265 |
| CruV-426 | 4368 | 39.2% | Standard | Ambisense | #N/A | | unnamed Arctic pond (78°02.935′N; 13°41.973′E) | Aguirre de Cárcer et al., 2015 | vDNA, MDA | Illumina HiSeq | IDBA-UD | Spliced Rep | PRJEB5265 |
| CruV-427 | 4371 | 50.3% | Standard | Unisense | CACTATTAC | | River (NZ) | | vDNA, MDA | Illumina HiSeq | metaSPAdes | | MT263632 |
| CruV-428 | 4379 | 53.2% | Standard | Unisense | AAATATTAA | Rep-based clusters | unnamed Arctic pond (78°02.935′N; 13°41.973′E) | Aguirre de Cárcer et al., 2015 | vDNA, MDA | Illumina HiSeq | IDBA-UD | | PRJEB5265 |
| CruV-429 | 4383 | 50.6% | Standard | Ambisense | #N/A | | River biofilm (NZ) | | vDNA, MDA | Illumina HiSeq | metaSPAdes | | MT263633 |
| CruV-430 | 4383 | 41.1% | Standard | Ambisense | #N/A | Rep-based clusters | unnamed Arctic pond (78°02.935′N; 13°41.973′E) | Aguirre de Cárcer et al., 2015 | vDNA, MDA | Illumina HiSeq | IDBA-UD | | PRJEB5265 |
| CruV-431 | 4384 | 44.8% | Standard | Ambisense | TAGTATTAC | | Lake Nordammen (78°38.279′N; 16°44.025′E) | Aguirre de Cárcer et al., 2015 | vDNA, MDA | Illumina HiSeq | IDBA-UD | | PRJEB5265 |

| Name | Length | %GC | Genetic Code | Genome organization | ori (nona) | Sequences Subset | Location | Publication | Source | Sequencing | Assembly software | Notes | Accession number* |
|---|---|---|---|---|---|---|---|---|---|---|---|---|---|
| CruV-432 | 4420 | 45.0% | Standard | Unisense | #N/A | Cp-based clusters | Sewage Oxydation Pond (NZ) | | vDNA, MDA | Illumina HiSeq | metaSPAdes | | MT263634 |
| CruV-433 | 4422 | 52.1% | Standard | Unisense | CACTAATAC | | Lake Aydat (45°39'52.859''N; 2°59'11.943''E) surface water | | vDNA, MDA | Illumina HiSeq | IDBA-UD | | MT478477 |
| CruV-434 | 4427 | 40.5% | Standard | Unisense | TAATTTTAC | | Lake Tunsjøen (78°03.375′N; 13°40.313′E) | Aguirre de Cárcer et al., 2015 | vDNA, MDA | Illumina HiSeq | IDBA-UD | Spliced Rep; Capsid contains an endomucin domain | PRJEB5265 |
| CruV-435 | 4428 | 39.3% | Standard | Ambisense | #N/A | Rep-based clusters | unnamed Arctic pond (78°02.935′N; 13°41.973′E) | Aguirre de Cárcer et al., 2015 | vDNA, MDA | Illumina HiSeq | IDBA-UD | | PRJEB5265 |
| CruCGE-436 | 4430 | 40.4% | Standard | Unisense | CATTAATAT | | | Aguirre de Cárcer et al., 2015 | vDNA, MDA | Illumina HiSeq | IDBA-UD | | PRJEB5265 |
| CruV-437 | 4436 | 46.7% | Standard | Unisense | #N/A | | unnamed Arctic pond (78°02.935′N; 13°41.973′E) | Aguirre de Cárcer et al., 2015 | vDNA, MDA | Illumina HiSeq | IDBA-UD | | PRJEB5265 |
| CruV-438 | 4441 | 40.2% | Standard | Ambisense | AAATAATAT | | Soil (NZ) | | vDNA, MDA | Illumina HiSeq | metaSPAdes | | MT263635 |
| CruV-439 | 4447 | 44.8% | Standard | Unisense | GATTACTAT | Rep-based clusters | unnamed Arctic pond (78°02.935′N; 13°41.973′E) | Aguirre de Cárcer et al., 2015 | vDNA, MDA | Illumina HiSeq | IDBA-UD | | PRJEB5265 |
| CruV-440 | 4448 | 50.4% | Standard | Unisense | #N/A | | unnamed Arctic pond (78°02.935′N; 13°41.973′E) | Aguirre de Cárcer et al., 2015 | vDNA, MDA | Illumina HiSeq | IDBA-UD | | PRJEB5265 |
| CruV-441 | 4451 | 46.1% | Standard | Ambisense | #N/A | | unnamed Arctic pond (78°02.935′N; 13°41.973′E) | Aguirre de Cárcer et al., 2015 | vDNA, MDA | Illumina HiSeq | IDBA-UD | | PRJEB5265 |
| CruV-442 | 4451 | 43.4% | Standard | Ambisense | TAGTATTAC | | unnamed Arctic pond (78°02.935′N; 13°41.973′E) | Aguirre de Cárcer et al., 2015 | vDNA, MDA | Illumina HiSeq | IDBA-UD | | PRJEB5265 |
| CruV-443 | 4452 | 37.1% | Standard | Ambisense | #N/A | | Lake Nordammen (78°38.279′N; 16°44.025′E) | Aguirre de Cárcer et al., 2015 | vDNA, MDA | Illumina HiSeq | IDBA-UD | | PRJEB5265 |
| CruV-444 | 4457 | 38.5% | Standard | Ambisense | TAGTATTAC | | unnamed Arctic pond (78°02.935′N; 13°41.973′E) | Aguirre de Cárcer et al., 2015 | vDNA, MDA | Illumina HiSeq | IDBA-UD | | PRJEB5265 |
| CruV-445 | 4457 | 40.3% | Standard | Ambisense | AAATACTAA | | Lake Nordammen (78°38.279′N; 16°44.025′E) | Aguirre de Cárcer et al., 2015 | vDNA, MDA | Illumina HiSeq | IDBA-UD | | PRJEB5265 |
| CruV-446 | 4459 | 44.7% | Standard | Unisense | #N/A | Cp-based clusters | Lake Tunsjøen (78°03.375′N; 13°40.313′E) | Aguirre de Cárcer et al., 2015 | vDNA, MDA | Illumina HiSeq | IDBA-UD | | PRJEB5265 |
| CruV-447 | 4462 | 48.7% | Standard | Ambisense | AAATATTAT | | unnamed Arctic pond (78°02.935′N; 13°41.973′E) | Aguirre de Cárcer et al., 2015 | vDNA, MDA | Illumina HiSeq | IDBA-UD | | PRJEB5265 |
| CruV-448 | 4464 | 42.7% | Standard | Unisense | #N/A | | Lake Nordammen (78°38.279′N; 16°44.025′E) | Aguirre de Cárcer et al., 2015 | vDNA, MDA | Illumina HiSeq | IDBA-UD | | PRJEB5265 |
| CruV-449 | 4482 | 38.5% | Standard | Unisense | #N/A | | Lake Nordammen (78°38.279′N; 16°44.025′E) | Aguirre de Cárcer et al., 2015 | vDNA, MDA | Illumina HiSeq | IDBA-UD | | PRJEB5265 |
| CruV-450 | 4485 | 44.9% | Standard | Unisense | TAAATAAAC / TAATATAAT | | unnamed Arctic pond (78°02.935′N; 13°41.973′E) | Aguirre de Cárcer et al., 2015 | vDNA, MDA | Illumina HiSeq | IDBA-UD | Spliced Rep | PRJEB5265 |
| CruV-451 | 4487 | 37.7% | Standard | Ambisense | TAGTATTAC | Rep-based clusters | Lake Aydat (45°39'52.859''N; 2°59'11.943''E) surface water | | vDNA, MDA | Illumina HiSeq | IDBA-UD | | MT478476 |
| CruV-452 | 4507 | 43.7% | Standard | Ambisense | TACTAGTAA / GACTAGTAT | Rep-based clusters | unnamed Arctic pond (78°02.935′N; 13°41.973′E) | Aguirre de Cárcer et al., 2015 | vDNA, MDA | Illumina HiSeq | IDBA-UD | | PRJEB5265 |
| CruV-453 | 4520 | 44.9% | Standard | Unisense | TAATACTAG | | Green lipped muscles (NZ) | | vDNA, MDA | Illumina HiSeq | metaSPAdes | | MT263534 |
| CruV-454 | 4520 | 44.8% | Standard | Unisense | TATATAAAT | | Green lipped muscles (NZ) | | vDNA, MDA | Illumina HiSeq | metaSPAdes | | MT263535 |
| CruV-455 | 4520 | 44.8% | Standard | Unisense | | | Green lipped muscles (NZ) | | vDNA, MDA | Illumina HiSeq | metaSPAdes | | MT263536 |
| CruV-456 | 4520 | 44.9% | Standard | Unisense | CAGTATTAC | | Green lipped muscles (NZ) | | vDNA, MDA | Illumina HiSeq | metaSPAdes | | MT263537 |
| CruV-457 | 4520 | 44.2% | Standard | Ambisense | TAGTATTAC | | Lake Pavin (45°29'45.11''N; 2°53'14.60''E), sampling depth = 22 meters | | vDNA, MDA | Illumina HiSeq | IDBA-UD | | MT478475 |
| CruV-458 | 4521 | 50.9% | Standard | Ambisense | TAGTATTAC | | | | vDNA, MDA | Illumina HiSeq | metaSPAdes | Spliced Rep | MT263636 |
| CruV-459 | 4525 | 46.1% | Standard | Unisense | TAGTATTAC | | Lake Tunsjøen (78°03.375′N; 13°40.313′E) | Aguirre de Cárcer et al., 2015 | vDNA, MDA | Illumina HiSeq | IDBA-UD | | PRJEB5265 |
| CruV-460 | 4538 | 44.0% | Standard | Unisense | #N/A | | unnamed Arctic pond (78°02.935′N; 13°41.973′E) | Aguirre de Cárcer et al., 2015 | vDNA, MDA | Illumina HiSeq | IDBA-UD | Spliced Rep | PRJEB5265 |
| CruV-461 | 4542 | 53.6% | Standard | Ambisense | TAGTATTAC | | unnamed Arctic pond (78°02.935′N; 13°41.973′E) | Aguirre de Cárcer et al., 2015 | vDNA, MDA | Illumina HiSeq | IDBA-UD | | PRJEB5265 |
| CruV-462 | 4547 | 49.8% | Standard | Ambisense | TAGTATTAC | | Lake Pavin (45°29'45.11''N; 2°53'14.60''E), sampling depth = 80 meters | | vDNA, MDA | Illumina HiSeq | IDBA-UD | Spliced Rep; CP similar to bufiviruses | MT478474 |
| CruV-463 | 4552 | 45.4% | Standard | Ambisense | TAGTATTAC | Cp-based clusters | Lake Pavin (45°29'45.11''N; 2°53'14.60''E), sampling depth = 22 meters | | vDNA, MDA | Illumina HiSeq | IDBA-UD | | MT478473 |
| CruV-464 | 4560 | 47.3% | Standard | Unisense | AAGTAGTAA | Rep-based clusters | unnamed Arctic pond (78°02.935′N; 13°41.973′E) | Aguirre de Cárcer et al., 2015 | vDNA, MDA | Illumina HiSeq | IDBA-UD | | PRJEB5265 |
| CruV-465 | 4568 | 43.9% | Standard | Unisense | #N/A | | unnamed Arctic pond (78°02.935′N; 13°41.973′E) | Aguirre de Cárcer et al., 2015 | vDNA, MDA | Illumina HiSeq | IDBA-UD | Spliced Rep | PRJEB5265 |
| CruV-466 | 4580 | 37.2% | Standard | Ambisense | AAGTATTAA | Rep-based clusters | unnamed Arctic pond (78°02.935′N; 13°41.973′E) | Aguirre de Cárcer et al., 2015 | vDNA, MDA | Illumina HiSeq | IDBA-UD | | PRJEB5265 |
| CruV-467 | 4588 | 39.5% | Standard | Ambisense | #N/A | | Lake Tunsjøen (78°03.375′N; 13°40.313′E) | Aguirre de Cárcer et al., 2015 | vDNA, MDA | Illumina HiSeq | IDBA-UD | | PRJEB5265 |
| CruV-468 | 4601 | 48.1% | Standard | Ambisense | #N/A | | River (NZ) | | vDNA, MDA | Illumina HiSeq | metaSPAdes | | MT263637 |
| CruV-469 | 4605 | 44.7% | Standard | Ambisense | TAGTATTAC | | unnamed Arctic pond (78°02.935′N; 13°41.973′E) | Aguirre de Cárcer et al., 2015 | vDNA, MDA | Illumina HiSeq | IDBA-UD | | PRJEB5265 |

| Name | Length | %GC | Genetic Code | Genome organization | ori (nona) | Sequences Subset | Location | Publication | Source | Sequencing | Assembly software | Notes | Accession number* |
|---|---|---|---|---|---|---|---|---|---|---|---|---|---|
| CruV-470 | 4619 | 44.6% | Standard | Unisense | #N/A | | Lake Aydat (45°39'52.859''N; 2°59'11.943''E) surface water | | vDNA, MDA | Illumina HiSeq | IDBA-UD | | MT478472 |
| CruCGE-471 | 4650 | 43.8% | Standard | Unisense | CAATATTAC | | River (NZ) | | vDNA, MDA | Illumina HiSeq | metaSPAdes | | MT263539 |
| CruV-472 | 4656 | 49.8% | Standard | Ambisense | #N/A | | Tern feces (Canada) | | vDNA, MDA | Illumina HiSeq | metaSPAdes | | MT263638 |
| CruV-473 | 4661 | 38.7% | Ciliate | Ambisense | #N/A | | unnamed Arctic pond (78°02.935′N; 13°41.973′E) | Aguirre de Cárcer et al., 2015 | vDNA, MDA | Illumina HiSeq | IDBA-UD | | PRJEB5265 |
| CruV-474 | 4673 | 41.1% | Standard | Ambisense | CAGTATTAT | | Odonatan larvae (NZ) | | vDNA, MDA | Illumina HiSeq | metaSPAdes | | MT263639 |
| CruV-475 | 4676 | 52.2% | Standard | Ambisense | #N/A | | Sewage Oxydation Pond (NZ) | | vDNA, MDA | Illumina HiSeq | metaSPAdes | | MT263640 |
| CruV-476 | 4679 | 47.8% | Standard | Ambisense | #N/A | | unnamed Arctic pond (78°02.935′N; 13°41.973′E) | Aguirre de Cárcer et al., 2015 | vDNA, MDA | Illumina HiSeq | IDBA-UD | | PRJEB5265 |
| CruV-477 | 4711 | 37.4% | Standard | Ambisense | #N/A | | unnamed Arctic pond (78°02.935′N; 13°41.973′E) | Aguirre de Cárcer et al., 2015 | vDNA, MDA | Illumina HiSeq | IDBA-UD | | PRJEB5265 |
| CruV-478 | 4712 | 43.0% | Standard | Unisense | #N/A | | unnamed Arctic pond (78°02.935′N; 13°41.973′E) | Aguirre de Cárcer et al., 2015 | vDNA, MDA | Illumina HiSeq | IDBA-UD | | PRJEB5265 |
| CruV-479 | 4714 | 49.1% | Standard | Ambisense | CAGTATTAC | | River (NZ) | | vDNA, MDA | Illumina HiSeq | metaSPAdes | | MT263641 |
| CruV-480 | 4727 | 54.2% | Standard | Unisense | TACTATTAC | Cp-based clusters | River (NZ) | | vDNA, MDA | Illumina HiSeq | metaSPAdes | | MT263642 |
| CruV-481 | 4764 | 43.0% | Standard | Ambisense | #N/A | | Lake Pavin (45°29'45.11''N; 2°53'14.60''E), sampling depth = 8 meters | | vDNA, MDA | Illumina HiSeq | IDBA-UD | | MT478471 |
| CruV-482 | 4765 | 45.7% | Standard | Ambisense | CAGTATTAC | | River (NZ) | | vDNA, MDA | Illumina HiSeq | metaSPAdes | | MT263643 |
| CruV-483 | 4768 | 39.5% | Standard | Unisense | TATTATTAC | | River bank soil (NZ) | | vDNA, MDA | Illumina HiSeq | metaSPAdes | Spliced Rep | MT263644 |
| CruV-484 | 4778 | 45.9% | Standard | Ambisense | AATTACTAG | | Lake Pavin (45°29'45.11''N; 2°53'14.60''E), sampling depth = 22 meters | | vDNA, MDA | Illumina HiSeq | IDBA-UD | | MT478470 |
| CruV-485 | 4778 | 49.3% | Standard | Ambisense | #N/A | | unnamed Arctic pond (78°02.935′N; 13°41.973′E) | Aguirre de Cárcer et al., 2015 | vDNA, MDA | Illumina HiSeq | IDBA-UD | | PRJEB5265 |
| CruV-486 | 4778 | 39.5% | Standard | Ambisense | CACTAATAG | | Lake Pavin (45°29'45.11''N; 2°53'14.60''E), sampling depth = 22 meters | | vDNA, MDA | Illumina HiSeq | IDBA-UD | | MT478469 |
| CruV-487 | 4779 | 38.3% | Standard | Unisense | AAGTATTAC | Cp-based clusters | Lake Tenndammen (78°06.118′N; 15°02.024′E) | Aguirre de Cárcer et al., 2015 | vDNA, MDA | Illumina HiSeq | IDBA-UD | | PRJEB5265 |
| CruV-488 | 4789 | 40.3% | Standard | Ambisense | #N/A | | Lake Nordammen (78°38.279′N; 16°44.025′E) | Aguirre de Cárcer et al., 2015 | vDNA, MDA | Illumina HiSeq | IDBA-UD | | PRJEB5265 |
| CruV-489 | 4817 | 35.3% | Ciliate | Unisense | CAGTACTAC | | Lake Tunsjøen (78°03.375′N; 13°40.313′E) | Aguirre de Cárcer et al., 2015 | vDNA, MDA | Illumina HiSeq | IDBA-UD | | PRJEB5265 |
| CruV-490 | 4826 | 40.7% | Standard | Unisense | #N/A | | unnamed Arctic pond (78°02.935′N; 13°41.973′E) | Aguirre de Cárcer et al., 2015 | vDNA, MDA | Illumina HiSeq | IDBA-UD | Spliced Rep | PRJEB5265 |
| CruV-491 | 4831 | 47.4% | Standard | Unisense | GATTAGTAC | | unnamed Arctic pond (78°02.935′N; 13°41.973′E) | Aguirre de Cárcer et al., 2015 | vDNA, MDA | Illumina HiSeq | IDBA-UD | | PRJEB5265 |
| CruV-492 | 4833 | 37.5% | Standard | Unisense | GATTAATAA | Cp-based clusters | unnamed Arctic pond (78°02.935′N; 13°41.973′E) | Aguirre de Cárcer et al., 2015 | vDNA, MDA | Illumina HiSeq | IDBA-UD | | PRJEB5265 |
| CruV-493 | 4860 | 42.0% | Standard | Unisense | TAATACTAA / GATTAATAT / TAATATTAA | | Lake Nordammen (78°38.279′N; 16°44.025′E) | Aguirre de Cárcer et al., 2015 | vDNA, MDA | Illumina HiSeq | IDBA-UD | | PRJEB5265 |
| CruV-494 | 4862 | 44.0% | Standard | Ambisense | AACTAATAA | | Lake Nordammen (78°38.279′N; 16°44.025′E) | Aguirre de Cárcer et al., 2015 | vDNA, MDA | Illumina HiSeq | IDBA-UD | | PRJEB5265 |
| CruV-495 | 4869 | 45.9% | Standard | Ambisense | #N/A | | Blackfly (NZ) | | vDNA, MDA | Illumina HiSeq | metaSPAdes | | MT263645 |
| CruV-496 | 4874 | 43.6% | Standard | Unisense | #N/A | | unnamed Arctic pond (78°02.935′N; 13°41.973′E) | Aguirre de Cárcer et al., 2015 | vDNA, MDA | Illumina HiSeq | IDBA-UD | | PRJEB5265 |
| CruV-497 | 4881 | 36.0% | Standard | Ambisense | #N/A | | River (NZ) | | vDNA, MDA | Illumina HiSeq | metaSPAdes | Spliced Rep | MT263646 |
| CruV-498 | 4899 | 40.6% | Standard | Unisense | AATTAATAC / GACTAATAT | | Lake Tunsjøen (78°03.375′N; 13°40.313′E) | Aguirre de Cárcer et al., 2015 | vDNA, MDA | Illumina HiSeq | IDBA-UD | | PRJEB5265 |
| CruV-499 | 4920 | 44.5% | Standard | Unisense | #N/A | | Penguin guano (Antarctica) | | vDNA, MDA | Illumina HiSeq | metaSPAdes | Spliced Rep | MT263647 |
| CruV-500 | 4940 | 43.6% | Ciliate | Unisense | TAGTATTAC | | unnamed Arctic pond (78°02.935′N; 13°41.973′E) | Aguirre de Cárcer et al., 2015 | vDNA, MDA | Illumina HiSeq | IDBA-UD | | PRJEB5265 |
| CruV-501 | 4942 | 42.7% | Standard | Unisense | CACTAGTAG | | Bivalve (NZ) | | vDNA, MDA | Illumina HiSeq | metaSPAdes | | MT263648 |
| CruV-502 | 4948 | 50.7% | Ciliate | Ambisense | TACTATTAC | | Brewers Bay, St. Thomas, U.S. Virgin Islands (18°20'34.526''N; 64°58'51.24''W) | Soffer et al., 2014 | vDNA, MDA | Roche 454 | newbler | | N/A |
| CruV-503 | 4955 | 45.6% | Standard | Unisense | #N/A | | Lake Aydat (45°39'52.859''N; 2°59'11.943''E) surface water | | vDNA, MDA | Illumina HiSeq | IDBA-UD | | MT478468 |
| CruV-504 | 4979 | 39.9% | Standard | Unisense | #N/A | | Lake Aydat (45°39'52.859''N; 2°59'11.943''E) surface water | | vDNA, MDA | Illumina HiSeq | IDBA-UD | | MT478467 |
| CruV-505 | 4991 | 47.7% | Standard | Ambisense | #N/A | | River (NZ) | | vDNA, MDA | Illumina HiSeq | metaSPAdes | | MT263649 |
| CruV-506 | 4991 | 47.7% | Standard | Ambisense | #N/A | | River (NZ) | | vDNA, MDA | Illumina HiSeq | metaSPAdes | | MT263650 |
| CruV-507 | 4992 | 42.8% | Standard | Ambisense | TAATATAAG / TATTATAAT / TAAATAAAT | Cp-based clusters | Lake Nordammen (78°38.279′N; 16°44.025′E) | Aguirre de Cárcer et al., 2015 | vDNA, MDA | Illumina HiSeq | IDBA-UD | | PRJEB5265 |
| CruV-508 | 4998 | 49.9% | Standard | Ambisense | TAATATAAG / TATTATAAT / TAAATAAAG | | Lake Aydat (45°39'52.859''N; 2°59'11.943''E) surface water | | vDNA, MDA | Illumina HiSeq | IDBA-UD | Spliced Rep; CP similar to bufiviruses | MT478466 |

| Name | Length | %GC | Genetic Code | Genome organization | ori (nona) | Sequences Subset | Location | Publication | Source | Sequencing | Assembly software | Notes | Accession number* |
|---|---|---|---|---|---|---|---|---|---|---|---|---|---|
| CruV-509 | 5004 | 43.2% | Standard | Unisense | TAATATAAG / TATTATAAT / TAAATAAAT | | River (NZ) | | vDNA, MDA | Illumina HiSeq | metaSPAdes | | MT263651 |
| CruV-510 | 5010 | 38.3% | Standard | Ambisense | TAATATAAG / TATTATAAT / TAAATAAAT | | Lake Aydat (45°39'52.859''N; 2°59'11.943''E) surface water | | vDNA, MDA | Illumina HiSeq | IDBA-UD | Rep is homologous to Smittium Rep | MT478465 |
| CruV-511 | 5014 | 44.1% | Standard | Unisense | CACTACTAG | | unnamed Arctic pond (78°02.935′N; 13°41.973′E) | Aguirre de Cárcer et al., 2015 | vDNA, MDA | Illumina HiSeq | IDBA-UD | | PRJEB5265 |
| CruV-512 | 5039 | 44.1% | Standard | Ambisense | CAGTATTAC | | unnamed Arctic pond (78°02.935′N; 13°41.973′E) | Aguirre de Cárcer et al., 2015 | vDNA, MDA | Illumina HiSeq | IDBA-UD | Spliced Rep | PRJEB5265 |
| CruV-513 | 5051 | 39.3% | Standard | Unisense | TAGTATTAC | | River bank soil (NZ) | | vDNA, MDA | Illumina HiSeq | metaSPAdes | Spliced Rep and CP | MT263652 |
| CruV-514 | 5087 | 40.7% | Ciliate | Ambisense | AACTAATAT | | Gastropods (NZ) | | vDNA, MDA | Illumina HiSeq | metaSPAdes | | MT263653 |
| CruV-515 | 5111 | 51.0% | Standard | Ambisense | #N/A | | Lake Pavin (45°29'45.11''N; 2°53'14.60''E), sampling depth = 80 meters | | vDNA, MDA | Illumina HiSeq | IDBA-UD | Spliced Rep | MT478464 |
| CruV-516 | 5115 | 46.4% | Ciliate | Unisense | CAGTATTAC | | unnamed Arctic pond (78°02.935′N; 13°41.973′E) | Aguirre de Cárcer et al., 2015 | vDNA, MDA | Illumina HiSeq | IDBA-UD | | PRJEB5265 |
| CruV-517 | 5120 | 50.9% | Standard | Ambisense | TAATTCAAG | | Lake Pavin (45°29'45.11''N; 2°53'14.60''E), sampling depth = 22 meters | | vDNA, MDA | Illumina HiSeq | IDBA-UD | Spliced Rep | MT478463 |
| CruV-518 | 5126 | 34.6% | Standard | Unisense | #N/A | | Lake Aydat (45°39'52.859''N; 2°59'11.943''E) surface water | | vDNA, MDA | Illumina HiSeq | IDBA-UD | Spliced Rep and CP | MT478462 |
| CruV-519 | 5164 | 56.6% | Standard | Unisense | #N/A | | Lake Nordammen (78°38.279′N; 16°44.025′E) | Aguirre de Cárcer et al., 2015 | vDNA, MDA | Illumina HiSeq | IDBA-UD | | PRJEB5265 |
| CruV-520 | 5206 | 48.9% | Standard | Unisense | TAATGTAAA | | River (NZ) | | vDNA, MDA | Illumina HiSeq | metaSPAdes | | MT263654 |
| CruV-521 | 5219 | 47.1% | Standard | Ambisense | #N/A | | Lake Aydat (45°39'52.859''N; 2°59'11.943''E) surface water | | vDNA, MDA | Illumina HiSeq | IDBA-UD | | MT478461 |
| CruV-522 | 5225 | 42.2% | Standard | Unisense | CATTAATAT | | Lake Aydat (45°39'52.859''N; 2°59'11.943''E) surface water | | vDNA, MDA | Illumina HiSeq | IDBA-UD | Spliced Rep | MT478460 |
| CruV-523 | 5257 | 50.8% | Standard | Unisense | GATTACTAC | | River (NZ) | | vDNA, MDA | Illumina HiSeq | metaSPAdes | Spliced Rep, low similarity CP | MT263655 |
| CruV-524 | 5271 | 43.6% | Standard | Ambisense | CATTATTAC | Cp-based clusters | Lake Aydat (45°39'52.859''N; 2°59'11.943''E) surface water | | vDNA, MDA | Illumina HiSeq | IDBA-UD | | MT478459 |
| CruV-525 | 5284 | 46.9% | Standard | Ambisense | TAAATCTAC | | Lake Aydat (45°39'52.859''N; 2°59'11.943''E) surface water | | vDNA, MDA | Illumina HiSeq | IDBA-UD | | MT478458 |
| CruV-526 | 5312 | 53.7% | Standard | Unisense | TAGTATTAC | | Lake Nordammen (78°38.279′N; 16°44.025′E) | Aguirre de Cárcer et al., 2015 | vDNA, MDA | Illumina HiSeq | IDBA-UD | Spliced Rep | PRJEB5265 |
| CruV-527 | 5318 | 32.7% | Standard | Ambisense | AAATAATAA | | Soil (NZ) | | vDNA, MDA | Illumina HiSeq | metaSPAdes | | MT263656 |
| CruV-528 | 5370 | 32.7% | Standard | Ambisense | AAGTAATAA | Cp-based clusters | Soil (NZ) | | vDNA, MDA | Illumina HiSeq | metaSPAdes | | MT263657 |
| CruV-529 | 5452 | 46.9% | Standard | Ambisense | TAGTATTAC | | Lake Aydat (45°39'52.859''N; 2°59'11.943''E) surface water | | vDNA, MDA | Illumina HiSeq | IDBA-UD | Spliced Rep | MT478457 |
| CruV-530 | 5549 | 38.4% | Standard | Unisense | #N/A | | Chirominidae (NZ) | | vDNA, MDA | Illumina HiSeq | metaSPAdes | | MT263658 |
| CruV-531 | 5582 | 42.3% | Standard | Ambisense | TAAATTTAA / TAAATTTAT | | Lake Aydat (45°39'52.859''N; 2°59'11.943''E) surface water | | vDNA, MDA | Illumina HiSeq | IDBA-UD | | MT478456 |
| CruV-532 | 5603 | 49.9% | Ciliate | Ambisense | #N/A | | unnamed Arctic pond (78°02.935′N; 13°41.973′E) | Aguirre de Cárcer et al., 2015 | vDNA, MDA | Illumina HiSeq | IDBA-UD | | PRJEB5265 |
| CruCGE-533 | 5688 | 42.9% | Standard | Unisense | CATTAATAT | | Lake Nordammen (78°38.279′N; 16°44.025′E) | Aguirre de Cárcer et al., 2015 | vDNA, MDA | Illumina HiSeq | IDBA-UD | | PRJEB5265 |
| CruV-534 | 5797 | 40.9% | Standard | Ambisense | TATATAAAA | | Soil (NZ) | | vDNA, MDA | Illumina HiSeq | metaSPAdes | | MT263659 |
| CruV-535 | 5803 | 42.7% | Standard | Ambisense | CAGTATTAC | | Lake Aydat (45°39'52.859''N; 2°59'11.943''E) surface water | | vDNA, MDA | Illumina HiSeq | IDBA-UD | | MT478455 |
| CruV-536 | 5856 | 37.3% | Standard | Unisense | TATTGAAAG | | River bank soil (NZ) | | vDNA, MDA | Illumina HiSeq | metaSPAdes | | MT263660 |
| CruV-537 | 6350 | 42.3% | Standard | Unisense | CAGTATTAC | | Lake Aydat (45°39'52.859''N; 2°59'11.943''E) surface water | | vDNA, MDA | Illumina HiSeq | IDBA-UD | Spliced Rep | MT478454 |
| CruV-538 | 6385 | 42.9% | Standard | Unisense | #N/A | Rep-based clusters | River (NZ) | | vDNA, MDA | Illumina HiSeq | metaSPAdes | | MT263661 |
| CruV-539 | 6421 | 39.3% | Standard | Ambisense | TGATATTAC | | River (NZ) | | vDNA, MDA | Illumina HiSeq | metaSPAdes | | MT263662 |
| CruV-540 | 6902 | 40.8% | Standard | Ambisense | #N/A | | River bank soil (NZ) | | vDNA, MDA | Illumina HiSeq | metaSPAdes | | MT263663 |
| CruV-541 | 7947 | 45.1% | Standard | Unisense | CAATATTAC / TAATGTTAT / TAATGAAAT / TAAAGTTAC | | Lake Nordammen (78°38.279′N; 16°44.025′E) | Aguirre de Cárcer et al., 2015 | vDNA, MDA | Illumina HiSeq | IDBA-UD | | PRJEB5265 |


*** GenBank accession numbers are provided for previously unpublished sequences. SRA and JGI's GOLD accession numbers are provided for previously published sequences 461 annotated cruciviral sequences are provided as a GenBank flatfile in Supp. File 1.**
