## Supplementary material for "Unveiling Crucivirus Diversity by Mining Metagenomic Data": Supp. Table 2

### Supplementary Table 2: Viromes analyzed

| Project Name | Virome Code | Habitat Type (code) | Ecosystem Type (text) | Location | GPS | Source | Sequencing Technology | Assembly Software | # crucivirus | Reference |
|---|---|---|---|---|---|---|---|---|---|---|
| Santiago_chemostatCulture | Donor10Day12 | Engineered_Bioreactor | Bioreactor | USA : California | 32.879696 -117.234507 | reads MG-RAST | ion torrent | Newbler | 0 | Santiago-Rodriguez TM. Ly M. Daigneault MC. Brown IH. McDonald JA. Bonilla N. Vercoe EA. Pride DT. Chemostat culture systems support diverse bacteriophage communities from human feces. Microbiome. 2015 Nov 9;3(1):58. doi: 10.1186/s40168-015-0124-3. |
| Santiago_chemostatCulture | Donor10Day16 | Engineered_Bioreactor | Bioreactor | USA : California | 32.879696 -117.234507 | reads MG-RAST | ion torrent | Newbler | 0 | |
| Santiago_chemostatCulture | Donor10Day4 | Engineered_Bioreactor | Bioreactor | USA : California | 32.879696 -117.234507 | reads MG-RAST | ion torrent | Newbler | 0 | |
| Santiago_chemostatCulture | Donor10Day8 | Engineered_Bioreactor | Bioreactor | USA : California | 32.879696 -117.234507 | reads MG-RAST | ion torrent | Newbler | 0 | |
| Santiago_chemostatCulture | Donor10Liq | Engineered_Bioreactor | Bioreactor | USA : California | 32.879696 -117.234507 | reads MG-RAST | ion torrent | Newbler | 0 | |
| Santiago_chemostatCulture | Donor10Stool | Engineered_Bioreactor | Bioreactor | USA : California | 32.879696 -117.234507 | reads MG-RAST | ion torrent | Newbler | 0 | |
| Santiago_chemostatCulture | Donor1Day12 | Engineered_Bioreactor | Bioreactor | USA : California | 32.879696 -117.234507 | reads MG-RAST | ion torrent | Newbler | 0 | |
| Santiago_chemostatCulture | Donor1Day16 | Engineered_Bioreactor | Bioreactor | USA : California | 32.879696 -117.234507 | reads MG-RAST | ion torrent | Newbler | 0 | |
| Santiago_chemostatCulture | Donor1Day4 | Engineered_Bioreactor | Bioreactor | USA : California | 32.879696 -117.234507 | reads MG-RAST | ion torrent | Newbler | 0 | |
| Santiago_chemostatCulture | Donor1Day8 | Engineered_Bioreactor | Bioreactor | USA : California | 32.879696 -117.234507 | reads MG-RAST | ion torrent | Newbler | 0 | |
| Santiago_chemostatCulture | Donor1Liq | Engineered_Bioreactor | Bioreactor | USA : California | 32.879696 -117.234507 | reads MG-RAST | ion torrent | Newbler | 0 | |
| Santiago_chemostatCulture | Donor1Stool | Engineered_Bioreactor | Bioreactor | USA : California | 32.879696 -117.234507 | reads MG-RAST | ion torrent | Newbler | 0 | |
| Santiago_chemostatCulture | Donor2Day12 | Engineered_Bioreactor | Bioreactor | USA : California | 32.879696 -117.234507 | reads MG-RAST | ion torrent | Newbler | 0 | |
| Santiago_chemostatCulture | Donor2Day16 | Engineered_Bioreactor | Bioreactor | USA : California | 32.879696 -117.234507 | reads MG-RAST | ion torrent | Newbler | 0 | |
| Santiago_chemostatCulture | Donor2Day4 | Engineered_Bioreactor | Bioreactor | USA : California | 32.879696 -117.234507 | reads MG-RAST | ion torrent | Newbler | 0 | |
| Santiago_chemostatCulture | Donor2Day8 | Engineered_Bioreactor | Bioreactor | USA : California | 32.879696 -117.234507 | reads MG-RAST | ion torrent | Newbler | 0 | |
| Santiago_chemostatCulture | Donor2Liq | Engineered_Bioreactor | Bioreactor | USA : California | 32.879696 -117.234507 | reads MG-RAST | ion torrent | Newbler | 0 | |
| Santiago_chemostatCulture | Donor2Stool | Engineered_Bioreactor | Bioreactor | USA : California | 32.879696 -117.234507 | reads MG-RAST | ion torrent | Newbler | 0 | |
| Santiago_chemostatCulture | Donor8Day12 | Engineered_Bioreactor | Bioreactor | USA : California | 32.879696 -117.234507 | reads MG-RAST | ion torrent | Newbler | 0 | |
| Santiago_chemostatCulture | Donor8Day18 | Engineered_Bioreactor | Bioreactor | USA : California | 32.879696 -117.234507 | reads MG-RAST | ion torrent | Newbler | 0 | |
| Santiago_chemostatCulture | Donor8Day3 | Engineered_Bioreactor | Bioreactor | USA : California | 32.879696 -117.234507 | reads MG-RAST | ion torrent | Newbler | 0 | |
| Santiago_chemostatCulture | Donor8Day6 | Engineered_Bioreactor | Bioreactor | USA : California | 32.879696 -117.234507 | reads MG-RAST | ion torrent | Newbler | 0 | |
| Santiago_chemostatCulture | Donor8Liq | Engineered_Bioreactor | Bioreactor | USA : California | 32.879696 -117.234507 | reads MG-RAST | ion torrent | Newbler | 0 | |
| Santiago_chemostatCulture | Donor8Stool | Engineered_Bioreactor | Bioreactor | USA : California | 32.879696 -117.234507 | reads MG-RAST | ion torrent | Newbler | 0 | |
| Santiago_chemostatCulture | Donor9Day12 | Engineered_Bioreactor | Bioreactor | USA : California | 32.879696 -117.234507 | reads MG-RAST | ion torrent | Newbler | 0 | |
| Santiago_chemostatCulture | Donor9Day18 | Engineered_Bioreactor | Bioreactor | USA : California | 32.879696 -117.234507 | reads MG-RAST | ion torrent | Newbler | 0 | |
| Santiago_chemostatCulture | Donor9Day3 | Engineered_Bioreactor | Bioreactor | USA : California | 32.879696 -117.234507 | reads MG-RAST | ion torrent | Newbler | 0 | |
| Santiago_chemostatCulture | Donor9Day6 | Engineered_Bioreactor | Bioreactor | USA : California | 32.879696 -117.234507 | reads MG-RAST | ion torrent | Newbler | 0 | |
| Santiago_chemostatCulture | Donor9Liq | Engineered_Bioreactor | Bioreactor | USA : California | 32.879696 -117.234507 | reads MG-RAST | ion torrent | Newbler | 0 | |
| Santiago_chemostatCulture | Donor9Stool | Engineered_Bioreactor | Bioreactor | USA : California | 32.879696 -117.234507 | reads MG-RAST | ion torrent | Newbler | 0 | |
| Park_FermentedFood | Cabbage | Engineered_FoodProduction | Food | Korea : Seoul | 37.528853 126.985222 | reads Metavir | 454 | Newbler | 0 | Park EJ. Kim KH. Abell GC. Kim MS. Roh SW. Bae JW. Metagenomic analysis of the viral communities in fermented foods. Appl Environ Microbiol. 2011 Feb;77(4):1284-91. doi: 10.1128/AEM.01859-10. |
| Park_FermentedFood | Sauerkraut | Engineered_FoodProduction | Food | Korea : Seoul | 37.528853 126.985222 | reads Metavir | 454 | Newbler | 0 | |
| Park_FermentedFood | Shrimp | Engineered_FoodProduction | Food | Korea : Seoul | 37.528853 126.985222 | reads Metavir | 454 | Newbler | 0 | |
| Bellas_CryoconiteArctic | Cryoconite | EnvAquatic_Freshwater | Polar Freshwater | Greenland sea | 67.161028 -50.014639 | contigs Metavir | ? | ? | 0 | Bellas CM. Anesio AM. Barker G. Analysis of virus genomes from glacial environments reveals novel virus groups with unusual host interactions. Front Microbiol. 2015 Jul 3;6:656. doi: 10.3389/fmicb.2015.00656. eCollection 2015. |
| Cai_SubtropicalJiulongRiverEstuary | JRE6750 | EnvAquatic_Freshwater | Tropical Freshwater | China : Jiulong River Estuary | 24.432207 117.930848 | reads Metavir/ contigs Metavir | ? | ? | 0 | Cai L. Zhang R. He Y. Feng X. Jiao N. Metagenomic Analysis of Virioplankton of the Subtropical Jiulong River Estuary. China. Viruses. 2016 Feb 2;8(2). pii: E35. doi: 10.3390/v8020035. |
| Carcer_ArcticFresh | ArctSumIII | EnvAquatic_Freshwater | Polar Freshwater | Borgdammane pond | 78.070900 13.794200 | contigs Metavir | Illumina HiSeq | IDBA_UD | 1 | Aguirre de Cárcer D. López-Bueno A. Pearce DA. Alcamí A. Biodiversity and distribution of polar freshwater DNA viruses. Sci Adv. 2015 Jun 19;1(5):e1400127. eCollection 2015 Jun. PMID:26601189 |
| Carcer_ArcticFresh | IR1 | EnvAquatic_Freshwater | Polar Freshwater | Lake Tunsjøen | 78.056250 13.671883 | contigs Metavir | Illumina HiSeq | IDBA_UD | 34 | |
| Carcer_ArcticFresh | IR2 | EnvAquatic_Freshwater | Polar Freshwater | unnamed Arctic pond | 78.048917 13.699550 | contigs Metavir | Illumina HiSeq | IDBA_UD | 114 | |
| Carcer_ArcticFresh | Lv1 | EnvAquatic_Freshwater | Polar Freshwater | Lake Linnevatnet | 78.064400 13.771800 | contigs Metavir | Illumina HiSeq | IDBA_UD | 1 | |

| Project Name | Virome Code | Habitat Type (code) | Ecosystem Type (text) | Location | GPS | Source | Sequencing Technology | Assembly Software | # crucivirus | Reference |
|---|---|---|---|---|---|---|---|---|---|---|
| Carcer_ArcticFresh | SvL1 | EnvAquatic_Freshwater | Polar Freshwater | Lake Nordammen | 78.637983 16.733750 | contigs Metavir | Illumina HiSeq | IDBA_UD | 40 | |
| Carcer_ArcticFresh | SvL2 | EnvAquatic_Freshwater | Polar Freshwater | Lake Tenndammen | 78.101967 15.033733 | contigs Metavir | Illumina HiSeq | IDBA_UD | 24 | |
| Fancello_Gueltas | ElBerbera | EnvAquatic_Freshwater | Freshwater | Mauritania | 20.03361111 13.85666667 | reads Metavir | 454 | Newbler | 0 | Fancello L. Trape S. Robert C. Boyer M. Popgeorgiev N. Raoult D. Desnues C. Viruses in the desert: a metagenomic survey of viral communities in four perennial ponds of the Mauritanian Sahara. ISME J. 2013 Feb;7(2):359-69. doi: 10.1038/ismej.2012.101. Epub 2012 Oct 4. |
| Fancello_Gueltas | Hamdoun | EnvAquatic_Freshwater | Freshwater | Mauritania | 20.42222222 13.28611111 | reads Metavir | 454 | Newbler | 0 | |
| Fancello_Gueltas | Ilij | EnvAquatic_Freshwater | Freshwater | Mauritania | 20.64611111 13.26944444 | reads Metavir | 454 | Newbler | 0 | |
| Fancello_Gueltas | Molomhar | EnvAquatic_Freshwater | Freshwater | Mauritania | 20.64694444 13.35388889 | reads Metavir | 454 | Newbler | 0 | |
| Ge_EastLakeChina | EastLake210 | EnvAquatic_Freshwater | Freshwater | China : Lake Donghu | 30.557213 114.391886 | contigs MG-RAST | solexa | ? | 0 | Ge X. Wu Y. Wang M. Wang J. Wu L. Yang X. Zhang Y. Shi Z. Viral metagenomics analysis of planktonic viruses in East Lake. Wuhan. China. Virol Sin. 2013 Oct;28(5):280-90. doi: 10.1007/s12250-013-3365-y. Epub 2013 Sep 30. |
| Ge_EastLakeChina | EastLake208 | EnvAquatic_Freshwater | Freshwater | China : Lake Donghu | 30.557213 114.391886 | contigs MG-RAST | solexa | Newbler | 0 | |
| Ge_EastLakeChina | EastLake209 | EnvAquatic_Freshwater | Freshwater | China : Lake Donghu | 30.557213 114.391886 | contigs MG-RAST | solexa | Newbler | 0 | |
| Ge_EastLakeChina | EastLake211 | EnvAquatic_Freshwater | Freshwater | China : Lake Donghu | 30.557213 114.391886 | contigs MG-RAST | solexa | ? | 0 | |
| Green_LakeMatoaka | CrimDell | EnvAquatic_Freshwater | Freshwater | USA : Virginia | 37.267189 -76.725779 | reads Metavir | 454 | Newbler | 0 | Jasmin C. Green. Faraz Rahman. Matthew A. Saxton. Kurt E. Williamson . Metagenomic assessment of viral diversity in Lake Matoaka. a temperate. eutrophic freshwater lake in southeastern Virginia. USA . AME 75:117-128 (2015) - doi:10.3354/ame0175 |
| Green_LakeMatoaka | Matoaka | EnvAquatic_Freshwater | Freshwater | USA : Virginia | 37.267189 -76.725779 | reads Metavir | 454 | Newbler | 0 | |
| Green_LakeMatoaka | Pogonian | EnvAquatic_Freshwater | Freshwater | USA : Virginia | 37.267189 -76.725779 | reads Metavir | 454 | Newbler | 0 | |
| Lopezbueno_RNAantarctic | L2006RNA | EnvAquatic_Freshwater | Polar Freshwater | Antarctica | -62.562386 -60.040952 | reads MG-RAST | 454 | Newbler | 0 | López-Bueno A1. Rastrojo A1. Peiró R1. Arenas M1. Alcamí A1. Ecological connectivity shapes quasispecies structure of RNA viruses in an Antarctic lake. Mol Ecol. 2015 Oct;24(19):4812-4825. doi: 10.1111/mec.13321. Epub 2015 Aug 11. |
| Lopezbueno_RNAantarctic | L2007RNA | EnvAquatic_Freshwater | Polar Freshwater | Antarctica | -62.562386 -60.040952 | reads Metavir | 454 | Newbler | 0 | |
| Lopezbueno_RNAantarctic | L2010RNA | EnvAquatic_Freshwater | Polar Freshwater | Antarctica | -62.562386 -60.040952 | reads Metavir | 454 | Newbler | 0 | |
| Mohiuddin_FreshLakes | 50pOnta3215e | EnvAquatic_Freshwater | Freshwater | Lake Ontario | 43.2247 -79.6183 | reads SRA | Illumina HiSeq2000 | IDBA_UD | 0 | Mohiuddin M. Schellhorn HE . Spatial and temporal dynamics of virus occurrence in two freshwater lakes captured through metagenomic analysis. Front Microbiol. 2015 Sep 15;6:960. doi: 10.3389/fmicb.2015.00960. eCollection 2015. |
| Mohiuddin_FreshLakes | 50pOnta3224v | EnvAquatic_Freshwater | Freshwater | Lake Ontario | 43.2247 -79.6183 | reads SRA | Illumina HiSeq2000 | IDBA_UD | 0 | |
| Mohiuddin_FreshLakes | Lb1Erie3223v | EnvAquatic_Freshwater | Freshwater | Lake Erie | 42.8638 -79.3862 | reads SRA | Illumina HiSeq2000 | IDBA_UD | 0 | |
| Mohiuddin_FreshLakes | Lb2Erie3222v | EnvAquatic_Freshwater | Freshwater | Lake Erie | 42.8698 -79.3982 | reads SRA | Illumina HiSeq2000 | IDBA_UD | 0 | |
| Mohiuddin_FreshLakes | Lb3Erie3221v | EnvAquatic_Freshwater | Freshwater | Lake Erie | 42.8723 -79.4187 | reads SRA | Illumina HiSeq2000 | IDBA_UD | 0 | |
| Mohiuddin_FreshLakes | Lb4Erie3220v | EnvAquatic_Freshwater | Freshwater | Lake Erie | 42.8638 -79.3862 | reads SRA | Illumina HiSeq2000 | IDBA_UD | 0 | |
| Mohiuddin_FreshLakes | LbCEErie2964e | EnvAquatic_Freshwater | Freshwater | Lake Erie | 42.8722 -79.4272 | reads SRA | Illumina HiSeq2000 | IDBA_UD | 0 | |
| Mohiuddin_FreshLakes | LbCEErie3218v | EnvAquatic_Freshwater | Freshwater | Lake Erie | 42.8722 -79.4272 | reads SRA | Illumina HiSeq2000 | IDBA_UD | 0 | |
| Mohiuddin_FreshLakes | LbErie3214e | EnvAquatic_Freshwater | Freshwater | Lake Erie | 42.8638 -79.3862 | reads SRA | Illumina HiSeq2000 | IDBA_UD | 0 | |
| Mohiuddin_FreshLakes | Ls1Onta3504v | EnvAquatic_Freshwater | Freshwater | Lake Ontario | 43.2043 -79.2669 | reads SRA | Illumina HiSeq2000 | IDBA_UD | 0 | |
| Mohiuddin_FreshLakes | Ls2Onta3231v | EnvAquatic_Freshwater | Freshwater | Lake Ontario | 43.2047 -79.266 | reads SRA | Illumina HiSeq2000 | IDBA_UD | 0 | |
| Mohiuddin_FreshLakes | Ls3Onta3230v | EnvAquatic_Freshwater | Freshwater | Lake Ontario | 43.2052 -79.2648 | reads SRA | Illumina HiSeq2000 | IDBA_UD | 0 | |
| Mohiuddin_FreshLakes | Ls4Onta3227v | EnvAquatic_Freshwater | Freshwater | Lake Ontario | 43.2043 -79.2669 | reads SRA | Illumina HiSeq2000 | IDBA_UD | 0 | |
| Mohiuddin_FreshLakes | LsOnta3217e | EnvAquatic_Freshwater | Freshwater | Lake Ontario | 43.2043 -79.2669 | reads SRA | Illumina HiSeq2000 | IDBA_UD | 0 | |
| Mohiuddin_FreshLakes | NbErie3213e | EnvAquatic_Freshwater | Freshwater | Lake Erie | 42.8746 -79.2323 | reads SRA | Illumina HiSeq2000 | IDBA_UD | 0 | |
| Mohiuddin_FreshLakes | NbErie3219v | EnvAquatic_Freshwater | Freshwater | Lake Erie | 42.8746 -79.2323 | reads SRA | Illumina HiSeq2000 | IDBA_UD | 0 | |
| Mohiuddin_FreshLakes | QrOnta3216e | EnvAquatic_Freshwater | Freshwater | Lake Ontario | 43.2579 -79.0687 | reads SRA | Illumina HiSeq2000 | IDBA_UD | 0 | |
| Mohiuddin_FreshLakes | QrOnta3225v | EnvAquatic_Freshwater | Freshwater | Lake Ontario | 43.2579 -79.0687 | reads SRA | Illumina HiSeq2000 | IDBA_UD | 0 | |
| Roux_Caviar | PavinA15 | EnvAquatic_Freshwater | Freshwater | France : Pavin lake | 45.495210 2.888074 | reads OwnSite | Illumina HiSeq2000 | IDBA_UD | 0 | Roux S, Enault F, Ravet V, Pereira O, Sullivan MB. Genomic characteristics and environmental distributions of the uncultivated Far-T4 phages. Front Microbiol. 2015 Mar 16;6:199. doi: 10.3389/fmicb.2015.00199. |
| Roux_Caviar | PavinA22 | EnvAquatic_Freshwater | Freshwater | France : Pavin lake | 45.495210 2.888074 | reads OwnSite | Illumina HiSeq2000 | IDBA_UD | 11 | |
| Roux_Caviar | PavinA4 | EnvAquatic_Freshwater | Freshwater | France : Pavin lake | 45.495210 2.888074 | reads OwnSite | Illumina HiSeq2000 | IDBA_UD | 0 | |
| Roux_Caviar | PavinA8 | EnvAquatic_Freshwater | Freshwater | France : Pavin lake | 45.495210 2.888074 | reads OwnSite | Illumina HiSeq2000 | IDBA_UD | 1 | |
| Roux_Caviar | PavinA80 | EnvAquatic_Freshwater | Freshwater | France : Pavin lake | 45.495210 2.888074 | reads OwnSite | Illumina HiSeq2000 | IDBA_UD | 2 | |
| Roux_Caviar | PavinART15 | EnvAquatic_Freshwater | Freshwater | France : Pavin lake | 45.495210 2.888074 | reads OwnSite | Illumina HiSeq2000 | IDBA_UD | 0 | |
| Roux_Caviar | PavinART22 | EnvAquatic_Freshwater | Freshwater | France : Pavin lake | 45.495210 2.888074 | reads OwnSite | Illumina HiSeq2000 | IDBA_UD | 0 | |
| Roux_Caviar | PavinART4 | EnvAquatic_Freshwater | Freshwater | France : Pavin lake | 45.495210 2.888074 | reads OwnSite | Illumina HiSeq2000 | IDBA_UD | 0 | |

| Project Name | Virome Code | Habitat Type (code) | Ecosystem Type (text) | Location | GPS | Source | Sequencing Technology | Assembly Software | # crucivirus | Reference |
|---|---|---|---|---|---|---|---|---|---|---|
| Roux_Caviar | PavinART8 | EnvAquatic_Freshwater | Freshwater | France : Pavin lake | 45.495210 2.888074 | reads OwnSite | Illumina HiSeq2000 | IDBA_UD | 0 | |
| Roux_Caviar | PavinARt80 | EnvAquatic_Freshwater | Freshwater | France : Pavin lake | 45.495210 2.888074 | reads OwnSite | Illumina HiSeq2000 | IDBA_UD | 0 | |
| Roux_Caviar | virusbaton | EnvAquatic_Freshwater | Freshwater | France : Pavin lake | 45.495210 2.888074 | reads OwnSite | Illumina HiSeq2000 | IDBA_UD | 95 | |
| Skvortsov_LakeLoughNeagh | LoughNeagh | EnvAquatic_Freshwater | Freshwater | UK : Lough Neagh | 54.591791 -6.434395 | reads SRA | Illumina | IDBA_UD | 0 | Skvortsov T. de Leeuwe C. Quinn JP. McGrath JW. Allen CC. McElarney Y. Watson C. Arkhipova K. Lavigne R. Kulakov LA. Metagenomic Characterisation of the Viral Community of Lough Neagh. the Largest Freshwater Lake in Ireland. PLoS One. 2016 Feb 29;11(2):e0150361. doi: 10.1371/journal.pone.0150361. |
| Tseng_SubTropicalFresh | subTropFresh74 | EnvAquatic_Freshwater | Freshwater | Taiwan : Taipei | 24.905627 121.553608 | reads SRA | 454 GS FLX | Newbler | 0 | Tseng CH. Chiang PW. Shiah FK. Chen YL. Liou JR. Hsu TC. Maheswararajah S. Saeed I. Halgamuge S. Tang SL. Microbial and viral metagenomes of a subtropical freshwater reservoir subject to climatic disturbances. ISME J. 2013 Dec;7(12):2374-86. doi: 10.1038/ismej.2013.118. |
| Tseng_SubTropicalFresh | SRR6483 | EnvAquatic_Freshwater | Freshwater | Taiwan : Taipei | 24.905627 121.553608 | reads SRA | 454 GS FLX | Newbler | 0 | |
| Tseng_SubTropicalFresh | subTropFresh11 | EnvAquatic_Freshwater | Freshwater | Taiwan : Taipei | 24.905627 121.553608 | reads SRA | 454 GS FLX | Newbler | 0 | |
| Tseng_SubTropicalFresh | subTropFresh13 | EnvAquatic_Freshwater | Freshwater | Taiwan : Taipei | 24.905627 121.553608 | reads SRA | 454 GS FLX | Newbler | 0 | |
| Tseng_SubTropicalFresh | subTropFresh14 | EnvAquatic_Freshwater | Freshwater | Taiwan : Taipei | 24.905627 121.553608 | reads SRA | 454 GS FLX | Newbler | 0 | |
| Tseng_SubTropicalFresh | subTropFresh73 | EnvAquatic_Freshwater | Freshwater | Taiwan : Taipei | 24.905627 121.553608 | reads SRA | 454 GS FLX | Newbler | 0 | |
| Watkins_LakeMichigan | Mont05Jul13 | EnvAquatic_Freshwater | Freshwater | USA : Michigan | 43.376762 -86.994278 | reads Metavir | ? | IDBA_UD | 0 | Watkins SC. Kuehnle N. Ruggeri CA. Malki K. Bruder K. Elayyan J. Damisch K. Vahora N. O'Malley P. Ruggles-Sage B. Romer Z. Putonti C. Assessment of a metaviromic dataset generated from nearshore Lake Michigan. Marine and Freshwater Research - http://dx.doi.org/10.1071/MF15172 |
| Watkins_LakeMichigan | Mont05Jun13 | EnvAquatic_Freshwater | Freshwater | USA : Michigan | 43.376762 -86.994278 | reads Metavir | ? | IDBA_UD | 0 | |
| Watkins_LakeMichigan | Mont15Jun13 | EnvAquatic_Freshwater | Freshwater | USA : Michigan | 43.376762 -86.994278 | reads Metavir | ? | IDBA_UD | 0 | |
| Watkins_LakeMichigan | Mont25Jul13 | EnvAquatic_Freshwater | Freshwater | USA : Michigan | 43.376762 -86.994278 | reads Metavir | ? | IDBA_UD | 0 | |
| Watkins_LakeMichigan | Mont25Jun13 | EnvAquatic_Freshwater | Freshwater | USA : Michigan | 43.376762 -86.994278 | reads Metavir | ? | IDBA_UD | 0 | |
| Watkins_LakeMichigan | s57th05Jun13 | EnvAquatic_Freshwater | Freshwater | USA : Michigan | 43.376762 -86.994278 | reads Metavir | ? | IDBA_UD | 0 | |
| Watkins_LakeMichigan | s57th14Aug13 | EnvAquatic_Freshwater | Freshwater | USA : Michigan | 43.376762 -86.994278 | reads Metavir | ? | IDBA_UD | 0 | |
| Watkins_LakeMichigan | s57th15Jul13 | EnvAquatic_Freshwater | Freshwater | USA : Michigan | 43.376762 -86.994278 | reads Metavir | ? | IDBA_UD | 0 | |
| Watkins_LakeMichigan | s57th25Jun13 | EnvAquatic_Freshwater | Freshwater | USA : Michigan | 43.376762 -86.994278 | reads Metavir | ? | IDBA_UD | 0 | |
| Chow_SaanichInletBC | Saanich10m | EnvAquatic_Marine | Marine water | Canada : British columbia | 48.591667 -123.506111 | reads Metavir | 2 × 250 bp paired-end reads MiSeq v2.0 : avail on Metavir no qual files | Newbler | 0 | Chow CE. Winget DM. White RA. Hallam SJ. Suttle CA. Combining genomic sequencing methods to explore viral diversity and reveal potential virus-host interactions. Front Microbiol. 2015 Apr 10;6:265. doi: 10.3389/fmicb.2015.00265. eCollection 2015. |
| Chow_SaanichInletBC | Saanich200m | EnvAquatic_Marine | Marine water | Canada : British columbia | 48.591667 -123.506111 | reads Metavir | 2 × 250 bp paired-end reads MiSeq v2.0 : avail on Metavir no qual files | Newbler | 0 | |
| Hurwitz_POV | Dunk | EnvAquatic_Marine | Marine water | Coral sea | -17.933420 146.162830 | reads Metavir | 454 | Newbler | 0 | |
| Hurwitz_POV | Fitzroy | EnvAquatic_Marine | Marine water | Coral sea | -16.928549 145.986021 | reads Metavir | 454 | Newbler | 0 | |
| Hurwitz_POV | LA26A1397 | EnvAquatic_Marine | Marine water | Pacific Ocean | 46.956867 -145.666980 | reads Metavir | 454 | Newbler | 0 | |
| Hurwitz_POV | LA26D1400 | EnvAquatic_Marine | Marine water | Pacific Ocean | 46.956867 -145.666980 | reads Metavir | 454 | Newbler | 0 | |
| Hurwitz_POV | LA26O1398 | EnvAquatic_Marine | Marine water | Pacific Ocean | 46.956867 -145.666980 | reads Metavir | 454 | Newbler | 0 | |
| Hurwitz_POV | LA26S1396 | EnvAquatic_Marine | Marine water | Pacific Ocean | 46.956867 -145.666980 | reads Metavir | 454 | Newbler | 0 | |
| Hurwitz_POV | LF26A1402 | EnvAquatic_Marine | Marine water | Pacific Ocean | 46.956867 -145.666980 | reads Metavir | 454 | Newbler | 0 | |
| Hurwitz_POV | LF26D1404 | EnvAquatic_Marine | Marine water | Pacific Ocean | 46.956867 -145.666980 | reads Metavir | 454 | Newbler | 0 | |
| Hurwitz_POV | LF26O1403 | EnvAquatic_Marine | Marine water | Pacific Ocean | 46.956867 -145.666980 | reads Metavir | 454 | Newbler | 0 | |
| Hurwitz_POV | LF26S1401 | EnvAquatic_Marine | Marine water | Pacific Ocean | 46.956867 -145.666980 | reads Metavir | 454 | Newbler | 0 | |
| Hurwitz_POV | LJ12A1393 | EnvAquatic_Marine | Marine water | Pacific Ocean | 46.956867 -145.666980 | reads Metavir | 454 | Newbler | 0 | |
| Hurwitz_POV | LJ12D1395 | EnvAquatic_Marine | Marine water | Pacific Ocean | 46.956867 -145.666980 | reads Metavir | 454 | Newbler | 0 | |
| Hurwitz_POV | LJ12O1394 | EnvAquatic_Marine | Marine water | Pacific Ocean | 46.956867 -145.666980 | reads Metavir | 454 | Newbler | 0 | |
| Hurwitz_POV | LJ12S1392 | EnvAquatic_Marine | Marine water | Pacific Ocean | 46.956867 -145.666980 | reads Metavir | 454 | Newbler | 0 | |
| Hurwitz_POV | LJ26D1408 | EnvAquatic_Marine | Marine water | Pacific Ocean | 46.956867 -145.666980 | reads Metavir | 454 | Newbler | 0 | |
| Hurwitz_POV | LJ26O1407 | EnvAquatic_Marine | Marine water | Pacific Ocean | 46.956867 -145.666980 | reads Metavir | 454 | Newbler | 0 | Hurwitz BL. Sullivan MB. The Pacific Ocean virome (POV): a marine viral metagenomic dataset and [illegible] |

| Project Name | Virome Code | Habitat Type (code) | Ecosystem Type (text) | Location | GPS | Source | Sequencing Technology | Assembly Software | # crucivirus | Reference |
|---|---|---|---|---|---|---|---|---|---|---|
| Hurwitz_POV | LJ26S1405 | EnvAquatic_Marine | Marine water | Pacific Ocean | 46.956867 -145.666980 | reads Metavir | 454 | Newbler | 0 | associated protein clusters for quantitative viral ecology. PLoS One. 2013;8(2):e57355. doi: 10.1371/ journal.pone.0057355. |
| Hurwitz_POV | LJ4A1410 | EnvAquatic_Marine | Marine water | Pacific Ocean | 46.956867 -145.666980 | reads Metavir | 454 | Newbler | 0 | |
| Hurwitz_POV | LJ4D1415 | EnvAquatic_Marine | Marine water | Pacific Ocean | 46.956867 -145.666980 | reads Metavir | 454 | Newbler | 0 | |
| Hurwitz_POV | LJ4O1414 | EnvAquatic_Marine | Marine water | Pacific Ocean | 46.956867 -145.666980 | reads Metavir | 454 | Newbler | 0 | |
| Hurwitz_POV | LJ4S1409 | EnvAquatic_Marine | Marine water | Pacific Ocean | 46.956867 -145.666980 | reads Metavir | 454 | Newbler | 0 | |
| Hurwitz_POV | M1CS1440 | EnvAquatic_Marine | Marine water | Pacific Ocean | 35.575213 -123.563863 | reads Metavir | 454 | Newbler | 0 | |
| Hurwitz_POV | M2MS1438 | EnvAquatic_Marine | Marine water | Pacific Ocean | 35.575213 -123.563863 | reads Metavir | 454 | Newbler | 0 | |
| Hurwitz_POV | M3MD1439 | EnvAquatic_Marine | Marine water | Pacific Ocean | 35.575213 -123.563863 | reads Metavir | 454 | Newbler | 0 | |
| Hurwitz_POV | M4OS1430 | EnvAquatic_Marine | Marine water | Pacific Ocean | 35.575213 -123.563863 | reads Metavir | 454 | Newbler | 0 | |
| Hurwitz_POV | M5OD1431 | EnvAquatic_Marine | Marine water | Pacific Ocean | 35.575213 -123.563863 | reads Metavir | 454 | Newbler | 0 | |
| Hurwitz_POV | M6O1K1432 | EnvAquatic_Marine | Marine water | Pacific Ocean | 35.575213 -123.563863 | reads Metavir | 454 | Newbler | 0 | |
| Hurwitz_POV | M7O4K1433 | EnvAquatic_Marine | Marine water | Pacific Ocean | 35.575213 -123.563863 | reads Metavir | 454 | Newbler | 0 | |
| Hurwitz_POV | SFCS1442 | EnvAquatic_Marine | Marine water | Pacific Ocean | 32.108588 -118.106031 | reads Metavir | 454 | Newbler | 0 | |
| Hurwitz_POV | SFDS1443 | EnvAquatic_Marine | Marine water | Pacific Ocean | 32.108588 -118.106031 | reads Metavir | 454 | Newbler | 0 | |
| Hurwitz_POV | SFSS1444 | EnvAquatic_Marine | Marine water | Pacific Ocean | 32.108588 -118.106031 | reads Metavir | 454 | Newbler | 0 | |
| Hurwitz_POV | STCS1445 | EnvAquatic_Marine | Marine water | Pacific Ocean | 32.108588 -118.106031 | reads Metavir | 454 | Newbler | 0 | |
| McDaniel_inducedProphage | 030409X2a | EnvAquatic_Marine | Marine water | USA : Florida | 27.694309 -82.636907 | reads Moore | 454 GS FLX pyrosequencer (Roche) | Newbler | 0 | McDaniel LD. Rosario K. Breitbart M. Paul JH. Comparative metagenomics: natural populations of induced prophages demonstrate highly unique. lower diversity viral sequences. Environ Microbiol. 2014 Feb;16(2):570-85. doi: 10.1111/1462-2920.12184. |
| McDaniel_inducedProphage | 030409X2i | EnvAquatic_Marine | Marine water | USA : Florida | 27.694309 -82.636907 | reads Moore | 454 GS FLX pyrosequencer (Roche) | Newbler | 0 | |
| McDaniel_inducedProphage | tampBay | EnvAquatic_Marine | Marine water | USA : Florida | 27.694309 -82.636907 | reads Moore | 454 GS FLX pyrosequencer (Roche) | Newbler | 0 | |
| Williamson_IndianOcean | indOcean1478 | EnvAquatic_Marine | Marine water | Indian Ocean | -8.63 95.38 | reads Metavir | 454 | Newbler | 0 | Williamson SJ. Allen LZ. Lorenzi HA. Fadrosh DW. Brami D. Thiagarajan M. McCrow JP. Tovchigrechko A. Yooseph S. Venter JC. Metagenomic exploration of viruses throughout the Indian Ocean. PLoS One. 2012;7(10):e42047. doi: 10.1371/journal.pone.0042047. |
| Williamson_IndianOcean | indOcean1477 | EnvAquatic_Marine | Marine water | Indian Ocean | -8.15 80.38 | reads Metavir | 454 | Newbler | 0 | |
| Williamson_IndianOcean | indOcean1479 | EnvAquatic_Marine | Marine water | Indian Ocean | -8.14 54.21 | reads Metavir | 454 | Newbler | 0 | |
| Williamson_IndianOcean | indOcean1480 | EnvAquatic_Marine | Marine water | Indian Ocean | -29.55 40.68 | reads Metavir | 454 | Newbler | 0 | |
| Adriaenssens_NamibDesertSaltPans | Gobabeb | EnvAquatic_Saline | Hypersaline | Namibia : Gobabeb | -23.507153 15.07125 | contigs NCBI | MiSeq | CLC 99 % | 0 | Adriaenssens EM. van Zyl LJ. Cowan DA. Trindade MI. Metaviromics of Namib Desert Salt Pans: A Novel Lineage of Haloarchaeal Salterproviruses and a Rich Source of ssDNA Viruses. Viruses. 2016 Jan 8;8(1). pii: E14. doi: 10.3390/v8010014. |
| Adriaenssens_NamibDesertSaltPans | Swakopmund | EnvAquatic_Saline | Hypersaline | Namibia : Dorob national park | -22.484728 14.571683 | contigs NCBI | MiSeq | CLC 99 % | 0 | |
| Adriaenssens_NamibHypoliths | hypolith | EnvAquatic_Saline | Hypersaline | Namibia : Gobabeb | -23.561111 15.041389 | reads SRA | Illumina | IDBA_UD | 0 | Adriaenssens EM. Van Zyl L. De Maayer P. Rubagotti E. Rybicki E. Tuffin M. Cowan DA. Metagenomic analysis of the viral community in Namib Desert hypoliths. Environ Microbiol. 2015 Feb;17(2):480-95. Doi: 10.1111/1462-2920.12528. &&&& Zablocki O1. Adriaenssens EM1. Cowan D2. Diversity and Ecology of Viruses in Hyperarid Desert Soils. Appl Environ Microbiol. 2015 Nov 20;82(3):770-7. doi: 10.1128/AEM.02651-15. |
| GarciaHeredia_SalineFosmids | SalFosmids | EnvAquatic_Saline | Hypersaline | Spain : Santa Pola | 38.191242 -0.615222 | contigs NCBI | ? | ? | 0 | Garcia-Heredia I. Martin-Cuadrado AB. Mojica FJ. Santos F. Mira A. Antón J. Rodriguez-Valera F. Reconstructing viral genomes from the environment using fosmid clones: the case of haloviruses. PLoS One. 2012;7(3):e33802. doi: 10.1371/journal.pone.0033802. |
| Pacton_LagoaVermelhaOrganoMineral | Lagoa | EnvAquatic_Saline | | Brazil | -23.083000 -43.950000 | reads Metavir | 454 | Newbler | 0 | Pacton M. Wacey D. Corinaldesi C. Tangherlini M. Kilburn MR. Gorin GE. Danovaro R. Vasconcelos C. Viruses as new agents of organomineralization in the geological record. Nat Commun. 2014 Jul 3;5:4298. doi: 10.1038/ ncomms5298. |
| Roux_Archevir | P2Saloon | EnvAquatic_Saline | Saline | Senegal | 14.131228 -16.471595 | contigs Metavir | Illumina HiSeq2000 | IDBA_UD | 0 | Roux S. Enault F. Ravet V. Colombet J. Bettarel Y. Auguet JC. Bouvier T. Lucas-Staat S. Vellet A. Prangishivili D. Forterre P. Debroas D. Sime-Ngando T. Analysis of metagenomic data reveals common features of halophilic viral communities across continents. Environ Microbiol. 2015 Oct 16. doi: 10.1111/1462-2920.13084. |
| Roux_Archevir | P5Ngallou | EnvAquatic_Saline | Hypersaline | Senegal | 14.077154 -16.737001 | contigs Metavir | Illumina HiSeq2000 | IDBA_UD | 0 | |
| Roux_Archevir | P6Retba | EnvAquatic_Saline | Hypersaline | Senegal | 14.837910 -17.234274 | contigs Metavir | Illumina HiSeq2000 | IDBA_UD | 0 | |
| Roux_Archevir | P7Ngallou | EnvAquatic_Saline | Hypersaline | Senegal | 14.077154 -16.737001 | contigs Metavir | Illumina HiSeq2000 | IDBA_UD | 0 | |
| Roux_Archevir | P8Ngallou | EnvAquatic_Saline | Hypersaline | Senegal | 14.077154 -16.737001 | contigs Metavir | Illumina HiSeq2000 | IDBA_UD | 0 | |
| Roux_Archevir | P9Ngallou | EnvAquatic_Saline | Hypersaline | Senegal | 14.077154 -16.737001 | contigs Metavir | Illumina HiSeq2000 | IDBA_UD | 0 | |
| Bolduc_YellowstoneHotSpring | NL10X0809 | EnvAquatic_ThermalSprings | Hot spring | USA : Yellowstone national park | 44.753600 -110.723700 | contigs NCBI | 454 GS FLX | Newbler | 0 | |

| Project Name | Virome Code | Habitat Type (code) | Ecosystem Type (text) | Location | GPS | Source | Sequencing Technology | Assembly Software | # crucivirus | Reference |
|---|---|---|---|---|---|---|---|---|---|---|
| Bolduc_YellowstoneHotSpring | NL10X0901 | EnvAquatic_ThermalSprings | Hot spring | USA : Yellowstone national park | 44.753600 -110.723700 | contigs NCBI | 454 GS FLX | Newbler | 0 | Bolduc B. Wirth JF. Mazurie A. Young MJ. Viral assemblage composition in Yellowstone acidic hot springs assessed by network analysis. ISME J. 2015 Oct;9(10):2162-77. doi: 10.1038/ismej.2015.28. |
| Bolduc_YellowstoneHotSpring | NL10X0903 | EnvAquatic_ThermalSprings | Hot spring | USA : Yellowstone national park | 44.753600 -110.723700 | contigs NCBI | 454 GS FLX | Newbler | 0 | |
| Bolduc_YellowstoneHotSpring | NL10X0904 | EnvAquatic_ThermalSprings | Hot spring | USA : Yellowstone national park | 44.753600 -110.723700 | contigs NCBI | 454 GS FLX | Newbler | 0 | |
| Bolduc_YellowstoneHotSpring | NL10X0908 | EnvAquatic_ThermalSprings | Hot spring | USA : Yellowstone national park | 44.753600 -110.723700 | contigs NCBI | 454 GS FLX | Newbler | 0 | |
| Bolduc_YellowstoneHotSpring | NL10X0909 | EnvAquatic_ThermalSprings | Hot spring | USA : Yellowstone national park | 44.753600 -110.723700 | contigs NCBI | 454 GS FLX | Newbler | 0 | |
| Bolduc_YellowstoneHotSpring | NL10X0910 | EnvAquatic_ThermalSprings | Hot spring | USA : Yellowstone national park | 44.753600 -110.723700 | contigs NCBI | 454 GS FLX | Newbler | 0 | |
| Bolduc_YellowstoneHotSpring | NL10X1002 | EnvAquatic_ThermalSprings | Hot spring | USA : Yellowstone national park | 44.753600 -110.723700 | contigs NCBI | 454 GS FLX | Newbler | 0 | |
| Bolduc_YellowstoneHotSpring | NL10X1006 | EnvAquatic_ThermalSprings | Hot spring | USA : Yellowstone national park | 44.753600 -110.723700 | contigs NCBI | 454 GS FLX | Newbler | 0 | |
| Bolduc_YellowstoneHotSpring | NL10X1X3 | EnvAquatic_ThermalSprings | Hot spring | USA : Yellowstone national park | 44.753600 -110.723700 | contigs NCBI | MiSeq | IDBA_UD | 0 | |
| Bolduc_YellowstoneHotSpring | NL10X1X4 | EnvAquatic_ThermalSprings | Hot spring | USA : Yellowstone national park | 44.753600 -110.723700 | contigs NCBI | MiSeq | IDBA_UD | 0 | |
| Bolduc_YellowstoneHotSpring | NL10X1X5 | EnvAquatic_ThermalSprings | Hot spring | USA : Yellowstone national park | 44.753600 -110.723700 | contigs NCBI | MiSeq | IDBA_UD | 0 | |
| Anderson_HydrothermalVents | G2810 | EnvAquatic_Vents | Hydrothermal Vents | Pacific Ocean | 45.999969 -129.999700 | reads Moore | 454 GS FLX | Newbler | 0 | Anderson RE. Brazelton WJ. Baross JA (2011) Using CRISPRs as a metagenomic tool to identify microbial hosts of a diffuse flow hydrothermal vent viral assemblage. FEMS Microbiology Ecology 77(1): 120-133. |
| Appelt_14thCoprolite | Coprolite | EnvTerrestrial_Other | | Belgium : Namur | 50.466667 4.866667 | reads Metavir | 454 GS FLX | IDBA_UD | 0 | Appelt S1. Fancello L. Le Bailly M. Raoult D. Drancourt M. Desnues C. Viruses in a 14th-century coprolite. Appl Environ Microbiol. 2014 May;80(9):2648-55. doi: 10.1128/AEM.03242-13. |
| Reavy_ssDNAIn2Soils | brownEarth | EnvTerrestrial_Soil | soil | Scotland | 55.279472 -3.8906851 | reads MG-RAST | 454 | Newbler | 0 | Reavy B. Swanson MM. Cock PJ. Dawson L. Freitag TE. Singh BK. Torrance L. Mushegian AR. Taliansky M. Distinct circular single-stranded DNA viruses exist in different soil types. Appl Environ Microbiol. 2015 Jun 15;81(12):3934-45. doi: 10.1128/AEM.03878-14. Epub 2015 Apr 3. |
| Reavy_ssDNAIn2Soils | machair | EnvTerrestrial_Soil | soil | Scotland | 56.599975 -6.6453090 | reads MG-RAST | 454 | Newbler | 0 | |
| Zablocki_AntarcticSoils | Antarchypolith | EnvTerrestrial_Soil | Polar soil | Antarctica | -78.093333 163.810000 | contigs Metavir | | CLC | 0 | Zablocki O. van Zyl L. Adriaenssens EM. Rubagotti E. Tuffin M. Cary SC. Cowan D. High-level diversity of tailed phages. eukaryote-associated viruses. and virophage-like elements in the metaviromes of antarctic soils. Appl Environ Microbiol. 2014 Nov;80(22):6888-97. doi: 10.1128/AEM.01525-14. Epub 2014 Aug 29. |
| Zablocki_AntarcticSoils | AntarcSoils | EnvTerrestrial_Soil | Polar soil | Antarctica | -78.093333 163.810000 | contigs Metavir | | CLC | 0 | |
| Carlos_CoralBrazil | Mdecactis | HostAssoc_Cnidaria | Coral | Brazil : Buzios Island | -22.748090 -41.881290 | reads MG-RAST | 454 | Newbler | 0 | Carlos C. Castro DB. Ottoboni LM. Comparative metagenomic analysis of coral microbial communities using a reference-independent approach. PLoS One. 2014 Nov 7;9(11):e111626. doi: 10.1371/journal.pone.0111626. eCollection 2014. |
| Carlos_CoralBrazil | Mhispida | HostAssoc_Cnidaria | Coral | Brazil : Buzios Island | -22.748090 -41.881290 | reads MG-RAST | 454 | Newbler | 0 | |
| Correa_CoralBleaching | coralBleach | HostAssoc_Cnidaria | Coral | Australia : Heron Island | -23.443075 151.908442 | reads ENA | MiSeq | IDBA_UD | 0 | AM Correa. TD Ainsworth. SM Rosales. AR Thurber. CR Butler and RL Vega Thurber. Viral outbreak in corals associated with an in situ bleaching event: atypical herpes-like viruses and a new megavirus infecting Symbiodinium. Front. Microbiol. I doi: 10.3389/fmicb.2016.00127 |
| Soffer_CoralPlague | A5Coral853 | HostAssoc_Cnidaria | Coral | USA : Virgin Islands | 18.342267 -64.984840 | reads Metavir | 454 GS FLX | Newbler | 0 | Soffer N. Brandt ME. Correa AM. Smith TB. Thurber RV. Potential role of viruses in white plague coral disease. ISME J. 2014 Feb;8(2):271-83. doi: 10.1038/ismej.2013.137. Epub 2013 Aug 15. |
| Soffer_CoralPlague | A10Coral879 | HostAssoc_Cnidaria | Coral | USA : Virgin Islands | 18.342267 -64.984840 | reads Metavir | 454 GS FLX | Newbler | 0 | |
| Soffer_CoralPlague | A2Coral851 | HostAssoc_Cnidaria | Coral | USA : Virgin Islands | 18.342267 -64.984840 | reads Metavir | 454 GS FLX | Newbler | 0 | |
| Soffer_CoralPlague | A6Coral854 | HostAssoc_Cnidaria | Coral | USA : Virgin Islands | 18.342267 -64.984840 | reads Metavir | 454 GS FLX | Newbler | 0 | |
| Soffer_CoralPlague | D10Coral886 | HostAssoc_Cnidaria | Coral | USA : Virgin Islands | 18.342267 -64.984840 | reads Metavir | 454 GS FLX | Newbler | 0 | |
| Soffer_CoralPlague | D1Coral880 | HostAssoc_Cnidaria | Coral | USA : Virgin Islands | 18.342267 -64.984840 | reads Metavir | 454 GS FLX | Newbler | 0 | |
| Soffer_CoralPlague | D2Coral881 | HostAssoc_Cnidaria | Coral | USA : Virgin Islands | 18.342267 -64.984840 | reads Metavir | 454 GS FLX | Newbler | 0 | |
| Soffer_CoralPlague | D3Coral882 | HostAssoc_Cnidaria | Coral | USA : Virgin Islands | 18.342267 -64.984840 | reads Metavir | 454 GS FLX | Newbler | 0 | |
| Soffer_CoralPlague | D6Coral884 | HostAssoc_Cnidaria | Coral | USA : Virgin Islands | 18.342267 -64.984840 | reads Metavir | 454 GS FLX | Newbler | 0 | |
| Soffer_CoralPlague | H10Coral892 | HostAssoc_Cnidaria | Coral | USA : Virgin Islands | 18.342267 -64.984840 | reads Metavir | 454 GS FLX | Newbler | 0 | |
| Soffer_CoralPlague | SWdCoral895 | HostAssoc_Cnidaria | Coral | USA : Virgin Islands | 18.342267 -64.984840 | reads Metavir | 454 GS FLX | Newbler | 1 | |
| Soffer_CoralPlague | swHCoral896 | HostAssoc_Cnidaria | Coral | USA : Virgin Islands | 18.342267 -64.984840 | reads Metavir | 454 GS FLX | Newbler | 0 | |
| Soffer_CoralPlague | Ub2Coral894 | HostAssoc_Cnidaria | Coral | USA : Virgin Islands | 18.342267 -64.984840 | reads Metavir | 454 GS FLX | Newbler | 0 | |
| Weynberg_7coral | AtenuisDSISPA | HostAssoc_Cnidaria | Coral | Coral sea | -18.496944 146.596111 | reads SRA | | Newbler | 0 | |
| Weynberg_7coral | AtenuisRSISPA | HostAssoc_Cnidaria | Coral | Coral sea | -18.496944 146.596111 | reads SRA | | Newbler | 0 | |
| Weynberg_7coral | PdamicornisCFM | HostAssoc_Cnidaria | Coral | Coral sea | -18.496944 146.596111 | reads SRA | | Newbler | 0 | Weynberg KD. Wood-Charlson EM. Suttle CA. van Oppen |

| Project Name | Virome Code | Habitat Type (code) | Ecosystem Type (text) | Location | GPS | Source | Sequencing Technology | Assembly Software | # crucivirus | Reference |
|---|---|---|---|---|---|---|---|---|---|---|
| Weynberg_7coral | PdamicornisLN2 | HostAssoc_Cnidaria | Coral | Coral sea | -18.496944 146.596111 | reads SRA | | Newbler | 0 | MJ. Generating viral metagenomes from the coral holobiont. Front Microbiol. 2014 May 7;5:206. doi: 10.3389/fmicb.2014.00206. |
| Weynberg_7coral | PdamicornisLN2S | HostAssoc_Cnidaria | Coral | Coral sea | -18.496944 146.596111 | reads SRA | | Newbler | 0 | |
| Weynberg_7coral | PdamicornisNLN | HostAssoc_Cnidaria | Coral | Coral sea | -18.496944 146.596111 | reads SRA | | Newbler | 0 | |
| Weynberg_7coral | PdamicornisNLNS | HostAssoc_Cnidaria | Coral | Coral sea | -18.496944 146.596111 | reads SRA | | Newbler | 0 | |
| WoodCharlson_CoralsRNA | CoralCtrlRNA | HostAssoc_Cnidaria | Coral | USA : Florida | 24.541370 -81.790761 | reads Metavir | 454 | Newbler | 0 | Wood-Charlson EM. Weynberg KD. Suttle CA. Roux S. van Oppen MJ. Metagenomic characterisation of viral communities in corals: Mining biological signal from methodological noise. Environ Microbiol Rep. 2015 Feb 13. doi: 10.1111/1758-2229.12275. |
| WoodCharlson_CoralsRNA | CoralStressRNA | HostAssoc_Cnidaria | Coral | USA : Florida | 24.541370 -81.790761 | reads Metavir | 454 | Newbler | 0 | |
| Lim_EarlyLifeDynamics | A109167M | HostAssoc_Human Feces | Human feces | USA : Missouri | 38.130000 -90.000000 | reads SRA | 2 x 250 paired-end, MiSeq v2 | Newbler | 0 | |
| Lim_EarlyLifeDynamics | A109167S | HostAssoc_Human Feces | Human feces | USA : Missouri | 38.130000 -90.000000 | reads SRA | 2 x 250 paired-end, MiSeq v2 | Newbler | 0 | |
| Lim_EarlyLifeDynamics | A1120834M | HostAssoc_Human Feces | Human feces | USA : Missouri | 38.130000 -90.000000 | reads SRA | 2 x 250 paired-end, MiSeq v2 | Newbler | 0 | |
| Lim_EarlyLifeDynamics | A1120834S | HostAssoc_Human Feces | Human feces | USA : Missouri | 38.130000 -90.000000 | reads SRA | 2 x 250 paired-end, MiSeq v2 | Newbler | 0 | |
| Lim_EarlyLifeDynamics | A1186364M | HostAssoc_Human Feces | Human feces | USA : Missouri | 38.130000 -90.000000 | reads SRA | 2 x 250 paired-end, MiSeq v2 | Newbler | 0 | |
| Lim_EarlyLifeDynamics | A1186364S | HostAssoc_Human Feces | Human feces | USA : Missouri | 38.130000 -90.000000 | reads SRA | 2 x 250 paired-end, MiSeq v2 | Newbler | 0 | |
| Lim_EarlyLifeDynamics | A1246377M | HostAssoc_Human Feces | Human feces | USA : Missouri | 38.130000 -90.000000 | reads SRA | 2 x 250 paired-end, MiSeq v2 | Newbler | 0 | |
| Lim_EarlyLifeDynamics | A1246377S | HostAssoc_Human Feces | Human feces | USA : Missouri | 38.130000 -90.000000 | reads SRA | 2 x 250 paired-end, MiSeq v2 | Newbler | 0 | |
| Lim_EarlyLifeDynamics | A133483M | HostAssoc_Human Feces | Human feces | USA : Missouri | 38.130000 -90.000000 | reads SRA | 2 x 250 paired-end, MiSeq v2 | Newbler | 0 | |
| Lim_EarlyLifeDynamics | A133483S | HostAssoc_Human Feces | Human feces | USA : Missouri | 38.130000 -90.000000 | reads SRA | 2 x 250 paired-end, MiSeq v2 | Newbler | 0 | |
| Lim_EarlyLifeDynamics | A163707M | HostAssoc_Human Feces | Human feces | USA : Missouri | 38.130000 -90.000000 | reads SRA | 2 x 250 paired-end, MiSeq v2 | Newbler | 0 | |
| Lim_EarlyLifeDynamics | A163707S | HostAssoc_Human Feces | Human feces | USA : Missouri | 38.130000 -90.000000 | reads SRA | 2 x 250 paired-end, MiSeq v2 | Newbler | 0 | |
| Lim_EarlyLifeDynamics | A208584M | HostAssoc_Human Feces | Human feces | USA : Missouri | 38.130000 -90.000000 | reads SRA | 2 x 250 paired-end, MiSeq v2 | Newbler | 0 | |
| Lim_EarlyLifeDynamics | A208584S | HostAssoc_Human Feces | Human feces | USA : Missouri | 38.130000 -90.000000 | reads SRA | 2 x 250 paired-end, MiSeq v2 | Newbler | 0 | |
| Lim_EarlyLifeDynamics | A2120860M | HostAssoc_Human Feces | Human feces | USA : Missouri | 38.130000 -90.000000 | reads SRA | 2 x 250 paired-end, MiSeq v2 | Newbler | 0 | |
| Lim_EarlyLifeDynamics | A2120860S | HostAssoc_Human Feces | Human feces | USA : Missouri | 38.130000 -90.000000 | reads SRA | 2 x 250 paired-end, MiSeq v2 | Newbler | 0 | |
| Lim_EarlyLifeDynamics | A2186574M | HostAssoc_Human Feces | Human feces | USA : Missouri | 38.130000 -90.000000 | reads SRA | 2 x 250 paired-end, MiSeq v2 | Newbler | 0 | |
| Lim_EarlyLifeDynamics | A2186574S | HostAssoc_Human Feces | Human feces | USA : Missouri | 38.130000 -90.000000 | reads SRA | 2 x 250 paired-end, MiSeq v2 | Newbler | 0 | |
| Lim_EarlyLifeDynamics | A2246583M | HostAssoc_Human Feces | Human feces | USA : Missouri | 38.130000 -90.000000 | reads SRA | 2 x 250 paired-end, MiSeq v2 | Newbler | 0 | |
| Lim_EarlyLifeDynamics | A2246583S | HostAssoc_Human Feces | Human feces | USA : Missouri | 38.130000 -90.000000 | reads SRA | 2 x 250 paired-end, MiSeq v2 | Newbler | 0 | |
| Lim_EarlyLifeDynamics | A233467M | HostAssoc_Human Feces | Human feces | USA : Missouri | 38.130000 -90.000000 | reads SRA | 2 x 250 paired-end, MiSeq v2 | Newbler | 0 | |
| Lim_EarlyLifeDynamics | A233467S | HostAssoc_Human Feces | Human feces | USA : Missouri | 38.130000 -90.000000 | reads SRA | 2 x 250 paired-end, MiSeq v2 | Newbler | 0 | |
| Lim_EarlyLifeDynamics | A263710M | HostAssoc_Human Feces | Human feces | USA : Missouri | 38.130000 -90.000000 | reads SRA | 2 x 250 paired-end, MiSeq v2 | Newbler | 0 | |
| Lim_EarlyLifeDynamics | A263710S | HostAssoc_Human Feces | Human feces | USA : Missouri | 38.130000 -90.000000 | reads SRA | 2 x 250 paired-end, MiSeq v2 | Newbler | 0 | |
| Lim_EarlyLifeDynamics | B108199M | HostAssoc_Human Feces | Human feces | USA : Missouri | 38.130000 -90.000000 | reads SRA | 2 x 250 paired-end, MiSeq v2 | Newbler | 0 | |
| Lim_EarlyLifeDynamics | B108199S | HostAssoc_Human Feces | Human feces | USA : Missouri | 38.130000 -90.000000 | reads SRA | 2 x 250 paired-end, MiSeq v2 | Newbler | 0 | |
| Lim_EarlyLifeDynamics | B1123480M | HostAssoc_Human Feces | Human feces | USA : Missouri | 38.130000 -90.000000 | reads SRA | 2 x 250 paired-end, MiSeq v2 | Newbler | 0 | |
| Lim_EarlyLifeDynamics | B1123480S | HostAssoc_Human Feces | Human feces | USA : Missouri | 38.130000 -90.000000 | reads SRA | 2 x 250 paired-end, MiSeq v2 | Newbler | 0 | |
| Lim_EarlyLifeDynamics | B1181568M | HostAssoc_Human Feces | Human feces | USA : Missouri | 38.130000 -90.000000 | reads SRA | 2 x 250 paired-end, MiSeq v2 | Newbler | 0 | |
| Lim_EarlyLifeDynamics | B1181568S | HostAssoc_Human Feces | Human feces | USA : Missouri | 38.130000 -90.000000 | reads SRA | 2 x 250 paired-end, MiSeq v2 | Newbler | 0 | |
| Lim_EarlyLifeDynamics | B1246604M | HostAssoc_Human Feces | Human feces | USA : Missouri | 38.130000 -90.000000 | reads SRA | 2 x 250 paired-end, MiSeq v2 | Newbler | 0 | |
| Lim_EarlyLifeDynamics | B1246604S | HostAssoc_Human Feces | Human feces | USA : Missouri | 38.130000 -90.000000 | reads SRA | 2 x 250 paired-end, MiSeq v2 | Newbler | 0 | |
| Lim_EarlyLifeDynamics | B134708M | HostAssoc_Human Feces | Human feces | USA : Missouri | 38.130000 -90.000000 | reads SRA | 2 x 250 paired-end, MiSeq v2 | Newbler | 0 | |
| Lim_EarlyLifeDynamics | B134708S | HostAssoc_Human Feces | Human feces | USA : Missouri | 38.130000 -90.000000 | reads SRA | 2 x 250 paired-end, MiSeq v2 | Newbler | 0 | |
| Lim_EarlyLifeDynamics | B160912M | HostAssoc_Human Feces | Human feces | USA : Missouri | 38.130000 -90.000000 | reads SRA | 2 x 250 paired-end, MiSeq v2 | Newbler | 0 | |
| Lim_EarlyLifeDynamics | B160912S | HostAssoc_Human Feces | Human feces | USA : Missouri | 38.130000 -90.000000 | reads SRA | 2 x 250 paired-end, MiSeq v2 | Newbler | 0 | |

| Project Name | Virome Code | Habitat Type (code) | Ecosystem Type (text) | Location | GPS | Source | Sequencing Technology | Assembly Software | # crucivirus | Reference |
|---|---|---|---|---|---|---|---|---|---|---|
| Lim_EarlyLifeDynamics | B208680M | HostAssoc_Human Feces | Human feces | USA : Missouri | 38.130000 -90.000000 | reads SRA | 2 × 250 paired-end, MiSeq v2 | Newbler | 0 | Lim ES. Zhou Y. Zhao G. Bauer IK. Droit L. Ndao IM. Warner BB. Tarr PI. Wang D. Holtz LR. Early life dynamics of the human gut virome and bacterial microbiome in infants. Nat Med. 2015 Oct;21(10):1228-34. doi: 10.1038/nm.3950. |
| Lim_EarlyLifeDynamics | B208680S | HostAssoc_Human Feces | Human feces | USA : Missouri | 38.130000 -90.000000 | reads SRA | 2 × 250 paired-end, MiSeq v2 | Newbler | 0 | |
| Lim_EarlyLifeDynamics | B2120802M | HostAssoc_Human Feces | Human feces | USA : Missouri | 38.130000 -90.000000 | reads SRA | 2 × 250 paired-end, MiSeq v2 | Newbler | 0 | |
| Lim_EarlyLifeDynamics | B2120802S | HostAssoc_Human Feces | Human feces | USA : Missouri | 38.130000 -90.000000 | reads SRA | 2 × 250 paired-end, MiSeq v2 | Newbler | 0 | |
| Lim_EarlyLifeDynamics | B2181797M | HostAssoc_Human Feces | Human feces | USA : Missouri | 38.130000 -90.000000 | reads SRA | 2 × 250 paired-end, MiSeq v2 | Newbler | 0 | |
| Lim_EarlyLifeDynamics | B2181797S | HostAssoc_Human Feces | Human feces | USA : Missouri | 38.130000 -90.000000 | reads SRA | 2 × 250 paired-end, MiSeq v2 | Newbler | 0 | |
| Lim_EarlyLifeDynamics | B2241824M | HostAssoc_Human Feces | Human feces | USA : Missouri | 38.130000 -90.000000 | reads SRA | 2 × 250 paired-end, MiSeq v2 | Newbler | 0 | |
| Lim_EarlyLifeDynamics | B2241824S | HostAssoc_Human Feces | Human feces | USA : Missouri | 38.130000 -90.000000 | reads SRA | 2 × 250 paired-end, MiSeq v2 | Newbler | 0 | |
| Lim_EarlyLifeDynamics | B234724M | HostAssoc_Human Feces | Human feces | USA : Missouri | 38.130000 -90.000000 | reads SRA | 2 × 250 paired-end, MiSeq v2 | Newbler | 0 | |
| Lim_EarlyLifeDynamics | B234724S | HostAssoc_Human Feces | Human feces | USA : Missouri | 38.130000 -90.000000 | reads SRA | 2 × 250 paired-end, MiSeq v2 | Newbler | 0 | |
| Lim_EarlyLifeDynamics | B263463M | HostAssoc_Human Feces | Human feces | USA : Missouri | 38.130000 -90.000000 | reads SRA | 2 × 250 paired-end, MiSeq v2 | Newbler | 0 | |
| Lim_EarlyLifeDynamics | B263463S | HostAssoc_Human Feces | Human feces | USA : Missouri | 38.130000 -90.000000 | reads SRA | 2 × 250 paired-end, MiSeq v2 | Newbler | 0 | |
| Lim_EarlyLifeDynamics | C100245M | HostAssoc_Human Feces | Human feces | USA : Missouri | 38.130000 -90.000000 | reads SRA | 2 × 250 paired-end, MiSeq v2 | Newbler | 0 | |
| Lim_EarlyLifeDynamics | C100245S | HostAssoc_Human Feces | Human feces | USA : Missouri | 38.130000 -90.000000 | reads SRA | 2 × 250 paired-end, MiSeq v2 | Newbler | 0 | |
| Lim_EarlyLifeDynamics | C1129721M | HostAssoc_Human Feces | Human feces | USA : Missouri | 38.130000 -90.000000 | reads SRA | 2 × 250 paired-end, MiSeq v2 | Newbler | 0 | |
| Lim_EarlyLifeDynamics | C1129721S | HostAssoc_Human Feces | Human feces | USA : Missouri | 38.130000 -90.000000 | reads SRA | 2 × 250 paired-end, MiSeq v2 | Newbler | 0 | |
| Lim_EarlyLifeDynamics | C1181815M | HostAssoc_Human Feces | Human feces | USA : Missouri | 38.130000 -90.000000 | reads SRA | 2 × 250 paired-end, MiSeq v2 | Newbler | 0 | |
| Lim_EarlyLifeDynamics | C1181815S | HostAssoc_Human Feces | Human feces | USA : Missouri | 38.130000 -90.000000 | reads SRA | 2 × 250 paired-end, MiSeq v2 | Newbler | 0 | |
| Lim_EarlyLifeDynamics | C1246608M | HostAssoc_Human Feces | Human feces | USA : Missouri | 38.130000 -90.000000 | reads SRA | 2 × 250 paired-end, MiSeq v2 | Newbler | 0 | |
| Lim_EarlyLifeDynamics | C1246608S | HostAssoc_Human Feces | Human feces | USA : Missouri | 38.130000 -90.000000 | reads SRA | 2 × 250 paired-end, MiSeq v2 | Newbler | 0 | |
| Lim_EarlyLifeDynamics | C130019M | HostAssoc_Human Feces | Human feces | USA : Missouri | 38.130000 -90.000000 | reads SRA | 2 × 250 paired-end, MiSeq v2 | Newbler | 0 | |
| Lim_EarlyLifeDynamics | C130019S | HostAssoc_Human Feces | Human feces | USA : Missouri | 38.130000 -90.000000 | reads SRA | 2 × 250 paired-end, MiSeq v2 | Newbler | 0 | |
| Lim_EarlyLifeDynamics | C161171M | HostAssoc_Human Feces | Human feces | USA : Missouri | 38.130000 -90.000000 | reads SRA | 2 × 250 paired-end, MiSeq v2 | Newbler | 0 | |
| Lim_EarlyLifeDynamics | C161171S | HostAssoc_Human Feces | Human feces | USA : Missouri | 38.130000 -90.000000 | reads SRA | 2 × 250 paired-end, MiSeq v2 | Newbler | 0 | |
| Lim_EarlyLifeDynamics | C200238M | HostAssoc_Human Feces | Human feces | USA : Missouri | 38.130000 -90.000000 | reads SRA | 2 × 250 paired-end, MiSeq v2 | Newbler | 0 | |
| Lim_EarlyLifeDynamics | C2129723M | HostAssoc_Human Feces | Human feces | USA : Missouri | 38.130000 -90.000000 | reads SRA | 2 × 250 paired-end, MiSeq v2 | Newbler | 0 | |
| Lim_EarlyLifeDynamics | C2129723S | HostAssoc_Human Feces | Human feces | USA : Missouri | 38.130000 -90.000000 | reads SRA | 2 × 250 paired-end, MiSeq v2 | Newbler | 0 | |
| Lim_EarlyLifeDynamics | C2186333M | HostAssoc_Human Feces | Human feces | USA : Missouri | 38.130000 -90.000000 | reads SRA | 2 × 250 paired-end, MiSeq v2 | Newbler | 0 | |
| Lim_EarlyLifeDynamics | C2186333S | HostAssoc_Human Feces | Human feces | USA : Missouri | 38.130000 -90.000000 | reads SRA | 2 × 250 paired-end, MiSeq v2 | Newbler | 0 | |
| Lim_EarlyLifeDynamics | C2249052M | HostAssoc_Human Feces | Human feces | USA : Missouri | 38.130000 -90.000000 | reads SRA | 2 × 250 paired-end, MiSeq v2 | Newbler | 0 | |
| Lim_EarlyLifeDynamics | C2249052S | HostAssoc_Human Feces | Human feces | USA : Missouri | 38.130000 -90.000000 | reads SRA | 2 × 250 paired-end, MiSeq v2 | Newbler | 0 | |
| Lim_EarlyLifeDynamics | C233337M | HostAssoc_Human Feces | Human feces | USA : Missouri | 38.130000 -90.000000 | reads SRA | 2 × 250 paired-end, MiSeq v2 | Newbler | 0 | |
| Lim_EarlyLifeDynamics | C233337S | HostAssoc_Human Feces | Human feces | USA : Missouri | 38.130000 -90.000000 | reads SRA | 2 × 250 paired-end, MiSeq v2 | Newbler | 0 | |
| Lim_EarlyLifeDynamics | C260838M | HostAssoc_Human Feces | Human feces | USA : Missouri | 38.130000 -90.000000 | reads SRA | 2 × 250 paired-end, MiSeq v2 | Newbler | 0 | |
| Lim_EarlyLifeDynamics | C260838S | HostAssoc_Human Feces | Human feces | USA : Missouri | 38.130000 -90.000000 | reads SRA | 2 × 250 paired-end, MiSeq v2 | Newbler | 0 | |
| Lim_EarlyLifeDynamics | D103848M | HostAssoc_Human Feces | Human feces | USA : Missouri | 38.130000 -90.000000 | reads SRA | 2 × 250 paired-end, MiSeq v2 | Newbler | 0 | |
| Lim_EarlyLifeDynamics | D103848S | HostAssoc_Human Feces | Human feces | USA : Missouri | 38.130000 -90.000000 | reads SRA | 2 × 250 paired-end, MiSeq v2 | Newbler | 0 | |
| Lim_EarlyLifeDynamics | D1128367M | HostAssoc_Human Feces | Human feces | USA : Missouri | 38.130000 -90.000000 | reads SRA | 2 × 250 paired-end, MiSeq v2 | Newbler | 0 | |
| Lim_EarlyLifeDynamics | D1128367S | HostAssoc_Human Feces | Human feces | USA : Missouri | 38.130000 -90.000000 | reads SRA | 2 × 250 paired-end, MiSeq v2 | Newbler | 0 | |
| Lim_EarlyLifeDynamics | D1188340M | HostAssoc_Human Feces | Human feces | USA : Missouri | 38.130000 -90.000000 | reads SRA | 2 × 250 paired-end, MiSeq v2 | Newbler | 0 | |
| Lim_EarlyLifeDynamics | D1188340S | HostAssoc_Human Feces | Human feces | USA : Missouri | 38.130000 -90.000000 | reads SRA | 2 × 250 paired-end, MiSeq v2 | Newbler | 0 | |
| Lim_EarlyLifeDynamics | D1247512M | HostAssoc_Human Feces | Human feces | USA : Missouri | 38.130000 -90.000000 | reads SRA | 2 × 250 paired-end, MiSeq v2 | Newbler | 0 | |

| Project Name | Virome Code | Habitat Type (code) | Ecosystem Type (text) | Location | GPS | Source | Sequencing Technology | Assembly Software | # crucivirus | Reference |
|---|---|---|---|---|---|---|---|---|---|---|
| Lim_EarlyLifeDynamics | D1247512S | HostAssoc_Human Feces | Human feces | USA : Missouri | 38.130000 -90.000000 | reads SRA | 2 x 250 paired-end, MiSeq v2 | Newbler | 0 | |
| Lim_EarlyLifeDynamics | D132131M | HostAssoc_Human Feces | Human feces | USA : Missouri | 38.130000 -90.000000 | reads SRA | 2 x 250 paired-end, MiSeq v2 | Newbler | 0 | |
| Lim_EarlyLifeDynamics | D132131S | HostAssoc_Human Feces | Human feces | USA : Missouri | 38.130000 -90.000000 | reads SRA | 2 x 250 paired-end, MiSeq v2 | Newbler | 0 | |
| Lim_EarlyLifeDynamics | D166691M | HostAssoc_Human Feces | Human feces | USA : Missouri | 38.130000 -90.000000 | reads SRA | 2 x 250 paired-end, MiSeq v2 | Newbler | 0 | |
| Lim_EarlyLifeDynamics | D166691S | HostAssoc_Human Feces | Human feces | USA : Missouri | 38.130000 -90.000000 | reads SRA | 2 x 250 paired-end, MiSeq v2 | Newbler | 0 | |
| Lim_EarlyLifeDynamics | D203840M | HostAssoc_Human Feces | Human feces | USA : Missouri | 38.130000 -90.000000 | reads SRA | 2 x 250 paired-end, MiSeq v2 | Newbler | 0 | |
| Lim_EarlyLifeDynamics | D203840S | HostAssoc_Human Feces | Human feces | USA : Missouri | 38.130000 -90.000000 | reads SRA | 2 x 250 paired-end, MiSeq v2 | Newbler | 0 | |
| Lim_EarlyLifeDynamics | D2122062M | HostAssoc_Human Feces | Human feces | USA : Missouri | 38.130000 -90.000000 | reads SRA | 2 x 250 paired-end, MiSeq v2 | Newbler | 0 | |
| Lim_EarlyLifeDynamics | D2122062S | HostAssoc_Human Feces | Human feces | USA : Missouri | 38.130000 -90.000000 | reads SRA | 2 x 250 paired-end, MiSeq v2 | Newbler | 0 | |
| Lim_EarlyLifeDynamics | D2188216M | HostAssoc_Human Feces | Human feces | USA : Missouri | 38.130000 -90.000000 | reads SRA | 2 x 250 paired-end, MiSeq v2 | Newbler | 0 | |
| Lim_EarlyLifeDynamics | D2188216S | HostAssoc_Human Feces | Human feces | USA : Missouri | 38.130000 -90.000000 | reads SRA | 2 x 250 paired-end, MiSeq v2 | Newbler | 0 | |
| Lim_EarlyLifeDynamics | D2247517M | HostAssoc_Human Feces | Human feces | USA : Missouri | 38.130000 -90.000000 | reads SRA | 2 x 250 paired-end, MiSeq v2 | Newbler | 0 | |
| Lim_EarlyLifeDynamics | D2247517S | HostAssoc_Human Feces | Human feces | USA : Missouri | 38.130000 -90.000000 | reads SRA | 2 x 250 paired-end, MiSeq v2 | Newbler | 0 | |
| Lim_EarlyLifeDynamics | D232028M | HostAssoc_Human Feces | Human feces | USA : Missouri | 38.130000 -90.000000 | reads SRA | 2 x 250 paired-end, MiSeq v2 | Newbler | 0 | |
| Lim_EarlyLifeDynamics | D232028S | HostAssoc_Human Feces | Human feces | USA : Missouri | 38.130000 -90.000000 | reads SRA | 2 x 250 paired-end, MiSeq v2 | Newbler | 0 | |
| Lim_EarlyLifeDynamics | D268614M | HostAssoc_Human Feces | Human feces | USA : Missouri | 38.130000 -90.000000 | reads SRA | 2 x 250 paired-end, MiSeq v2 | Newbler | 0 | |
| Lim_EarlyLifeDynamics | D268614S | HostAssoc_Human Feces | Human feces | USA : Missouri | 38.130000 -90.000000 | reads SRA | 2 x 250 paired-end, MiSeq v2 | Newbler | 0 | |
| Minot_RapidEvolution | d00X1 | HostAssoc_Human Feces | Human feces | USA : Pennsylvania | 39.947802 -75.193702 | reads SRA | Illumina HiSeq2000 | IDBA_UD | 0 | |
| Minot_RapidEvolution | d00X2 | HostAssoc_Human Feces | Human feces | USA : Pennsylvania | 39.947802 -75.193702 | reads SRA | Illumina HiSeq2000 | IDBA_UD | 0 | |
| Minot_RapidEvolution | d01X1 | HostAssoc_Human Feces | Human feces | USA : Pennsylvania | 39.947802 -75.193702 | reads SRA | Illumina HiSeq2000 | IDBA_UD | 0 | |
| Minot_RapidEvolution | d02X1 | HostAssoc_Human Feces | Human feces | USA : Pennsylvania | 39.947802 -75.193702 | reads SRA | Illumina HiSeq2000 | IDBA_UD | 0 | |
| Minot_RapidEvolution | d03X1 | HostAssoc_Human Feces | Human feces | USA : Pennsylvania | 39.947802 -75.193702 | reads SRA | Illumina HiSeq2000 | IDBA_UD | 0 | |
| Minot_RapidEvolution | d03X2 | HostAssoc_Human Feces | Human feces | USA : Pennsylvania | 39.947802 -75.193702 | reads SRA | Illumina HiSeq2000 | IDBA_UD | 0 | |
| Minot_RapidEvolution | d04X1 | HostAssoc_Human Feces | Human feces | USA : Pennsylvania | 39.947802 -75.193702 | reads SRA | Illumina HiSeq2000 | IDBA_UD | 0 | |
| Minot_RapidEvolution | d05X1 | HostAssoc_Human Feces | Human feces | USA : Pennsylvania | 39.947802 -75.193702 | reads SRA | Illumina HiSeq2000 | IDBA_UD | 0 | |
| Minot_RapidEvolution | d11X1 | HostAssoc_Human Feces | Human feces | USA : Pennsylvania | 39.947802 -75.193702 | reads SRA | Illumina HiSeq2000 | IDBA_UD | 0 | |
| Minot_RapidEvolution | d11X2 | HostAssoc_Human Feces | Human feces | USA : Pennsylvania | 39.947802 -75.193702 | reads SRA | Illumina HiSeq2000 | IDBA_UD | 0 | |
| Minot_RapidEvolution | d12X1 | HostAssoc_Human Feces | Human feces | USA : Pennsylvania | 39.947802 -75.193702 | reads SRA | Illumina HiSeq2000 | IDBA_UD | 0 | |
| Minot_RapidEvolution | d12X2 | HostAssoc_Human Feces | Human feces | USA : Pennsylvania | 39.947802 -75.193702 | reads SRA | Illumina HiSeq2000 | IDBA_UD | 0 | |
| Minot_RapidEvolution | d13X1 | HostAssoc_Human Feces | Human feces | USA : Pennsylvania | 39.947802 -75.193702 | reads SRA | Illumina HiSeq2000 | IDBA_UD | 0 | Minot S. Bryson A. Chehoud C. Wu GD. Lewis JD. Bushman FD. (2013) Rapid evolution of the human gut virome. Proc Natl Acad Sci U S A. 110(30):12450-5. doi: 10.1073/pnas.1300833110. |
| Minot_RapidEvolution | d13X2 | HostAssoc_Human Feces | Human feces | USA : Pennsylvania | 39.947802 -75.193702 | reads SRA | Illumina HiSeq2000 | IDBA_UD | 0 | |
| Minot_RapidEvolution | d14X1 | HostAssoc_Human Feces | Human feces | USA : Pennsylvania | 39.947802 -75.193702 | reads SRA | Illumina HiSeq2000 | IDBA_UD | 0 | |
| Minot_RapidEvolution | d15X1 | HostAssoc_Human Feces | Human feces | USA : Pennsylvania | 39.947802 -75.193702 | reads SRA | Illumina HiSeq2000 | IDBA_UD | 0 | |
| Minot_RapidEvolution | d21X1 | HostAssoc_Human Feces | Human feces | USA : Pennsylvania | 39.947802 -75.193702 | reads SRA | Illumina HiSeq2000 | IDBA_UD | 0 | |
| Minot_RapidEvolution | d21X2 | HostAssoc_Human Feces | Human feces | USA : Pennsylvania | 39.947802 -75.193702 | reads SRA | Illumina HiSeq2000 | IDBA_UD | 0 | |
| Minot_RapidEvolution | d22X1 | HostAssoc_Human Feces | Human feces | USA : Pennsylvania | 39.947802 -75.193702 | reads SRA | Illumina HiSeq2000 | IDBA_UD | 0 | |
| Minot_RapidEvolution | d22X2 | HostAssoc_Human Feces | Human feces | USA : Pennsylvania | 39.947802 -75.193702 | reads SRA | Illumina HiSeq2000 | IDBA_UD | 0 | |
| Minot_RapidEvolution | d23X1 | HostAssoc_Human Feces | Human feces | USA : Pennsylvania | 39.947802 -75.193702 | reads SRA | Illumina HiSeq2000 | IDBA_UD | 0 | |
| Minot_RapidEvolution | d23X2 | HostAssoc_Human Feces | Human feces | USA : Pennsylvania | 39.947802 -75.193702 | reads SRA | Illumina HiSeq2000 | IDBA_UD | 0 | |
| Minot_RapidEvolution | d24X1 | HostAssoc_Human Feces | Human feces | USA : Pennsylvania | 39.947802 -75.193702 | reads SRA | Illumina HiSeq2000 | IDBA_UD | 0 | |
| Minot_RapidEvolution | d25X1 | HostAssoc_Human Feces | Human feces | USA : Pennsylvania | 39.947802 -75.193702 | reads SRA | Illumina HiSeq2000 | IDBA_UD | 0 | |
| Minot_RapidEvolution | Mic1014X01 | HostAssoc_Human Feces | Human feces | USA : Pennsylvania | 39.947802 -75.193702 | reads SRA | Illumina HiSeq2000 | IDBA_UD | 0 | |

| Project Name | Virome Code | Habitat Type (code) | Ecosystem Type (text) | Location | GPS | Source | Sequencing Technology | Assembly Software | # crucivirus | Reference |
|---|---|---|---|---|---|---|---|---|---|---|
| Minot_RapidEvolution | Mic1014X02 | HostAssoc_Human Feces | Human feces | USA : Pennsylvania | 39.947802 -75.193702 | reads SRA | Illumina HiSeq2000 | IDBA_UD | 0 | |
| Minot_RapidEvolution | Mic1014X03 | HostAssoc_Human Feces | Human feces | USA : Pennsylvania | 39.947802 -75.193702 | reads SRA | Illumina HiSeq2000 | IDBA_UD | 0 | |
| Minot_VariableLoci | Loci1470 | HostAssoc_Human Feces | Human feces | USA : Pennsylvania | 39.947802 -75.193702 | contigs Metavir | Illumina HiSeq2000 | ? | 0 | Minot S. Grunberg S. Wu GD. Lewis JD. Bushman FD. Hypervariable loci in the human gut virome. Proc Natl Acad Sci U S A. 2012 Mar 6;109(10):3962-6. doi: 10.1073/pnas.1119061109. |
| Norman_EntericCrohnUlcere | 10568 | HostAssoc_Human Feces | Human feces | UK : Cambridge | 52.175148 0.140482 | reads ENA | Miseq paired end | IDBA_UD | 0 | |
| Norman_EntericCrohnUlcere | 2409 | HostAssoc_Human Feces | Human feces | USA : Illinois | 41.874204 -87.668708 | reads ENA | Miseq paired end | IDBA_UD | 0 | |
| Norman_EntericCrohnUlcere | 2411 | HostAssoc_Human Feces | Human feces | USA : Illinois | 41.874204 -87.668708 | reads ENA | Miseq paired end | IDBA_UD | 0 | |
| Norman_EntericCrohnUlcere | 2412 | HostAssoc_Human Feces | Human feces | USA : Illinois | 41.874204 -87.668708 | reads ENA | Miseq paired end | IDBA_UD | 0 | |
| Norman_EntericCrohnUlcere | 2413 | HostAssoc_Human Feces | Human feces | USA : Illinois | 41.874204 -87.668708 | reads ENA | Miseq paired end | IDBA_UD | 0 | |
| Norman_EntericCrohnUlcere | 2414 | HostAssoc_Human Feces | Human feces | USA : Illinois | 41.874204 -87.668708 | reads ENA | Miseq paired end | IDBA_UD | 0 | |
| Norman_EntericCrohnUlcere | 2415 | HostAssoc_Human Feces | Human feces | USA : Illinois | 41.874204 -87.668708 | reads ENA | Miseq paired end | IDBA_UD | 0 | |
| Norman_EntericCrohnUlcere | 2419 | HostAssoc_Human Feces | Human feces | USA : Illinois | 41.874204 -87.668708 | reads ENA | Miseq paired end | IDBA_UD | 0 | |
| Norman_EntericCrohnUlcere | 2420 | HostAssoc_Human Feces | Human feces | USA : Illinois | 41.874204 -87.668708 | reads ENA | Miseq paired end | IDBA_UD | 0 | |
| Norman_EntericCrohnUlcere | 2422 | HostAssoc_Human Feces | Human feces | USA : Illinois | 41.874204 -87.668708 | reads ENA | Miseq paired end | IDBA_UD | 0 | |
| Norman_EntericCrohnUlcere | 2424 | HostAssoc_Human Feces | Human feces | USA : Illinois | 41.874204 -87.668708 | reads ENA | Miseq paired end | IDBA_UD | 0 | |
| Norman_EntericCrohnUlcere | 2426 | HostAssoc_Human Feces | Human feces | USA : Illinois | 41.874204 -87.668708 | reads ENA | Miseq paired end | IDBA_UD | 0 | |
| Norman_EntericCrohnUlcere | 2448 | HostAssoc_Human Feces | Human feces | USA : Illinois | 41.874204 -87.668708 | reads ENA | Miseq paired end | IDBA_UD | 0 | |
| Norman_EntericCrohnUlcere | 7544 | HostAssoc_Human Feces | Human feces | UK : Cambridge | 52.175148 0.140482 | reads ENA | Miseq paired end | IDBA_UD | 0 | |
| Norman_EntericCrohnUlcere | 7547 | HostAssoc_Human Feces | Human feces | UK : Cambridge | 52.175148 0.140482 | reads ENA | Miseq paired end | IDBA_UD | 0 | |
| Norman_EntericCrohnUlcere | 7550 | HostAssoc_Human Feces | Human feces | UK : Cambridge | 52.175148 0.140482 | reads ENA | Miseq paired end | IDBA_UD | 0 | |
| Norman_EntericCrohnUlcere | 7552 | HostAssoc_Human Feces | Human feces | UK : Cambridge | 52.175148 0.140482 | reads ENA | Miseq paired end | IDBA_UD | 0 | |
| Norman_EntericCrohnUlcere | 7554 | HostAssoc_Human Feces | Human feces | UK : Cambridge | 52.175148 0.140482 | reads ENA | Miseq paired end | IDBA_UD | 0 | |
| Norman_EntericCrohnUlcere | 7556 | HostAssoc_Human Feces | Human feces | UK : Cambridge | 52.175148 0.140482 | reads ENA | Miseq paired end | IDBA_UD | 0 | |
| Norman_EntericCrohnUlcere | 7560 | HostAssoc_Human Feces | Human feces | UK : Cambridge | 52.175148 0.140482 | reads ENA | Miseq paired end | IDBA_UD | 0 | |
| Norman_EntericCrohnUlcere | 7562 | HostAssoc_Human Feces | Human feces | UK : Cambridge | 52.175148 0.140482 | reads ENA | Miseq paired end | IDBA_UD | 0 | |
| Norman_EntericCrohnUlcere | 7564 | HostAssoc_Human Feces | Human feces | UK : Cambridge | 52.175148 0.140482 | reads ENA | Miseq paired end | IDBA_UD | 0 | |
| Norman_EntericCrohnUlcere | 7577 | HostAssoc_Human Feces | Human feces | UK : Cambridge | 52.175148 0.140482 | reads ENA | Miseq paired end | IDBA_UD | 0 | |
| Norman_EntericCrohnUlcere | 7589 | HostAssoc_Human Feces | Human feces | UK : Cambridge | 52.175148 0.140482 | reads ENA | Miseq paired end | IDBA_UD | 0 | |
| Norman_EntericCrohnUlcere | 7605 | HostAssoc_Human Feces | Human feces | UK : Cambridge | 52.175148 0.140482 | reads ENA | Miseq paired end | IDBA_UD | 0 | |
| Norman_EntericCrohnUlcere | 7612 | HostAssoc_Human Feces | Human feces | UK : Cambridge | 52.175148 0.140482 | reads ENA | Miseq paired end | IDBA_UD | 0 | |
| Norman_EntericCrohnUlcere | 7614 | HostAssoc_Human Feces | Human feces | UK : Cambridge | 52.175148 0.140482 | reads ENA | Miseq paired end | IDBA_UD | 0 | |
| Norman_EntericCrohnUlcere | 7631 | HostAssoc_Human Feces | Human feces | UK : Cambridge | 52.175148 0.140482 | reads ENA | Miseq paired end | IDBA_UD | 0 | |
| Norman_EntericCrohnUlcere | 7632 | HostAssoc_Human Feces | Human feces | UK : Cambridge | 52.175148 0.140482 | reads ENA | Miseq paired end | IDBA_UD | 0 | |
| Norman_EntericCrohnUlcere | 7634 | HostAssoc_Human Feces | Human feces | UK : Cambridge | 52.175148 0.140482 | reads ENA | Miseq paired end | IDBA_UD | 0 | |
| Norman_EntericCrohnUlcere | 7647 | HostAssoc_Human Feces | Human feces | UK : Cambridge | 52.175148 0.140482 | reads ENA | Miseq paired end | IDBA_UD | 0 | |
| Norman_EntericCrohnUlcere | 7650 | HostAssoc_Human Feces | Human feces | UK : Cambridge | 52.175148 0.140482 | reads ENA | Miseq paired end | IDBA_UD | 0 | |
| Norman_EntericCrohnUlcere | 7652 | HostAssoc_Human Feces | Human feces | UK : Cambridge | 52.175148 0.140482 | reads ENA | Miseq paired end | IDBA_UD | 0 | |
| Norman_EntericCrohnUlcere | 10569 | HostAssoc_Human Feces | Human feces | UK : Cambridge | 52.175148 0.140482 | reads ENA | Miseq paired end | IDBA_UD | 0 | |
| Norman_EntericCrohnUlcere | 10571 | HostAssoc_Human Feces | Human feces | UK : Cambridge | 52.175148 0.140482 | reads ENA | Miseq paired end | IDBA_UD | 0 | |
| Norman_EntericCrohnUlcere | 10577 | HostAssoc_Human Feces | Human feces | UK : Cambridge | 52.175148 0.140482 | reads ENA | Miseq paired end | IDBA_UD | 0 | |
| Norman_EntericCrohnUlcere | 10578 | HostAssoc_Human Feces | Human feces | UK : Cambridge | 52.175148 0.140482 | reads ENA | Miseq paired end | IDBA_UD | 0 | |
| Norman_EntericCrohnUlcere | 10579 | HostAssoc_Human Feces | Human feces | UK : Cambridge | 52.175148 0.140482 | reads ENA | Miseq paired end | IDBA_UD | 0 | |
| Norman_EntericCrohnUlcere | 10586 | HostAssoc_Human Feces | Human feces | UK : Cambridge | 52.175148 0.140482 | reads ENA | Miseq paired end | IDBA_UD | 0 | |

| Project Name | Virome Code | Habitat Type (code) | Ecosystem Type (text) | Location | GPS | Source | Sequencing Technology | Assembly Software | # crucivirus | Reference |
|---|---|---|---|---|---|---|---|---|---|---|
| Norman_EntericCrohnUlcere | 10587 | HostAssoc_Human Feces | Human feces | UK : Cambridge | 52.175148 0.140482 | reads ENA | Miseq paired end | IDBA_UD | 0 | |
| Norman_EntericCrohnUlcere | 10599 | HostAssoc_Human Feces | Human feces | UK : Cambridge | 52.175148 0.140482 | reads ENA | Miseq paired end | IDBA_UD | 0 | |
| Norman_EntericCrohnUlcere | 10613 | HostAssoc_Human Feces | Human feces | UK : Cambridge | 52.175148 0.140482 | reads ENA | Miseq paired end | IDBA_UD | 0 | |
| Norman_EntericCrohnUlcere | 10615 | HostAssoc_Human Feces | Human feces | UK : Cambridge | 52.175148 0.140482 | reads ENA | Miseq paired end | IDBA_UD | 0 | |
| Norman_EntericCrohnUlcere | 10618 | HostAssoc_Human Feces | Human feces | UK : Cambridge | 52.175148 0.140482 | reads ENA | Miseq paired end | IDBA_UD | 0 | |
| Norman_EntericCrohnUlcere | 12274 | HostAssoc_Human Feces | Human feces | UK : Cambridge | 52.175148 0.140482 | reads ENA | Miseq paired end | IDBA_UD | 0 | |
| Norman_EntericCrohnUlcere | 13463 | HostAssoc_Human Feces | Human feces | USA : Illinois | 41.874204 -87.668708 | reads ENA | Miseq paired end | IDBA_UD | 0 | |
| Norman_EntericCrohnUlcere | 13469 | HostAssoc_Human Feces | Human feces | USA : Illinois | 41.874204 -87.668708 | reads ENA | Miseq paired end | IDBA_UD | 0 | |
| Norman_EntericCrohnUlcere | 13475 | HostAssoc_Human Feces | Human feces | USA : Illinois | 41.874204 -87.668708 | reads ENA | Miseq paired end | IDBA_UD | 0 | |
| Norman_EntericCrohnUlcere | 13478 | HostAssoc_Human Feces | Human feces | USA : Illinois | 41.874204 -87.668708 | reads ENA | Miseq paired end | IDBA_UD | 0 | |
| Norman_EntericCrohnUlcere | 13481 | HostAssoc_Human Feces | Human feces | USA : Illinois | 41.874204 -87.668708 | reads ENA | Miseq paired end | IDBA_UD | 0 | |
| Norman_EntericCrohnUlcere | 13484 | HostAssoc_Human Feces | Human feces | USA : Illinois | 41.874204 -87.668708 | reads ENA | Miseq paired end | IDBA_UD | 0 | |
| Norman_EntericCrohnUlcere | 13487 | HostAssoc_Human Feces | Human feces | USA : Illinois | 41.874204 -87.668708 | reads ENA | Miseq paired end | IDBA_UD | 0 | |
| Norman_EntericCrohnUlcere | 13490 | HostAssoc_Human Feces | Human feces | USA : Illinois | 41.874204 -87.668708 | reads ENA | Miseq paired end | IDBA_UD | 0 | |
| Norman_EntericCrohnUlcere | 13519 | HostAssoc_Human Feces | Human feces | UK : Cambridge | 52.175148 0.140482 | reads ENA | Miseq paired end | IDBA_UD | 0 | |
| Norman_EntericCrohnUlcere | 13520 | HostAssoc_Human Feces | Human feces | UK : Cambridge | 52.175148 0.140482 | reads ENA | Miseq paired end | IDBA_UD | 0 | |
| Norman_EntericCrohnUlcere | 13527 | HostAssoc_Human Feces | Human feces | UK : Cambridge | 52.175148 0.140482 | reads ENA | Miseq paired end | IDBA_UD | 0 | |
| Norman_EntericCrohnUlcere | 13530 | HostAssoc_Human Feces | Human feces | UK : Cambridge | 52.175148 0.140482 | reads ENA | Miseq paired end | IDBA_UD | 0 | |
| Norman_EntericCrohnUlcere | 14881 | HostAssoc_Human Feces | Human feces | UK : Cambridge | 52.175148 0.140482 | reads ENA | Miseq paired end | IDBA_UD | 0 | |
| Norman_EntericCrohnUlcere | 14882 | HostAssoc_Human Feces | Human feces | UK : Cambridge | 52.175148 0.140482 | reads ENA | Miseq paired end | IDBA_UD | 0 | |
| Norman_EntericCrohnUlcere | 14887 | HostAssoc_Human Feces | Human feces | UK : Cambridge | 52.175148 0.140482 | reads ENA | Miseq paired end | IDBA_UD | 0 | |
| Norman_EntericCrohnUlcere | 14888 | HostAssoc_Human Feces | Human feces | UK : Cambridge | 52.175148 0.140482 | reads ENA | Miseq paired end | IDBA_UD | 0 | |
| Norman_EntericCrohnUlcere | 14894 | HostAssoc_Human Feces | Human feces | UK : Cambridge | 52.175148 0.140482 | reads ENA | Miseq paired end | IDBA_UD | 0 | |
| Norman_EntericCrohnUlcere | 14898 | HostAssoc_Human Feces | Human feces | UK : Cambridge | 52.175148 0.140482 | reads ENA | Miseq paired end | IDBA_UD | 0 | |
| Norman_EntericCrohnUlcere | 14900 | HostAssoc_Human Feces | Human feces | UK : Cambridge | 52.175148 0.140482 | reads ENA | Miseq paired end | IDBA_UD | 0 | |
| Norman_EntericCrohnUlcere | 14918 | HostAssoc_Human Feces | Human feces | UK : Cambridge | 52.175148 0.140482 | reads ENA | Miseq paired end | IDBA_UD | 0 | |
| Norman_EntericCrohnUlcere | 14950 | HostAssoc_Human Feces | Human feces | UK : Cambridge | 52.175148 0.140482 | reads ENA | Miseq paired end | IDBA_UD | 0 | |
| Norman_EntericCrohnUlcere | 14978 | HostAssoc_Human Feces | Human feces | UK : Cambridge | 52.175148 0.140482 | reads ENA | Miseq paired end | IDBA_UD | 0 | |
| Norman_EntericCrohnUlcere | 14986 | HostAssoc_Human Feces | Human feces | UK : Cambridge | 52.175148 0.140482 | reads ENA | Miseq paired end | IDBA_UD | 0 | |
| Norman_EntericCrohnUlcere | 14998 | HostAssoc_Human Feces | Human feces | UK : Cambridge | 52.175148 0.140482 | reads ENA | Miseq paired end | IDBA_UD | 0 | |
| Norman_EntericCrohnUlcere | 14999 | HostAssoc_Human Feces | Human feces | UK : Cambridge | 52.175148 0.140482 | reads ENA | Miseq paired end | IDBA_UD | 0 | |
| Norman_EntericCrohnUlcere | 15011 | HostAssoc_Human Feces | Human feces | UK : Cambridge | 52.175148 0.140482 | reads ENA | Miseq paired end | IDBA_UD | 0 | |
| Norman_EntericCrohnUlcere | 15019 | HostAssoc_Human Feces | Human feces | UK : Cambridge | 52.175148 0.140482 | reads ENA | Miseq paired end | IDBA_UD | 0 | |
| Norman_EntericCrohnUlcere | 15027 | HostAssoc_Human Feces | Human feces | UK : Cambridge | 52.175148 0.140482 | reads ENA | Miseq paired end | IDBA_UD | 0 | |
| Norman_EntericCrohnUlcere | 15078 | HostAssoc_Human Feces | Human feces | UK : Cambridge | 52.175148 0.140482 | reads ENA | Miseq paired end | IDBA_UD | 0 | |
| Norman_EntericCrohnUlcere | 15105 | HostAssoc_Human Feces | Human feces | UK : Cambridge | 52.175148 0.140482 | reads ENA | Miseq paired end | IDBA_UD | 0 | |
| Norman_EntericCrohnUlcere | 15763 | HostAssoc_Human Feces | Human feces | UK : Cambridge | 52.175148 0.140482 | reads ENA | Miseq paired end | IDBA_UD | 0 | |
| Norman_EntericCrohnUlcere | 15767 | HostAssoc_Human Feces | Human feces | UK : Cambridge | 52.175148 0.140482 | reads ENA | Miseq paired end | IDBA_UD | 0 | |
| Norman_EntericCrohnUlcere | 15777 | HostAssoc_Human Feces | Human feces | UK : Cambridge | 52.175148 0.140482 | reads ENA | Miseq paired end | IDBA_UD | 0 | |
| Norman_EntericCrohnUlcere | 15783 | HostAssoc_Human Feces | Human feces | UK : Cambridge | 52.175148 0.140482 | reads ENA | Miseq paired end | IDBA_UD | 0 | |
| Norman_EntericCrohnUlcere | 15785 | HostAssoc_Human Feces | Human feces | UK : Cambridge | 52.175148 0.140482 | reads ENA | Miseq paired end | IDBA_UD | 0 | |
| Norman_EntericCrohnUlcere | 15787 | HostAssoc_Human Feces | Human feces | UK : Cambridge | 52.175148 0.140482 | reads ENA | Miseq paired end | IDBA_UD | 0 | |

| Project Name | Virome Code | Habitat Type (code) | Ecosystem Type (text) | Location | GPS | Source | Sequencing Technology | Assembly Software | # crucivirus | Reference |
|---|---|---|---|---|---|---|---|---|---|---|
| Norman_EntericCrohnUlcere | 15789 | HostAssoc_Human Feces | Human feces | UK : Cambridge | 52.175148 0.140482 | reads ENA | Miseq paired end | IDBA_UD | 0 | Norman JM. Handley SA. Baldridge MT. Droit L. Liu CY. Keller BC. Kambal A. Monaco CL. Zhao G. Fleshner P. Stappenbeck TS. McGovern DP. Keshavarzian A. Mutlu EA. Sauk J. Gevers D. Xavier RJ. Wang D. Parkes M. Virgin HW. Disease-specific alterations in the enteric virome in inflammatory bowel disease. Cell. 2015 Jan 29;160(3):447-60. doi: 10.1016/j.cell.2015.01.002. |
| Norman_EntericCrohnUlcere | 15791 | HostAssoc_Human Feces | Human feces | UK : Cambridge | 52.175148 0.140482 | reads ENA | Miseq paired end | IDBA_UD | 0 | |
| Norman_EntericCrohnUlcere | 15800 | HostAssoc_Human Feces | Human feces | UK : Cambridge | 52.175148 0.140482 | reads ENA | Miseq paired end | IDBA_UD | 0 | |
| Norman_EntericCrohnUlcere | 15802 | HostAssoc_Human Feces | Human feces | UK : Cambridge | 52.175148 0.140482 | reads ENA | Miseq paired end | IDBA_UD | 0 | |
| Norman_EntericCrohnUlcere | 15804 | HostAssoc_Human Feces | Human feces | UK : Cambridge | 52.175148 0.140482 | reads ENA | Miseq paired end | IDBA_UD | 0 | |
| Norman_EntericCrohnUlcere | 15819 | HostAssoc_Human Feces | Human feces | UK : Cambridge | 52.175148 0.140482 | reads ENA | Miseq paired end | IDBA_UD | 0 | |
| Norman_EntericCrohnUlcere | 15821 | HostAssoc_Human Feces | Human feces | UK : Cambridge | 52.175148 0.140482 | reads ENA | Miseq paired end | IDBA_UD | 0 | |
| Norman_EntericCrohnUlcere | 15825 | HostAssoc_Human Feces | Human feces | UK : Cambridge | 52.175148 0.140482 | reads ENA | Miseq paired end | IDBA_UD | 0 | |
| Norman_EntericCrohnUlcere | 15828 | HostAssoc_Human Feces | Human feces | UK : Cambridge | 52.175148 0.140482 | reads ENA | Miseq paired end | IDBA_UD | 0 | |
| Norman_EntericCrohnUlcere | 15829 | HostAssoc_Human Feces | Human feces | UK : Cambridge | 52.175148 0.140482 | reads ENA | Miseq paired end | IDBA_UD | 0 | |
| Norman_EntericCrohnUlcere | 15830 | HostAssoc_Human Feces | Human feces | UK : Cambridge | 52.175148 0.140482 | reads ENA | Miseq paired end | IDBA_UD | 0 | |
| Norman_EntericCrohnUlcere | 15831 | HostAssoc_Human Feces | Human feces | UK : Cambridge | 52.175148 0.140482 | reads ENA | Miseq paired end | IDBA_UD | 0 | |
| Norman_EntericCrohnUlcere | 15832 | HostAssoc_Human Feces | Human feces | UK : Cambridge | 52.175148 0.140482 | reads ENA | Miseq paired end | IDBA_UD | 0 | |
| Norman_EntericCrohnUlcere | 15834 | HostAssoc_Human Feces | Human feces | UK : Cambridge | 52.175148 0.140482 | reads ENA | Miseq paired end | IDBA_UD | 0 | |
| Norman_EntericCrohnUlcere | 15837 | HostAssoc_Human Feces | Human feces | UK : Cambridge | 52.175148 0.140482 | reads ENA | Miseq paired end | IDBA_UD | 0 | |
| Norman_EntericCrohnUlcere | 15839 | HostAssoc_Human Feces | Human feces | UK : Cambridge | 52.175148 0.140482 | reads ENA | Miseq paired end | IDBA_UD | 0 | |
| Norman_EntericCrohnUlcere | 15841 | HostAssoc_Human Feces | Human feces | UK : Cambridge | 52.175148 0.140482 | reads ENA | Miseq paired end | IDBA_UD | 0 | |
| Norman_EntericCrohnUlcere | 16043 | HostAssoc_Human Feces | Human feces | USA : Massachusetts | 42.363482 -71.068472 | reads ENA | Miseq paired end | IDBA_UD | 0 | |
| Norman_EntericCrohnUlcere | 16045 | HostAssoc_Human Feces | Human feces | USA : Massachusetts | 42.363482 -71.068472 | reads ENA | Miseq paired end | IDBA_UD | 0 | |
| Norman_EntericCrohnUlcere | 16046 | HostAssoc_Human Feces | Human feces | USA : Massachusetts | 42.363482 -71.068472 | reads ENA | Miseq paired end | IDBA_UD | 0 | |
| Norman_EntericCrohnUlcere | 16047 | HostAssoc_Human Feces | Human feces | USA : Massachusetts | 42.363482 -71.068472 | reads ENA | Miseq paired end | IDBA_UD | 0 | |
| Norman_EntericCrohnUlcere | 16048 | HostAssoc_Human Feces | Human feces | USA : Massachusetts | 42.363482 -71.068472 | reads ENA | Miseq paired end | IDBA_UD | 0 | |
| Norman_EntericCrohnUlcere | 16049 | HostAssoc_Human Feces | Human feces | USA : Massachusetts | 42.363482 -71.068472 | reads ENA | Miseq paired end | IDBA_UD | 0 | |
| Norman_EntericCrohnUlcere | 16050 | HostAssoc_Human Feces | Human feces | USA : Massachusetts | 42.363482 -71.068472 | reads ENA | Miseq paired end | IDBA_UD | 0 | |
| Norman_EntericCrohnUlcere | 16051 | HostAssoc_Human Feces | Human feces | USA : Massachusetts | 42.363482 -71.068472 | reads ENA | Miseq paired end | IDBA_UD | 0 | |
| Norman_EntericCrohnUlcere | 16052 | HostAssoc_Human Feces | Human feces | USA : Massachusetts | 42.363482 -71.068472 | reads ENA | Miseq paired end | IDBA_UD | 0 | |
| Norman_EntericCrohnUlcere | 16053 | HostAssoc_Human Feces | Human feces | USA : Massachusetts | 42.363482 -71.068472 | reads ENA | Miseq paired end | IDBA_UD | 0 | |
| Norman_EntericCrohnUlcere | 16054 | HostAssoc_Human Feces | Human feces | USA : Massachusetts | 42.363482 -71.068472 | reads ENA | Miseq paired end | IDBA_UD | 0 | |
| Norman_EntericCrohnUlcere | 16055 | HostAssoc_Human Feces | Human feces | USA : Massachusetts | 42.363482 -71.068472 | reads ENA | Miseq paired end | IDBA_UD | 0 | |
| Norman_EntericCrohnUlcere | 16057 | HostAssoc_Human Feces | Human feces | USA : Massachusetts | 42.363482 -71.068472 | reads ENA | Miseq paired end | IDBA_UD | 0 | |
| Norman_EntericCrohnUlcere | 16058 | HostAssoc_Human Feces | Human feces | USA : Massachusetts | 42.363482 -71.068472 | reads ENA | Miseq paired end | IDBA_UD | 0 | |
| Norman_EntericCrohnUlcere | 16060 | HostAssoc_Human Feces | Human feces | USA : Massachusetts | 42.363482 -71.068472 | reads ENA | Miseq paired end | IDBA_UD | 0 | |
| Norman_EntericCrohnUlcere | 16061 | HostAssoc_Human Feces | Human feces | USA : Massachusetts | 42.363482 -71.068472 | reads ENA | Miseq paired end | IDBA_UD | 0 | |
| Norman_EntericCrohnUlcere | 16063 | HostAssoc_Human Feces | Human feces | USA : Massachusetts | 42.363482 -71.068472 | reads ENA | Miseq paired end | IDBA_UD | 0 | |
| Norman_EntericCrohnUlcere | 16064 | HostAssoc_Human Feces | Human feces | USA : Massachusetts | 42.363482 -71.068472 | reads ENA | Miseq paired end | IDBA_UD | 0 | |
| Norman_EntericCrohnUlcere | 16065 | HostAssoc_Human Feces | Human feces | USA : Massachusetts | 42.363482 -71.068472 | reads ENA | Miseq paired end | IDBA_UD | 0 | |
| Norman_EntericCrohnUlcere | 16067 | HostAssoc_Human Feces | Human feces | USA : Massachusetts | 42.363482 -71.068472 | reads ENA | Miseq paired end | IDBA_UD | 0 | |
| Norman_EntericCrohnUlcere | 16070 | HostAssoc_Human Feces | Human feces | USA : Massachusetts | 42.363482 -71.068472 | reads ENA | Miseq paired end | IDBA_UD | 0 | |
| Norman_EntericCrohnUlcere | 16071 | HostAssoc_Human Feces | Human feces | USA : Massachusetts | 42.363482 -71.068472 | reads ENA | Miseq paired end | IDBA_UD | 0 | |
| Norman_EntericCrohnUlcere | 16073 | HostAssoc_Human Feces | Human feces | USA : Massachusetts | 42.363482 -71.068472 | reads ENA | Miseq paired end | IDBA_UD | 0 | |
| Norman_EntericCrohnUlcere | 16074 | HostAssoc_Human Feces | Human feces | USA : Massachusetts | 42.363482 -71.068472 | reads ENA | Miseq paired end | IDBA_UD | 0 | |
| Norman_EntericCrohnUlcere | 16075 | HostAssoc_Human Feces | Human feces | USA : Massachusetts | 42.363482 -71.068472 | reads ENA | Miseq paired end | IDBA_UD | 0 | |

| Project Name | Virome Code | Habitat Type (code) | Ecosystem Type (text) | Location | GPS | Source | Sequencing Technology | Assembly Software | # crucivirus | Reference |
|---|---|---|---|---|---|---|---|---|---|---|
| Norman_EntericCrohnUlcere | 16076 | HostAssoc_Human Feces | Human feces | USA : Massachusetts | 42.363482 -71.068472 | reads ENA | Miseq paired end | IDBA_UD | 0 | |
| Norman_EntericCrohnUlcere | 16079 | HostAssoc_Human Feces | Human feces | USA : Massachusetts | 42.363482 -71.068472 | reads ENA | Miseq paired end | IDBA_UD | 0 | |
| Norman_EntericCrohnUlcere | 16080 | HostAssoc_Human Feces | Human feces | USA : Massachusetts | 42.363482 -71.068472 | reads ENA | Miseq paired end | IDBA_UD | 0 | |
| Norman_EntericCrohnUlcere | 16081 | HostAssoc_Human Feces | Human feces | USA : Massachusetts | 42.363482 -71.068472 | reads ENA | Miseq paired end | IDBA_UD | 0 | |
| Norman_EntericCrohnUlcere | 16082 | HostAssoc_Human Feces | Human feces | USA : Massachusetts | 42.363482 -71.068472 | reads ENA | Miseq paired end | IDBA_UD | 0 | |
| Norman_EntericCrohnUlcere | 16083 | HostAssoc_Human Feces | Human feces | USA : Massachusetts | 42.363482 -71.068472 | reads ENA | Miseq paired end | IDBA_UD | 0 | |
| Norman_EntericCrohnUlcere | 16084 | HostAssoc_Human Feces | Human feces | USA : Massachusetts | 42.363482 -71.068472 | reads ENA | Miseq paired end | IDBA_UD | 0 | |
| Norman_EntericCrohnUlcere | 16085 | HostAssoc_Human Feces | Human feces | USA : Massachusetts | 42.363482 -71.068472 | reads ENA | Miseq paired end | IDBA_UD | 0 | |
| Norman_EntericCrohnUlcere | 16087 | HostAssoc_Human Feces | Human feces | USA : Massachusetts | 42.363482 -71.068472 | reads ENA | Miseq paired end | IDBA_UD | 0 | |
| Norman_EntericCrohnUlcere | 16089 | HostAssoc_Human Feces | Human feces | USA : Illinois | 41.874204 -87.668708 | reads ENA | Miseq paired end | IDBA_UD | 0 | |
| Norman_EntericCrohnUlcere | 16090 | HostAssoc_Human Feces | Human feces | USA : Illinois | 41.874204 -87.668708 | reads ENA | Miseq paired end | IDBA_UD | 0 | |
| Norman_EntericCrohnUlcere | 16092 | HostAssoc_Human Feces | Human feces | USA : Illinois | 41.874204 -87.668708 | reads ENA | Miseq paired end | IDBA_UD | 0 | |
| Norman_EntericCrohnUlcere | 16094 | HostAssoc_Human Feces | Human feces | USA : Illinois | 41.874204 -87.668708 | reads ENA | Miseq paired end | IDBA_UD | 0 | |
| Norman_EntericCrohnUlcere | 16096 | HostAssoc_Human Feces | Human feces | USA : Illinois | 41.874204 -87.668708 | reads ENA | Miseq paired end | IDBA_UD | 0 | |
| Norman_EntericCrohnUlcere | 16097 | HostAssoc_Human Feces | Human feces | USA : Illinois | 41.874204 -87.668708 | reads ENA | Miseq paired end | IDBA_UD | 0 | |
| Norman_EntericCrohnUlcere | 16098 | HostAssoc_Human Feces | Human feces | USA : Illinois | 41.874204 -87.668708 | reads ENA | Miseq paired end | IDBA_UD | 0 | |
| Norman_EntericCrohnUlcere | 16099 | HostAssoc_Human Feces | Human feces | USA : Illinois | 41.874204 -87.668708 | reads ENA | Miseq paired end | IDBA_UD | 0 | |
| Norman_EntericCrohnUlcere | 16100 | HostAssoc_Human Feces | Human feces | USA : Illinois | 41.874204 -87.668708 | reads ENA | Miseq paired end | IDBA_UD | 0 | |
| Norman_EntericCrohnUlcere | 16101 | HostAssoc_Human Feces | Human feces | USA : Illinois | 41.874204 -87.668708 | reads ENA | Miseq paired end | IDBA_UD | 0 | |
| Norman_EntericCrohnUlcere | 16102 | HostAssoc_Human Feces | Human feces | USA : Illinois | 41.874204 -87.668708 | reads ENA | Miseq paired end | IDBA_UD | 0 | |
| Norman_EntericCrohnUlcere | 16103 | HostAssoc_Human Feces | Human feces | USA : Illinois | 41.874204 -87.668708 | reads ENA | Miseq paired end | IDBA_UD | 0 | |
| Norman_EntericCrohnUlcere | 16104 | HostAssoc_Human Feces | Human feces | USA : Illinois | 41.874204 -87.668708 | reads ENA | Miseq paired end | IDBA_UD | 0 | |
| Norman_EntericCrohnUlcere | 16105 | HostAssoc_Human Feces | Human feces | USA : Illinois | 41.874204 -87.668708 | reads ENA | Miseq paired end | IDBA_UD | 0 | |
| Norman_EntericCrohnUlcere | 16106 | HostAssoc_Human Feces | Human feces | USA : Illinois | 41.874204 -87.668708 | reads ENA | Miseq paired end | IDBA_UD | 0 | |
| Norman_EntericCrohnUlcere | 16107 | HostAssoc_Human Feces | Human feces | USA : Illinois | 41.874204 -87.668708 | reads ENA | Miseq paired end | IDBA_UD | 0 | |
| Norman_EntericCrohnUlcere | 16108 | HostAssoc_Human Feces | Human feces | USA : Illinois | 41.874204 -87.668708 | reads ENA | Miseq paired end | IDBA_UD | 0 | |
| Norman_EntericCrohnUlcere | 16110 | HostAssoc_Human Feces | Human feces | USA : Illinois | 41.874204 -87.668708 | reads ENA | Miseq paired end | IDBA_UD | 0 | |
| Norman_EntericCrohnUlcere | 16111 | HostAssoc_Human Feces | Human feces | USA : Illinois | 41.874204 -87.668708 | reads ENA | Miseq paired end | IDBA_UD | 0 | |
| Norman_EntericCrohnUlcere | 16112 | HostAssoc_Human Feces | Human feces | USA : Illinois | 41.874204 -87.668708 | reads ENA | Miseq paired end | IDBA_UD | 0 | |
| Norman_EntericCrohnUlcere | 16113 | HostAssoc_Human Feces | Human feces | USA : Illinois | 41.874204 -87.668708 | reads ENA | Miseq paired end | IDBA_UD | 0 | |
| Norman_EntericCrohnUlcere | 16114 | HostAssoc_Human Feces | Human feces | USA : Illinois | 41.874204 -87.668708 | reads ENA | Miseq paired end | IDBA_UD | 0 | |
| Norman_EntericCrohnUlcere | 16115 | HostAssoc_Human Feces | Human feces | USA : Illinois | 41.874204 -87.668708 | reads ENA | Miseq paired end | IDBA_UD | 0 | |
| Norman_EntericCrohnUlcere | 16116 | HostAssoc_Human Feces | Human feces | USA : Illinois | 41.874204 -87.668708 | reads ENA | Miseq paired end | IDBA_UD | 0 | |
| Norman_EntericCrohnUlcere | 16117 | HostAssoc_Human Feces | Human feces | USA : Illinois | 41.874204 -87.668708 | reads ENA | Miseq paired end | IDBA_UD | 0 | |
| Norman_EntericCrohnUlcere | 16118 | HostAssoc_Human Feces | Human feces | USA : Illinois | 41.874204 -87.668708 | reads ENA | Miseq paired end | IDBA_UD | 0 | |
| Norman_EntericCrohnUlcere | 16119 | HostAssoc_Human Feces | Human feces | USA : Illinois | 41.874204 -87.668708 | reads ENA | Miseq paired end | IDBA_UD | 0 | |
| Norman_EntericCrohnUlcere | 16120 | HostAssoc_Human Feces | Human feces | USA : Illinois | 41.874204 -87.668708 | reads ENA | Miseq paired end | IDBA_UD | 0 | |
| Norman_EntericCrohnUlcere | 16121 | HostAssoc_Human Feces | Human feces | USA : Massachusetts | 42.363482 -71.068472 | reads ENA | Miseq paired end | IDBA_UD | 0 | |
| Norman_EntericCrohnUlcere | 16122 | HostAssoc_Human Feces | Human feces | USA : Massachusetts | 42.363482 -71.068472 | reads ENA | Miseq paired end | IDBA_UD | 0 | |
| Norman_EntericCrohnUlcere | 16123 | HostAssoc_Human Feces | Human feces | USA : Massachusetts | 42.363482 -71.068472 | reads ENA | Miseq paired end | IDBA_UD | 0 | |
| Norman_EntericCrohnUlcere | 16124 | HostAssoc_Human Feces | Human feces | USA : Massachusetts | 42.363482 -71.068472 | reads ENA | Miseq paired end | IDBA_UD | 0 | |
| Norman_EntericCrohnUlcere | 16125 | HostAssoc_Human Feces | Human feces | USA : Massachusetts | 42.363482 -71.068472 | reads ENA | Miseq paired end | IDBA_UD | 0 | |

| Project Name | Virome Code | Habitat Type (code) | Ecosystem Type (text) | Location | GPS | Source | Sequencing Technology | Assembly Software | # crucivirus | Reference |
|---|---|---|---|---|---|---|---|---|---|---|
| Norman_EntericCrohnUlcere | 16126 | HostAssoc_Human Feces | Human feces | USA : Massachusetts | 42.363482 -71.068472 | reads ENA | Miseq paired end | IDBA_UD | 0 | |
| Norman_EntericCrohnUlcere | 16127 | HostAssoc_Human Feces | Human feces | USA : Massachusetts | 42.363482 -71.068472 | reads ENA | Miseq paired end | IDBA_UD | 0 | |
| Norman_EntericCrohnUlcere | 16128 | HostAssoc_Human Feces | Human feces | USA : Massachusetts | 42.363482 -71.068472 | reads ENA | Miseq paired end | IDBA_UD | 0 | |
| Norman_EntericCrohnUlcere | 16129 | HostAssoc_Human Feces | Human feces | USA : Massachusetts | 42.363482 -71.068472 | reads ENA | Miseq paired end | IDBA_UD | 0 | |
| Norman_EntericCrohnUlcere | 16130 | HostAssoc_Human Feces | Human feces | USA : Massachusetts | 42.363482 -71.068472 | reads ENA | Miseq paired end | IDBA_UD | 0 | |
| Norman_EntericCrohnUlcere | 16131 | HostAssoc_Human Feces | Human feces | USA : Massachusetts | 42.363482 -71.068472 | reads ENA | Miseq paired end | IDBA_UD | 0 | |
| PerezBrocal_CDViromes | CDv5176 | HostAssoc_Human Feces | Human feces | Spain : Valencia | 39.468228 -0.378304 | reads ENA | 454 GS-FLX Titanium | Newbler | 0 | |
| PerezBrocal_CDViromes | CDv5177 | HostAssoc_Human Feces | Human feces | Spain : Valencia | 39.468228 -0.378304 | reads ENA | 454 GS-FLX Titanium | Newbler | 0 | |
| PerezBrocal_CDViromes | CDv5178 | HostAssoc_Human Feces | Human feces | Spain : Valencia | 39.468228 -0.378304 | reads ENA | 454 GS-FLX Titanium | Newbler | 0 | |
| PerezBrocal_CDViromes | CDv5180 | HostAssoc_Human Feces | Human feces | Spain : Valencia | 39.468228 -0.378304 | reads ENA | 454 GS-FLX Titanium | Newbler | 0 | |
| PerezBrocal_CDViromes | CDv5181 | HostAssoc_Human Feces | Human feces | Spain : Valencia | 39.468228 -0.378304 | reads ENA | 454 GS-FLX Titanium | Newbler | 0 | |
| PerezBrocal_CDViromes | CDv5182 | HostAssoc_Human Feces | Human feces | Spain : Valencia | 39.468228 -0.378304 | reads ENA | 454 GS-FLX Titanium | Newbler | 0 | |
| PerezBrocal_CDViromes | CDv5184 | HostAssoc_Human Feces | Human feces | Spain : Valencia | 39.468228 -0.378304 | reads ENA | 454 GS-FLX Titanium | Newbler | 0 | |
| PerezBrocal_CDViromes | CDv5185 | HostAssoc_Human Feces | Human feces | Spain : Valencia | 39.468228 -0.378304 | reads ENA | 454 GS-FLX Titanium | Newbler | 0 | |
| PerezBrocal_CDViromes | CDv5186 | HostAssoc_Human Feces | Human feces | Spain : Valencia | 39.468228 -0.378304 | reads ENA | 454 GS-FLX Titanium | Newbler | 0 | |
| PerezBrocal_CDViromes | CDv5188 | HostAssoc_Human Feces | Human feces | Spain : Valencia | 39.468228 -0.378304 | reads ENA | 454 GS-FLX Titanium | Newbler | 0 | |
| PerezBrocal_CDViromes | CDv5189 | HostAssoc_Human Feces | Human feces | Spain : Valencia | 39.468228 -0.378304 | reads ENA | 454 GS-FLX Titanium | Newbler | 0 | |
| PerezBrocal_CDViromes | CDv5190 | HostAssoc_Human Feces | Human feces | Spain : Valencia | 39.468228 -0.378304 | reads ENA | 454 GS-FLX Titanium | Newbler | 0 | |
| PerezBrocal_CDViromes | CDv5192 | HostAssoc_Human Feces | Human feces | Spain : Valencia | 39.468228 -0.378304 | reads ENA | 454 GS-FLX Titanium | Newbler | 0 | |
| PerezBrocal_CDViromes | CDv5193 | HostAssoc_Human Feces | Human feces | Spain : Valencia | 39.468228 -0.378304 | reads ENA | 454 GS-FLX Titanium | Newbler | 0 | |
| PerezBrocal_CDViromes | CDv5194 | HostAssoc_Human Feces | Human feces | Spain : Valencia | 39.468228 -0.378304 | reads ENA | 454 GS-FLX Titanium | Newbler | 0 | |
| PerezBrocal_CDViromes | CDv5196 | HostAssoc_Human Feces | Human feces | Spain : Valencia | 39.468228 -0.378304 | reads ENA | 454 GS-FLX Titanium | Newbler | 0 | |
| PerezBrocal_CDViromes | CDv5197 | HostAssoc_Human Feces | Human feces | Spain : Valencia | 39.468228 -0.378304 | reads ENA | 454 GS-FLX Titanium | Newbler | 0 | |
| PerezBrocal_CDViromes | CDv5198 | HostAssoc_Human Feces | Human feces | Spain : Valencia | 39.468228 -0.378304 | reads ENA | 454 GS-FLX Titanium | Newbler | 0 | |
| PerezBrocal_CDViromes | CDv5200 | HostAssoc_Human Feces | Human feces | Spain : Valencia | 39.468228 -0.378304 | reads ENA | 454 GS-FLX Titanium | Newbler | 0 | |
| PerezBrocal_CDViromes | CDv5202 | HostAssoc_Human Feces | Human feces | Spain : Valencia | 39.468228 -0.378304 | reads ENA | 454 GS-FLX Titanium | Newbler | 0 | |
| PerezBrocal_CDViromes | CDv5204 | HostAssoc_Human Feces | Human feces | Spain : Valencia | 39.468228 -0.378304 | reads ENA | 454 GS-FLX Titanium | Newbler | 0 | |
| PerezBrocal_CDViromes | CDv5205 | HostAssoc_Human Feces | Human feces | Spain : Valencia | 39.468228 -0.378304 | reads ENA | 454 GS-FLX Titanium | Newbler | 0 | |
| PerezBrocal_CDViromes | CDv5206 | HostAssoc_Human Feces | Human feces | Spain : Valencia | 39.468228 -0.378304 | reads ENA | 454 GS-FLX Titanium | Newbler | 0 | |
| PerezBrocal_CDViromes | CDv5208 | HostAssoc_Human Feces | Human feces | Spain : Valencia | 39.468228 -0.378304 | reads ENA | 454 GS-FLX Titanium | Newbler | 0 | |
| PerezBrocal_CDViromes | CDv5210 | HostAssoc_Human Feces | Human feces | Spain : Valencia | 39.468228 -0.378304 | reads ENA | 454 GS-FLX Titanium | Newbler | 0 | |
| PerezBrocal_CDViromes | CDv5212 | HostAssoc_Human Feces | Human feces | Spain : Valencia | 39.468228 -0.378304 | reads ENA | 454 GS-FLX Titanium | Newbler | 0 | |
| PerezBrocal_CDViromes | CDv5213 | HostAssoc_Human Feces | Human feces | Spain : Valencia | 39.468228 -0.378304 | reads ENA | 454 GS-FLX Titanium | Newbler | 0 | |
| PerezBrocal_CDViromes | CDv5214 | HostAssoc_Human Feces | Human feces | Spain : Valencia | 39.468228 -0.378304 | reads ENA | 454 GS-FLX Titanium | Newbler | 0 | |
| PerezBrocal_CDViromes | CDv5216 | HostAssoc_Human Feces | Human feces | Spain : Valencia | 39.468228 -0.378304 | reads ENA | 454 GS-FLX Titanium | Newbler | 0 | |
| PerezBrocal_CDViromes | CDv5217 | HostAssoc_Human Feces | Human feces | Spain : Valencia | 39.468228 -0.378304 | reads ENA | 454 GS-FLX Titanium | Newbler | 0 | |
| PerezBrocal_CDViromes | CDv5218 | HostAssoc_Human Feces | Human feces | Spain : Valencia | 39.468228 -0.378304 | reads ENA | 454 GS-FLX Titanium | Newbler | 0 | |
| PerezBrocal_CDViromes | CDv5220 | HostAssoc_Human Feces | Human feces | Spain : Valencia | 39.468228 -0.378304 | reads ENA | 454 GS-FLX Titanium | Newbler | 0 | |
| PerezBrocal_CDViromes | CDv5222 | HostAssoc_Human Feces | Human feces | Spain : Valencia | 39.468228 -0.378304 | reads ENA | 454 GS-FLX Titanium | Newbler | 0 | |
| PerezBrocal_CDViromes | CDv5224 | HostAssoc_Human Feces | Human feces | Spain : Valencia | 39.468228 -0.378304 | reads ENA | 454 GS-FLX Titanium | Newbler | 0 | |
| PerezBrocal_CDViromes | CDv5225 | HostAssoc_Human Feces | Human feces | Spain : Valencia | 39.468228 -0.378304 | reads ENA | 454 GS-FLX Titanium | Newbler | 0 | |
| PerezBrocal_CDViromes | CDv5226 | HostAssoc_Human Feces | Human feces | Spain : Valencia | 39.468228 -0.378304 | reads ENA | 454 GS-FLX Titanium | Newbler | 0 | |

| Project Name | Virome Code | Habitat Type (code) | Ecosystem Type (text) | Location | GPS | Source | Sequencing Technology | Assembly Software | # crucivirus | Reference |
|---|---|---|---|---|---|---|---|---|---|---|
| PerezBrocal_CDViromes | CDv5228 | HostAssoc_Human Feces | Human feces | Spain : Valencia | 39.468228 -0.378304 | reads ENA | 454 GS-FLX Titanium | Newbler | 0 | Pérez-Brocal V. García-López R. Nos P. Beltrán B. Moret I. Moya A. Metagenomic Analysis of Crohn's Disease Patients Identifies Changes in the Virome and Microbiome Related to Disease Status and Therapy. and Detects Potential Interactions and Biomarkers. Inflamm Bowel Dis. 2015 Nov;21(11):2515-32. doi: 10.1097/MIB. |
| PerezBrocal_CDViromes | CDv5229 | HostAssoc_Human Feces | Human feces | Spain : Valencia | 39.468228 -0.378304 | reads ENA | 454 GS-FLX Titanium | Newbler | 0 | |
| PerezBrocal_CDViromes | CDv5231 | HostAssoc_Human Feces | Human feces | Spain : Valencia | 39.468228 -0.378304 | reads ENA | 454 GS-FLX Titanium | Newbler | 0 | |
| PerezBrocal_CDViromes | CDv5232 | HostAssoc_Human Feces | Human feces | Spain : Valencia | 39.468228 -0.378304 | reads ENA | 454 GS-FLX Titanium | Newbler | 0 | |
| PerezBrocal_CDViromes | CDv5233 | HostAssoc_Human Feces | Human feces | Spain : Valencia | 39.468228 -0.378304 | reads ENA | 454 GS-FLX Titanium | Newbler | 0 | |
| PerezBrocal_CDViromes | CDv5235 | HostAssoc_Human Feces | Human feces | Spain : Valencia | 39.468228 -0.378304 | reads ENA | 454 GS-FLX Titanium | Newbler | 0 | |
| PerezBrocal_CDViromes | CDv5236 | HostAssoc_Human Feces | Human feces | Spain : Valencia | 39.468228 -0.378304 | reads ENA | 454 GS-FLX Titanium | Newbler | 0 | |
| PerezBrocal_CDViromes | CDv5237 | HostAssoc_Human Feces | Human feces | Spain : Valencia | 39.468228 -0.378304 | reads ENA | 454 GS-FLX Titanium | Newbler | 0 | |
| PerezBrocal_CDViromes | CDv5239 | HostAssoc_Human Feces | Human feces | Spain : Valencia | 39.468228 -0.378304 | reads ENA | 454 GS-FLX Titanium | Newbler | 0 | |
| PerezBrocal_CDViromes | CDv5240 | HostAssoc_Human Feces | Human feces | Spain : Valencia | 39.468228 -0.378304 | reads ENA | 454 GS-FLX Titanium | Newbler | 0 | |
| PerezBrocal_CDViromes | CDv5241 | HostAssoc_Human Feces | Human feces | Spain : Valencia | 39.468228 -0.378304 | reads ENA | 454 GS-FLX Titanium | Newbler | 0 | |
| PerezBrocal_CDViromes | CDv5243 | HostAssoc_Human Feces | Human feces | Spain : Valencia | 39.468228 -0.378304 | reads ENA | 454 GS-FLX Titanium | Newbler | 0 | |
| PerezBrocal_CDViromes | CDv5245 | HostAssoc_Human Feces | Human feces | Spain : Valencia | 39.468228 -0.378304 | reads ENA | 454 GS-FLX Titanium | Newbler | 0 | |
| PerezBrocal_CDViromes | CDv5247 | HostAssoc_Human Feces | Human feces | Spain : Valencia | 39.468228 -0.378304 | reads ENA | 454 GS-FLX Titanium | Newbler | 0 | |
| PerezBrocal_CDViromes | CDv5248 | HostAssoc_Human Feces | Human feces | Spain : Valencia | 39.468228 -0.378304 | reads ENA | 454 GS-FLX Titanium | Newbler | 0 | |
| PerezBrocal_CDViromes | CDv5249 | HostAssoc_Human Feces | Human feces | Spain : Valencia | 39.468228 -0.378304 | reads ENA | 454 GS-FLX Titanium | Newbler | 0 | |
| PerezBrocal_CDViromes | CDv5251 | HostAssoc_Human Feces | Human feces | Spain : Valencia | 39.468228 -0.378304 | reads ENA | 454 GS-FLX Titanium | Newbler | 0 | |
| PerezBrocal_CDViromes | CDv5252 | HostAssoc_Human Feces | Human feces | Spain : Valencia | 39.468228 -0.378304 | reads ENA | 454 GS-FLX Titanium | Newbler | 0 | |
| PerezBrocal_CDViromes | CDv5253 | HostAssoc_Human Feces | Human feces | Spain : Valencia | 39.468228 -0.378304 | reads ENA | 454 GS-FLX Titanium | Newbler | 0 | |
| PerezBrocal_CDViromes | CDv5255 | HostAssoc_Human Feces | Human feces | Spain : Valencia | 39.468228 -0.378304 | reads ENA | 454 GS-FLX Titanium | Newbler | 0 | |
| PerezBrocal_CDViromes | CDv5256 | HostAssoc_Human Feces | Human feces | Spain : Valencia | 39.468228 -0.378304 | reads ENA | 454 GS-FLX Titanium | Newbler | 0 | |
| PerezBrocal_CDViromes | CDv5257 | HostAssoc_Human Feces | Human feces | Spain : Valencia | 39.468228 -0.378304 | reads ENA | 454 GS-FLX Titanium | Newbler | 0 | |
| PerezBrocal_CDViromes | CDv5259 | HostAssoc_Human Feces | Human feces | Spain : Valencia | 39.468228 -0.378304 | reads ENA | 454 GS-FLX Titanium | Newbler | 0 | |
| PerezBrocal_CDViromes | CDv5260 | HostAssoc_Human Feces | Human feces | Spain : Valencia | 39.468228 -0.378304 | reads ENA | 454 GS-FLX Titanium | Newbler | 0 | |
| PerezBrocal_CDViromes | CDv5261 | HostAssoc_Human Feces | Human feces | Spain : Valencia | 39.468228 -0.378304 | reads ENA | 454 GS-FLX Titanium | Newbler | 0 | |
| PerezBrocal_CDViromes | CDv5263 | HostAssoc_Human Feces | Human feces | Spain : Valencia | 39.468228 -0.378304 | reads ENA | 454 GS-FLX Titanium | Newbler | 0 | |
| PerezBrocal_CDViromes | CDv5264 | HostAssoc_Human Feces | Human feces | Spain : Valencia | 39.468228 -0.378304 | reads ENA | 454 GS-FLX Titanium | Newbler | 0 | |
| PerezBrocal_CDViromes | CDv5265 | HostAssoc_Human Feces | Human feces | Spain : Valencia | 39.468228 -0.378304 | reads ENA | 454 GS-FLX Titanium | Newbler | 0 | |
| PerezBrocal_CDViromes | CDv5267 | HostAssoc_Human Feces | Human feces | Spain : Valencia | 39.468228 -0.378304 | reads ENA | 454 GS-FLX Titanium | Newbler | 0 | |
| PerezBrocal_CDViromes | CDv5268 | HostAssoc_Human Feces | Human feces | Spain : Valencia | 39.468228 -0.378304 | reads ENA | 454 GS-FLX Titanium | Newbler | 0 | |
| PerezBrocal_CDViromes | CDv5269 | HostAssoc_Human Feces | Human feces | Spain : Valencia | 39.468228 -0.378304 | reads ENA | 454 GS-FLX Titanium | Newbler | 0 | |
| PerezBrocal_CDViromes | CDv5271 | HostAssoc_Human Feces | Human feces | Spain : Valencia | 39.468228 -0.378304 | reads ENA | 454 GS-FLX Titanium | Newbler | 0 | |
| PerezBrocal_CDViromes | CDv5272 | HostAssoc_Human Feces | Human feces | Spain : Valencia | 39.468228 -0.378304 | reads ENA | 454 GS-FLX Titanium | Newbler | 0 | |
| PerezBrocal_CDViromes | CDv5273 | HostAssoc_Human Feces | Human feces | Spain : Valencia | 39.468228 -0.378304 | reads ENA | 454 GS-FLX Titanium | Newbler | 0 | |
| PerezBrocal_CDViromes | CDv5275 | HostAssoc_Human Feces | Human feces | Spain : Valencia | 39.468228 -0.378304 | reads ENA | 454 GS-FLX Titanium | Newbler | 0 | |
| PerezBrocal_CDViromes | CDv5276 | HostAssoc_Human Feces | Human feces | Spain : Valencia | 39.468228 -0.378304 | reads ENA | 454 GS-FLX Titanium | Newbler | 0 | |
| PerezBrocal_CDViromes | CDv5277 | HostAssoc_Human Feces | Human feces | Spain : Valencia | 39.468228 -0.378304 | reads ENA | 454 GS-FLX Titanium | Newbler | 0 | |
| PerezBrocal_CDViromes | CDv5279 | HostAssoc_Human Feces | Human feces | Spain : Valencia | 39.468228 -0.378304 | reads ENA | 454 GS-FLX Titanium | Newbler | 0 | |
| PerezBrocal_CDViromes | CDv5280 | HostAssoc_Human Feces | Human feces | Spain : Valencia | 39.468228 -0.378304 | reads ENA | 454 GS-FLX Titanium | Newbler | 0 | |
| PerezBrocal_CDViromes | CDv5281 | HostAssoc_Human Feces | Human feces | Spain : Valencia | 39.468228 -0.378304 | reads ENA | 454 GS-FLX Titanium | Newbler | 0 | |
| PerezBrocal_CDViromes | CDv5283 | HostAssoc_Human Feces | Human feces | Spain : Valencia | 39.468228 -0.378304 | reads ENA | 454 GS-FLX Titanium | Newbler | 0 | |
| PerezBrocal_CDViromes | CDv5284 | HostAssoc_Human Feces | Human feces | Spain : Valencia | 39.468228 -0.378304 | reads ENA | 454 GS-FLX Titanium | Newbler | 0 | |

| Project Name | Virome Code | Habitat Type (code) | Ecosystem Type (text) | Location | GPS | Source | Sequencing Technology | Assembly Software | # crucivirus | Reference |
|---|---|---|---|---|---|---|---|---|---|---|
| PerezBrocal_CDViromes | CDv5285 | HostAssoc_Human Feces | Human feces | Spain : Valencia | 39.468228 -0.378304 | reads ENA | 454 GS-FLX Titanium | Newbler | 0 | |
| PerezBrocal_CDViromes | CDv5287 | HostAssoc_Human Feces | Human feces | Spain : Valencia | 39.468228 -0.378304 | reads ENA | 454 GS-FLX Titanium | Newbler | 0 | |
| PerezBrocal_CDViromes | CDv5288 | HostAssoc_Human Feces | Human feces | Spain : Valencia | 39.468228 -0.378304 | reads ENA | 454 GS-FLX Titanium | Newbler | 0 | |
| PerezBrocal_CDViromes | CDv5289 | HostAssoc_Human Feces | Human feces | Spain : Valencia | 39.468228 -0.378304 | reads ENA | 454 GS-FLX Titanium | Newbler | 0 | |
| PerezBrocal_CDViromes | CDv5291 | HostAssoc_Human Feces | Human feces | Spain : Valencia | 39.468228 -0.378304 | reads ENA | 454 GS-FLX Titanium | Newbler | 0 | |
| PerezBrocal_CDViromes | CDv5292 | HostAssoc_Human Feces | Human feces | Spain : Valencia | 39.468228 -0.378304 | reads ENA | 454 GS-FLX Titanium | Newbler | 0 | |
| PerezBrocal_CDViromes | CDv5293 | HostAssoc_Human Feces | Human feces | Spain : Valencia | 39.468228 -0.378304 | reads ENA | 454 GS-FLX Titanium | Newbler | 0 | |
| PerezBrocal_CDViromes | CDv5295 | HostAssoc_Human Feces | Human feces | Spain : Valencia | 39.468228 -0.378304 | reads ENA | 454 GS-FLX Titanium | Newbler | 0 | |
| PerezBrocal_CDViromes | CDv5296 | HostAssoc_Human Feces | Human feces | Spain : Valencia | 39.468228 -0.378304 | reads ENA | 454 GS-FLX Titanium | Newbler | 0 | |
| PerezBrocal_CDViromes | CDv5298 | HostAssoc_Human Feces | Human feces | Spain : Valencia | 39.468228 -0.378304 | reads ENA | 454 GS-FLX Titanium | Newbler | 0 | |
| PerezBrocal_CDViromes | CDv5299 | HostAssoc_Human Feces | Human feces | Spain : Valencia | 39.468228 -0.378304 | reads ENA | 454 GS-FLX Titanium | Newbler | 0 | |
| PerezBrocal_CDViromes | CDv5300 | HostAssoc_Human Feces | Human feces | Spain : Valencia | 39.468228 -0.378304 | reads ENA | 454 GS-FLX Titanium | Newbler | 0 | |
| PerezBrocal_CDViromes | CDv5302 | HostAssoc_Human Feces | Human feces | Spain : Valencia | 39.468228 -0.378304 | reads ENA | 454 GS-FLX Titanium | Newbler | 0 | |
| PerezBrocal_CDViromes | CDv5303 | HostAssoc_Human Feces | Human feces | Spain : Valencia | 39.468228 -0.378304 | reads ENA | 454 GS-FLX Titanium | Newbler | 0 | |
| PerezBrocal_CDViromes | CDv5304 | HostAssoc_Human Feces | Human feces | Spain : Valencia | 39.468228 -0.378304 | reads ENA | 454 GS-FLX Titanium | Newbler | 0 | |
| PerezBrocal_CDViromes | CDv5306 | HostAssoc_Human Feces | Human feces | Spain : Valencia | 39.468228 -0.378304 | reads ENA | 454 GS-FLX Titanium | Newbler | 0 | |
| PerezBrocal_CDViromes | CDv5307 | HostAssoc_Human Feces | Human feces | Spain : Valencia | 39.468228 -0.378304 | reads ENA | 454 GS-FLX Titanium | Newbler | 0 | |
| PerezBrocal_CDViromes | CDv5308 | HostAssoc_Human Feces | Human feces | Spain : Valencia | 39.468228 -0.378304 | reads ENA | 454 GS-FLX Titanium | Newbler | 0 | |
| PerezBrocal_CDViromes | CDv5310 | HostAssoc_Human Feces | Human feces | Spain : Valencia | 39.468228 -0.378304 | reads ENA | 454 GS-FLX Titanium | Newbler | 0 | |
| PerezBrocal_CDViromes | CDv5312 | HostAssoc_Human Feces | Human feces | Spain : Valencia | 39.468228 -0.378304 | reads ENA | 454 GS-FLX Titanium | Newbler | 0 | |
| PerezBrocal_CDViromes | CDv5314 | HostAssoc_Human Feces | Human feces | Spain : Valencia | 39.468228 -0.378304 | reads ENA | 454 GS-FLX Titanium | Newbler | 0 | |
| PerezBrocal_CDViromes | CDv5315 | HostAssoc_Human Feces | Human feces | Spain : Valencia | 39.468228 -0.378304 | reads ENA | 454 GS-FLX Titanium | Newbler | 0 | |
| PerezBrocal_CDViromes | CDv5316 | HostAssoc_Human Feces | Human feces | Spain : Valencia | 39.468228 -0.378304 | reads ENA | 454 GS-FLX Titanium | Newbler | 0 | |
| PerezBrocal_CDViromes | CDv5318 | HostAssoc_Human Feces | Human feces | Spain : Valencia | 39.468228 -0.378304 | reads ENA | 454 GS-FLX Titanium | Newbler | 0 | |
| PerezBrocal_CDViromes | CDv5319 | HostAssoc_Human Feces | Human feces | Spain : Valencia | 39.468228 -0.378304 | reads ENA | 454 GS-FLX Titanium | Newbler | 0 | |
| PerezBrocal_CDViromes | CDv5320 | HostAssoc_Human Feces | Human feces | Spain : Valencia | 39.468228 -0.378304 | reads ENA | 454 GS-FLX Titanium | Newbler | 0 | |
| PerezBrocal_CDViromes | CDv5322 | HostAssoc_Human Feces | Human feces | Spain : Valencia | 39.468228 -0.378304 | reads ENA | 454 GS-FLX Titanium | Newbler | 0 | |
| PerezBrocal_CDViromes | CDv5323 | HostAssoc_Human Feces | Human feces | Spain : Valencia | 39.468228 -0.378304 | reads ENA | 454 GS-FLX Titanium | Newbler | 0 | |
| PerezBrocal_CDViromes | CDv5324 | HostAssoc_Human Feces | Human feces | Spain : Valencia | 39.468228 -0.378304 | reads ENA | 454 GS-FLX Titanium | Newbler | 0 | |
| PerezBrocal_CDViromes | CDv5326 | HostAssoc_Human Feces | Human feces | Spain : Valencia | 39.468228 -0.378304 | reads ENA | 454 GS-FLX Titanium | Newbler | 0 | |
| PerezBrocal_CDViromes | CDv5327 | HostAssoc_Human Feces | Human feces | Spain : Valencia | 39.468228 -0.378304 | reads ENA | 454 GS-FLX Titanium | Newbler | 0 | |
| PerezBrocal_CDViromes | CDv5328 | HostAssoc_Human Feces | Human feces | Spain : Valencia | 39.468228 -0.378304 | reads ENA | 454 GS-FLX Titanium | Newbler | 0 | |
| PerezBrocal_CDViromes | CDv5330 | HostAssoc_Human Feces | Human feces | Spain : Valencia | 39.468228 -0.378304 | reads ENA | 454 GS-FLX Titanium | Newbler | 0 | |
| PerezBrocal_CDViromes | CDv5331 | HostAssoc_Human Feces | Human feces | Spain : Valencia | 39.468228 -0.378304 | reads ENA | 454 GS-FLX Titanium | Newbler | 0 | |
| PerezBrocal_CDViromes | CDv5332 | HostAssoc_Human Feces | Human feces | Spain : Valencia | 39.468228 -0.378304 | reads ENA | 454 GS-FLX Titanium | Newbler | 0 | |
| PerezBrocal_CDViromes | CDv5334 | HostAssoc_Human Feces | Human feces | Spain : Valencia | 39.468228 -0.378304 | reads ENA | 454 GS-FLX Titanium | Newbler | 0 | |
| PerezBrocal_CDViromes | CDv5336 | HostAssoc_Human Feces | Human feces | Spain : Valencia | 39.468228 -0.378304 | reads ENA | 454 GS-FLX Titanium | Newbler | 0 | |
| Pride_LongTermAntibio | ELA100A | HostAssoc_Human Feces | Human feces | USA : California | 32.876609 -117.236885 | reads MG-RAST | Ion Torrent | Newbler -large | 0 | |
| Pride_LongTermAntibio | ELA100B | HostAssoc_Human Feces | Human feces | USA : California | 32.876609 -117.236885 | reads MG-RAST | Ion Torrent | Newbler -large | 0 | |
| Pride_LongTermAntibio | ELA100C | HostAssoc_Human Feces | Human feces | USA : California | 32.876609 -117.236885 | reads MG-RAST | Ion Torrent | Newbler -large | 0 | |
| Pride_LongTermAntibio | ELA1A | HostAssoc_Human Feces | Human feces | USA : California | 32.876609 -117.236885 | reads MG-RAST | Ion Torrent | Newbler -large | 0 | |
| Pride_LongTermAntibio | ELA1B | HostAssoc_Human Feces | Human feces | USA : California | 32.876609 -117.236885 | reads MG-RAST | Ion Torrent | Newbler -large | 0 | |

| Project Name | Virome Code | Habitat Type (code) | Ecosystem Type (text) | Location | GPS | Source | Sequencing Technology | Assembly Software | # crucivirus | Reference |
|---|---|---|---|---|---|---|---|---|---|---|
| Pride_LongTermAntibio | ELA1C | HostAssoc_Human Feces | Human feces | USA : California | 32.876609 -117.236885 | reads MG-RAST | Ion Torrent | Newbler -large | 0 | Abeles SR. Ly M. Santiago-Rodriguez TM. Pride DT. Effects of Long Term Antibiotic Therapy on Human Oral and Fecal Viromes. PLoS One. 2015 Aug 26;10(8):e0134941. doi: 10.1371/journal.pone.0134941. eCollection 2015. |
| Pride_LongTermAntibio | ELA2B | HostAssoc_Human Feces | Human feces | USA : California | 32.876609 -117.236885 | reads MG-RAST | Ion Torrent | Newbler -large | 0 | |
| Pride_LongTermAntibio | ELA2C | HostAssoc_Human Feces | Human feces | USA : California | 32.876609 -117.236885 | reads MG-RAST | Ion Torrent | Newbler -large | 0 | |
| Pride_LongTermAntibio | ELA33A | HostAssoc_Human Feces | Human feces | USA : California | 32.876609 -117.236885 | reads MG-RAST | Ion Torrent | Newbler -large | 0 | |
| Pride_LongTermAntibio | ELA33B | HostAssoc_Human Feces | Human feces | USA : California | 32.876609 -117.236885 | reads MG-RAST | Ion Torrent | Newbler -large | 0 | |
| Pride_LongTermAntibio | ELA33C | HostAssoc_Human Feces | Human feces | USA : California | 32.876609 -117.236885 | reads MG-RAST | Ion Torrent | Newbler -large | 0 | |
| Pride_LongTermAntibio | ELA3A | HostAssoc_Human Feces | Human feces | USA : California | 32.876609 -117.236885 | reads MG-RAST | Ion Torrent | Newbler -large | 0 | |
| Pride_LongTermAntibio | ELA3B | HostAssoc_Human Feces | Human feces | USA : California | 32.876609 -117.236885 | reads MG-RAST | Ion Torrent | Newbler -large | 0 | |
| Pride_LongTermAntibio | ELA3C | HostAssoc_Human Feces | Human feces | USA : California | 32.876609 -117.236885 | reads MG-RAST | Ion Torrent | Newbler -large | 0 | |
| Pride_LongTermAntibio | ELA4A | HostAssoc_Human Feces | Human feces | USA : California | 32.876609 -117.236885 | reads MG-RAST | Ion Torrent | Newbler -large | 0 | |
| Pride_LongTermAntibio | ELA4B | HostAssoc_Human Feces | Human feces | USA : California | 32.876609 -117.236885 | reads MG-RAST | Ion Torrent | Newbler -large | 0 | |
| Pride_LongTermAntibio | ELA4C | HostAssoc_Human Feces | Human feces | USA : California | 32.876609 -117.236885 | reads MG-RAST | Ion Torrent | Newbler -large | 0 | |
| Pride_LongTermAntibio | ELA7A | HostAssoc_Human Feces | Human feces | USA : California | 32.876609 -117.236885 | reads MG-RAST | Ion Torrent | Newbler -large | 0 | |
| Pride_LongTermAntibio | ELA7B | HostAssoc_Human Feces | Human feces | USA : California | 32.876609 -117.236885 | reads MG-RAST | Ion Torrent | Newbler -large | 0 | |
| Pride_LongTermAntibio | ELA7C | HostAssoc_Human Feces | Human feces | USA : California | 32.876609 -117.236885 | reads MG-RAST | Ion Torrent | Newbler -large | 0 | |
| Pride_LongTermAntibio | ELA8A | HostAssoc_Human Feces | Human feces | USA : California | 32.876609 -117.236885 | reads MG-RAST | Ion Torrent | Newbler -large | 0 | |
| Pride_LongTermAntibio | ELA8B | HostAssoc_Human Feces | Human feces | USA : California | 32.876609 -117.236885 | reads MG-RAST | Ion Torrent | Newbler -large | 0 | |
| Pride_LongTermAntibio | ELA8C | HostAssoc_Human Feces | Human feces | USA : California | 32.876609 -117.236885 | reads MG-RAST | Ion Torrent | Newbler -large | 0 | |
| Pride_LongTermAntibio | ELA9A | HostAssoc_Human Feces | Human feces | USA : California | 32.876609 -117.236885 | reads MG-RAST | Ion Torrent | Newbler -large | 0 | |
| Pride_LongTermAntibio | ELA9B | HostAssoc_Human Feces | Human feces | USA : California | 32.876609 -117.236885 | reads MG-RAST | Ion Torrent | Newbler -large | 0 | |
| Pride_LongTermAntibio | ELA9C | HostAssoc_Human Feces | Human feces | USA : California | 32.876609 -117.236885 | reads MG-RAST | Ion Torrent | Newbler -large | 0 | |
| Reyes_Malnutrition | Malnut4990 | HostAssoc_Human Feces | Human feces | Malawi | -13.754313 33.929313 | reads ENA | Illumina MiSeq | Newbler | 0 | |
| Reyes_Malnutrition | Malnut4991 | HostAssoc_Human Feces | Human feces | Malawi | -13.754313 33.929313 | reads ENA | Illumina MiSeq | Newbler | 0 | |
| Reyes_Malnutrition | Malnut4992 | HostAssoc_Human Feces | Human feces | Malawi | -13.754313 33.929313 | reads ENA | Illumina MiSeq | Newbler | 0 | |
| Reyes_Malnutrition | Malnut4993 | HostAssoc_Human Feces | Human feces | Malawi | -13.754313 33.929313 | reads ENA | Illumina MiSeq | Newbler | 0 | |
| Reyes_Malnutrition | Malnut4994 | HostAssoc_Human Feces | Human feces | Malawi | -13.754313 33.929313 | reads ENA | Illumina MiSeq | Newbler | 0 | |
| Reyes_Malnutrition | Malnut4995 | HostAssoc_Human Feces | Human feces | Malawi | -13.754313 33.929313 | reads ENA | Illumina MiSeq | Newbler | 0 | |
| Reyes_Malnutrition | Malnut4996 | HostAssoc_Human Feces | Human feces | Malawi | -13.754313 33.929313 | reads ENA | Illumina MiSeq | Newbler | 0 | |
| Reyes_Malnutrition | Malnut4997 | HostAssoc_Human Feces | Human feces | Malawi | -13.754313 33.929313 | reads ENA | Illumina MiSeq | Newbler | 0 | |
| Reyes_Malnutrition | Malnut4998 | HostAssoc_Human Feces | Human feces | Malawi | -13.754313 33.929313 | reads ENA | Illumina MiSeq | Newbler | 0 | |
| Reyes_Malnutrition | Malnut4999 | HostAssoc_Human Feces | Human feces | Malawi | -13.754313 33.929313 | reads ENA | Illumina MiSeq | Newbler | 0 | |
| Reyes_Malnutrition | Malnut5000 | HostAssoc_Human Feces | Human feces | Malawi | -13.754313 33.929313 | reads ENA | Illumina MiSeq | Newbler | 0 | |
| Reyes_Malnutrition | Malnut5002 | HostAssoc_Human Feces | Human feces | Malawi | -13.754313 33.929313 | reads ENA | Illumina MiSeq | Newbler | 0 | |
| Reyes_Malnutrition | Malnut5003 | HostAssoc_Human Feces | Human feces | Malawi | -13.754313 33.929313 | reads ENA | Illumina MiSeq | Newbler | 0 | |
| Reyes_Malnutrition | Malnut5004 | HostAssoc_Human Feces | Human feces | Malawi | -13.754313 33.929313 | reads ENA | Illumina MiSeq | Newbler | 0 | |
| Reyes_Malnutrition | Malnut5005 | HostAssoc_Human Feces | Human feces | Malawi | -13.754313 33.929313 | reads ENA | Illumina MiSeq | Newbler | 0 | |
| Reyes_Malnutrition | Malnut5006 | HostAssoc_Human Feces | Human feces | Malawi | -13.754313 33.929313 | reads ENA | Illumina MiSeq | Newbler | 0 | |
| Reyes_Malnutrition | Malnut5007 | HostAssoc_Human Feces | Human feces | Malawi | -13.754313 33.929313 | reads ENA | Illumina MiSeq | Newbler | 0 | |
| Reyes_Malnutrition | Malnut5008 | HostAssoc_Human Feces | Human feces | Malawi | -13.754313 33.929313 | reads ENA | Illumina MiSeq | Newbler | 0 | |
| Reyes_Malnutrition | Malnut5009 | HostAssoc_Human Feces | Human feces | Malawi | -13.754313 33.929313 | reads ENA | Illumina MiSeq | Newbler | 0 | |
| Reyes_Malnutrition | Malnut5010 | HostAssoc_Human Feces | Human feces | Malawi | -13.754313 33.929313 | reads ENA | Illumina MiSeq | Newbler | 0 | |
| Reyes_Malnutrition | Malnut5011 | HostAssoc_Human Feces | Human feces | Malawi | -13.754313 33.929313 | reads ENA | Illumina MiSeq | Newbler | 0 | |

| Project Name | Virome Code | Habitat Type (code) | Ecosystem Type (text) | Location | GPS | Source | Sequencing Technology | Assembly Software | # crucivirus | Reference |
|---|---|---|---|---|---|---|---|---|---|---|
| Reyes_Malnutrition | Malnut5012 | HostAssoc_Human Feces | Human feces | Malawi | -13.754313 33.929313 | reads ENA | Illumina MiSeq | Newbler | 0 | |
| Reyes_Malnutrition | Malnut5013 | HostAssoc_Human Feces | Human feces | Malawi | -13.754313 33.929313 | reads ENA | Illumina MiSeq | Newbler | 0 | |
| Reyes_Malnutrition | Malnut5014 | HostAssoc_Human Feces | Human feces | Malawi | -13.754313 33.929313 | reads ENA | Illumina MiSeq | Newbler | 0 | |
| Reyes_Malnutrition | Malnut5015 | HostAssoc_Human Feces | Human feces | Malawi | -13.754313 33.929313 | reads ENA | Illumina MiSeq | Newbler | 0 | |
| Reyes_Malnutrition | Malnut5016 | HostAssoc_Human Feces | Human feces | Malawi | -13.754313 33.929313 | reads ENA | Illumina MiSeq | Newbler | 0 | |
| Reyes_Malnutrition | Malnut5017 | HostAssoc_Human Feces | Human feces | Malawi | -13.754313 33.929313 | reads ENA | Illumina MiSeq | Newbler | 0 | |
| Reyes_Malnutrition | Malnut5018 | HostAssoc_Human Feces | Human feces | Malawi | -13.754313 33.929313 | reads ENA | Illumina MiSeq | Newbler | 0 | |
| Reyes_Malnutrition | Malnut5019 | HostAssoc_Human Feces | Human feces | Malawi | -13.754313 33.929313 | reads ENA | Illumina MiSeq | Newbler | 0 | |
| Reyes_Malnutrition | Malnut5020 | HostAssoc_Human Feces | Human feces | Malawi | -13.754313 33.929313 | reads ENA | Illumina MiSeq | Newbler | 0 | |
| Reyes_Malnutrition | Malnut5021 | HostAssoc_Human Feces | Human feces | Malawi | -13.754313 33.929313 | reads ENA | Illumina MiSeq | Newbler | 0 | |
| Reyes_Malnutrition | Malnut5022 | HostAssoc_Human Feces | Human feces | Malawi | -13.754313 33.929313 | reads ENA | Illumina MiSeq | Newbler | 0 | |
| Reyes_Malnutrition | Malnut5023 | HostAssoc_Human Feces | Human feces | Malawi | -13.754313 33.929313 | reads ENA | Illumina MiSeq | Newbler | 0 | |
| Reyes_Malnutrition | Malnut5024 | HostAssoc_Human Feces | Human feces | Malawi | -13.754313 33.929313 | reads ENA | Illumina MiSeq | Newbler | 0 | |
| Reyes_Malnutrition | Malnut5025 | HostAssoc_Human Feces | Human feces | Malawi | -13.754313 33.929313 | reads ENA | Illumina MiSeq | Newbler | 0 | |
| Reyes_Malnutrition | Malnut5026 | HostAssoc_Human Feces | Human feces | Malawi | -13.754313 33.929313 | reads ENA | Illumina MiSeq | Newbler | 0 | |
| Reyes_Malnutrition | Malnut5027 | HostAssoc_Human Feces | Human feces | Malawi | -13.754313 33.929313 | reads ENA | Illumina MiSeq | Newbler | 0 | |
| Reyes_Malnutrition | Malnut5028 | HostAssoc_Human Feces | Human feces | Malawi | -13.754313 33.929313 | reads ENA | Illumina MiSeq | Newbler | 0 | |
| Reyes_Malnutrition | Malnut5029 | HostAssoc_Human Feces | Human feces | Malawi | -13.754313 33.929313 | reads ENA | Illumina MiSeq | Newbler | 0 | |
| Reyes_Malnutrition | Malnut5030 | HostAssoc_Human Feces | Human feces | Malawi | -13.754313 33.929313 | reads ENA | Illumina MiSeq | Newbler | 0 | |
| Reyes_Malnutrition | Malnut5031 | HostAssoc_Human Feces | Human feces | Malawi | -13.754313 33.929313 | reads ENA | Illumina MiSeq | Newbler | 0 | |
| Reyes_Malnutrition | Malnut5032 | HostAssoc_Human Feces | Human feces | Malawi | -13.754313 33.929313 | reads ENA | Illumina MiSeq | Newbler | 0 | |
| Reyes_Malnutrition | Malnut5033 | HostAssoc_Human Feces | Human feces | Malawi | -13.754313 33.929313 | reads ENA | Illumina MiSeq | Newbler | 0 | |
| Reyes_Malnutrition | Malnut5034 | HostAssoc_Human Feces | Human feces | Malawi | -13.754313 33.929313 | reads ENA | Illumina MiSeq | Newbler | 0 | |
| Reyes_Malnutrition | Malnut5035 | HostAssoc_Human Feces | Human feces | Malawi | -13.754313 33.929313 | reads ENA | Illumina MiSeq | Newbler | 0 | |
| Reyes_Malnutrition | Malnut5036 | HostAssoc_Human Feces | Human feces | Malawi | -13.754313 33.929313 | reads ENA | Illumina MiSeq | Newbler | 0 | |
| Reyes_Malnutrition | Malnut5037 | HostAssoc_Human Feces | Human feces | Malawi | -13.754313 33.929313 | reads ENA | Illumina MiSeq | Newbler | 0 | |
| Reyes_Malnutrition | Malnut5038 | HostAssoc_Human Feces | Human feces | Malawi | -13.754313 33.929313 | reads ENA | Illumina MiSeq | Newbler | 0 | |
| Reyes_Malnutrition | Malnut5039 | HostAssoc_Human Feces | Human feces | Malawi | -13.754313 33.929313 | reads ENA | Illumina MiSeq | Newbler | 0 | |
| Reyes_Malnutrition | Malnut5040 | HostAssoc_Human Feces | Human feces | Malawi | -13.754313 33.929313 | reads ENA | Illumina MiSeq | Newbler | 0 | |
| Reyes_Malnutrition | Malnut5041 | HostAssoc_Human Feces | Human feces | Malawi | -13.754313 33.929313 | reads ENA | Illumina MiSeq | Newbler | 0 | |
| Reyes_Malnutrition | Malnut5042 | HostAssoc_Human Feces | Human feces | Malawi | -13.754313 33.929313 | reads ENA | Illumina MiSeq | Newbler | 0 | |
| Reyes_Malnutrition | Malnut5043 | HostAssoc_Human Feces | Human feces | Malawi | -13.754313 33.929313 | reads ENA | Illumina MiSeq | Newbler | 0 | |
| Reyes_Malnutrition | Malnut5044 | HostAssoc_Human Feces | Human feces | Malawi | -13.754313 33.929313 | reads ENA | Illumina MiSeq | Newbler | 0 | |
| Reyes_Malnutrition | Malnut5045 | HostAssoc_Human Feces | Human feces | Malawi | -13.754313 33.929313 | reads ENA | Illumina MiSeq | Newbler | 0 | |
| Reyes_Malnutrition | Malnut5046 | HostAssoc_Human Feces | Human feces | Malawi | -13.754313 33.929313 | reads ENA | Illumina MiSeq | Newbler | 0 | |
| Reyes_Malnutrition | Malnut5047 | HostAssoc_Human Feces | Human feces | Malawi | -13.754313 33.929313 | reads ENA | Illumina MiSeq | Newbler | 0 | |
| Reyes_Malnutrition | Malnut5048 | HostAssoc_Human Feces | Human feces | Malawi | -13.754313 33.929313 | reads ENA | Illumina MiSeq | Newbler | 0 | |
| Reyes_Malnutrition | Malnut5049 | HostAssoc_Human Feces | Human feces | Malawi | -13.754313 33.929313 | reads ENA | Illumina MiSeq | Newbler | 0 | |
| Reyes_Malnutrition | Malnut5050 | HostAssoc_Human Feces | Human feces | Malawi | -13.754313 33.929313 | reads ENA | Illumina MiSeq | Newbler | 0 | |
| Reyes_Malnutrition | Malnut5051 | HostAssoc_Human Feces | Human feces | Malawi | -13.754313 33.929313 | reads ENA | Illumina MiSeq | Newbler | 0 | |
| Reyes_Malnutrition | Malnut5052 | HostAssoc_Human Feces | Human feces | Malawi | -13.754313 33.929313 | reads ENA | Illumina MiSeq | Newbler | 0 | |
| Reyes_Malnutrition | Malnut5053 | HostAssoc_Human Feces | Human feces | Malawi | -13.754313 33.929313 | reads ENA | Illumina MiSeq | Newbler | 0 | |

| Project Name | Virome Code | Habitat Type (code) | Ecosystem Type (text) | Location | GPS | Source | Sequencing Technology | Assembly Software | # crucivirus | Reference |
|---|---|---|---|---|---|---|---|---|---|---|
| Reyes_Malnutrition | Malnut5054 | HostAssoc_Human Feces | Human feces | Malawi | -13.754313 33.929313 | reads ENA | Illumina MiSeq | Newbler | 0 | |
| Reyes_Malnutrition | Malnut5055 | HostAssoc_Human Feces | Human feces | Malawi | -13.754313 33.929313 | reads ENA | Illumina MiSeq | Newbler | 0 | |
| Reyes_Malnutrition | Malnut5056 | HostAssoc_Human Feces | Human feces | Malawi | -13.754313 33.929313 | reads ENA | Illumina MiSeq | Newbler | 0 | |
| Reyes_Malnutrition | Malnut5057 | HostAssoc_Human Feces | Human feces | Malawi | -13.754313 33.929313 | reads ENA | Illumina MiSeq | Newbler | 0 | |
| Reyes_Malnutrition | Malnut5058 | HostAssoc_Human Feces | Human feces | Malawi | -13.754313 33.929313 | reads ENA | Illumina MiSeq | Newbler | 0 | |
| Reyes_Malnutrition | Malnut5059 | HostAssoc_Human Feces | Human feces | Malawi | -13.754313 33.929313 | reads ENA | Illumina MiSeq | Newbler | 0 | |
| Reyes_Malnutrition | Malnut5060 | HostAssoc_Human Feces | Human feces | Malawi | -13.754313 33.929313 | reads ENA | Illumina MiSeq | Newbler | 0 | |
| Reyes_Malnutrition | Malnut5061 | HostAssoc_Human Feces | Human feces | Malawi | -13.754313 33.929313 | reads ENA | Illumina MiSeq | Newbler | 0 | |
| Reyes_Malnutrition | Malnut5062 | HostAssoc_Human Feces | Human feces | Malawi | -13.754313 33.929313 | reads ENA | Illumina MiSeq | Newbler | 0 | |
| Reyes_Malnutrition | Malnut5063 | HostAssoc_Human Feces | Human feces | Malawi | -13.754313 33.929313 | reads ENA | Illumina MiSeq | Newbler | 0 | |
| Reyes_Malnutrition | Malnut5064 | HostAssoc_Human Feces | Human feces | Malawi | -13.754313 33.929313 | reads ENA | Illumina MiSeq | Newbler | 0 | |
| Reyes_Malnutrition | Malnut5065 | HostAssoc_Human Feces | Human feces | Malawi | -13.754313 33.929313 | reads ENA | Illumina MiSeq | Newbler | 0 | |
| Reyes_Malnutrition | Malnut5066 | HostAssoc_Human Feces | Human feces | Malawi | -13.754313 33.929313 | reads ENA | Illumina MiSeq | Newbler | 0 | |
| Reyes_Malnutrition | Malnut5067 | HostAssoc_Human Feces | Human feces | Malawi | -13.754313 33.929313 | reads ENA | Illumina MiSeq | Newbler | 0 | |
| Reyes_Malnutrition | Malnut5068 | HostAssoc_Human Feces | Human feces | Malawi | -13.754313 33.929313 | reads ENA | Illumina MiSeq | Newbler | 0 | |
| Reyes_Malnutrition | Malnut5069 | HostAssoc_Human Feces | Human feces | Malawi | -13.754313 33.929313 | reads ENA | Illumina MiSeq | Newbler | 0 | |
| Reyes_Malnutrition | Malnut5070 | HostAssoc_Human Feces | Human feces | Malawi | -13.754313 33.929313 | reads ENA | Illumina MiSeq | Newbler | 0 | |
| Reyes_Malnutrition | Malnut5071 | HostAssoc_Human Feces | Human feces | Malawi | -13.754313 33.929313 | reads ENA | Illumina MiSeq | Newbler | 0 | |
| Reyes_Malnutrition | Malnut5072 | HostAssoc_Human Feces | Human feces | Malawi | -13.754313 33.929313 | reads ENA | Illumina MiSeq | Newbler | 0 | |
| Reyes_Malnutrition | Malnut5073 | HostAssoc_Human Feces | Human feces | Malawi | -13.754313 33.929313 | reads ENA | Illumina MiSeq | Newbler | 0 | |
| Reyes_Malnutrition | Malnut5074 | HostAssoc_Human Feces | Human feces | Malawi | -13.754313 33.929313 | reads ENA | Illumina MiSeq | Newbler | 0 | |
| Reyes_Malnutrition | Malnut5075 | HostAssoc_Human Feces | Human feces | Malawi | -13.754313 33.929313 | reads ENA | Illumina MiSeq | Newbler | 0 | |
| Reyes_Malnutrition | Malnut5076 | HostAssoc_Human Feces | Human feces | Malawi | -13.754313 33.929313 | reads ENA | Illumina MiSeq | Newbler | 0 | |
| Reyes_Malnutrition | Malnut5077 | HostAssoc_Human Feces | Human feces | Malawi | -13.754313 33.929313 | reads ENA | Illumina MiSeq | Newbler | 0 | |
| Reyes_Malnutrition | Malnut5078 | HostAssoc_Human Feces | Human feces | Malawi | -13.754313 33.929313 | reads ENA | Illumina MiSeq | Newbler | 0 | |
| Reyes_Malnutrition | Malnut5079 | HostAssoc_Human Feces | Human feces | Malawi | -13.754313 33.929313 | reads ENA | Illumina MiSeq | Newbler | 0 | |
| Reyes_Malnutrition | Malnut5080 | HostAssoc_Human Feces | Human feces | Malawi | -13.754313 33.929313 | reads ENA | Illumina MiSeq | Newbler | 0 | |
| Reyes_Malnutrition | Malnut5081 | HostAssoc_Human Feces | Human feces | Malawi | -13.754313 33.929313 | reads ENA | Illumina MiSeq | Newbler | 0 | |
| Reyes_Malnutrition | Malnut5082 | HostAssoc_Human Feces | Human feces | Malawi | -13.754313 33.929313 | reads ENA | Illumina MiSeq | Newbler | 0 | |
| Reyes_Malnutrition | Malnut5084 | HostAssoc_Human Feces | Human feces | Malawi | -13.754313 33.929313 | reads ENA | Illumina MiSeq | Newbler | 0 | |
| Reyes_Malnutrition | Malnut5086 | HostAssoc_Human Feces | Human feces | Malawi | -13.754313 33.929313 | reads ENA | Illumina MiSeq | Newbler | 0 | |
| Reyes_Malnutrition | Malnut5087 | HostAssoc_Human Feces | Human feces | Malawi | -13.754313 33.929313 | reads ENA | Illumina MiSeq | Newbler | 0 | |
| Reyes_Malnutrition | Malnut5088 | HostAssoc_Human Feces | Human feces | Malawi | -13.754313 33.929313 | reads ENA | Illumina MiSeq | Newbler | 0 | |
| Reyes_Malnutrition | Malnut5089 | HostAssoc_Human Feces | Human feces | Malawi | -13.754313 33.929313 | reads ENA | Illumina MiSeq | Newbler | 0 | |
| Reyes_Malnutrition | Malnut5090 | HostAssoc_Human Feces | Human feces | Malawi | -13.754313 33.929313 | reads ENA | Illumina MiSeq | Newbler | 0 | |
| Reyes_Malnutrition | Malnut5091 | HostAssoc_Human Feces | Human feces | Malawi | -13.754313 33.929313 | reads ENA | Illumina MiSeq | Newbler | 0 | |
| Reyes_Malnutrition | Malnut5092 | HostAssoc_Human Feces | Human feces | Malawi | -13.754313 33.929313 | reads ENA | Illumina MiSeq | Newbler | 0 | |
| Reyes_Malnutrition | Malnut5093 | HostAssoc_Human Feces | Human feces | Malawi | -13.754313 33.929313 | reads ENA | Illumina MiSeq | Newbler | 0 | |
| Reyes_Malnutrition | Malnut5094 | HostAssoc_Human Feces | Human feces | Malawi | -13.754313 33.929313 | reads ENA | Illumina MiSeq | Newbler | 0 | |
| Reyes_Malnutrition | Malnut5095 | HostAssoc_Human Feces | Human feces | Malawi | -13.754313 33.929313 | reads ENA | Illumina MiSeq | Newbler | 0 | |
| Reyes_Malnutrition | Malnut5096 | HostAssoc_Human Feces | Human feces | Malawi | -13.754313 33.929313 | reads ENA | Illumina MiSeq | Newbler | 0 | |
| Reyes_Malnutrition | Malnut5097 | HostAssoc_Human Feces | Human feces | Malawi | -13.754313 33.929313 | reads ENA | Illumina MiSeq | Newbler | 0 | |

| Project Name | Virome Code | Habitat Type (code) | Ecosystem Type (text) | Location | GPS | Source | Sequencing Technology | Assembly Software | # crucivirus | Reference |
|---|---|---|---|---|---|---|---|---|---|---|
| Reyes_Malnutrition | Malnut5098 | HostAssoc_Human Feces | Human feces | Malawi | -13.754313 33.929313 | reads ENA | Illumina MiSeq | Newbler | 0 | |
| Reyes_Malnutrition | Malnut5099 | HostAssoc_Human Feces | Human feces | Malawi | -13.754313 33.929313 | reads ENA | Illumina MiSeq | Newbler | 0 | |
| Reyes_Malnutrition | Malnut5100 | HostAssoc_Human Feces | Human feces | Malawi | -13.754313 33.929313 | reads ENA | Illumina MiSeq | Newbler | 0 | |
| Reyes_Malnutrition | Malnut5101 | HostAssoc_Human Feces | Human feces | Malawi | -13.754313 33.929313 | reads ENA | Illumina MiSeq | Newbler | 0 | |
| Reyes_Malnutrition | Malnut5102 | HostAssoc_Human Feces | Human feces | Malawi | -13.754313 33.929313 | reads ENA | Illumina MiSeq | Newbler | 0 | |
| Reyes_Malnutrition | Malnut5103 | HostAssoc_Human Feces | Human feces | Malawi | -13.754313 33.929313 | reads ENA | Illumina MiSeq | Newbler | 0 | |
| Reyes_Malnutrition | Malnut5104 | HostAssoc_Human Feces | Human feces | Malawi | -13.754313 33.929313 | reads ENA | Illumina MiSeq | Newbler | 0 | |
| Reyes_Malnutrition | Malnut5105 | HostAssoc_Human Feces | Human feces | Malawi | -13.754313 33.929313 | reads ENA | Illumina MiSeq | Newbler | 0 | |
| Reyes_Malnutrition | Malnut5106 | HostAssoc_Human Feces | Human feces | Malawi | -13.754313 33.929313 | reads ENA | Illumina MiSeq | Newbler | 0 | |
| Reyes_Malnutrition | Malnut5107 | HostAssoc_Human Feces | Human feces | Malawi | -13.754313 33.929313 | reads ENA | Illumina MiSeq | Newbler | 0 | |
| Reyes_Malnutrition | Malnut5108 | HostAssoc_Human Feces | Human feces | Malawi | -13.754313 33.929313 | reads ENA | Illumina MiSeq | Newbler | 0 | |
| Reyes_Malnutrition | Malnut5109 | HostAssoc_Human Feces | Human feces | Malawi | -13.754313 33.929313 | reads ENA | Illumina MiSeq | Newbler | 0 | |
| Reyes_Malnutrition | Malnut5110 | HostAssoc_Human Feces | Human feces | Malawi | -13.754313 33.929313 | reads ENA | Illumina MiSeq | Newbler | 0 | |
| Reyes_Malnutrition | Malnut5111 | HostAssoc_Human Feces | Human feces | Malawi | -13.754313 33.929313 | reads ENA | Illumina MiSeq | Newbler | 0 | |
| Reyes_Malnutrition | Malnut5112 | HostAssoc_Human Feces | Human feces | Malawi | -13.754313 33.929313 | reads ENA | Illumina MiSeq | Newbler | 0 | |
| Reyes_Malnutrition | Malnut5113 | HostAssoc_Human Feces | Human feces | Malawi | -13.754313 33.929313 | reads ENA | Illumina MiSeq | Newbler | 0 | |
| Reyes_Malnutrition | Malnut5114 | HostAssoc_Human Feces | Human feces | Malawi | -13.754313 33.929313 | reads ENA | Illumina MiSeq | Newbler | 0 | |
| Reyes_Malnutrition | Malnut5115 | HostAssoc_Human Feces | Human feces | Malawi | -13.754313 33.929313 | reads ENA | Illumina MiSeq | Newbler | 0 | |
| Reyes_Malnutrition | Malnut5116 | HostAssoc_Human Feces | Human feces | Malawi | -13.754313 33.929313 | reads ENA | Illumina MiSeq | Newbler | 0 | |
| Reyes_Malnutrition | Malnut5117 | HostAssoc_Human Feces | Human feces | Malawi | -13.754313 33.929313 | reads ENA | Illumina MiSeq | Newbler | 0 | |
| Reyes_Malnutrition | Malnut5118 | HostAssoc_Human Feces | Human feces | Malawi | -13.754313 33.929313 | reads ENA | Illumina MiSeq | Newbler | 0 | |
| Reyes_Malnutrition | Malnut5119 | HostAssoc_Human Feces | Human feces | Malawi | -13.754313 33.929313 | reads ENA | Illumina MiSeq | Newbler | 0 | |
| Reyes_Malnutrition | Malnut5120 | HostAssoc_Human Feces | Human feces | Malawi | -13.754313 33.929313 | reads ENA | Illumina MiSeq | Newbler | 0 | |
| Reyes_Malnutrition | Malnut5121 | HostAssoc_Human Feces | Human feces | Malawi | -13.754313 33.929313 | reads ENA | Illumina MiSeq | Newbler | 0 | |
| Reyes_Malnutrition | Malnut5122 | HostAssoc_Human Feces | Human feces | Malawi | -13.754313 33.929313 | reads ENA | Illumina MiSeq | Newbler | 0 | |
| Reyes_Malnutrition | Malnut5123 | HostAssoc_Human Feces | Human feces | Malawi | -13.754313 33.929313 | reads ENA | Illumina MiSeq | Newbler | 0 | |
| Reyes_Malnutrition | Malnut5124 | HostAssoc_Human Feces | Human feces | Malawi | -13.754313 33.929313 | reads ENA | Illumina MiSeq | Newbler | 0 | |
| Reyes_Malnutrition | Malnut5125 | HostAssoc_Human Feces | Human feces | Malawi | -13.754313 33.929313 | reads ENA | Illumina MiSeq | Newbler | 0 | |
| Reyes_Malnutrition | Malnut5126 | HostAssoc_Human Feces | Human feces | Malawi | -13.754313 33.929313 | reads ENA | Illumina MiSeq | Newbler | 0 | |
| Reyes_Malnutrition | Malnut5127 | HostAssoc_Human Feces | Human feces | Malawi | -13.754313 33.929313 | reads ENA | Illumina MiSeq | Newbler | 0 | |
| Reyes_Malnutrition | Malnut5128 | HostAssoc_Human Feces | Human feces | Malawi | -13.754313 33.929313 | reads ENA | Illumina MiSeq | Newbler | 0 | |
| Reyes_Malnutrition | Malnut5129 | HostAssoc_Human Feces | Human feces | Malawi | -13.754313 33.929313 | reads ENA | Illumina MiSeq | Newbler | 0 | |
| Reyes_Malnutrition | Malnut5130 | HostAssoc_Human Feces | Human feces | Malawi | -13.754313 33.929313 | reads ENA | Illumina MiSeq | Newbler | 0 | |
| Reyes_Malnutrition | Malnut5131 | HostAssoc_Human Feces | Human feces | Malawi | -13.754313 33.929313 | reads ENA | Illumina MiSeq | Newbler | 0 | |
| Reyes_Malnutrition | Malnut5132 | HostAssoc_Human Feces | Human feces | Malawi | -13.754313 33.929313 | reads ENA | Illumina MiSeq | Newbler | 0 | |
| Reyes_Malnutrition | Malnut5133 | HostAssoc_Human Feces | Human feces | Malawi | -13.754313 33.929313 | reads ENA | Illumina MiSeq | Newbler | 0 | |
| Reyes_Malnutrition | Malnut5134 | HostAssoc_Human Feces | Human feces | Malawi | -13.754313 33.929313 | reads ENA | Illumina MiSeq | Newbler | 0 | |
| Reyes_Malnutrition | Malnut5135 | HostAssoc_Human Feces | Human feces | Malawi | -13.754313 33.929313 | reads ENA | Illumina MiSeq | Newbler | 0 | |
| Reyes_Malnutrition | Malnut5136 | HostAssoc_Human Feces | Human feces | Malawi | -13.754313 33.929313 | reads ENA | Illumina MiSeq | Newbler | 0 | |
| Reyes_Malnutrition | Malnut5137 | HostAssoc_Human Feces | Human feces | Malawi | -13.754313 33.929313 | reads ENA | Illumina MiSeq | Newbler | 0 | |
| Reyes_Malnutrition | Malnut5138 | HostAssoc_Human Feces | Human feces | Malawi | -13.754313 33.929313 | reads ENA | Illumina MiSeq | Newbler | 0 | |
| Reyes_Malnutrition | Malnut5139 | HostAssoc_Human Feces | Human feces | Malawi | -13.754313 33.929313 | reads ENA | Illumina MiSeq | Newbler | 0 | |

| Project Name | Virome Code | Habitat Type (code) | Ecosystem Type (text) | Location | GPS | Source | Sequencing Technology | Assembly Software | # crucivirus | Reference |
|---|---|---|---|---|---|---|---|---|---|---|
| Reyes_Malnutrition | Malnut5140 | HostAssoc_Human Feces | Human feces | Malawi | -13.754313 33.929313 | reads ENA | Illumina MiSeq | Newbler | 0 | Reyes A. Blanton LV. Cao S. Zhao G. Manary M. Trehan I. Smith MI. Wang D. Virgin HW. Rohwer F. Gordon JI. Gut DNA viromes of Malawian twins discordant for severe acute malnutrition. Proc Natl Acad Sci U S A. 2015 Sep 22;112(38):11941-6. doi: 10.1073/pnas.1514285112. Epub 2015 Sep 8. |
| Reyes_Malnutrition | Malnut5141 | HostAssoc_Human Feces | Human feces | Malawi | -13.754313 33.929313 | reads ENA | Illumina MiSeq | Newbler | 0 | |
| Reyes_Malnutrition | Malnut5142 | HostAssoc_Human Feces | Human feces | Malawi | -13.754313 33.929313 | reads ENA | Illumina MiSeq | Newbler | 0 | |
| Reyes_Malnutrition | Malnut5143 | HostAssoc_Human Feces | Human feces | Malawi | -13.754313 33.929313 | reads ENA | Illumina MiSeq | Newbler | 0 | |
| Reyes_Malnutrition | Malnut5144 | HostAssoc_Human Feces | Human feces | Malawi | -13.754313 33.929313 | reads ENA | Illumina MiSeq | Newbler | 0 | |
| Reyes_Malnutrition | Malnut5145 | HostAssoc_Human Feces | Human feces | Malawi | -13.754313 33.929313 | reads ENA | Illumina MiSeq | Newbler | 0 | |
| Reyes_Malnutrition | Malnut5146 | HostAssoc_Human Feces | Human feces | Malawi | -13.754313 33.929313 | reads ENA | Illumina MiSeq | Newbler | 0 | |
| Reyes_Malnutrition | Malnut5147 | HostAssoc_Human Feces | Human feces | Malawi | -13.754313 33.929313 | reads ENA | Illumina MiSeq | Newbler | 0 | |
| Reyes_Malnutrition | Malnut5148 | HostAssoc_Human Feces | Human feces | Malawi | -13.754313 33.929313 | reads ENA | Illumina MiSeq | Newbler | 0 | |
| Reyes_Malnutrition | Malnut5149 | HostAssoc_Human Feces | Human feces | Malawi | -13.754313 33.929313 | reads ENA | Illumina MiSeq | Newbler | 0 | |
| Reyes_Malnutrition | Malnut5150 | HostAssoc_Human Feces | Human feces | Malawi | -13.754313 33.929313 | reads ENA | Illumina MiSeq | Newbler | 0 | |
| Reyes_Malnutrition | Malnut5151 | HostAssoc_Human Feces | Human feces | Malawi | -13.754313 33.929313 | reads ENA | Illumina MiSeq | Newbler | 0 | |
| Reyes_Malnutrition | Malnut5152 | HostAssoc_Human Feces | Human feces | Malawi | -13.754313 33.929313 | reads ENA | Illumina MiSeq | Newbler | 0 | |
| Reyes_Malnutrition | Malnut5153 | HostAssoc_Human Feces | Human feces | Malawi | -13.754313 33.929313 | reads ENA | Illumina MiSeq | Newbler | 0 | |
| Reyes_Malnutrition | Malnut5154 | HostAssoc_Human Feces | Human feces | Malawi | -13.754313 33.929313 | reads ENA | Illumina MiSeq | Newbler | 0 | |
| Reyes_Malnutrition | Malnut5155 | HostAssoc_Human Feces | Human feces | Malawi | -13.754313 33.929313 | reads ENA | Illumina MiSeq | Newbler | 0 | |
| Reyes_Malnutrition | Malnut5156 | HostAssoc_Human Feces | Human feces | Malawi | -13.754313 33.929313 | reads ENA | Illumina MiSeq | Newbler | 0 | |
| Reyes_Malnutrition | Malnut5157 | HostAssoc_Human Feces | Human feces | Malawi | -13.754313 33.929313 | reads ENA | Illumina MiSeq | Newbler | 0 | |
| Reyes_Malnutrition | Malnut5158 | HostAssoc_Human Feces | Human feces | Malawi | -13.754313 33.929313 | reads ENA | Illumina MiSeq | Newbler | 0 | |
| Reyes_Malnutrition | Malnut5159 | HostAssoc_Human Feces | Human feces | Malawi | -13.754313 33.929313 | reads ENA | Illumina MiSeq | Newbler | 0 | |
| Reyes_Malnutrition | Malnut5160 | HostAssoc_Human Feces | Human feces | Malawi | -13.754313 33.929313 | reads ENA | Illumina MiSeq | Newbler | 0 | |
| Reyes_Malnutrition | Malnut5161 | HostAssoc_Human Feces | Human feces | Malawi | -13.754313 33.929313 | reads ENA | Illumina MiSeq | Newbler | 0 | |
| Reyes_Malnutrition | Malnut5162 | HostAssoc_Human Feces | Human feces | Malawi | -13.754313 33.929313 | reads ENA | Illumina MiSeq | Newbler | 0 | |
| Reyes_Malnutrition | Malnut5163 | HostAssoc_Human Feces | Human feces | Malawi | -13.754313 33.929313 | reads ENA | Illumina MiSeq | Newbler | 0 | |
| Reyes_Malnutrition | Malnut5164 | HostAssoc_Human Feces | Human feces | Malawi | -13.754313 33.929313 | reads ENA | Illumina MiSeq | Newbler | 0 | |
| Reyes_Malnutrition | Malnut5165 | HostAssoc_Human Feces | Human feces | Malawi | -13.754313 33.929313 | reads ENA | Illumina MiSeq | Newbler | 0 | |
| Reyes_Malnutrition | Malnut5166 | HostAssoc_Human Feces | Human feces | Malawi | -13.754313 33.929313 | reads ENA | Illumina MiSeq | Newbler | 0 | |
| Reyes_Malnutrition | Malnut5167 | HostAssoc_Human Feces | Human feces | Malawi | -13.754313 33.929313 | reads ENA | Illumina MiSeq | Newbler | 0 | |
| Reyes_Malnutrition | Malnut5168 | HostAssoc_Human Feces | Human feces | Malawi | -13.754313 33.929313 | reads ENA | Illumina MiSeq | Newbler | 0 | |
| Reyes_Malnutrition | Malnut5169 | HostAssoc_Human Feces | Human feces | Malawi | -13.754313 33.929313 | reads ENA | Illumina MiSeq | Newbler | 0 | |
| Reyes_Malnutrition | Malnut5170 | HostAssoc_Human Feces | Human feces | Malawi | -13.754313 33.929313 | reads ENA | Illumina MiSeq | Newbler | 0 | |
| Reyes_Malnutrition | Malnut5171 | HostAssoc_Human Feces | Human feces | Malawi | -13.754313 33.929313 | reads ENA | Illumina MiSeq | Newbler | 0 | |
| Reyes_Malnutrition | Malnut5172 | HostAssoc_Human Feces | Human feces | Malawi | -13.754313 33.929313 | reads ENA | Illumina MiSeq | Newbler | 0 | |
| Reyes_Malnutrition | Malnut5173 | HostAssoc_Human Feces | Human feces | Malawi | -13.754313 33.929313 | reads ENA | Illumina MiSeq | Newbler | 0 | |
| Reyes_Malnutrition | Malnut5174 | HostAssoc_Human Feces | Human feces | Malawi | -13.754313 33.929313 | reads ENA | Illumina MiSeq | Newbler | 0 | |
| Reyes_Malnutrition | Malnut5175 | HostAssoc_Human Feces | Human feces | Malawi | -13.754313 33.929313 | reads ENA | Illumina MiSeq | Newbler | 0 | |
| Reyes_Malnutrition | Malnut5176 | HostAssoc_Human Feces | Human feces | Malawi | -13.754313 33.929313 | reads ENA | Illumina MiSeq | Newbler | 0 | |
| Reyes_Malnutrition | Malnut5177 | HostAssoc_Human Feces | Human feces | Malawi | -13.754313 33.929313 | reads ENA | Illumina MiSeq | Newbler | 0 | |
| Reyes_Malnutrition | Malnut5178 | HostAssoc_Human Feces | Human feces | Malawi | -13.754313 33.929313 | reads ENA | Illumina MiSeq | Newbler | 0 | |
| Reyes_Malnutrition | Malnut5179 | HostAssoc_Human Feces | Human feces | Malawi | -13.754313 33.929313 | reads ENA | Illumina MiSeq | Newbler | 0 | |
| Reyes_Malnutrition | Malnut5180 | HostAssoc_Human Feces | Human feces | Malawi | -13.754313 33.929313 | reads ENA | Illumina MiSeq | Newbler | 0 | |
| Reyes_Malnutrition | Malnut5181 | HostAssoc_Human Feces | Human feces | Malawi | -13.754313 33.929313 | reads ENA | Illumina MiSeq | Newbler | 0 | |

| Project Name | Virome Code | Habitat Type (code) | Ecosystem Type (text) | Location | GPS | Source | Sequencing Technology | Assembly Software | # crucivirus | Reference |
|---|---|---|---|---|---|---|---|---|---|---|
| Reyes_Malnutrition | Malnut5182 | HostAssoc_Human Feces | Human feces | Malawi | -13.754313 33.929313 | reads ENA | Illumina MiSeq | Newbler | 0 | |
| Reyes_Malnutrition | Malnut5183 | HostAssoc_Human Feces | Human feces | Malawi | -13.754313 33.929313 | reads ENA | Illumina MiSeq | Newbler | 0 | |
| Reyes_Malnutrition | Malnut5184 | HostAssoc_Human Feces | Human feces | Malawi | -13.754313 33.929313 | reads ENA | Illumina MiSeq | Newbler | 0 | |
| Reyes_Malnutrition | Malnut5185 | HostAssoc_Human Feces | Human feces | Malawi | -13.754313 33.929313 | reads ENA | Illumina MiSeq | Newbler | 0 | |
| Reyes_Malnutrition | Malnut5186 | HostAssoc_Human Feces | Human feces | Malawi | -13.754313 33.929313 | reads ENA | Illumina MiSeq | Newbler | 0 | |
| Reyes_Malnutrition | Malnut5187 | HostAssoc_Human Feces | Human feces | Malawi | -13.754313 33.929313 | reads ENA | Illumina MiSeq | Newbler | 0 | |
| Reyes_Malnutrition | Malnut5188 | HostAssoc_Human Feces | Human feces | Malawi | -13.754313 33.929313 | reads ENA | Illumina MiSeq | Newbler | 0 | |
| Reyes_Malnutrition | Malnut5189 | HostAssoc_Human Feces | Human feces | Malawi | -13.754313 33.929313 | reads ENA | Illumina MiSeq | Newbler | 0 | |
| Reyes_Malnutrition | Malnut5190 | HostAssoc_Human Feces | Human feces | Malawi | -13.754313 33.929313 | reads ENA | Illumina MiSeq | Newbler | 0 | |
| Reyes_Malnutrition | Malnut5191 | HostAssoc_Human Feces | Human feces | Malawi | -13.754313 33.929313 | reads ENA | Illumina MiSeq | Newbler | 0 | |
| Reyes_Malnutrition | Malnut5192 | HostAssoc_Human Feces | Human feces | Malawi | -13.754313 33.929313 | reads ENA | Illumina MiSeq | Newbler | 0 | |
| Reyes_Malnutrition | Malnut5193 | HostAssoc_Human Feces | Human feces | Malawi | -13.754313 33.929313 | reads ENA | Illumina MiSeq | Newbler | 0 | |
| Reyes_Malnutrition | Malnut5194 | HostAssoc_Human Feces | Human feces | Malawi | -13.754313 33.929313 | reads ENA | Illumina MiSeq | Newbler | 0 | |
| Reyes_Malnutrition | Malnut5195 | HostAssoc_Human Feces | Human feces | Malawi | -13.754313 33.929313 | reads ENA | Illumina MiSeq | Newbler | 0 | |
| Reyes_Malnutrition | Malnut5196 | HostAssoc_Human Feces | Human feces | Malawi | -13.754313 33.929313 | reads ENA | Illumina MiSeq | Newbler | 0 | |
| Reyes_Malnutrition | Malnut5197 | HostAssoc_Human Feces | Human feces | Malawi | -13.754313 33.929313 | reads ENA | Illumina MiSeq | Newbler | 0 | |
| Reyes_Malnutrition | Malnut5198 | HostAssoc_Human Feces | Human feces | Malawi | -13.754313 33.929313 | reads ENA | Illumina MiSeq | Newbler | 0 | |
| Reyes_Malnutrition | Malnut5199 | HostAssoc_Human Feces | Human feces | Malawi | -13.754313 33.929313 | reads ENA | Illumina MiSeq | Newbler | 0 | |
| Reyes_Malnutrition | Malnut5200 | HostAssoc_Human Feces | Human feces | Malawi | -13.754313 33.929313 | reads ENA | Illumina MiSeq | Newbler | 0 | |
| Reyes_Malnutrition | Malnut5201 | HostAssoc_Human Feces | Human feces | Malawi | -13.754313 33.929313 | reads ENA | Illumina MiSeq | Newbler | 0 | |
| Reyes_Malnutrition | Malnut5202 | HostAssoc_Human Feces | Human feces | Malawi | -13.754313 33.929313 | reads ENA | Illumina MiSeq | Newbler | 0 | |
| Reyes_Malnutrition | Malnut5203 | HostAssoc_Human Feces | Human feces | Malawi | -13.754313 33.929313 | reads ENA | Illumina MiSeq | Newbler | 0 | |
| Reyes_Malnutrition | Malnut5204 | HostAssoc_Human Feces | Human feces | Malawi | -13.754313 33.929313 | reads ENA | Illumina MiSeq | Newbler | 0 | |
| Reyes_Malnutrition | Malnut5205 | HostAssoc_Human Feces | Human feces | Malawi | -13.754313 33.929313 | reads ENA | Illumina MiSeq | Newbler | 0 | |
| Reyes_Malnutrition | Malnut5206 | HostAssoc_Human Feces | Human feces | Malawi | -13.754313 33.929313 | reads ENA | Illumina MiSeq | Newbler | 0 | |
| Reyes_Malnutrition | Malnut5207 | HostAssoc_Human Feces | Human feces | Malawi | -13.754313 33.929313 | reads ENA | Illumina MiSeq | Newbler | 0 | |
| Reyes_Malnutrition | Malnut5208 | HostAssoc_Human Feces | Human feces | Malawi | -13.754313 33.929313 | reads ENA | Illumina MiSeq | Newbler | 0 | |
| Reyes_Malnutrition | Malnut5209 | HostAssoc_Human Feces | Human feces | Malawi | -13.754313 33.929313 | reads ENA | Illumina MiSeq | Newbler | 0 | |
| Reyes_Malnutrition | Malnut5210 | HostAssoc_Human Feces | Human feces | Malawi | -13.754313 33.929313 | reads ENA | Illumina MiSeq | Newbler | 0 | |
| Reyes_Malnutrition | Malnut5211 | HostAssoc_Human Feces | Human feces | Malawi | -13.754313 33.929313 | reads ENA | Illumina MiSeq | Newbler | 0 | |
| Reyes_Malnutrition | Malnut5212 | HostAssoc_Human Feces | Human feces | Malawi | -13.754313 33.929313 | reads ENA | Illumina MiSeq | Newbler | 0 | |
| Reyes_Malnutrition | Malnut5213 | HostAssoc_Human Feces | Human feces | Malawi | -13.754313 33.929313 | reads ENA | Illumina MiSeq | Newbler | 0 | |
| Reyes_Malnutrition | Malnut5214 | HostAssoc_Human Feces | Human feces | Malawi | -13.754313 33.929313 | reads ENA | Illumina MiSeq | Newbler | 0 | |
| Reyes_Malnutrition | Malnut5215 | HostAssoc_Human Feces | Human feces | Malawi | -13.754313 33.929313 | reads ENA | Illumina MiSeq | Newbler | 0 | |
| Reyes_Malnutrition | Malnut5216 | HostAssoc_Human Feces | Human feces | Malawi | -13.754313 33.929313 | reads ENA | Illumina MiSeq | Newbler | 0 | |
| Reyes_Malnutrition | Malnut5217 | HostAssoc_Human Feces | Human feces | Malawi | -13.754313 33.929313 | reads ENA | Illumina MiSeq | Newbler | 0 | |
| Reyes_Malnutrition | Malnut5218 | HostAssoc_Human Feces | Human feces | Malawi | -13.754313 33.929313 | reads ENA | Illumina MiSeq | Newbler | 0 | |
| Reyes_Malnutrition | Malnut5219 | HostAssoc_Human Feces | Human feces | Malawi | -13.754313 33.929313 | reads ENA | Illumina MiSeq | Newbler | 0 | |
| Reyes_Malnutrition | Malnut5220 | HostAssoc_Human Feces | Human feces | Malawi | -13.754313 33.929313 | reads ENA | Illumina MiSeq | Newbler | 0 | |
| Reyes_Malnutrition | Malnut5221 | HostAssoc_Human Feces | Human feces | Malawi | -13.754313 33.929313 | reads ENA | Illumina MiSeq | Newbler | 0 | |
| Reyes_Malnutrition | Malnut5222 | HostAssoc_Human Feces | Human feces | Malawi | -13.754313 33.929313 | reads ENA | Illumina MiSeq | Newbler | 0 | |
| Reyes_Malnutrition | Malnut5223 | HostAssoc_Human Feces | Human feces | Malawi | -13.754313 33.929313 | reads ENA | Illumina MiSeq | Newbler | 0 | |

| Project Name | Virome Code | Habitat Type (code) | Ecosystem Type (text) | Location | GPS | Source | Sequencing Technology | Assembly Software | # crucivirus | Reference |
|---|---|---|---|---|---|---|---|---|---|---|
| Reyes_Malnutrition | Malnut5224 | HostAssoc_Human Feces | Human feces | Malawi | -13.754313 33.929313 | reads ENA | Illumina MiSeq | Newbler | 0 | |
| Reyes_Malnutrition | Malnut5225 | HostAssoc_Human Feces | Human feces | Malawi | -13.754313 33.929313 | reads ENA | Illumina MiSeq | Newbler | 0 | |
| Reyes_Malnutrition | Malnut5226 | HostAssoc_Human Feces | Human feces | Malawi | -13.754313 33.929313 | reads ENA | Illumina MiSeq | Newbler | 0 | |
| Reyes_Malnutrition | Malnut5227 | HostAssoc_Human Feces | Human feces | Malawi | -13.754313 33.929313 | reads ENA | Illumina MiSeq | Newbler | 0 | |
| Reyes_Malnutrition | Malnut5228 | HostAssoc_Human Feces | Human feces | Malawi | -13.754313 33.929313 | reads ENA | Illumina MiSeq | Newbler | 0 | |
| Reyes_Malnutrition | Malnut5229 | HostAssoc_Human Feces | Human feces | Malawi | -13.754313 33.929313 | reads ENA | Illumina MiSeq | Newbler | 0 | |
| Reyes_Malnutrition | Malnut5230 | HostAssoc_Human Feces | Human feces | Malawi | -13.754313 33.929313 | reads ENA | Illumina MiSeq | Newbler | 0 | |
| Reyes_Malnutrition | Malnut5231 | HostAssoc_Human Feces | Human feces | Malawi | -13.754313 33.929313 | reads ENA | Illumina MiSeq | Newbler | 0 | |
| Reyes_Malnutrition | Malnut5232 | HostAssoc_Human Feces | Human feces | Malawi | -13.754313 33.929313 | reads ENA | Illumina MiSeq | Newbler | 0 | |
| Reyes_Malnutrition | Malnut5233 | HostAssoc_Human Feces | Human feces | Malawi | -13.754313 33.929313 | reads ENA | Illumina MiSeq | Newbler | 0 | |
| Reyes_Malnutrition | Malnut5234 | HostAssoc_Human Feces | Human feces | Malawi | -13.754313 33.929313 | reads ENA | Illumina MiSeq | Newbler | 0 | |
| Reyes_Malnutrition | Malnut5235 | HostAssoc_Human Feces | Human feces | Malawi | -13.754313 33.929313 | reads ENA | Illumina MiSeq | Newbler | 0 | |
| Reyes_Malnutrition | Malnut5236 | HostAssoc_Human Feces | Human feces | Malawi | -13.754313 33.929313 | reads ENA | Illumina MiSeq | Newbler | 0 | |
| Reyes_Malnutrition | Malnut5237 | HostAssoc_Human Feces | Human feces | Malawi | -13.754313 33.929313 | reads ENA | Illumina MiSeq | Newbler | 0 | |
| Reyes_Malnutrition | Malnut5238 | HostAssoc_Human Feces | Human feces | Malawi | -13.754313 33.929313 | reads ENA | Illumina MiSeq | Newbler | 0 | |
| Reyes_Malnutrition | Malnut5239 | HostAssoc_Human Feces | Human feces | Malawi | -13.754313 33.929313 | reads ENA | Illumina MiSeq | Newbler | 0 | |
| Reyes_Malnutrition | Malnut5240 | HostAssoc_Human Feces | Human feces | Malawi | -13.754313 33.929313 | reads ENA | Illumina MiSeq | Newbler | 0 | |
| Reyes_Malnutrition | Malnut5241 | HostAssoc_Human Feces | Human feces | Malawi | -13.754313 33.929313 | reads ENA | Illumina MiSeq | Newbler | 0 | |
| Reyes_Malnutrition | Malnut5242 | HostAssoc_Human Feces | Human feces | Malawi | -13.754313 33.929313 | reads ENA | Illumina MiSeq | Newbler | 0 | |
| Reyes_Malnutrition | Malnut5243 | HostAssoc_Human Feces | Human feces | Malawi | -13.754313 33.929313 | reads ENA | Illumina MiSeq | Newbler | 0 | |
| Reyes_Malnutrition | Malnut5244 | HostAssoc_Human Feces | Human feces | Malawi | -13.754313 33.929313 | reads ENA | Illumina MiSeq | Newbler | 0 | |
| Reyes_Malnutrition | Malnut5245 | HostAssoc_Human Feces | Human feces | Malawi | -13.754313 33.929313 | reads ENA | Illumina MiSeq | Newbler | 0 | |
| Reyes_Malnutrition | Malnut5246 | HostAssoc_Human Feces | Human feces | Malawi | -13.754313 33.929313 | reads ENA | Illumina MiSeq | Newbler | 0 | |
| Reyes_Malnutrition | Malnut5247 | HostAssoc_Human Feces | Human feces | Malawi | -13.754313 33.929313 | reads ENA | Illumina MiSeq | Newbler | 0 | |
| Reyes_Malnutrition | Malnut5248 | HostAssoc_Human Feces | Human feces | Malawi | -13.754313 33.929313 | reads ENA | Illumina MiSeq | Newbler | 0 | |
| Reyes_Malnutrition | Malnut5249 | HostAssoc_Human Feces | Human feces | Malawi | -13.754313 33.929313 | reads ENA | Illumina MiSeq | Newbler | 0 | |
| Reyes_Malnutrition | Malnut5250 | HostAssoc_Human Feces | Human feces | Malawi | -13.754313 33.929313 | reads ENA | Illumina MiSeq | Newbler | 0 | |
| Reyes_Malnutrition | Malnut5251 | HostAssoc_Human Feces | Human feces | Malawi | -13.754313 33.929313 | reads ENA | Illumina MiSeq | Newbler | 0 | |
| Reyes_Malnutrition | Malnut5252 | HostAssoc_Human Feces | Human feces | Malawi | -13.754313 33.929313 | reads ENA | Illumina MiSeq | Newbler | 0 | |
| Reyes_Malnutrition | Malnut5253 | HostAssoc_Human Feces | Human feces | Malawi | -13.754313 33.929313 | reads ENA | Illumina MiSeq | Newbler | 0 | |
| Reyes_Malnutrition | Malnut5254 | HostAssoc_Human Feces | Human feces | Malawi | -13.754313 33.929313 | reads ENA | Illumina MiSeq | Newbler | 0 | |
| Reyes_Malnutrition | Malnut5255 | HostAssoc_Human Feces | Human feces | Malawi | -13.754313 33.929313 | reads ENA | Illumina MiSeq | Newbler | 0 | |
| Reyes_Malnutrition | Malnut5256 | HostAssoc_Human Feces | Human feces | Malawi | -13.754313 33.929313 | reads ENA | Illumina MiSeq | Newbler | 0 | |
| Reyes_Malnutrition | Malnut5257 | HostAssoc_Human Feces | Human feces | Malawi | -13.754313 33.929313 | reads ENA | Illumina MiSeq | Newbler | 0 | |
| Reyes_Malnutrition | Malnut5258 | HostAssoc_Human Feces | Human feces | Malawi | -13.754313 33.929313 | reads ENA | Illumina MiSeq | Newbler | 0 | |
| Reyes_Malnutrition | Malnut5259 | HostAssoc_Human Feces | Human feces | Malawi | -13.754313 33.929313 | reads ENA | Illumina MiSeq | Newbler | 0 | |
| Reyes_Malnutrition | Malnut5260 | HostAssoc_Human Feces | Human feces | Malawi | -13.754313 33.929313 | reads ENA | Illumina MiSeq | Newbler | 0 | |
| Reyes_Malnutrition | Malnut5261 | HostAssoc_Human Feces | Human feces | Malawi | -13.754313 33.929313 | reads ENA | Illumina MiSeq | Newbler | 0 | |
| Reyes_Malnutrition | Malnut5262 | HostAssoc_Human Feces | Human feces | Malawi | -13.754313 33.929313 | reads ENA | Illumina MiSeq | Newbler | 0 | |
| Reyes_Malnutrition | Malnut5263 | HostAssoc_Human Feces | Human feces | Malawi | -13.754313 33.929313 | reads ENA | Illumina MiSeq | Newbler | 0 | |
| Reyes_Malnutrition | Malnut5264 | HostAssoc_Human Feces | Human feces | Malawi | -13.754313 33.929313 | reads ENA | Illumina MiSeq | Newbler | 0 | |
| Reyes_Malnutrition | Malnut5265 | HostAssoc_Human Feces | Human feces | Malawi | -13.754313 33.929313 | reads ENA | Illumina MiSeq | Newbler | 0 | |

| Project Name | Virome Code | Habitat Type (code) | Ecosystem Type (text) | Location | GPS | Source | Sequencing Technology | Assembly Software | # crucivirus | Reference |
|---|---|---|---|---|---|---|---|---|---|---|
| Reyes_Malnutrition | Malnut5266 | HostAssoc_Human Feces | Human feces | Malawi | -13.754313 33.929313 | reads ENA | Illumina MiSeq | Newbler | 0 | |
| Reyes_Malnutrition | Malnut5267 | HostAssoc_Human Feces | Human feces | Malawi | -13.754313 33.929313 | reads ENA | Illumina MiSeq | Newbler | 0 | |
| Reyes_Malnutrition | Malnut5268 | HostAssoc_Human Feces | Human feces | Malawi | -13.754313 33.929313 | reads ENA | Illumina MiSeq | Newbler | 0 | |
| Reyes_Malnutrition | Malnut5269 | HostAssoc_Human Feces | Human feces | Malawi | -13.754313 33.929313 | reads ENA | Illumina MiSeq | Newbler | 0 | |
| Reyes_Malnutrition | Malnut5270 | HostAssoc_Human Feces | Human feces | Malawi | -13.754313 33.929313 | reads ENA | Illumina MiSeq | Newbler | 0 | |
| Reyes_Malnutrition | Malnut5271 | HostAssoc_Human Feces | Human feces | Malawi | -13.754313 33.929313 | reads ENA | Illumina MiSeq | Newbler | 0 | |
| Reyes_Malnutrition | Malnut5272 | HostAssoc_Human Feces | Human feces | Malawi | -13.754313 33.929313 | reads ENA | Illumina MiSeq | Newbler | 0 | |
| Reyes_Malnutrition | Malnut5273 | HostAssoc_Human Feces | Human feces | Malawi | -13.754313 33.929313 | reads ENA | Illumina MiSeq | Newbler | 0 | |
| Reyes_Malnutrition | Malnut5274 | HostAssoc_Human Feces | Human feces | Malawi | -13.754313 33.929313 | reads ENA | Illumina MiSeq | Newbler | 0 | |
| Reyes_Malnutrition | Malnut5275 | HostAssoc_Human Feces | Human feces | Malawi | -13.754313 33.929313 | reads ENA | Illumina MiSeq | Newbler | 0 | |
| Reyes_Malnutrition | Malnut5276 | HostAssoc_Human Feces | Human feces | Malawi | -13.754313 33.929313 | reads ENA | Illumina MiSeq | Newbler | 0 | |
| Reyes_Malnutrition | Malnut5277 | HostAssoc_Human Feces | Human feces | Malawi | -13.754313 33.929313 | reads ENA | Illumina MiSeq | Newbler | 0 | |
| Reyes_Malnutrition | Malnut5278 | HostAssoc_Human Feces | Human feces | Malawi | -13.754313 33.929313 | reads ENA | Illumina MiSeq | Newbler | 0 | |
| Reyes_Malnutrition | Malnut5279 | HostAssoc_Human Feces | Human feces | Malawi | -13.754313 33.929313 | reads ENA | Illumina MiSeq | Newbler | 0 | |
| Reyes_Malnutrition | Malnut5280 | HostAssoc_Human Feces | Human feces | Malawi | -13.754313 33.929313 | reads ENA | Illumina MiSeq | Newbler | 0 | |
| Reyes_Malnutrition | Malnut5281 | HostAssoc_Human Feces | Human feces | Malawi | -13.754313 33.929313 | reads ENA | Illumina MiSeq | Newbler | 0 | |
| Reyes_Malnutrition | Malnut5282 | HostAssoc_Human Feces | Human feces | Malawi | -13.754313 33.929313 | reads ENA | Illumina MiSeq | Newbler | 0 | |
| Reyes_Malnutrition | Malnut5283 | HostAssoc_Human Feces | Human feces | Malawi | -13.754313 33.929313 | reads ENA | Illumina MiSeq | Newbler | 0 | |
| Reyes_Malnutrition | Malnut5284 | HostAssoc_Human Feces | Human feces | Malawi | -13.754313 33.929313 | reads ENA | Illumina MiSeq | Newbler | 0 | |
| Reyes_Malnutrition | Malnut5285 | HostAssoc_Human Feces | Human feces | Malawi | -13.754313 33.929313 | reads ENA | Illumina MiSeq | Newbler | 0 | |
| Reyes_Malnutrition | Malnut5286 | HostAssoc_Human Feces | Human feces | Malawi | -13.754313 33.929313 | reads ENA | Illumina MiSeq | Newbler | 0 | |
| Reyes_Malnutrition | Malnut5287 | HostAssoc_Human Feces | Human feces | Malawi | -13.754313 33.929313 | reads ENA | Illumina MiSeq | Newbler | 0 | |
| Reyes_Malnutrition | Malnut5288 | HostAssoc_Human Feces | Human feces | Malawi | -13.754313 33.929313 | reads ENA | Illumina MiSeq | Newbler | 0 | |
| Reyes_Malnutrition | Malnut5289 | HostAssoc_Human Feces | Human feces | Malawi | -13.754313 33.929313 | reads ENA | Illumina MiSeq | Newbler | 0 | |
| Reyes_Malnutrition | Malnut5290 | HostAssoc_Human Feces | Human feces | Malawi | -13.754313 33.929313 | reads ENA | Illumina MiSeq | Newbler | 0 | |
| Reyes_Malnutrition | Malnut5291 | HostAssoc_Human Feces | Human feces | Malawi | -13.754313 33.929313 | reads ENA | Illumina MiSeq | Newbler | 0 | |
| Reyes_Malnutrition | Malnut5292 | HostAssoc_Human Feces | Human feces | Malawi | -13.754313 33.929313 | reads ENA | Illumina MiSeq | Newbler | 0 | |
| Reyes_Malnutrition | Malnut5293 | HostAssoc_Human Feces | Human feces | Malawi | -13.754313 33.929313 | reads ENA | Illumina MiSeq | Newbler | 0 | |
| Reyes_Malnutrition | Malnut5294 | HostAssoc_Human Feces | Human feces | Malawi | -13.754313 33.929313 | reads ENA | Illumina MiSeq | Newbler | 0 | |
| Reyes_Malnutrition | Malnut5295 | HostAssoc_Human Feces | Human feces | Malawi | -13.754313 33.929313 | reads ENA | Illumina MiSeq | Newbler | 0 | |
| Reyes_Malnutrition | Malnut5296 | HostAssoc_Human Feces | Human feces | Malawi | -13.754313 33.929313 | reads ENA | Illumina MiSeq | Newbler | 0 | |
| Reyes_Malnutrition | Malnut5297 | HostAssoc_Human Feces | Human feces | Malawi | -13.754313 33.929313 | reads ENA | Illumina MiSeq | Newbler | 0 | |
| Reyes_Malnutrition | Malnut5298 | HostAssoc_Human Feces | Human feces | Malawi | -13.754313 33.929313 | reads ENA | Illumina MiSeq | Newbler | 0 | |
| Reyes_Malnutrition | Malnut5299 | HostAssoc_Human Feces | Human feces | Malawi | -13.754313 33.929313 | reads ENA | Illumina MiSeq | Newbler | 0 | |
| Reyes_Malnutrition | Malnut5300 | HostAssoc_Human Feces | Human feces | Malawi | -13.754313 33.929313 | reads ENA | Illumina MiSeq | Newbler | 0 | |
| Reyes_Malnutrition | Malnut5301 | HostAssoc_Human Feces | Human feces | Malawi | -13.754313 33.929313 | reads ENA | Illumina MiSeq | Newbler | 0 | |
| Reyes_Malnutrition | Malnut5302 | HostAssoc_Human Feces | Human feces | Malawi | -13.754313 33.929313 | reads ENA | Illumina MiSeq | Newbler | 0 | |
| Reyes_Malnutrition | Malnut5303 | HostAssoc_Human Feces | Human feces | Malawi | -13.754313 33.929313 | reads ENA | Illumina MiSeq | Newbler | 0 | |
| Reyes_Malnutrition | Malnut5304 | HostAssoc_Human Feces | Human feces | Malawi | -13.754313 33.929313 | reads ENA | Illumina MiSeq | Newbler | 0 | |
| Reyes_Malnutrition | Malnut5305 | HostAssoc_Human Feces | Human feces | Malawi | -13.754313 33.929313 | reads ENA | Illumina MiSeq | Newbler | 0 | |
| Reyes_Malnutrition | Malnut5306 | HostAssoc_Human Feces | Human feces | Malawi | -13.754313 33.929313 | reads ENA | Illumina MiSeq | Newbler | 0 | |
| Reyes_Malnutrition | Malnut5307 | HostAssoc_Human Feces | Human feces | Malawi | -13.754313 33.929313 | reads ENA | Illumina MiSeq | Newbler | 0 | |

| Project Name | Virome Code | Habitat Type (code) | Ecosystem Type (text) | Location | GPS | Source | Sequencing Technology | Assembly Software | # crucivirus | Reference |
|---|---|---|---|---|---|---|---|---|---|---|
| Reyes_Malnutrition | Malnut5308 | HostAssoc_Human Feces | Human feces | Malawi | -13.754313 33.929313 | reads ENA | Illumina MiSeq | Newbler | 0 | |
| Reyes_Malnutrition | Malnut5309 | HostAssoc_Human Feces | Human feces | Malawi | -13.754313 33.929313 | reads ENA | Illumina MiSeq | Newbler | 0 | |
| Reyes_Malnutrition | Malnut5310 | HostAssoc_Human Feces | Human feces | Malawi | -13.754313 33.929313 | reads ENA | Illumina MiSeq | Newbler | 0 | |
| Reyes_Malnutrition | Malnut5311 | HostAssoc_Human Feces | Human feces | Malawi | -13.754313 33.929313 | reads ENA | Illumina MiSeq | Newbler | 0 | |
| Reyes_Malnutrition | Malnut5312 | HostAssoc_Human Feces | Human feces | Malawi | -13.754313 33.929313 | reads ENA | Illumina MiSeq | Newbler | 0 | |
| Reyes_Malnutrition | Malnut5313 | HostAssoc_Human Feces | Human feces | Malawi | -13.754313 33.929313 | reads ENA | Illumina MiSeq | Newbler | 0 | |
| Reyes_Malnutrition | Malnut5314 | HostAssoc_Human Feces | Human feces | Malawi | -13.754313 33.929313 | reads ENA | Illumina MiSeq | Newbler | 0 | |
| Reyes_Malnutrition | Malnut5315 | HostAssoc_Human Feces | Human feces | Malawi | -13.754313 33.929313 | reads ENA | Illumina MiSeq | Newbler | 0 | |
| Reyes_Malnutrition | Malnut5316 | HostAssoc_Human Feces | Human feces | Malawi | -13.754313 33.929313 | reads ENA | Illumina MiSeq | Newbler | 0 | |
| Reyes_Malnutrition | Malnut5317 | HostAssoc_Human Feces | Human feces | Malawi | -13.754313 33.929313 | reads ENA | Illumina MiSeq | Newbler | 0 | |
| Reyes_Malnutrition | Malnut5318 | HostAssoc_Human Feces | Human feces | Malawi | -13.754313 33.929313 | reads ENA | Illumina MiSeq | Newbler | 0 | |
| Reyes_Malnutrition | Malnut5319 | HostAssoc_Human Feces | Human feces | Malawi | -13.754313 33.929313 | reads ENA | Illumina MiSeq | Newbler | 0 | |
| Reyes_Malnutrition | Malnut5320 | HostAssoc_Human Feces | Human feces | Malawi | -13.754313 33.929313 | reads ENA | Illumina MiSeq | Newbler | 0 | |
| Reyes_Malnutrition | Malnut5321 | HostAssoc_Human Feces | Human feces | Malawi | -13.754313 33.929313 | reads ENA | Illumina MiSeq | Newbler | 0 | |
| Reyes_TwinsFeces | TwinFeces | HostAssoc_Human Feces | Human feces | USA : Missouri | 38.622948 -92.550521 | contigs Metavir | ? | Newbler | 0 | Reyes A. Haynes M. Hanson N. Angly FE. Heath AC. Rohwer F. Gordon JI. Viruses in the faecal microbiota of monozygotic twins and their mothers. Nature. 2010 Jul 15;466(7304):334-8. doi: 10.1038/nature09199. |
| Willner_CFibrosis | CF10 | HostAssoc_Human Feces | Human feces | USA : California | 32.754288 -117.166328 | reads Metavir | 454 | Newbler | 0 | |
| Willner_CFibrosis | CF6 | HostAssoc_Human Feces | Human feces | USA : California | 32.754288 -117.166328 | reads Metavir | 454 | Newbler | 0 | |
| Willner_CFibrosis | CF7 | HostAssoc_Human Feces | Human feces | USA : California | 32.754288 -117.166328 | reads Metavir | 454 | Newbler | 0 | |
| Willner_CFibrosis | CF8 | HostAssoc_Human Feces | Human feces | USA : California | 32.754288 -117.166328 | reads Metavir | 454 | Newbler | 0 | |
| Willner_CFibrosis | CF9 | HostAssoc_Human Feces | Human feces | USA : California | 32.754288 -117.166328 | reads Metavir | 454 | Newbler | 0 | Willner D, Furlan M, Haynes M, et al. Metagenomic analysis of respiratory tract DNA viral communities in cystic fibrosis and non-cystic fibrosis individuals. PLoS One. 2009;4(10). doi:10.1371/journal.pone.0007370. |
| Willner_CFibrosis | Norm3 | HostAssoc_Human Feces | Human feces | USA : California | 32.754288 -117.166328 | reads Metavir | 454 | Newbler | 0 | |
| Willner_CFibrosis | Norm4 | HostAssoc_Human Feces | Human feces | USA : California | 32.754288 -117.166328 | reads Metavir | 454 | Newbler | 0 | |
| Willner_CFibrosis | Norm5 | HostAssoc_Human Feces | Human feces | USA : California | 32.754288 -117.166328 | reads Metavir | 454 | Newbler | 0 | |
| Willner_CFibrosis | Norm6 | HostAssoc_Human Feces | Human feces | USA : California | 32.754288 -117.166328 | reads Metavir | 454 | Newbler | 0 | |
| Willner_CFibrosis | Norm7 | HostAssoc_Human Feces | Human feces | USA : California | 32.754288 -117.166328 | reads Metavir | 454 | Newbler | 0 | |
| Dinakaran_CardioDiseaseCircDNA | CON029 | HostAssoc_Human Other | | India : Madurai, Tamil Nadu | 9.926890 78.132085 | contigs Metavir | Ion Torrent (PGM) | ? | 0 | |
| Dinakaran_CardioDiseaseCircDNA | CON030 | HostAssoc_Human Other | | India : Madurai, Tamil Nadu | 9.926890 78.132085 | contigs Metavir | Ion Torrent (PGM) | ? | 0 | |
| Dinakaran_CardioDiseaseCircDNA | CON064 | HostAssoc_Human Other | | India : Madurai, Tamil Nadu | 9.926890 78.132085 | contigs Metavir | Ion Torrent (PGM) | ? | 0 | Dinakaran V. Rathinavel A. Pushpanathan M. Sivakumar R. Gunasekaran P. Rajendhran J. Elevated levels of circulating DNA in cardiovascular disease patients: metagenomic profiling of microbiome in the circulation. PLoS One. 2014 Aug 18;9(8):e105221. doi: 10.1371/journal.pone.0105221. |
| Dinakaran_CardioDiseaseCircDNA | CVD008 | HostAssoc_Human Other | | India : Madurai, Tamil Nadu | 9.926890 78.132085 | contigs Metavir | Ion Torrent (PGM) | ? | 0 | |
| Dinakaran_CardioDiseaseCircDNA | CVD010 | HostAssoc_Human Other | | India : Madurai, Tamil Nadu | 9.926890 78.132085 | contigs Metavir | Ion Torrent (PGM) | ? | 0 | |
| Dinakaran_CardioDiseaseCircDNA | CVD014 | HostAssoc_Human Other | | India : Madurai, Tamil Nadu | 9.926890 78.132085 | contigs Metavir | Ion Torrent (PGM) | ? | 0 | |
| Fancello_PeriCardialFluids | PeriCardCtrl2 | HostAssoc_Human Other | human fluids | France | 46.445546 2.888748 | reads Metavir | 454 | Newbler | 0 | |
| Fancello_PeriCardialFluids | PeriCard1 | HostAssoc_Human Other | human fluids | France | 46.445546 2.888748 | reads Metavir | 454 | Newbler | 0 | |
| Fancello_PeriCardialFluids | PeriCard2 | HostAssoc_Human Other | human fluids | France | 46.445546 2.888748 | reads Metavir | 454 | Newbler | 0 | |
| Fancello_PeriCardialFluids | PeriCard3 | HostAssoc_Human Other | human fluids | France | 46.445546 2.888748 | reads Metavir | 454 | Newbler | 0 | |
| Fancello_PeriCardialFluids | PeriCard4 | HostAssoc_Human Other | human fluids | France | 46.445546 2.888748 | reads Metavir | 454 | Newbler | 0 | |
| Fancello_PeriCardialFluids | PeriCard5 | HostAssoc_Human Other | human fluids | France | 46.445546 2.888748 | reads Metavir | 454 | Newbler | 0 | Fancello L. Monteil S. Popgeorgiev N. Rivet R. Gouriet F. Fournier PE. Raoult D. Desnues C. Viral communities associated with human pericardial fluids in idiopathic pericarditis. PLoS One. 2014 Apr 1;9(4):e93367. doi: 10.1371/journal.pone.0093367. ECollection 2014. |
| Fancello_PeriCardialFluids | PeriCard6 | HostAssoc_Human Other | human fluids | France | 46.445546 2.888748 | reads Metavir | 454 | Newbler | 0 | |
| Fancello_PeriCardialFluids | PeriCard7 | HostAssoc_Human Other | human fluids | France | 46.445546 2.888748 | reads Metavir | 454 | Newbler | 0 | |
| Fancello_PeriCardialFluids | PeriCard8 | HostAssoc_Human Other | human fluids | France | 46.445546 2.888748 | reads Metavir | 454 | Newbler | 0 | |
| Fancello_PeriCardialFluids | PeriCardCtrl1 | HostAssoc_Human Other | human fluids | France | 46.445546 2.888748 | reads Metavir | 454 | Newbler | 0 | |
| Fancello_PeriCardialFluids | PeriCardCtrlpos | HostAssoc_Human Other | human fluids | France | 46.445546 2.888748 | reads Metavir | 454 | Newbler | 0 | |

| Project Name | Virome Code | Habitat Type (code) | Ecosystem Type (text) | Location | GPS | Source | Sequencing Technology | Assembly Software | # crucivirus | Reference |
|---|---|---|---|---|---|---|---|---|---|---|
| Gibbons_restrooms | F1FH81130 | HostAssoc_Human Other | human fluids | USA : San Diego | 32.777655 -117.071384 | reads MG-RAST | | Newbler | 0 | Gibbons SM. Schwartz T. Fouquier J. Mitchell M. Sangwan N. Gilbert JA. Kelley ST. Ecological succession and viability of human-associated microbiota on restroom surfaces. Appl Environ Microbiol. 2015 Jan;81(2):765-73. doi: 10.1128/AEM.03117-14. |
| Gibbons_restrooms | F1MH21130 | HostAssoc_Human Other | human fluids | USA : San Diego | 32.777655 -117.071384 | reads MG-RAST | | Newbler | 0 | |
| Gibbons_restrooms | F3FH41205 | HostAssoc_Human Other | human fluids | USA : San Diego | 32.777655 -117.071384 | reads MG-RAST | | Newbler | 0 | |
| Gibbons_restrooms | F3FH71130 | HostAssoc_Human Other | human fluids | USA : San Diego | 32.777655 -117.071384 | reads MG-RAST | | Newbler | 0 | |
| Gibbons_restrooms | F3MH41205 | HostAssoc_Human Other | human fluids | USA : San Diego | 32.777655 -117.071384 | reads MG-RAST | | Newbler | 0 | |
| Gibbons_restrooms | F3MH61130 | HostAssoc_Human Other | human fluids | USA : San Diego | 32.777655 -117.071384 | reads MG-RAST | | Newbler | 0 | |
| Gibbons_restrooms | F3MH81130 | HostAssoc_Human Other | human fluids | USA : San Diego | 32.777655 -117.071384 | reads MG-RAST | | Newbler | 0 | |
| Pride_LongTermAntibio | ELA100ASal | HostAssoc_Human Other | Human saliva | USA : California | 32.876609 -117.236885 | reads MG-RAST | Ion Torrent | Newbler -large | 0 | |
| Pride_LongTermAntibio | ELA100BSal | HostAssoc_Human Other | Human saliva | USA : California | 32.876609 -117.236885 | reads MG-RAST | Ion Torrent | Newbler -large | 0 | |
| Pride_LongTermAntibio | ELA100CSal | HostAssoc_Human Other | Human saliva | USA : California | 32.876609 -117.236885 | reads MG-RAST | Ion Torrent | Newbler -large | 0 | |
| Pride_LongTermAntibio | ELA1ASal | HostAssoc_Human Other | Human saliva | USA : California | 32.876609 -117.236885 | reads MG-RAST | Ion Torrent | Newbler -large | 0 | |
| Pride_LongTermAntibio | ELA1BSal | HostAssoc_Human Other | Human saliva | USA : California | 32.876609 -117.236885 | reads MG-RAST | Ion Torrent | Newbler -large | 0 | |
| Pride_LongTermAntibio | ELA1CSal | HostAssoc_Human Other | Human saliva | USA : California | 32.876609 -117.236885 | reads MG-RAST | Ion Torrent | Newbler -large | 0 | |
| Pride_LongTermAntibio | ELA2BSal | HostAssoc_Human Other | Human saliva | USA : California | 32.876609 -117.236885 | reads MG-RAST | Ion Torrent | Newbler -large | 0 | |
| Pride_LongTermAntibio | ELA2CSal | HostAssoc_Human Other | Human saliva | USA : California | 32.876609 -117.236885 | reads MG-RAST | Ion Torrent | Newbler -large | 0 | |
| Pride_LongTermAntibio | ELA33ASal | HostAssoc_Human Other | Human saliva | USA : California | 32.876609 -117.236885 | reads MG-RAST | Ion Torrent | Newbler -large | 0 | |
| Pride_LongTermAntibio | ELA33BSal | HostAssoc_Human Other | Human saliva | USA : California | 32.876609 -117.236885 | reads MG-RAST | Ion Torrent | Newbler -large | 0 | |
| Pride_LongTermAntibio | ELA33CSal | HostAssoc_Human Other | Human saliva | USA : California | 32.876609 -117.236885 | reads MG-RAST | Ion Torrent | Newbler -large | 0 | |
| Pride_LongTermAntibio | ELA3ASal | HostAssoc_Human Other | Human saliva | USA : California | 32.876609 -117.236885 | reads MG-RAST | Ion Torrent | Newbler -large | 0 | |
| Pride_LongTermAntibio | ELA3BSal | HostAssoc_Human Other | Human saliva | USA : California | 32.876609 -117.236885 | reads MG-RAST | Ion Torrent | Newbler -large | 0 | |
| Pride_LongTermAntibio | ELA3CSal | HostAssoc_Human Other | Human saliva | USA : California | 32.876609 -117.236885 | reads MG-RAST | Ion Torrent | Newbler -large | 0 | |
| Pride_LongTermAntibio | ELA4ASal | HostAssoc_Human Other | Human saliva | USA : California | 32.876609 -117.236885 | reads MG-RAST | Ion Torrent | Newbler -large | 0 | |
| Pride_LongTermAntibio | ELA4BSal | HostAssoc_Human Other | Human saliva | USA : California | 32.876609 -117.236885 | reads MG-RAST | Ion Torrent | Newbler -large | 0 | |
| Pride_LongTermAntibio | ELA4CSal | HostAssoc_Human Other | Human saliva | USA : California | 32.876609 -117.236885 | reads MG-RAST | Ion Torrent | Newbler -large | 0 | |
| Pride_LongTermAntibio | ELA7ASal | HostAssoc_Human Other | Human saliva | USA : California | 32.876609 -117.236885 | reads MG-RAST | Ion Torrent | Newbler -large | 0 | |
| Pride_LongTermAntibio | ELA7BSal | HostAssoc_Human Other | Human saliva | USA : California | 32.876609 -117.236885 | reads MG-RAST | Ion Torrent | Newbler -large | 0 | |
| Pride_LongTermAntibio | ELA7CSal | HostAssoc_Human Other | Human saliva | USA : California | 32.876609 -117.236885 | reads MG-RAST | Ion Torrent | Newbler -large | 0 | |
| Pride_LongTermAntibio | ELA8ASal | HostAssoc_Human Other | Human saliva | USA : California | 32.876609 -117.236885 | reads MG-RAST | Ion Torrent | Newbler -large | 0 | |
| Pride_LongTermAntibio | ELA8BSal | HostAssoc_Human Other | Human saliva | USA : California | 32.876609 -117.236885 | reads MG-RAST | Ion Torrent | Newbler -large | 0 | |
| Pride_LongTermAntibio | ELA8CSal | HostAssoc_Human Other | Human saliva | USA : California | 32.876609 -117.236885 | reads MG-RAST | Ion Torrent | Newbler -large | 0 | |
| Pride_LongTermAntibio | ELA9ASal | HostAssoc_Human Other | Human saliva | USA : California | 32.876609 -117.236885 | reads MG-RAST | Ion Torrent | Newbler -large | 0 | |
| Pride_LongTermAntibio | ELA9BSal | HostAssoc_Human Other | Human saliva | USA : California | 32.876609 -117.236885 | reads MG-RAST | Ion Torrent | Newbler -large | 0 | |
| Pride_LongTermAntibio | ELA9CSal | HostAssoc_Human Other | Human saliva | USA : California | 32.876609 -117.236885 | reads MG-RAST | Ion Torrent | Newbler -large | 0 | |
| Rosario_RNAwhitefly | Whitefly89 | HostAssoc_Insecta | Animal feces | USA : Florida | 29.411571 -82.108277 | reads Metavir | Solexa | IDBA_UD | 0 | Karyna Rosario. Heather Capobianco. Terry Fei Fan Ng. Mya Breitbart. and Jane E. Polston. RNA Viral Metagenome of Whiteflies Leads to the Discovery and Characterization of a Whitefly-Transmitted Carlavirus in North America. PLoS One. 2014; 9(1): e86748. |
| Rosario_RNAwhitefly | Whitefly90 | HostAssoc_Insecta | Animal feces | USA : Florida | 29.411571 -82.108277 | reads Metavir | Solexa | IDBA_UD | 0 | |
| Temmam_ArthroVirome | midges43 | HostAssoc_Insecta | Animal feces | Senegal | 14.059243 -16.288288 | reads Metavir | Illumina | IDBA_UD | 0 | Temmam S. Monteil-Bouchard S. Sambou M. Aubadie-Ladrix M. Azza S. Decloquement P. Bou Khalil JY. Baudoin J-P. Jardot P. Robert C. La Scola B. Mediannikov OY. Raoult D and Desnues C (2015) Faustovirus-Like Asfarvirus in Hematophagous Biting Midges and Their Vertebrate Hosts. Front. Microbiol. 6:1406. doi: 10.3389/fmicb.2015.01406 |
| Temmam_ArthroVirome | imicola44 | HostAssoc_Insecta | Animal feces | Senegal | 14.059243 -16.288288 | reads Metavir | Illumina | IDBA_UD | 0 | |
| Temmam_ArthroVirome | imicola45 | HostAssoc_Insecta | Animal feces | Senegal | 14.059243 -16.288288 | reads Metavir | Illumina | IDBA_UD | 0 | |
| Chen_yakDiarrhea | JNVQ011 | HostAssoc_MammalFeces | Animal feces | Tibet | 30.152745 88.787819 | contigs NCBI | no reads | SOAP De Novo | 0 | Chen X. Zhang B. Yue H. Wang Y. Zhou F. Zhang Q. Tang C. A novel astrovirus species in the gut of yaks with diarrhea in the Qinghai Tibetan plateau. 2013. J Gen Virol. 2015 Sep 29. doi: 10.1099/jgv.0.000303. |
| Sachsenroder_pigFeces | PigletC12 | HostAssoc_MammalFeces | Animal feces | Germany | 52.528393 13.400384 | reads SRA | 454 | Newbler | 0 | |
| Sachsenroder_pigFeces | AllVirPig | HostAssoc_MammalFeces | Animal feces | Germany | 52.528393 13.400384 | reads SRA | 454 | Newbler | 0 | |

| Project Name | Virome Code | Habitat Type (code) | Ecosystem Type (text) | Location | GPS | Source | Sequencing Technology | Assembly Software | # crucivirus | Reference |
|---|---|---|---|---|---|---|---|---|---|---|
| Sachsenroder_pigFeces | PigletC54 | HostAssoc_MammalFeces | Animal feces | Germany | 52.528393 13.400384 | reads SRA | 454 | Newbler | 0 | Sachsenröder J. Twardziok SO. Scheuch M. Johne R. The general composition of the faecal virome of pigs depends on age. but not on feeding with a probiotic bacterium. PLoS One. 2014 Feb 19;9(2):e88888. Doi: 10.1371/journal.pone.0088888. eCollection 2014. |
| Sachsenroder_pigFeces | PigletP12 | HostAssoc_MammalFeces | Animal feces | Germany | 52.528393 13.400384 | reads SRA | 454 | Newbler | 0 | |
| Sachsenroder_pigFeces | PigletP54 | HostAssoc_MammalFeces | Animal feces | Germany | 52.528393 13.400384 | reads SRA | 454 | Newbler | 0 | |
| Sachsenroder_pigFeces | SowC14pp | HostAssoc_MammalFeces | Animal feces | Germany | 52.528393 13.400384 | reads SRA | 454 | Newbler | 0 | |
| Sachsenroder_pigFeces | SowC28ap | HostAssoc_MammalFeces | Animal feces | Germany | 52.528393 13.400384 | reads SRA | 454 | Newbler | 0 | |
| Sachsenroder_pigFeces | SowP14pp | HostAssoc_MammalFeces | Animal feces | Germany | 52.528393 13.400384 | reads SRA | 454 | Newbler | 0 | |
| Shan_Pigfeces | Pig136828 | HostAssoc_MammalFeces | Animal feces | USA : North Carolina | 35.587952 -79.988361 | reads SRA | 454 | IDBA_UD | 0 | Shan T. Li L. Simmonds P. Wang C. Moeser A. Delwart E. The fecal virome of pigs on a high-density farm. J Virol. 2011 Nov;85(22):11697-708. doi: 10.1128/JVI.05217-11. |
| Woo_CamelFeces | camelFeces | HostAssoc_MammalFeces | Animal feces | UAE | 25.210798 55.289773 | reads SRA | HiSeq 2500 paired-end | IDBA_UD | 0 | Woo PC1. Lau SK2. Teng JL3. Tsang AK3. Joseph M4. Wong EY3. Tang Y3. Sivakumar S4. Bai R3. Wernery R4. Wernery U5. Yuen KY2. Metagenomic analysis of viromes of dromedary camel fecal samples reveals large number and high diversity of circoviruses and picobirnaviruses. Virology. 2014 Dec;471-473:117-25. doi: 10.1016/j.virol.2014.09.020. Epub 2014 Oct 29. |
| Wu_batGuano | Bat2063878 | HostAssoc_MammalFeces | Animal feces | China : Beijing | 39.920239 116.415238 | reads SRA | Illumina | IDBA_UD IGS | 0 | |
| Wu_batGuano | Bat2063879 | HostAssoc_MammalFeces | Animal feces | China : Beijing | 39.920239 116.415238 | reads SRA | Illumina | IDBA_UD IGS | 0 | |
| Wu_batGuano | Bat2063880 | HostAssoc_MammalFeces | Animal feces | China : Beijing | 39.920239 116.415238 | reads SRA | Illumina | IDBA_UD IGS | 0 | |
| Wu_batGuano | Bat2063881 | HostAssoc_MammalFeces | Animal feces | China : Beijing | 39.920239 116.415238 | reads SRA | Illumina | IDBA_UD IGS | 0 | |
| Wu_batGuano | Bat2063882 | HostAssoc_MammalFeces | Animal feces | China : Beijing | 39.920239 116.415238 | reads SRA | Illumina | IDBA_UD IGS | 0 | |
| Wu_batGuano | Bat2063883 | HostAssoc_MammalFeces | Animal feces | China : Beijing | 39.920239 116.415238 | reads SRA | Illumina | IDBA_UD IGS | 0 | |
| Wu_batGuano | Bat2063884 | HostAssoc_MammalFeces | Animal feces | China : Beijing | 39.920239 116.415238 | reads SRA | Illumina | IDBA_UD IGS | 0 | |
| Wu_batGuano | Bat2063885 | HostAssoc_MammalFeces | Animal feces | China : Beijing | 39.920239 116.415238 | reads SRA | Illumina | IDBA_UD IGS | 0 | |
| Wu_batGuano | Bat2063886 | HostAssoc_MammalFeces | Animal feces | China : Beijing | 39.920239 116.415238 | reads SRA | Illumina | IDBA_UD IGS | 0 | |
| Wu_batGuano | Bat2063887 | HostAssoc_MammalFeces | Animal feces | China : Beijing | 39.920239 116.415238 | reads SRA | Illumina | IDBA_UD IGS | 0 | |
| Wu_batGuano | Bat2063888 | HostAssoc_MammalFeces | Animal feces | China : Beijing | 39.920239 116.415238 | reads SRA | Illumina | IDBA_UD IGS | 0 | |
| Wu_batGuano | Bat2063891 | HostAssoc_MammalFeces | Animal feces | China : Beijing | 39.920239 116.415238 | reads SRA | Illumina | IDBA_UD IGS | 0 | |
| Wu_batGuano | Bat2063894 | HostAssoc_MammalFeces | Animal feces | China : Beijing | 39.920239 116.415238 | reads SRA | Illumina | IDBA_UD IGS | 0 | |
| Wu_batGuano | Bat2063898 | HostAssoc_MammalFeces | Animal feces | China : Beijing | 39.920239 116.415238 | reads SRA | Illumina | IDBA_UD IGS | 0 | |
| Wu_batGuano | Bat2063899 | HostAssoc_MammalFeces | Animal feces | China : Beijing | 39.920239 116.415238 | reads SRA | Illumina | IDBA_UD IGS | 0 | |
| Wu_batGuano | Bat2063900 | HostAssoc_MammalFeces | Animal feces | China : Beijing | 39.920239 116.415238 | reads SRA | Illumina | IDBA_UD IGS | 0 | |
| Wu_batGuano | Bat2063901 | HostAssoc_MammalFeces | Animal feces | China : Beijing | 39.920239 116.415238 | reads SRA | Illumina | IDBA_UD IGS | 0 | |
| Wu_batGuano | Bat2063902 | HostAssoc_MammalFeces | Animal feces | China : Beijing | 39.920239 116.415238 | reads SRA | Illumina | IDBA_UD IGS | 0 | |
| Wu_batGuano | Bat2063903 | HostAssoc_MammalFeces | Animal feces | China : Beijing | 39.920239 116.415238 | reads SRA | Illumina | IDBA_UD IGS | 0 | |
| Wu_batGuano | Bat2063904 | HostAssoc_MammalFeces | Animal feces | China : Beijing | 39.920239 116.415238 | reads SRA | Illumina | IDBA_UD IGS | 0 | |
| Wu_batGuano | Bat2063905 | HostAssoc_MammalFeces | Animal feces | China : Beijing | 39.920239 116.415238 | reads SRA | Illumina | IDBA_UD IGS | 0 | |
| Wu_batGuano | Bat2063906 | HostAssoc_MammalFeces | Animal feces | China : Beijing | 39.920239 116.415238 | reads SRA | Illumina | IDBA_UD IGS | 0 | |
| Wu_batGuano | Bat2063907 | HostAssoc_MammalFeces | Animal feces | China : Beijing | 39.920239 116.415238 | reads SRA | Illumina | IDBA_UD IGS | 0 | |
| Wu_batGuano | Bat2063908 | HostAssoc_MammalFeces | Animal feces | China : Beijing | 39.920239 116.415238 | reads SRA | Illumina | IDBA_UD IGS | 0 | |
| Wu_batGuano | Bat2063909 | HostAssoc_MammalFeces | Animal feces | China : Beijing | 39.920239 116.415238 | reads SRA | Illumina | IDBA_UD IGS | 0 | |
| Wu_batGuano | Bat2063910 | HostAssoc_MammalFeces | Animal feces | China : Beijing | 39.920239 116.415238 | reads SRA | Illumina | IDBA_UD IGS | 0 | |
| Wu_batGuano | Bat2063911 | HostAssoc_MammalFeces | Animal feces | China : Beijing | 39.920239 116.415238 | reads SRA | Illumina | IDBA_UD IGS | 0 | |
| Wu_batGuano | Bat2063912 | HostAssoc_MammalFeces | Animal feces | China : Beijing | 39.920239 116.415238 | reads SRA | Illumina | IDBA_UD IGS | 0 | |
| Wu_batGuano | Bat2063913 | HostAssoc_MammalFeces | Animal feces | China : Beijing | 39.920239 116.415238 | reads SRA | Illumina | IDBA_UD IGS | 0 | |
| Wu_batGuano | Bat2063914 | HostAssoc_MammalFeces | Animal feces | China : Beijing | 39.920239 116.415238 | reads SRA | Illumina | IDBA_UD IGS | 0 | |
| Wu_batGuano | Bat2063915 | HostAssoc_MammalFeces | Animal feces | China : Beijing | 39.920239 116.415238 | reads SRA | Illumina | IDBA_UD IGS | 0 | |
| Wu_batGuano | Bat2063916 | HostAssoc_MammalFeces | Animal feces | China : Beijing | 39.920239 116.415238 | reads SRA | Illumina | IDBA_UD IGS | 0 | |

| Project Name | Virome Code | Habitat Type (code) | Ecosystem Type (text) | Location | GPS | Source | Sequencing Technology | Assembly Software | # crucivirus | Reference |
|---|---|---|---|---|---|---|---|---|---|---|
| Wu_batGuano | Bat2063917 | HostAssoc_MammalFeces | Animal feces | China : Beijing | 39.920239 116.415238 | reads SRA | Illumina | IDBA_UD IGS | 0 | Wu Z. Yang L. Ren X. He G. Zhang J. Yang J. Qian Z. Dong J. Sun L. Zhu Y. Du J. Yang F. Zhang S. Jin Q. Deciphering the bat virome catalog to better understand the ecological diversity of bat viruses and the bat origin of emerging infectious diseases. ISME J. 2015 Aug 11. doi: 10.1038/ismej.2015.138. |
| Wu_batGuano | Bat2063918 | HostAssoc_MammalFeces | Animal feces | China : Beijing | 39.920239 116.415238 | reads SRA | Illumina | IDBA_UD IGS | 0 | |
| Wu_batGuano | Bat2063919 | HostAssoc_MammalFeces | Animal feces | China : Beijing | 39.920239 116.415238 | reads SRA | Illumina | IDBA_UD IGS | 0 | |
| Wu_batGuano | Bat2063920 | HostAssoc_MammalFeces | Animal feces | China : Beijing | 39.920239 116.415238 | reads SRA | Illumina | IDBA_UD IGS | 0 | |
| Wu_batGuano | Bat2063921 | HostAssoc_MammalFeces | Animal feces | China : Beijing | 39.920239 116.415238 | reads SRA | Illumina | IDBA_UD IGS | 1 | |
| Wu_batGuano | Bat2063922 | HostAssoc_MammalFeces | Animal feces | China : Beijing | 39.920239 116.415238 | reads SRA | Illumina | IDBA_UD IGS | 0 | |
| Wu_batGuano | Bat2063923 | HostAssoc_MammalFeces | Animal feces | China : Beijing | 39.920239 116.415238 | reads SRA | Illumina | IDBA_UD IGS | 0 | |
| Wu_batGuano | Bat2063924 | HostAssoc_MammalFeces | Animal feces | China : Beijing | 39.920239 116.415238 | reads SRA | Illumina | IDBA_UD IGS | 0 | |
| Wu_batGuano | Bat2063925 | HostAssoc_MammalFeces | Animal feces | China : Beijing | 39.920239 116.415238 | reads SRA | Illumina | IDBA_UD IGS | 0 | |
| Wu_batGuano | Bat2063926 | HostAssoc_MammalFeces | Animal feces | China : Beijing | 39.920239 116.415238 | reads SRA | Illumina | IDBA_UD IGS | 0 | |
| Wu_batGuano | Bat2063927 | HostAssoc_MammalFeces | Animal feces | China : Beijing | 39.920239 116.415238 | reads SRA | Illumina | IDBA_UD IGS | 0 | |
| Wu_batGuano | Bat2063928 | HostAssoc_MammalFeces | Animal feces | China : Beijing | 39.920239 116.415238 | reads SRA | Illumina | IDBA_UD IGS | 0 | |
| Wu_batGuano | Bat2063929 | HostAssoc_MammalFeces | Animal feces | China : Beijing | 39.920239 116.415238 | reads SRA | Illumina | IDBA_UD IGS | 0 | |
| Wu_batGuano | Bat2063931 | HostAssoc_MammalFeces | Animal feces | China : Beijing | 39.920239 116.415238 | reads SRA | Illumina | IDBA_UD IGS | 0 | |
| Wu_batGuano | Bat2063933 | HostAssoc_MammalFeces | Animal feces | China : Beijing | 39.920239 116.415238 | reads SRA | Illumina | IDBA_UD IGS | 0 | |
| Wu_batGuano | Bat2063934 | HostAssoc_MammalFeces | Animal feces | China : Beijing | 39.920239 116.415238 | reads SRA | Illumina | IDBA_UD IGS | 0 | |
| Wu_batGuano | Bat2063935 | HostAssoc_MammalFeces | Animal feces | China : Beijing | 39.920239 116.415238 | reads SRA | Illumina | IDBA_UD IGS | 0 | |
| Wu_batGuano | Bat2063936 | HostAssoc_MammalFeces | Animal feces | China : Beijing | 39.920239 116.415238 | reads SRA | Illumina | IDBA_UD IGS | 0 | |
| Wu_batGuano | Bat2063937 | HostAssoc_MammalFeces | Animal feces | China : Beijing | 39.920239 116.415238 | reads SRA | Illumina | IDBA_UD IGS | 0 | |
| Wu_batGuano | Bat2063938 | HostAssoc_MammalFeces | Animal feces | China : Beijing | 39.920239 116.415238 | reads SRA | Illumina | IDBA_UD IGS | 0 | |
| Wu_batGuano | Bat2063939 | HostAssoc_MammalFeces | Animal feces | China : Beijing | 39.920239 116.415238 | reads SRA | Illumina | IDBA_UD IGS | 0 | |
| Wu_batGuano | Bat2063940 | HostAssoc_MammalFeces | Animal feces | China : Beijing | 39.920239 116.415238 | reads SRA | Illumina | IDBA_UD IGS | 0 | |
| Wu_batGuano | Bat2063942 | HostAssoc_MammalFeces | Animal feces | China : Beijing | 39.920239 116.415238 | reads SRA | Illumina | IDBA_UD IGS | 0 | |
| Wu_batGuano | Bat2063943 | HostAssoc_MammalFeces | Animal feces | China : Beijing | 39.920239 116.415238 | reads SRA | Illumina | IDBA_UD IGS | 0 | |
| Wu_batGuano | Bat2063944 | HostAssoc_MammalFeces | Animal feces | China : Beijing | 39.920239 116.415238 | reads SRA | Illumina | IDBA_UD IGS | 0 | |
| Wu_batGuano | Bat2063945 | HostAssoc_MammalFeces | Animal feces | China : Beijing | 39.920239 116.415238 | reads SRA | Illumina | IDBA_UD IGS | 0 | |
| Wu_batGuano | Bat2063946 | HostAssoc_MammalFeces | Animal feces | China : Beijing | 39.920239 116.415238 | reads SRA | Illumina | IDBA_UD IGS | 0 | |
| Wu_batGuano | Bat2063948 | HostAssoc_MammalFeces | Animal feces | China : Beijing | 39.920239 116.415238 | reads SRA | Illumina | IDBA_UD IGS | 0 | |
| Wu_batGuano | Bat2063950 | HostAssoc_MammalFeces | Animal feces | China : Beijing | 39.920239 116.415238 | reads SRA | Illumina | IDBA_UD IGS | 0 | |
| Wu_batGuano | Bat2063951 | HostAssoc_MammalFeces | Animal feces | China : Beijing | 39.920239 116.415238 | reads SRA | Illumina | IDBA_UD IGS | 0 | |
| Wu_batGuano | Bat2063952 | HostAssoc_MammalFeces | Animal feces | China : Beijing | 39.920239 116.415238 | reads SRA | Illumina | IDBA_UD IGS | 0 | |
| Wu_batGuano | Bat2063954 | HostAssoc_MammalFeces | Animal feces | China : Beijing | 39.920239 116.415238 | reads SRA | Illumina | IDBA_UD IGS | 0 | |
| Wu_batGuano | Bat2063955 | HostAssoc_MammalFeces | Animal feces | China : Beijing | 39.920239 116.415238 | reads SRA | Illumina | IDBA_UD IGS | 0 | |
| Wu_batGuano | Bat2063956 | HostAssoc_MammalFeces | Animal feces | China : Beijing | 39.920239 116.415238 | reads SRA | Illumina | IDBA_UD IGS | 0 | |
| Wu_batGuano | Bat2063957 | HostAssoc_MammalFeces | Animal feces | China : Beijing | 39.920239 116.415238 | reads SRA | Illumina | IDBA_UD IGS | 0 | |
| Wu_batGuano | Bat2063958 | HostAssoc_MammalFeces | Animal feces | China : Beijing | 39.920239 116.415238 | reads SRA | Illumina | IDBA_UD IGS | 0 | |
| Wu_batGuano | Bat2063960 | HostAssoc_MammalFeces | Animal feces | China : Beijing | 39.920239 116.415238 | reads SRA | Illumina | IDBA_UD IGS | 0 | |
| Wu_batGuano | Bat2063961 | HostAssoc_MammalFeces | Animal feces | China : Beijing | 39.920239 116.415238 | reads SRA | Illumina | IDBA_UD IGS | 0 | |
| Wu_batGuano | Bat2063962 | HostAssoc_MammalFeces | Animal feces | China : Beijing | 39.920239 116.415238 | reads SRA | Illumina | IDBA_UD IGS | 0 | |
| Wu_batGuano | Bat2063967 | HostAssoc_MammalFeces | Animal feces | China : Beijing | 39.920239 116.415238 | reads SRA | Illumina | IDBA_UD IGS | 0 | |
| Wu_batGuano | Bat546088 | HostAssoc_MammalFeces | Animal feces | China : Beijing | 39.920239 116.415238 | reads SRA | Illumina | IDBA_UD IGS | 0 | |
| Wu_batGuano | Bat546089 | HostAssoc_MammalFeces | Animal feces | China : Beijing | 39.920239 116.415238 | reads SRA | Illumina | IDBA_UD IGS | 0 | |

| Project Name | Virome Code | Habitat Type (code) | Ecosystem Type (text) | Location | GPS | Source | Sequencing Technology | Assembly Software | # crucivirus | Reference |
|---|---|---|---|---|---|---|---|---|---|---|
| Zhang_catFeces | catFeces | HostAssoc_MammalFeces | Animal feces | USA : California | 37.778896 -122.446804 | reads SRA | | Newbler | 0 | Zhang W. Li L. Deng X. Kapusinszky B. Pesavento PA. Delwart E. Faecal virome of cats in an animal shelter. J Gen Virol. 2014 Nov;95(Pt 11):2553-64. doi: 10.1099/ vir.0.069674-0. Epub 2014 Jul 30. |
| Coetzee_VineyardRNA | RNAVine | HostAssoc_Plants | | South Africa : Stellenbosch | -33.935322 18.855379 | contigs Metavir | no reads | | 0 | Coetzee B1. Freeborough MJ. Maree HJ. Celton JM. Rees DJ. Burger JT. Deep sequencing analysis of viruses infecting grapevines: Virome of a vineyard. Virology. 2010 May 10;400(2):157-63. doi: 10.1016/j.virol.2010.01.023. |
| Hansen_CressCompletRattusFeces | cressComp | HostAssoc_Rodent Feces | Animal feces | Malaysia and Denmark | | contigs NCBI | | | 0 | Hansen TA. Fridholm H. Frøslev TG. Kjartansdóttir KR. Willerslev E. Nielsen LP. Hansen AJ. New Type of Papillomavirus and Novel Circular Single Stranded DNA Virus Discovered in Urban Rattus norvegicus Using Circular DNA Enrichment and Metagenomics. PLoS One. 2015 Nov 11;10(11):e0141952. doi: 10.1371/ journal.pone.0141952 |
| Modi_MouseFeces | AmpiCtrl | HostAssoc_Rodent Feces | Animal feces | USA : Massachusetts | 41.732542 -72.793500 | reads SRA | 454 GS FLX | Newbler | 0 | Modi SR. Lee HH. Spina CS. Collins JJ. Antibiotic treatment expands the resistance reservoir and ecological network of the phage metagenome. Nature. 2013 Jul 11;499(7457):219-22. doi: 10.1038/nature12212. |
| Modi_MouseFeces | AmpiTreat | HostAssoc_Rodent Feces | Animal feces | USA : Massachusetts | 41.732542 -72.793500 | reads SRA | 454 GS FLX | Newbler | 0 | |
| Modi_MouseFeces | CiproCtrl | HostAssoc_Rodent Feces | Animal feces | USA : Massachusetts | 41.732542 -72.793500 | reads SRA | 454 GS FLX | Newbler | 0 | |
| Modi_MouseFeces | CiproTreat | HostAssoc_Rodent Feces | Animal feces | USA : Massachusetts | 41.732542 -72.793500 | reads SRA | 454 GS FLX | Newbler | 0 | |
| Reyes_Gnotobiotic Mouse | Coproseq | HostAssoc_Rodent Feces | Animal feces | USA : Missouri | 38.635007 -90.262387 | reads ENA | Illumina | IDBA_UD | 0 | |
| Reyes_Gnotobiotic Mouse | 01HAE4ODE | HostAssoc_Rodent Feces | Animal feces | USA : Missouri | 38.635007 -90.262387 | reads ENA | 454 GS FLX pyrosequencer (Roche) | Newbler | 0 | |
| Reyes_Gnotobiotic Mouse | 08G3ZYMMR | HostAssoc_Rodent Feces | Animal feces | USA : Missouri | 38.635007 -90.262387 | reads ENA | 454 GS FLX pyrosequencer (Roche) | Newbler | 0 | |
| Reyes_Gnotobiotic Mouse | 09HAE4ODE | HostAssoc_Rodent Feces | Animal feces | USA : Missouri | 38.635007 -90.262387 | reads ENA | 454 GS FLX pyrosequencer (Roche) | Newbler | 0 | |
| Reyes_Gnotobiotic Mouse | 09HAE4ODE2 | HostAssoc_Rodent Feces | Animal feces | USA : Missouri | 38.635007 -90.262387 | reads ENA | 454 GS FLX pyrosequencer (Roche) | Newbler | 0 | |
| Reyes_Gnotobiotic Mouse | 09HCZ1WD0 | HostAssoc_Rodent Feces | Animal feces | USA : Missouri | 38.635007 -90.262387 | reads ENA | 454 GS FLX pyrosequencer (Roche) | Newbler | 0 | |
| Reyes_Gnotobiotic Mouse | 10GVVF7D3 | HostAssoc_Rodent Feces | Animal feces | USA : Missouri | 38.635007 -90.262387 | reads ENA | 454 GS FLX pyrosequencer (Roche) | Newbler | 0 | |
| Reyes_Gnotobiotic Mouse | 10HAE4ODE | HostAssoc_Rodent Feces | Animal feces | USA : Missouri | 38.635007 -90.262387 | reads ENA | 454 GS FLX pyrosequencer (Roche) | Newbler | 0 | |
| Reyes_Gnotobiotic Mouse | 10HAE4ODE2 | HostAssoc_Rodent Feces | Animal feces | USA : Missouri | 38.635007 -90.262387 | reads ENA | 454 GS FLX pyrosequencer (Roche) | Newbler | 0 | |
| Reyes_Gnotobiotic Mouse | 10HAE4ODE3 | HostAssoc_Rodent Feces | Animal feces | USA : Missouri | 38.635007 -90.262387 | reads ENA | 454 GS FLX pyrosequencer (Roche) | Newbler | 0 | |
| Reyes_Gnotobiotic Mouse | 10HCZ1WD0 | HostAssoc_Rodent Feces | Animal feces | USA : Missouri | 38.635007 -90.262387 | reads ENA | 454 GS FLX pyrosequencer (Roche) | Newbler | 0 | |
| Reyes_Gnotobiotic Mouse | 10HDVKH9M | HostAssoc_Rodent Feces | Animal feces | USA : Missouri | 38.635007 -90.262387 | reads ENA | 454 GS FLX pyrosequencer (Roche) | Newbler | 0 | |
| Reyes_Gnotobiotic Mouse | 12GVVF7D3 | HostAssoc_Rodent Feces | Animal feces | USA : Missouri | 38.635007 -90.262387 | reads ENA | 454 GS FLX pyrosequencer (Roche) | Newbler | 0 | |
| Reyes_Gnotobiotic Mouse | 12GVVF7D32 | HostAssoc_Rodent Feces | Animal feces | USA : Missouri | 38.635007 -90.262387 | reads ENA | 454 GS FLX pyrosequencer (Roche) | Newbler | 0 | |
| Reyes_Gnotobiotic Mouse | 12GVVF7D33 | HostAssoc_Rodent Feces | Animal feces | USA : Missouri | 38.635007 -90.262387 | reads ENA | 454 GS FLX pyrosequencer (Roche) | Newbler | 0 | |
| Reyes_Gnotobiotic Mouse | 23GVL26ZJ | HostAssoc_Rodent Feces | Animal feces | USA : Missouri | 38.635007 -90.262387 | reads ENA | 454 GS FLX pyrosequencer (Roche) | Newbler | 0 | |
| Reyes_Gnotobiotic Mouse | 23GVRF1TC | HostAssoc_Rodent Feces | Animal feces | USA : Missouri | 38.635007 -90.262387 | reads ENA | 454 GS FLX pyrosequencer (Roche) | Newbler | 0 | |
| Reyes_Gnotobiotic Mouse | 23GVVF7D3 | HostAssoc_Rodent Feces | Animal feces | USA : Missouri | 38.635007 -90.262387 | reads ENA | 454 GS FLX pyrosequencer (Roche) | Newbler | 0 | |
| Reyes_Gnotobiotic Mouse | 24HAE4ODE | HostAssoc_Rodent Feces | Animal feces | USA : Missouri | 38.635007 -90.262387 | reads ENA | 454 GS FLX pyrosequencer (Roche) | Newbler | 0 | |
| Reyes_Gnotobiotic Mouse | 24HCZ1WD0 | HostAssoc_Rodent Feces | Animal feces | USA : Missouri | 38.635007 -90.262387 | reads ENA | 454 GS FLX pyrosequencer (Roche) | Newbler | 0 | |
| Reyes_Gnotobiotic Mouse | 34GVVF7D3 | HostAssoc_Rodent Feces | Animal feces | USA : Missouri | 38.635007 -90.262387 | reads ENA | 454 GS FLX pyrosequencer (Roche) | Newbler | 0 | |

| Project Name | Virome Code | Habitat Type (code) | Ecosystem Type (text) | Location | GPS | Source | Sequencing Technology | Assembly Software | # crucivirus | Reference |
|---|---|---|---|---|---|---|---|---|---|---|
| Reyes_Gnotobiotic Mouse | 34GVVF7D32 | HostAssoc_Rodent Feces | Animal feces | USA : Missouri | 38.635007 -90.262387 | reads ENA | 454 GS FLX pyrosequencer (Roche) | Newbler | 0 | Reyes A. Wu M. McNulty NP. Rohwer FL. Gordon JI. Gnotobiotic mouse model of phage-bacterial host dynamics in the human gut. Proc Natl Acad Sci U S A. 2013 Dec 10;110(50):20236-41. doi: 10.1073/pnas.1319470110. Epub 2013 Nov 20. |
| Reyes_Gnotobiotic Mouse | 34HAE4ODE | HostAssoc_Rodent Feces | Animal feces | USA : Missouri | 38.635007 -90.262387 | reads ENA | 454 GS FLX pyrosequencer (Roche) | Newbler | 0 | |
| Reyes_Gnotobiotic Mouse | 45GVL26ZJ | HostAssoc_Rodent Feces | Animal feces | USA : Missouri | 38.635007 -90.262387 | reads ENA | 454 GS FLX pyrosequencer (Roche) | Newbler | 0 | |
| Reyes_Gnotobiotic Mouse | 45GVRF1TC | HostAssoc_Rodent Feces | Animal feces | USA : Missouri | 38.635007 -90.262387 | reads ENA | 454 GS FLX pyrosequencer (Roche) | Newbler | 0 | |
| Reyes_Gnotobiotic Mouse | 45GVVF7D3 | HostAssoc_Rodent Feces | Animal feces | USA : Missouri | 38.635007 -90.262387 | reads ENA | 454 GS FLX pyrosequencer (Roche) | Newbler | 0 | |
| Reyes_Gnotobiotic Mouse | 45GVVF7D32 | HostAssoc_Rodent Feces | Animal feces | USA : Missouri | 38.635007 -90.262387 | reads ENA | 454 GS FLX pyrosequencer (Roche) | Newbler | 0 | |
| Reyes_Gnotobiotic Mouse | 51HAE4ODE | HostAssoc_Rodent Feces | Animal feces | USA : Missouri | 38.635007 -90.262387 | reads ENA | 454 GS FLX pyrosequencer (Roche) | Newbler | 0 | |
| Reyes_Gnotobiotic Mouse | 67G3ZYMMR | HostAssoc_Rodent Feces | Animal feces | USA : Missouri | 38.635007 -90.262387 | reads ENA | 454 GS FLX pyrosequencer (Roche) | Newbler | 0 | |
| Reyes_Gnotobiotic Mouse | 67G3ZYMMR2 | HostAssoc_Rodent Feces | Animal feces | USA : Missouri | 38.635007 -90.262387 | reads ENA | 454 GS FLX pyrosequencer (Roche) | Newbler | 0 | |
| Reyes_Gnotobiotic Mouse | 78GVVF7D3 | HostAssoc_Rodent Feces | Animal feces | USA : Missouri | 38.635007 -90.262387 | reads ENA | 454 GS FLX pyrosequencer (Roche) | Newbler | 0 | |
| Reyes_Gnotobiotic Mouse | 78GVVF7D32 | HostAssoc_Rodent Feces | Animal feces | USA : Missouri | 38.635007 -90.262387 | reads ENA | 454 GS FLX pyrosequencer (Roche) | Newbler | 0 | |
| Reyes_Gnotobiotic Mouse | 78HAE4ODE | HostAssoc_Rodent Feces | Animal feces | USA : Missouri | 38.635007 -90.262387 | reads ENA | 454 GS FLX pyrosequencer (Roche) | Newbler | 0 | |
| Reyes_Gnotobiotic Mouse | 78HCZ1WD0 | HostAssoc_Rodent Feces | Animal feces | USA : Missouri | 38.635007 -90.262387 | reads ENA | 454 GS FLX pyrosequencer (Roche) | Newbler | 0 | |
| Reyes_Gnotobiotic Mouse | 89GVVF7D3 | HostAssoc_Rodent Feces | Animal feces | USA : Missouri | 38.635007 -90.262387 | reads ENA | 454 GS FLX pyrosequencer (Roche) | Newbler | 0 | |
| Reyes_Gnotobiotic Mouse | 89HAE4ODE | HostAssoc_Rodent Feces | Animal feces | USA : Missouri | 38.635007 -90.262387 | reads ENA | 454 GS FLX pyrosequencer (Roche) | Newbler | 0 | |
| Reyes_Gnotobiotic Mouse | lumGVL26ZJ | HostAssoc_Rodent Feces | Animal feces | USA : Missouri | 38.635007 -90.262387 | reads ENA | 454 GS FLX pyrosequencer (Roche) | Newbler | 0 | |
| Reyes_Gnotobiotic Mouse | lumGVRF1TC | HostAssoc_Rodent Feces | Animal feces | USA : Missouri | 38.635007 -90.262387 | reads ENA | 454 GS FLX pyrosequencer (Roche) | Newbler | 0 | |
| Reyes_Gnotobiotic Mouse | M10G4PULPT | HostAssoc_Rodent Feces | Animal feces | USA : Missouri | 38.635007 -90.262387 | reads ENA | 454 GS FLX pyrosequencer (Roche) | Newbler | 0 | |
| Reyes_Gnotobiotic Mouse | M1G3ZYMMR | HostAssoc_Rodent Feces | Animal feces | USA : Missouri | 38.635007 -90.262387 | reads ENA | 454 GS FLX pyrosequencer (Roche) | Newbler | 0 | |
| Reyes_Gnotobiotic Mouse | M2G3ZYMMR | HostAssoc_Rodent Feces | Animal feces | USA : Missouri | 38.635007 -90.262387 | reads ENA | 454 GS FLX pyrosequencer (Roche) | Newbler | 0 | |
| Reyes_Gnotobiotic Mouse | M3G4PULPT | HostAssoc_Rodent Feces | Animal feces | USA : Missouri | 38.635007 -90.262387 | reads ENA | 454 GS FLX pyrosequencer (Roche) | Newbler | 0 | |
| Reyes_Gnotobiotic Mouse | M3HCZ1WD0 | HostAssoc_Rodent Feces | Animal feces | USA : Missouri | 38.635007 -90.262387 | reads ENA | 454 GS FLX pyrosequencer (Roche) | Newbler | 0 | |
| Reyes_Gnotobiotic Mouse | M4G4PULPT | HostAssoc_Rodent Feces | Animal feces | USA : Missouri | 38.635007 -90.262387 | reads ENA | 454 GS FLX pyrosequencer (Roche) | Newbler | 0 | |
| Reyes_Gnotobiotic Mouse | M5G4PULPT | HostAssoc_Rodent Feces | Animal feces | USA : Missouri | 38.635007 -90.262387 | reads ENA | 454 GS FLX pyrosequencer (Roche) | Newbler | 0 | |
| Reyes_Gnotobiotic Mouse | M5HAE4ODE | HostAssoc_Rodent Feces | Animal feces | USA : Missouri | 38.635007 -90.262387 | reads ENA | 454 GS FLX pyrosequencer (Roche) | Newbler | 0 | |
| Reyes_Gnotobiotic Mouse | M5HCZ1WD0 | HostAssoc_Rodent Feces | Animal feces | USA : Missouri | 38.635007 -90.262387 | reads ENA | 454 GS FLX pyrosequencer (Roche) | Newbler | 0 | |
| Reyes_Gnotobiotic Mouse | M6G4PULPT | HostAssoc_Rodent Feces | Animal feces | USA : Missouri | 38.635007 -90.262387 | reads ENA | 454 GS FLX pyrosequencer (Roche) | Newbler | 0 | |
| Reyes_Gnotobiotic Mouse | M7G4PULPT | HostAssoc_Rodent Feces | Animal feces | USA : Missouri | 38.635007 -90.262387 | reads ENA | 454 GS FLX pyrosequencer (Roche) | Newbler | 0 | |
| Reyes_Gnotobiotic Mouse | M8G4PULPT | HostAssoc_Rodent Feces | Animal feces | USA : Missouri | 38.635007 -90.262387 | reads ENA | 454 GS FLX pyrosequencer (Roche) | Newbler | 0 | |

| Project Name | Virome Code | Habitat Type (code) | Ecosystem Type (text) | Location | GPS | Source | Sequencing Technology | Assembly Software | # crucivirus | Reference |
|---|---|---|---|---|---|---|---|---|---|---|
| Reyes_Gnotobiotic Mouse | M9G4PULPT | HostAssoc_Rodent Feces | Animal feces | USA : Missouri | 38.635007 -90.262387 | reads ENA | 454 GS FLX pyrosequencer (Roche) | Newbler | 0 | |
