## Supplementary material for "Unveiling Crucivirus Diversity by Mining Metagenomic Data": Supp. Table 3

### Supplementary Table S3: Custom library for crucivirus CDS annotation

## CP

YP_009337040
ABV21601
ABV21605
ABV21600
KX388503.1 - capsid protein translation
KX388519.1 - capsid protein translation
KX388520.1 - capsid protein translation
KX388513.1 - capsid protein translation
KC248412.1 - Capsid protein translation
KX388525.1 - capsid protein translation
KX388523.1 - capsid protein translation
KX388524.1 - capsid protein translation
KX388526.1 - capsid protein translation
KX388521.1 - capsid protein translation
KX388522.1 - capsid protein translation
KF133808 - Capsid protein translation
KX388512.1 - capsid protein translation
KX388494.1 - capsid protein translation
KX388496.1 - capsid protein translation
KX388511.1 - capsid protein translation
KX388517.1 - capsid protein translation
KX388518.1 - capsid protein translation
KX388497.1 - capsid protein translation
KX388510.1 - capsid protein translation
KX388509.1 - capsid protein translation
KX388527.1 - capsid protein translation
KX388528.1 - capsid protein translation
KX388529.1 - capsid protein translation
KX388530.1 - capsid protein translation
KX388507.1 - capsid protein translation
KX388505.1 - capsid protein translation
KX388506.1 - capsid protein translation
KX388500.1 - capsid protein translation
KX388501.1 - capsid protein translation
KX388498.1 - capsid protein translation
KX388499.1 - capsid protein translation
KX388508.1 - capsid protein translation
KX388516.1 - capsid protein translation
KX388502.1 - capsid protein translation
KX388495.1 - capsid protein translation
KX388504.1 - capsid protein translation
KX388498.1 - capsid protein translation
KX388515.1 - Capsid protein translation
AQU11754
KX388514.1 - Capsid protein translation
YP_009336611
AHA86933
AHA86933
YP_009337040
YP_009337040
YP_009342257
YP_009342257
ABV21601
ABV21605
ABV21600
AHA86933
YP_009337040
YP_009342257
AQU11790
AXH74444
AQR57899
BAF37070
AXH74994
AQU11760
AXH74298
AUF34965
AQU11721
AQU11716
AQU11732
AQU11735
AQU11713
AHA86933
YP_009342257
NP_995579
NP_995579
AQU11727
AQU11727
BAF37070
BAF37070
AXH77252
AXH77252
AXH74444
AXH74444
YP_009237532
YP_009237532
AQU11772
AQU11772
AQU11775
AQU11775
YP_009337040
YP_009337040
AXH74994
AXH74994
AHA86933
AHA86933
YP_009252339
YP_009252339
AXH73059
AXH73059
YP_009336611
YP_009336611
AQU11782
AQU11782
AQU11784
AQU11784
AQU11778
AQU11778
AQR57899
AQR57899
ADK55584
ADK55584
ADK55585
ADK55585
ADK55582
ADK55582
ADK55583
ADK55583
YP_009342300
YP_009342300
YP_009336592
YP_009336592
YP_009337418
YP_009337418
YP_009109631
YP_009109631
YP_009337312
YP_009337312
AQU11750
AQU11750
AUM61786
AUM61786
AUM61959
AUM61959
AQU11701
AQU11701
AQU11708
AQU11708
AUM61905
AUM61905
AQU11748
AQU11748
AQU11763
AQU11763
AQU11713
AQU11713
AQU11745
AQU11745
AQU11742
AQU11742
AQU11790
AQU11790
AXH77579
AXH77579
AUM61616
AUM61616
AQU11735
AQU11735
APG76598
APG76598
AXL65908
AXL65908
AUF34965
AUF34965
AKM12421
AKM12421
YP_009094499
YP_009094499
AQU11732
AQU11732
AQU11734
AQU11734
AXH77953
AXH77953
AUM61901
AUM61901
AXH76175
AXH76175
AQU11721
AQU11721
AQU11716
AQU11716
AIF34804
AIF34804
YP_009142777
YP_009142777
AQU11739
AQU11739
AXH74298
AXH74298
AXH74708
AXH74708
YP_009336686
YP_009336686
AQU11760
AQU11760
AYF34965
AYF34965
YP_009342257
YP_009342257
AHA86928
AHA86928
AOV86262
AOV86262
AQU11725
AQU11725
AQU11706
AQU11706
AQU11730
AQU11730
AXH76175
YP_009094499
YP_009336686
AQU11701
YP_009237532
AIF34804
AKM12421
AQU11734
AQU11745
AQU11727
AQU11775
AXH77953
AXH77252
AQU11748
YP_009337418
NP_995579
AHA86933
AQU11754
AXH73059
AQU11730
AHA86928
AQU11725
YP_009109631
AQU11750
AQU11706
ABV21600
ABV21601
ADK55583
ADK55584
ADK55585
ABV21605
ADK55582
YP_009342300
YP_009336592
YP_009337312
AQU11742
AQU11763
AQU11782
AQU11784
AQU11778
AQU11754
AQU11772
AXH74708
AQU11716
AQU11721
AQU11708
YP_009142777
YP_009342257
AQU11739
YP_009337040

### Rep

KP005453 (modified) - Replication associated protein interval 2 translation - Replication associated protein
KP005454 (modified) - Replication associated protein interval 2 translation - Replication associated protein
KT149398 (modified) - hypothetical protein CDS translation - hypothetical protein CDS
KT149398 (modified) - hypothetical protein CDS translation 3 - hypothetical protein CDS
KT149398 (modified) - hypothetical protein CDS translation 3 - hypothetical protein CDS
KX618694 - TrAP translation - Ren
KT149408 (modified) - hypothetical protein CDS translation 2 - hypothetical protein CDS
KT149408 (modified) - Replication associated protein translation - hypothetical protein CDS
KT149408 (modified) - hypothetical protein CDS translation 2 - Replication associated protein
MH425572.1 - Replication associated protein interval 2 translation - Replication associated protein
U49907 (modified) - Replication-associated protein translation - C2 CDS
KX618694 - Replication associated protein translation - TrAP
YP_009021245 - Viral Rep
YP_009021245 - Viral Rep
KT149395 (modified) - hypothetical protein CDS translation 2 - hypothetical protein CDS
KT149395 (modified) - Replication associated protein translation - hypothetical protein CDS
KT149395 (modified) - hypothetical protein CDS translation 2 - Replication associated protein
AOV86234 - Viral Rep
AOV86234 - Viral Rep
AXH73056 - Viral Rep
AXH73056 - Viral Rep
YP_009109643 - Viral Rep
YP_009109643 - Viral Rep
ARI44308 - Viral Rep
ARI44308 - Viral Rep
AHH31400 - Viral Rep
AHH31400 - Viral Rep
AEL28791 - Viral Rep
AEL28791 - Viral Rep
YP_764455 - Viral Rep
YP_764455 - Viral Rep
KC248420 (modified) - Replication associated protein translation - hypothetical protein CDS
KY487822.1 - Replication associated protein translation - Replication associated protein
AIF34812 - Gemini AL1 M
AIF34812 - Gemini AL1 M
KT945154 (modified) - Hypothetical 2 translation - hypothetical protein CDS
KT945154 (modified) - Replication associated protein translation - Hypothetical 2
MG748718.1 - Replication associated protein translation - hypothetical protein CDS
MG748719.1 - Replication associated protein translation - hypothetical protein CDS
MG748720.1 - Replication associated protein translation - hypothetical protein CDS
MG748718.1 - Replication associated protein translation - Replication associated protein
MG748719.1 - Replication associated protein translation - Replication associated protein
MG748720.1 - Replication associated protein translation - Replication associated protein
MG748715.1 - Replication associated protein translation - hypothetical protein CDS
MG748715.1 - Replication associated protein translation - Replication associated protein
MG748716.1 - Replication associated protein translation - hypothetical protein CDS
MG748716.1 - Replication associated protein translation - Replication associated protein
MG748717.1 - Replication associated protein translation - hypothetical protein CDS
MG748717.1 - Replication associated protein translation - Replication associated protein
KT149403 (modified) - hypothetical protein CDS translation 2 - hypothetical protein CDS
KT149403 (modified) - Replication associated protein translation - hypothetical protein CDS
KT149403 (modified) - hypothetical protein CDS translation 2 - Replication associated protein
AUF34964 - CDS
AUF34964 - putative replication-associated protein
AUF34964 - Viral Rep
AUF34964 - Viral Rep
AUF34964 - Viral Rep
YP_009116902 - Viral Rep
YP_009116902 - Viral Rep
AEL28793 - Viral Rep
AEL28793 - Viral Rep
AWW06057 - Viral Rep
AWW06057 - Viral Rep
KJ437671 (modified) - Replication-associated protein interval 2 translation - RepA CDS
KJ437671 (modified) - Replication-associated protein interval 2 translation - Replication-associated protein
ALE29827 - Viral Rep
ALE29827 - Viral Rep
AMD39533 - Viral Rep
AMD39533 - Viral Rep
AVX29443 - Viral Rep
AVX29443 - Viral Rep
AXH74443 - Viral Rep
AXH74443 - Viral Rep
AGA18441 - Viral Rep
AGA18441 - Viral Rep
KC248414 (modified) - Replication associated protein translation - ORF 2 (frame 1)
AIF76259 - Viral Rep
AIF76259 - Viral Rep
KT214386 (modified) - Replication-associated protein interval 1 translation - C3 CDS
KT214386 (modified) - C3 CDS translation - Replication-associated protein
KT214386 (modified) - C3 CDS translation - C3 CDS
KT214386 (modified) - RepA CDS translation - C3 CDS
KT214386 (modified) - C3 CDS translation - RepA CDS
YP_009163920 - Viral Rep
YP_009163920 - Viral Rep
AIF34802 - Viral Rep
AIF34802 - Viral Rep
GU734126 (modified) - Replication-associated protein translation - C2 CDS
KM386645 (modified) - Replication-associated protein translation - TrAP
AIF76277 - Viral Rep
AIF76277 - Viral Rep
KP153500 (modified) - Hypothetical 1 translation - Hypothetical 1
KP153500 (modified) - Hypothetical 2 translation - Hypothetical 1
KP153500 (modified) - Hypothetical 1 translation - Hypothetical 2
KF133827 - hypothetical protein CDS translation - hypothetical protein CDS
KT149401 (modified) - hypothetical protein CDS translation 2 - Replication associated protein
YP_009126898 - Viral Rep
YP_009126898 - Viral Rep
AGA18265 - Viral Rep
AGA18265 - Viral Rep
AIX11626 - Viral Rep
AIX11626 - Viral Rep
AUW34331 - Viral Rep
AUW34331 - Viral Rep
YP_009109675 - Viral Rep
YP_009109675 - Viral Rep
KU203351 (modified) - ORF 6 (frame 3) translation - ORF 7 (frame 2)
AF379637 (modified) - L4 CDS translation - Replication-associated protein
MH545516.1 - Replication associated protein interval 2 translation - Replication associated protein
AGG39817 - Viral Rep
AGG39817 - Viral Rep
AJM89742 - Viral Rep
AJM89742 - Viral Rep
WP_027090230 - other interval 2
WP_027090230 - other interval 2
WP_027090230 - other interval 2
WP_027090230 - other interval 2
WP_027090230 - other interval 2
YP_009237578 - other interval 2
YP_009237578 - other interval 2
YP_009237578 - other interval 2
YP_009237578 - other interval 2
KY487791.1 - Replication associated protein translation - Replication associated protein
KT149398 (modified) - hypothetical protein CDS translation 2 - hypothetical protein CDS
KT149398 (modified) - Replication associated protein translation - hypothetical protein CDS
KT149398 (modified) - hypothetical protein CDS translation 2 - Replication associated protein
KT149396 (modified) - hypothetical protein CDS translation 2 - hypothetical protein CDS
KT149396 (modified) - Replication associated protein translation - hypothetical protein CDS
KT149396 (modified) - hypothetical protein CDS translation 2 - Replication associated protein
KR134313 (modified) - RNA-binding protein CDS translation - Replication associated protein
KR134313 (modified) - Replication associated protein translation - RNA-binding protein CDS
KR134313 (modified) - RNA-binding protein CDS translation - RNA-binding protein CDS
KT149404 (modified) - hypothetical protein CDS translation 2 - hypothetical protein CDS
KT149404 (modified) - Replication associated protein translation - hypothetical protein CDS
KT149404 (modified) - hypothetical protein CDS translation 2 - Replication associated protein
YP_009126881 - CDS
YP_009126881 - replication-associated protein
YP_009126881 - Viral Rep
YP_009126881 - Viral Rep
YP_009126881 - Viral Rep
KT214389 (modified) - Replication-associated protein interval 2 translation - RepA CDS
KT214389 (modified) - Replication-associated protein interval 2 translation - Replication-associated protein
KT214389 (modified) - Replication-associated protein interval 1 translation - C3 CDS
KT214389 (modified) - C3 CDS translation - Replication-associated protein
KT214389 (modified) - C3 CDS translation - C3 CDS
KT214389 (modified) - RepA CDS translation - C3 CDS
KT214389 (modified) - C3 CDS translation - RepA CDS
KX388527.1 - Replication associated protein translation - Replication associated protein
KX388527.1 - Replication associated protein translation - stem loop
KX388527.1 - Replication associated protein translation - stem loop
KJ955447 (modified) - viral RNA-binding protein CDS translation - Replication associated protein
KJ955448 (modified) - viral RNA-binding protein CDS translation - Replication associated protein
KJ955449 (modified) - viral RNA-binding protein CDS translation - Replication associated protein
KJ955451 (modified) - viral RNA-binding protein CDS translation - Replication associated protein
KR134311 (modified) - RNA-binding protein CDS translation - Replication associated protein
KR134312 (modified) - RNA-binding protein CDS translation - Replication associated protein
KR134314 (modified) - RNA-binding protein CDS translation - Replication associated protein
KR134315 (modified) - RNA-binding protein CDS translation - Replication associated protein
KR134316 (modified) - RNA-binding protein CDS translation - Replication associated protein
KR134317 (modified) - RNA-binding protein CDS translation - Replication associated protein
KR134318 (modified) - RNA-binding protein CDS translation - Replication associated protein
KR134319 (modified) - RNA-binding protein CDS translation - Replication associated protein
KR134320 (modified) - RNA-binding protein CDS translation - Replication associated protein
KR134322 (modified) - RNA-binding protein CDS translation - Replication associated protein
KR134323 (modified) - RNA-binding protein CDS translation - Replication associated protein
KR134324 (modified) - RNA-binding protein CDS translation - Replication associated protein
KR134343 (modified) - RNA-binding protein CDS translation - Replication associated protein
KR134344 (modified) - RNA-binding protein CDS translation - Replication associated protein
KR134345 (modified) - RNA-binding protein CDS translation - Replication associated protein
KR134346 (modified) - RNA-binding protein CDS translation - Replication associated protein
KR134347 (modified) - RNA-binding protein CDS translation - Replication associated protein
KR134348 (modified) - RNA-binding protein CDS translation - Replication associated protein
KR134349 (modified) - RNA-binding protein CDS translation - Replication associated protein
KR134350 (modified) - RNA-binding protein CDS translation - Replication associated protein
KR134311 (modified) - Replication associated protein translation - RNA-binding protein CDS
KR134311 (modified) - RNA-binding protein CDS translation - RNA-binding protein CDS
KR134312 (modified) - Replication associated protein translation - RNA-binding protein CDS
KR134312 (modified) - RNA-binding protein CDS translation - RNA-binding protein CDS
KR134314 (modified) - Replication associated protein translation - RNA-binding protein CDS
KR134314 (modified) - RNA-binding protein CDS translation - RNA-binding protein CDS
KR134315 (modified) - Replication associated protein translation - RNA-binding protein CDS
KR134315 (modified) - RNA-binding protein CDS translation - RNA-binding protein CDS
KR134316 (modified) - Replication associated protein translation - RNA-binding protein CDS
KR134316 (modified) - RNA-binding protein CDS translation - RNA-binding protein CDS
KR134317 (modified) - Replication associated protein translation - RNA-binding protein CDS
KR134317 (modified) - RNA-binding protein CDS translation - RNA-binding protein CDS
KR134318 (modified) - Replication associated protein translation - RNA-binding protein CDS
KR134318 (modified) - RNA-binding protein CDS translation - RNA-binding protein CDS
KR134319 (modified) - Replication associated protein translation - RNA-binding protein CDS
KR134319 (modified) - RNA-binding protein CDS translation - RNA-binding protein CDS
KR134320 (modified) - Replication associated protein translation - RNA-binding protein CDS
KR134320 (modified) - RNA-binding protein CDS translation - RNA-binding protein CDS
KR134322 (modified) - Replication associated protein translation - RNA-binding protein CDS
KR134322 (modified) - RNA-binding protein CDS translation - RNA-binding protein CDS
KR134323 (modified) - Replication associated protein translation - RNA-binding protein CDS
KR134323 (modified) - RNA-binding protein CDS translation - RNA-binding protein CDS
KR134324 (modified) - Replication associated protein translation - RNA-binding protein CDS
KR134324 (modified) - RNA-binding protein CDS translation - RNA-binding protein CDS
KR134343 (modified) - Replication associated protein translation - RNA-binding protein CDS
KR134343 (modified) - RNA-binding protein CDS translation - RNA-binding protein CDS
KR134344 (modified) - Replication associated protein translation - RNA-binding protein CDS
KR134344 (modified) - RNA-binding protein CDS translation - RNA-binding protein CDS
KR134345 (modified) - Replication associated protein translation - RNA-binding protein CDS
KR134345 (modified) - RNA-binding protein CDS translation - RNA-binding protein CDS
KR134346 (modified) - Replication associated protein translation - RNA-binding protein CDS
KR134346 (modified) - RNA-binding protein CDS translation - RNA-binding protein CDS
KR134347 (modified) - Replication associated protein translation - RNA-binding protein CDS
KR134347 (modified) - RNA-binding protein CDS translation - RNA-binding protein CDS

KR134348 (modified) - Replication associated protein translation - RNA-binding protein CDS
KR134348 (modified) - RNA-binding protein CDS translation - RNA-binding protein CDS
KR134349 (modified) - Replication associated protein translation - RNA-binding protein CDS
KR134349 (modified) - RNA-binding protein CDS translation - RNA-binding protein CDS
KR134350 (modified) - Replication associated protein translation - RNA-binding protein CDS
KR134350 (modified) - RNA-binding protein CDS translation - RNA-binding protein CDS
KJ955447 (modified) - Replication associated protein translation - viral RNA-binding protein CDS
KJ955447 (modified) - viral RNA-binding protein CDS translation - viral RNA-binding protein CDS
KJ955448 (modified) - Replication associated protein translation - viral RNA-binding protein CDS
KJ955448 (modified) - viral RNA-binding protein CDS translation - viral RNA-binding protein CDS
KJ955449 (modified) - Replication associated protein translation - viral RNA-binding protein CDS
KJ955449 (modified) - viral RNA-binding protein CDS translation - viral RNA-binding protein CDS
KJ955451 (modified) - Replication associated protein translation - viral RNA-binding protein CDS
KJ955451 (modified) - viral RNA-binding protein CDS translation - viral RNA-binding protein CDS
KJ955450 (modified) - viral RNA-binding protein CDS translation - Replication associated protein
KR134325 (modified) - RNA-binding protein CDS translation - Replication associated protein
KR134326 (modified) - RNA-binding protein CDS translation - Replication associated protein
KR134327 (modified) - RNA-binding protein CDS translation - Replication associated protein
KR134328 (modified) - RNA-binding protein CDS translation - Replication associated protein
KR134329 (modified) - RNA-binding protein CDS translation - Replication associated protein
KR134330 (modified) - RNA-binding protein CDS translation - Replication associated protein
KR134331 (modified) - RNA-binding protein CDS translation - Replication associated protein
KR134332 (modified) - RNA-binding protein CDS translation - Replication associated protein
KR134333 (modified) - RNA-binding protein CDS translation - Replication associated protein
KR134334 (modified) - RNA-binding protein CDS translation - Replication associated protein
KR134335 (modified) - RNA-binding protein CDS translation - Replication associated protein
KR134336 (modified) - RNA-binding protein CDS translation - Replication associated protein
KR134337 (modified) - RNA-binding protein CDS translation - Replication associated protein
KR134338 (modified) - RNA-binding protein CDS translation - Replication associated protein
KR134339 (modified) - RNA-binding protein CDS translation - Replication associated protein
KR134340 (modified) - RNA-binding protein CDS translation - Replication associated protein
KR134341 (modified) - RNA-binding protein CDS translation - Replication associated protein
KR134325 (modified) - Replication associated protein translation - RNA-binding protein CDS
KR134325 (modified) - RNA-binding protein CDS translation - RNA-binding protein CDS
KR134326 (modified) - Replication associated protein translation - RNA-binding protein CDS
KR134326 (modified) - RNA-binding protein CDS translation - RNA-binding protein CDS
KR134327 (modified) - Replication associated protein translation - RNA-binding protein CDS
KR134327 (modified) - RNA-binding protein CDS translation - RNA-binding protein CDS
KR134328 (modified) - Replication associated protein translation - RNA-binding protein CDS
KR134328 (modified) - RNA-binding protein CDS translation - RNA-binding protein CDS
KR134329 (modified) - Replication associated protein translation - RNA-binding protein CDS
KR134329 (modified) - RNA-binding protein CDS translation - RNA-binding protein CDS
KR134330 (modified) - Replication associated protein translation - RNA-binding protein CDS
KR134330 (modified) - RNA-binding protein CDS translation - RNA-binding protein CDS
KR134331 (modified) - Replication associated protein translation - RNA-binding protein CDS
KR134331 (modified) - RNA-binding protein CDS translation - RNA-binding protein CDS
KR134332 (modified) - Replication associated protein translation - RNA-binding protein CDS
KR134332 (modified) - RNA-binding protein CDS translation - RNA-binding protein CDS
KR134333 (modified) - Replication associated protein translation - RNA-binding protein CDS
KR134333 (modified) - RNA-binding protein CDS translation - RNA-binding protein CDS
KR134334 (modified) - Replication associated protein translation - RNA-binding protein CDS
KR134334 (modified) - RNA-binding protein CDS translation - RNA-binding protein CDS
KR134335 (modified) - Replication associated protein translation - RNA-binding protein CDS
KR134335 (modified) - RNA-binding protein CDS translation - RNA-binding protein CDS
KR134336 (modified) - Replication associated protein translation - RNA-binding protein CDS
KR134336 (modified) - RNA-binding protein CDS translation - RNA-binding protein CDS
KR134337 (modified) - Replication associated protein translation - RNA-binding protein CDS
KR134337 (modified) - RNA-binding protein CDS translation - RNA-binding protein CDS
KR134338 (modified) - Replication associated protein translation - RNA-binding protein CDS
KR134338 (modified) - RNA-binding protein CDS translation - RNA-binding protein CDS
KR134339 (modified) - Replication associated protein translation - RNA-binding protein CDS
KR134339 (modified) - RNA-binding protein CDS translation - RNA-binding protein CDS
KR134340 (modified) - Replication associated protein translation - RNA-binding protein CDS
KR134340 (modified) - RNA-binding protein CDS translation - RNA-binding protein CDS
KR134341 (modified) - Replication associated protein translation - RNA-binding protein CDS
KR134341 (modified) - RNA-binding protein CDS translation - RNA-binding protein CDS
KJ955450 (modified) - Replication associated protein translation - viral RNA-binding protein CDS
KJ955450 (modified) - viral RNA-binding protein CDS translation - viral RNA-binding protein CDS
KR134321 (modified) - RNA-binding protein CDS translation - Replication associated protein
KR134321 (modified) - Replication associated protein translation - RNA-binding protein CDS
KR134321 (modified) - RNA-binding protein CDS translation - RNA-binding protein CDS
KR134342 (modified) - RNA-binding protein CDS translation - Replication associated protein
KR134342 (modified) - Replication associated protein translation - RNA-binding protein CDS
KR134342 (modified) - RNA-binding protein CDS translation - RNA-binding protein CDS
KU058671.1 - Capsid protein translation - Hypothetical protein
KU058671.1 - Hypothetical protein translation - Hypothetical protein
JQ023166 (modified) - Capsid protein translation - hypothetical protein CDS
JQ023166 (modified) - hypothetical protein CDS translation - hypothetical protein CDS
JX305991 (modified) - Capsid protein translation - hypothetical protein CDS
JX305991 (modified) - hypothetical protein CDS translation - hypothetical protein CDS
JX305992 (modified) - Capsid protein translation - hypothetical protein CDS
JX305992 (modified) - hypothetical protein CDS translation - hypothetical protein CDS
JX305993 (modified) - Capsid protein translation - hypothetical protein CDS
JX305993 (modified) - hypothetical protein CDS translation - hypothetical protein CDS
JX305994 (modified) - Capsid protein translation - hypothetical protein CDS
JX305994 (modified) - hypothetical protein CDS translation - hypothetical protein CDS
JX305995 (modified) - Capsid protein translation - hypothetical protein CDS
JX305995 (modified) - hypothetical protein CDS translation - hypothetical protein CDS
JX305996 (modified) - Capsid protein translation - hypothetical protein CDS
JX305996 (modified) - hypothetical protein CDS translation - hypothetical protein CDS
JX305997 (modified) - Capsid protein translation - hypothetical protein CDS
JX305997 (modified) - hypothetical protein CDS translation - hypothetical protein CDS
JX305998 (modified) - Capsid protein translation - hypothetical protein CDS
JX305998 (modified) - hypothetical protein CDS translation - hypothetical protein CDS
KX388515.1 - Capsid protein translation - hypothetical protein CDS
KX388515.1 - hypothetical protein CDS translation - hypothetical protein CDS
DQ458791 (modified) - Replication-associated protein interval 2 translation - C1 CDS
DQ458791 (modified) - Replication-associated protein interval 2 translation - Replication-associated protein
Y00514 (modified) - Movement protein interval 2 translation - Movement protein
KT214386 (modified) - Replication-associated protein interval 2 translation - RepA CDS
KT214386 (modified) - Replication-associated protein interval 2 translation - Replication-associated protein
KT214373 (modified) - Replication-associated protein interval 2 translation - RepA CDS
KT214373 (modified) - Replication-associated protein interval 2 translation - Replication-associated protein
KF147918 (modified) - Replication-associated protein interval 2 translation - Replication-associated protein
KC248422 (modified) - Replication associated protein translation - hypothetical protein CDS
YP_009051960 - CDS
YP_009051960 - replication-associated protein
YP_009051960 - Viral Rep
YP_009051960 - Viral Rep
YP_009051960 - Viral Rep
KT149402 (modified) - hypothetical protein CDS translation 2 - hypothetical protein CDS
KT149402 (modified) - Replication associated protein translation - hypothetical protein CDS
KT149402 (modified) - hypothetical protein CDS translation 2 - Replication associated protein
KT149399 (modified) - hypothetical protein CDS translation 2 - Replication associated protein
KC108902 (modified) - Replication-associated protein translation - TrAP
GU456685 (modified) - Replication-associated protein translation - TrAP
KT388086 (modified) - Replication-associated protein translation - TrAP
EU921828 (modified) - Replication-associated protein translation - TrAP
JX559642 (modified) - Replication-associated protein interval 2 translation - Replication-associated protein
KC108902 (modified) - TrAP translation - Ren
GU456685 (modified) - TrAP translation - Ren
EU921828 (modified) - TrAP translation - Ren
AF379637 (modified) - TrAP translation - Ren
KT149397 (modified) - Replication associated protein translation - hypothetical protein CDS
KT149405 (modified) - Replication associated protein translation - hypothetical protein CDS
KT388088 (modified) - C4 CDS translation - C4 CDS
KT388088 (modified) - Replication-associated protein translation - C4 CDS
KT388088 (modified) - C4 CDS translation - Replication-associated protein
KR528558 (modified) - Replication associated protein interval 1 translation - contains nonanucleotide motif stem loop
KR528558 (modified) - Replication associated protein interval 1 translation - contains nonanucleotide motif stem loop
KR528558 (modified) - Replication associated protein interval 1 translation - Replication associated protein
KT149400 (modified) - hypothetical protein CDS translation - hypothetical protein CDS
KT149400 (modified) - hypothetical protein CDS translation 3 - hypothetical protein CDS
KT149400 (modified) - hypothetical protein CDS translation 3 - hypothetical protein CDS
AAK73450 - Gemini AL1 M
AAK73450 - Gemini AL1 M
U49907 (modified) - C4 CDS translation - C4 CDS
U49907 (modified) - Replication-associated protein translation - C4 CDS
U49907 (modified) - C4 CDS translation - Replication-associated protein
KF133807 - Replication associated protein translation - hypothetical protein CDS
FJ959079 (modified) - Replication associated protein interval 1 translation - Replication associated protein
FJ959079 (modified) - Replication associated protein interval 1 translation - stem loop
FJ959079 (modified) - Replication associated protein interval 1 translation - stem loop
KR131749 (modified) - Replication-associated protein interval 2 translation - putative RepA-like protein CDS
KR131749 (modified) - Replication-associated protein interval 2 translation - Replication-associated protein
KJ547646 (modified) - hypothetical protein CDS translation - hypothetical protein CDS
KJ547646 (modified) - Replication associated protein translation - hypothetical protein CDS
KJ547646 (modified) - hypothetical protein CDS translation - Replication associated protein
EF536860 (modified) - Replication-associated protein interval 2 translation - RepA
EF536860 (modified) - Replication-associated protein interval 2 translation - Replication-associated protein
KP153395 (modified) - Capsid protein translation - hypothetical protein CDS
KP153396 (modified) - Capsid protein translation - hypothetical protein CDS
KT388088 (modified) - Replication-associated protein translation - TrAP
KT945162 (modified) - Replication associated protein translation - hypothetical protein CDS
GU734126 (modified) - C4 CDS translation - C4 CDS
GU734126 (modified) - Replication-associated protein translation - C4 CDS
GU734126 (modified) - C4 CDS translation - Replication-associated protein
FJ959080 (modified) - Replication associated protein translation - Replication associated protein
FJ959080 (modified) - Replication associated protein translation - stem loop
FJ959080 (modified) - Replication associated protein translation - stem loop
YP_009237571 - CDS
YP_009237571 - replication associated protein
YP_009237571 - Viral Rep
YP_009237571 - Viral Rep
YP_009237571 - Viral Rep
KM821770 (modified) - Replication associated protein interval 2 translation - Replication associated protein
YP_009126879 - Viral Rep
YP_009126879 - Viral Rep
MH545515.1 - Replication associated protein interval 2 translation - Replication associated protein
YP_009142778 - Gemini AL1
YP_009142778 - Gemini AL1
KF133815 - hypothetical protein CDS interval 1 translation - hypothetical protein CDS
KF133815 - hypothetical protein CDS interval 1 translation - stem loop
KF133815 - hypothetical protein CDS interval 1 translation - stem loop
YP_009237504 - Gemini AL1
YP_009237504 - Gemini AL1
AIF34803 - Gemini AL1
AIF34803 - Gemini AL1
YP_009237555 - Gemini AL1
YP_009237555 - Gemini AL1
AXH77952 - Gemini AL1
AXH77952 - Gemini AL1
AXH74707 - Gemini AL1
AXH74707 - Gemini AL1
AAK73450 - Gemini AL1

AAK73450 - Gemini AL1
FJ959079 (modified) - Capsid protein translation - stem
loop
FJ959079 (modified) - Capsid protein translation - stem
loop
AXH78100 - Viral Rep
AXH78100 - Viral Rep
AJD07498 - Viral Rep
AJD07498 - Viral Rep
AXH73056 - P-loop NTPase
AXH73056 - P-loop NTPase
ARE68406 - CDS
ARE68406 - replication associated protein
KY487865.1 - Replication associated protein translation -
Replication associated protein
KR528547 (modified) - Replication associated protein
interval 2 translation - Replication associated protein
KT388086 (modified) - TrAP translation - Ren
KT388088 (modified) - TrAP translation - Ren
YP_009237554 - Gemini AL1 M
KY487886.1 - Replication associated protein translation -
Replication associated protein
JX559642 (modified) - Replication-associated protein
interval 1 translation - 17.9 kDa protein CDS
KF147918 (modified) - Replication-associated protein
interval 1 translation - ORF 5 (frame 3)
KF147918 (modified) - Replication-associated protein
interval 1 translation - ORF 5 (frame 3)
KF147918 (modified) - Replication-associated protein
interval 1 translation - Replication-associated protein
X84735 (modified) - ORF C4 CDS translation - ORF C4
CDS
X84735 (modified) - Replication-associated protein
translation - ORF C4 CDS
X84735 (modified) - ORF C4 CDS translation -
Replication-associated protein
GU456685 (modified) - C4 CDS translation - C4 CDS
GU456685 (modified) - Replication-associated protein
translation - C4 CDS
GU456685 (modified) - C4 CDS translation - Replication-
associated protein
KC108902 (modified) - C4 CDS translation - C4 CDS
KC108902 (modified) - Replication-associated protein
translation - C4 CDS
KC108902 (modified) - C4 CDS translation - Replication-
associated protein
KT388086 (modified) - C4 CDS translation - C4 CDS
KT388086 (modified) - Replication-associated protein
translation - C4 CDS
KT388086 (modified) - C4 CDS translation - Replication-
associated protein
KM386645 (modified) - C4 CDS translation - C4 CDS
KM386645 (modified) - Replication-associated protein
translation - C4 CDS
KM386645 (modified) - C4 CDS translation - Replication-
associated protein
YP_009237554 - Gemini AL1 M
YP_009237554 - Gemini AL1 M
AWR89667 - CDS
AWR89667 - replication initiation protein
AWR89667 - Viral Rep
AWR89667 - Viral Rep
AWR89667 - Viral Rep
KM386645 (modified) - TrAP translation - Ren
X84735 (modified) - TrAP translation - Ren
KT214386 (modified) - V2 CDS translation - V4 CDS
AF379637 (modified) - Replication-associated protein
translation - TrAP
KT945165 (modified) - Replication associated protein
interval 2 translation - Replication associated protein
KT945165 (modified) - Replication associated protein
interval 2 translation - stem loop
KT945165 (modified) - Replication associated protein
interval 2 translation - stem loop
KT945165 (modified) - Replication associated protein
interval 2 translation - Replication associated protein
KT214373 (modified) - V4 CDS translation - V3 CDS
KT214386 (modified) - V4 CDS translation - V3 CDS
KT214389 (modified) - V4 CDS translation - V3 CDS
KT149408 (modified) - hypothetical protein CDS
translation - hypothetical protein CDS
KT149408 (modified) - hypothetical protein CDS
translation 3 - hypothetical protein CDS
KT149408 (modified) - hypothetical protein CDS
translation 3 - hypothetical protein CDS
AMD39533 - Walker A/P-loop
AMD39533 - Walker A/P-loop
AMD39533 - Walker A/P-loop
AMD39533 - Walker A/P-loop
AMD39533 - Walker A/P-loop
AMD39533 - Walker A/P-loop
AMD39533 - Walker A/P-loop
AMD39533 - Walker A/P-loop
AIF76255 - Walker A/P-loop
AIF76255 - Walker A/P-loop
AIF76255 - Walker A/P-loop
YP_009237578 - Walker A motif
YP_009237578 - Walker A motif
YP_009237578 - Walker A motif
WP_027090230 - Walker A motif
WP_027090230 - Walker A motif
WP_027090230 - Walker A motif
WP_027090230 - Walker A motif
WP_027090230 - Walker A motif
KX618694 - HAP/C4 CDS translation - HAP/C4 CDS
KX618694 - Replication associated protein translation -
HAP/C4 CDS
KX618694 - HAP/C4 CDS translation - Replication
associated protein
KT149405 (modified) - hypothetical protein CDS
translation 2 - hypothetical protein CDS
KT149405 (modified) - Replication associated protein
translation - hypothetical protein CDS
KT149405 (modified) - hypothetical protein CDS
translation 2 - Replication associated protein
AGA19549 - Gemini AL1 M
AGA19549 - Gemini AL1 M
JF755410 (modified) - ORF 3 (frame 2) translation - ORF 4
(frame 3)
MH545525.1 - Replication associated protein translation -
Replication associated protein
MH545525.1 - Replication associated protein translation -
stem loop
MH545525.1 - Replication associated protein translation -
stem loop
AKR53286 - P-loop NTPase
AKR53286 - P-loop NTPase
KC248418 (modified) - Replication associated protein
translation - hypothetical protein CDS
U49907 (modified) - V3 CDS translation - V2 CDS
KP410285 (modified) - V3 CDS translation - V2 CDS
EU921828 (modified) - V3 CDS translation - V2 CDS
AF379637 (modified) - V3 translation - V2
HQ443515 (modified) - V3 CDS translation - V2 CDS
GU734126 (modified) - V3 CDS translation - V2 CDS
EU921828 (modified) - C4 CDS translation - C4 CDS
EU921828 (modified) - Replication-associated protein
translation - C4 CDS
EU921828 (modified) - C4 CDS translation - Replication-
associated protein
X84735 (modified) - Replication-associated protein
translation - TrAP
KF147918 (modified) - Capsid protein translation - ORF 3
(frame 1)
KF147918 (modified) - Capsid protein translation - ORF 3
(frame 1)
JX559642 (modified) - Capsid protein translation - 14.7
kDa protein CDS
KX388507.1 - Replication associated protein translation -
stem loop
KT214373 (modified) - Replication-associated protein
interval 1 translation - C3 CDS
KT214373 (modified) - C3 CDS translation - Replication-
associated protein
KT214373 (modified) - C3 CDS translation - C3 CDS
KT214373 (modified) - RepA CDS translation - C3 CDS
KT214373 (modified) - C3 CDS translation - RepA CDS
JX559621 (modified) - hypothetical protein CDS translation
2 - stem loop
JX559622 (modified) - hypothetical protein CDS translation
2 - stem loop
KU043397.1 - Replication associated protein translation -
Replication associated protein
KJ206566 (modified) - Replication associated protein
translation - hypothetical protein CDS
AJD20393 - Gemini AL1
AJD20393 - Gemini AL1
KF133820 - Replication associated protein interval 2
translation - Replication associated protein
KU043420.1 - Hypothetical protein translation -
Hypothetical protein
Y00514 (modified) - RepA translation - Replication-
associated protein interval 2
JF755415 (modified) - Replication associated protein
interval 2 translation - Replication associated protein
MH617545.1 - Replication associated protein translation -
hypothetical protein CDS
MH617545.1 - Replication associated protein translation -
Replication associated protein
JN857329 (modified) - Capsid protein translation -
Replication associated protein
AVH76405 - CDS
AVH76405 - Parvo NS1
AVH76405 - putative Rep Protein
AVH76405 - Parvo NS1
AVH76405 - Parvo NS1
KT732819 (modified) - Capsid protein translation -
Replication associated protein
KF133827 - hypothetical protein CDS translation 2 -
hypothetical protein CDS
KR528567 (modified) - Replication associated protein
translation - Replication associated protein
AGG39817 - P-loop NTPase
AGG39817 - P-loop NTPase
WP_027090230 - P-loop NTPase
WP_027090230 - P-loop NTPase
WP_027090230 - Walker B motif
WP_027090230 - Walker B motif
WP_027090230 - Walker B motif
WP_027090230 - Walker B motif
AQR57902 - RNA helicase
AQR57902 - RNA helicase
AQR57902 - RNA helicase
AXH73290 - P-loop NTPase
AXH73290 - P-loop NTPase
YP_009126892 - CDS
YP_009126892 - replication-associated protein
YP_009126892 - Viral Rep
YP_009126892 - Viral Rep
YP_009126892 - Viral Rep
FJ959079 (modified) - Replication associated protein
interval 2 translation - Replication associated protein
FJ959079 (modified) - Replication associated protein
interval 2 translation - Replication associated protein
FJ959079 (modified) - Replication associated protein
interval 2 translation - stem loop
FJ959079 (modified) - Replication associated protein
interval 2 translation - stem loop
YP_009237541 - P-loop NTPase
YP_009237541 - P-loop NTPase
YP_009237541 - P-loop NTPase
AUF34964 - P-loop NTPase
AUF34964 - P-loop NTPase
KF738881 (modified) - Capsid protein translation -
Replication associated protein
YP_009126892 - RNA helicase
YP_009126892 - RNA helicase
KM821767 (modified) - Replication associated protein
interval 2 translation - Replication associated protein
FJ959083 (modified) - hypothetical protein CDS translation
- Replication associated protein
AF071878 (modified) - Replication associated protein
translation - Replication associated protein
AF071878 (modified) - Replication associated protein
translation - Replication associated protein
KR134311 (modified) - viral protein RNase Z CDS
translation - Replication associated protein
KR134312 (modified) - viral protein RNase Z CDS
translation - Replication associated protein
KR134321 (modified) - viral protein RNase Z CDS
translation - Replication associated protein
KR134322 (modified) - viral protein RNase Z CDS
translation - Replication associated protein
KR134311 (modified) - Replication associated protein
translation - viral protein RNase Z CDS
KR134312 (modified) - Replication associated protein
translation - viral protein RNase Z CDS
KR134321 (modified) - Replication associated protein
translation - viral protein RNase Z CDS
KR134322 (modified) - Replication associated protein
translation - viral protein RNase Z CDS
AQU11729 - P-loop NTPase
AQU11729 - P-loop NTPase
AHH31400 - RNA helicase
AHH31400 - RNA helicase
ARI44308 - RNA helicase
ARI44308 - RNA helicase
AMD39533 - RNA helicase
AMD39533 - RNA helicase
AMD39533 - RNA helicase
AUT13975 - RNA helicase
AUT13975 - RNA helicase
YP_009170674 - RNA helicase
YP_009170674 - RNA helicase
YP_009237578 - RNA helicase
YP_009237578 - RNA helicase
YP_764455 - RNA helicase
YP_764455 - RNA helicase
YP_764455 - RNA helicase
AGA18409 - P-loop NTPase
AGA18409 - P-loop NTPase
ARE67375 - RNA helicase
ARE67375 - RNA helicase
NP_955176 - RNA helicase
NP_955176 - RNA helicase
AEL28813 - RNA helicase
AEL28813 - RNA helicase
AFH02742 - P-loop NTPase
AFH02742 - P-loop NTPase
AQU11733 - P-loop NTPase
AQU11733 - P-loop NTPase
AIF76255 - P-loop NTPase
AIF76255 - P-loop NTPase
AXH77121 - RNA helicase
AXH77121 - RNA helicase
AJP36430 - RNA helicase
AJP36430 - RNA helicase
YP_009126938 - P-loop NTPase
YP_009126938 - P-loop NTPase
AXG50856 - P-loop NTPase
AXG50856 - P-loop NTPase
NP_619761 - RNA helicase
NP_619761 - RNA helicase
AKO71368 - RNA helicase
AKO71368 - RNA helicase
ALA65733 - RNA helicase
ALA65733 - RNA helicase
AHC72271 - RNA helicase
AHC72271 - RNA helicase
YP_008997794 - RNA helicase
YP_008997794 - RNA helicase
AHC72177 - RNA helicase
AHC72177 - RNA helicase
AHC72167 - P-loop NTPase
AHC72167 - P-loop NTPase
CBK25810 - RNA helicase
CBK25810 - RNA helicase
ATY70087 - RNA helicase
ATY70087 - RNA helicase
ADC79191 - RNA helicase
ADC79191 - RNA helicase
ADC79191 - RNA helicase
AGA18391 - P-loop NTPase
AGA18391 - P-loop NTPase
AWR89667 - P-loop NTPase
AWR89667 - P-loop NTPase
AQU11717 - P-loop NTPase
AQU11717 - P-loop NTPase
AQU11724 - P-loop NTPase
AQU11724 - P-loop NTPase
AEL87784 - P-loop NTPase
AEL87784 - P-loop NTPase
KU043415.1 - Replication associated protein translation -
Replication associated protein
YP_009109660 - replication-associated protein
YP_009109660 - start codon not determined CDS
KF133817 - Replication associated protein translation -
Replication associated protein
KX388528.1 - hypothetical protein CDS translation - stem
loop
KF133813 - Replication associated protein interval 2
translation - Replication associated protein
AQR57902 - Rep CDS
AQR57902 - replicase Protein
AQR57902 - TIP49
AQR57902 - TIP49
AQR57902 - TIP49
KR528564 (modified) - Replication associated protein
interval 2 translation - Replication associated protein

KT149404 (modified) - Replication associated protein translation - hypothetical protein CDS
KT149404 (modified) - hypothetical protein CDS translation 3 - Replication associated protein
KF133827 - hypothetical protein CDS translation 2 - hypothetical protein CDS
YP_009237571 - P-loop NTPase
YP_009237571 - P-loop NTPase
AWR89667 - CDS
AWR89667 - replication initiation protein
KF738883 (modified) - Replication associated protein interval 3 translation - Replication associated protein
KR131749 (modified) - ORF4 CDS translation - Replication-associated protein
MH425570.1 - Capsid protein translation - Replication associated protein
KJ955447 (modified) - viral protein RNase Z CDS translation - Replication associated protein
KJ955448 (modified) - viral protein RNase Z CDS translation - Replication associated protein
KJ955449 (modified) - viral protein RNase Z CDS translation - Replication associated protein
KJ955450 (modified) - viral protein RNase Z CDS translation - Replication associated protein
KJ955451 (modified) - viral protein RNase Z CDS translation - Replication associated protein
KR134313 (modified) - viral protein RNase Z CDS translation - Replication associated protein
KR134314 (modified) - viral protein RNase Z CDS translation - Replication associated protein
KR134315 (modified) - viral protein RNase Z CDS translation - Replication associated protein
KR134316 (modified) - viral protein RNase Z CDS translation - Replication associated protein
KR134317 (modified) - viral protein RNase Z CDS translation - Replication associated protein
KR134318 (modified) - viral protein RNase Z CDS translation - Replication associated protein
KR134319 (modified) - viral protein RNase Z CDS translation - Replication associated protein
KR134320 (modified) - viral protein RNase Z CDS translation - Replication associated protein
KR134323 (modified) - viral protein RNase Z CDS translation - Replication associated protein
KR134324 (modified) - viral protein RNase Z CDS translation - Replication associated protein
KR134325 (modified) - viral protein RNase Z CDS translation - Replication associated protein
KR134326 (modified) - viral protein RNase Z CDS translation - Replication associated protein
KR134327 (modified) - viral protein RNase Z CDS translation - Replication associated protein
KR134328 (modified) - viral protein RNase Z CDS translation - Replication associated protein
KR134329 (modified) - viral protein RNase Z CDS translation - Replication associated protein
KR134330 (modified) - viral protein RNase Z CDS translation - Replication associated protein
KR134331 (modified) - viral protein RNase Z CDS translation - Replication associated protein
KR134332 (modified) - viral protein RNase Z CDS translation - Replication associated protein
KR134333 (modified) - viral protein RNase Z CDS translation - Replication associated protein
KR134334 (modified) - viral protein RNase Z CDS translation - Replication associated protein
KR134335 (modified) - viral protein RNase Z CDS translation - Replication associated protein
KR134336 (modified) - viral protein RNase Z CDS translation - Replication associated protein
KR134337 (modified) - viral protein RNase Z CDS translation - Replication associated protein
KR134338 (modified) - viral protein RNase Z CDS translation - Replication associated protein
KR134339 (modified) - viral protein RNase Z CDS translation - Replication associated protein
KR134340 (modified) - viral protein RNase Z CDS translation - Replication associated protein
KR134341 (modified) - viral protein RNase Z CDS translation - Replication associated protein
KR134342 (modified) - viral protein RNase Z CDS translation - Replication associated protein
KR134343 (modified) - viral protein RNase Z CDS translation - Replication associated protein
KR134344 (modified) - viral protein RNase Z CDS translation - Replication associated protein
KR134345 (modified) - viral protein RNase Z CDS translation - Replication associated protein
KR134346 (modified) - viral protein RNase Z CDS translation - Replication associated protein
KR134347 (modified) - viral protein RNase Z CDS translation - Replication associated protein
KR134348 (modified) - viral protein RNase Z CDS translation - Replication associated protein
KR134349 (modified) - viral protein RNase Z CDS translation - Replication associated protein
KR134350 (modified) - viral protein RNase Z CDS translation - Replication associated protein
YP_003084282 - RNA helicase
YP_003084282 - RNA helicase
KM598394 (modified) - Replication associated protein interval 2 translation - Replication associated protein
AOV86234 - DNA pol3 delta2
AOV86234 - DNA pol3 delta2
KJ547620 (modified) - Replication associated protein interval 2 translation - Replication associated protein
KP153434 (modified) - Capsid protein translation - Hypothetical protein
KP153436 (modified) - Capsid protein translation - Hypothetical protein
KP153435 (modified) - Capsid protein translation - Hypothetical protein
KT149408 (modified) - hypothetical protein CDS translation - hypothetical protein CDS
KT149408 (modified) - hypothetical protein CDS translation 4 - hypothetical protein CDS
KT149408 (modified) - hypothetical protein CDS translation 4 - hypothetical protein CDS
AIW81537 - RNA helicase
AIW81537 - RNA helicase
AQU11726 - P-loop NTPase
AQU11726 - P-loop NTPase
AQU11726 - P-loop NTPase
KM598412 (modified) - Capsid protein translation - Replication associated protein
KF133823 - hypothetical protein CDS translation - hypothetical protein CDS
KR131749 (modified) - Replication-associated protein interval 2 translation - ORF4 CDS
AGA18265 - P-loop NTPase
AGA18265 - P-loop NTPase
AJD07498 - RNA helicase
AJD07498 - RNA helicase
AXH73792 - P-loop NTPase
AXH73792 - P-loop NTPase
YP_009237564 - P-loop NTPase
YP_009237564 - P-loop NTPase
ALE29688 - RNA helicase
ALE29688 - RNA helicase
YP_009001742 - P-loop NTPase
YP_009001742 - P-loop NTPase
YP_009448204 - RNA helicase
YP_009448204 - RNA helicase
AGA18286 - P-loop NTPase
AGA18286 - P-loop NTPase
KP410285 (modified) - Replication-associated protein interval 2 translation - Replication-associated protein
KM510190 (modified) - Capsid protein translation - hypothetical protein CDS
KM510189 (modified) - Capsid protein translation - hypothetical protein CDS
KF133819 - hypothetical protein CDS translation - hypothetical protein CDS
KF133827 - Replication associated protein translation - Replication associated protein
KF133827 - Replication associated protein translation - stem loop
KF133827 - Replication associated protein translation - stem loop
KT945162 (modified) - Replication associated protein translation - Replication associated protein
KT945162 (modified) - Replication associated protein translation - stem loop
KT945162 (modified) - Replication associated protein translation - stem loop
KR528568 (modified) - Replication associated protein translation - Replication associated protein
KM874301 (modified) - Replication associated protein translation - Replication associated protein
KM874302 (modified) - Replication associated protein translation - Replication associated protein
KM874303 (modified) - Replication associated protein translation - Replication associated protein
MH545528.1 - Replication associated protein translation - Replication associated protein
MH545527.1 - Replication associated protein translation - Replication associated protein
KX618694 - Replication associated protein translation - Replication associated protein
KP153527 (modified) - Replication associated protein translation - Replication associated protein
KF738880 (modified) - Replication associated protein interval 1 translation - Replication associated protein
KP153434 (modified) - Replication associated protein translation - Replication associated protein
KP153435 (modified) - Replication associated protein translation - Replication associated protein
KP153436 (modified) - Replication associated protein translation - Replication associated protein
KF738878 (modified) - Replication associated protein interval 1 translation - Replication associated protein
KF738879 (modified) - Replication associated protein interval 1 translation - Replication associated protein
KP153524 (modified) - Replication associated protein translation - Replication associated protein
KP153525 (modified) - Replication associated protein translation - Replication associated protein
KM874341 (modified) - Replication associated protein translation - Replication associated protein
KM874340 (modified) - Replication associated protein translation - Replication associated protein
KM874342 (modified) - Replication associated protein translation - Replication associated protein
KT149402 (modified) - Replication associated protein translation - Replication associated protein
KM598407 (modified) - Replication associated protein translation - Replication associated protein
KT149396 (modified) - hypothetical protein CDS translation - Replication associated protein
JX904231 (modified) - Replication associated protein translation - Replication associated protein
KT149410 (modified) - hypothetical protein CDS translation - hypothetical protein CDS
MH545530.1 - Replication associated protein interval 1 translation - Replication associated protein
KF133827 - hypothetical protein CDS translation - hypothetical protein CDS
KF133812 - hypothetical protein CDS translation - hypothetical protein CDS
KC248420 (modified) - Replication associated protein translation - Replication associated protein
KR528554 (modified) - Replication associated protein translation - Replication associated protein
KX388511.1 - Replication associated protein translation - Replication associated protein
KR131749 (modified) - Capsid protein translation - putative movement protein CDS
KP153422 (modified) - Replication associated protein translation - Replication associated protein
KP153505 (modified) - Replication associated protein translation - Replication associated protein
KP153506 (modified) - Replication associated protein translation - Replication associated protein
KP153502 (modified) - Capsid protein translation - Replication associated protein
KP153503 (modified) - Capsid protein translation - Replication associated protein
AJ304456 (modified) - Replication associated protein translation - Replication associated protein
AJ304456 (modified) - Replication associated protein translation - Replication associated protein
KT214389 (modified) - V3 CDS translation - V4 CDS
KT214389 (modified) - V3 CDS translation - V3 CDS
DQ100076 (modified) - Replication associated protein translation - Replication associated protein
DQ100076 (modified) - Replication associated protein translation - Replication associated protein
KY884302.1 - Capsid protein translation - Replication associated protein
KP406530 (modified) - Replication associated protein translation - Replication associated protein
KP153516 (modified) - Replication associated protein translation - Replication associated protein
JF755408 (modified) - ORF 3 (frame 3) translation - ORF 3 (frame 3)
KT149412 (modified) - hypothetical protein CDS translation - hypothetical protein CDS
KF738881 (modified) - Replication associated protein translation - Replication associated protein
KJ547623 (modified) - Replication associated protein translation - Replication associated protein
KF133816 - hypothetical protein CDS translation - hypothetical protein CDS
AF379637 (modified) - L4 CDS translation - L4 CDS
MH545537.1 - Replication associated protein translation - Replication associated protein
KC248418 (modified) - hypothetical protein CDS translation - Replication associated protein
KC248418 (modified) - hypothetical protein CDS translation - hypothetical protein CDS
KP153494 (modified) - Replication associated protein translation - Replication associated protein
KP153495 (modified) - Replication associated protein translation - Replication associated protein
KU043423.1 - Replication associated protein translation - Replication associated protein
KX388502.1 - Replication associated protein translation - Replication associated protein
KP153526 (modified) - Replication associated protein translation - Replication associated protein
KF738882 (modified) - Replication associated protein translation - Replication associated protein
KP153424 (modified) - Replication associated protein translation - Replication associated protein
KP153425 (modified) - Replication associated protein translation - Replication associated protein
KP153426 (modified) - Replication associated protein translation - Replication associated protein
KF738876 (modified) - Replication associated protein translation - Replication associated protein
KP153449 (modified) - Replication associated protein translation - Replication associated protein
KP153447 (modified) - Replication associated protein translation - Replication associated protein
KP153448 (modified) - Replication associated protein translation - Replication associated protein
KP153450 (modified) - Replication associated protein translation - Replication associated protein
KF738875 (modified) - Replication associated protein translation - Replication associated protein
KJ547633 (modified) - Replication associated protein translation - Replication associated protein
KU043448.1 - Replication associated protein translation - Replication associated protein
KP153420 (modified) - Replication associated protein translation - Replication associated protein
KP153421 (modified) - Replication associated protein translation - Replication associated protein
AF252610 (modified) - Replication associated protein translation - Replication associated protein
AF252610 (modified) - Replication associated protein translation - Replication associated protein
KJ955448 (modified) - Replication associated protein translation - Replication associated protein
KR134314 (modified) - Replication associated protein translation - Replication associated protein
KJ955450 (modified) - Replication associated protein translation - Replication associated protein
KR134325 (modified) - Replication associated protein translation - Replication associated protein
KR134326 (modified) - Replication associated protein translation - Replication associated protein
KR134327 (modified) - Replication associated protein translation - Replication associated protein
KR134328 (modified) - Replication associated protein translation - Replication associated protein
KR134329 (modified) - Replication associated protein translation - Replication associated protein
KR134330 (modified) - Replication associated protein translation - Replication associated protein
KR134331 (modified) - Replication associated protein translation - Replication associated protein
KR134332 (modified) - Replication associated protein translation - Replication associated protein
KR134333 (modified) - Replication associated protein translation - Replication associated protein
KR134334 (modified) - Replication associated protein translation - Replication associated protein

KR134335 (modified) - Replication associated protein translation - Replication associated protein
KR134336 (modified) - Replication associated protein translation - Replication associated protein
KR134337 (modified) - Replication associated protein translation - Replication associated protein
KR134338 (modified) - Replication associated protein translation - Replication associated protein
KR134339 (modified) - Replication associated protein translation - Replication associated protein
KR134340 (modified) - Replication associated protein translation - Replication associated protein
KR134341 (modified) - Replication associated protein translation - Replication associated protein
KR134342 (modified) - Replication associated protein translation - Replication associated protein
KJ955447 (modified) - Replication associated protein translation - Replication associated protein
KJ955449 (modified) - Replication associated protein translation - Replication associated protein
KJ955451 (modified) - Replication associated protein translation - Replication associated protein
KR134313 (modified) - Replication associated protein translation - Replication associated protein
KR134315 (modified) - Replication associated protein translation - Replication associated protein
KR134316 (modified) - Replication associated protein translation - Replication associated protein
KR134317 (modified) - Replication associated protein translation - Replication associated protein
KR134318 (modified) - Replication associated protein translation - Replication associated protein
KR134319 (modified) - Replication associated protein translation - Replication associated protein
KR134320 (modified) - Replication associated protein translation - Replication associated protein
KR134323 (modified) - Replication associated protein translation - Replication associated protein
KR134324 (modified) - Replication associated protein translation - Replication associated protein
KR134343 (modified) - Replication associated protein translation - Replication associated protein
KR134344 (modified) - Replication associated protein translation - Replication associated protein
KR134345 (modified) - Replication associated protein translation - Replication associated protein
KR134346 (modified) - Replication associated protein translation - Replication associated protein
KR134347 (modified) - Replication associated protein translation - Replication associated protein
KR134348 (modified) - Replication associated protein translation - Replication associated protein
KR134349 (modified) - Replication associated protein translation - Replication associated protein
KR134350 (modified) - Replication associated protein translation - Replication associated protein
KR134311 (modified) - Replication associated protein translation - Replication associated protein
KR134312 (modified) - Replication associated protein translation - Replication associated protein
KR134322 (modified) - Replication associated protein translation - Replication associated protein
KR134321 (modified) - Replication associated protein translation - Replication associated protein
KT388088 (modified) - Replication-associated protein translation - Replication-associated protein
KP153409 (modified) - Replication associated protein translation - Replication associated protein
KR528553 (modified) - Replication associated protein translation - Replication associated protein
KR528549 (modified) - Replication associated protein translation - Replication associated protein
KM598406 (modified) - Replication associated protein translation - Replication associated protein
KM510189 (modified) - Replication associated protein interval 1 translation - Replication associated protein
KM510190 (modified) - Replication associated protein interval 1 translation - Replication associated protein
MH545516.1 - Replication associated protein interval 1 translation - Replication associated protein
KP153502 (modified) - Replication associated protein translation - Replication associated protein
KC172652 (modified) - Replication associated protein translation - Replication associated protein
KM874354 (modified) - Replication associated protein translation - Replication associated protein
KX388528.1 - Replication associated protein translation - Replication associated protein
KX388529.1 - Replication associated protein translation - Replication associated protein
KX388530.1 - Replication associated protein translation - Replication associated protein
MH545531.1 - Replication associated protein interval 1 translation - Replication associated protein
KM874355 (modified) - Replication associated protein translation - Replication associated protein
KM874356 (modified) - Replication associated protein translation - Replication associated protein
KM874357 (modified) - Replication associated protein translation - Replication associated protein
KM874358 (modified) - Replication associated protein translation - Replication associated protein
KJ547628 (modified) - Replication associated protein translation - Replication associated protein
KP153470 (modified) - Replication associated protein translation - Replication associated protein
KP153471 (modified) - Replication associated protein translation - Replication associated protein
MF118166.1 - hypothetical protein CDS translation 2 - hypothetical protein CDS
MF118167.1 - hypothetical protein CDS translation 2 - hypothetical protein CDS
KC543331 (modified) - Capsid protein translation - hypothetical protein CDS
KC794539 (modified) - Capsid protein translation - hypothetical protein CDS
KC794540 (modified) - Capsid protein translation - hypothetical protein CDS
KP153487 (modified) - Replication associated protein translation - Replication associated protein
MF118169.1 - hypothetical protein CDS translation - hypothetical protein CDS
KP153376 (modified) - Replication associated protein translation - Replication associated protein
KP153375 (modified) - Replication associated protein translation - Replication associated protein
KP153374 (modified) - Replication associated protein translation - Replication associated protein
KF738874 (modified) - Replication associated protein translation - Replication associated protein
KJ547631 (modified) - Replication associated protein translation - hypothetical protein CDS
KJ547631 (modified) - Replication associated protein translation - Replication associated protein
KC248417 (modified) - Replication associated protein translation - Replication associated protein
KM821755 (modified) - Replication associated protein translation - Replication associated protein
KM874362 (modified) - Replication associated protein translation - Replication associated protein
KM874363 (modified) - Replication associated protein translation - Replication associated protein
KM874364 (modified) - Replication associated protein translation - Replication associated protein
KM874365 (modified) - Replication associated protein translation - Replication associated protein
KM821748 (modified) - Replication associated protein interval 1 translation - Replication associated protein
HQ443515 (modified) - Replication-associated protein interval 1 translation - Replication-associated protein
JX094280 (modified) - Capsid protein translation - ORF 2 (frame 2)
JX094280 (modified) - Capsid protein translation - ORF 2 (frame 2)
JQ898333 (modified) - Capsid protein translation - hypothetical protein CDS
JF755411 (modified) - ORF 6 (frame 2) translation - ORF 6 (frame 2)
JF755412 (modified) - ORF 6 (frame 2) translation - ORF 6 (frame 2)
JF755413 (modified) - ORF 6 (frame 2) translation - ORF 6 (frame 2)
JF755414 (modified) - ORF 6 (frame 2) translation - ORF 6 (frame 2)
JF755402 (modified) - ORF 9 (frame 1) translation - ORF 9 (frame 1)
JX559642 (modified) - 17.9 kDa protein CDS translation - Replication-associated protein
JX559642 (modified) - 17.9 kDa protein CDS translation - 17.9 kDa protein CDS
KR131749 (modified) - Replication-associated protein interval 1 translation - putative RepA-like protein CDS
KR131749 (modified) - putative RepA-like protein CDS translation - Replication-associated protein interval 1
KR131749 (modified) - Replication-associated protein interval 1 translation - Replication-associated protein
KR131749 (modified) - putative RepA-like protein CDS translation - putative RepA-like protein CDS
KT732831 (modified) - Replication associated protein translation - Replication associated protein
KT732829 (modified) - Replication associated protein translation - Replication associated protein
FJ959085 (modified) - Replication associated protein translation - Replication associated protein
KP153452 (modified) - hypothetical protein CDS translation - hypothetical protein CDS
KP153453 (modified) - hypothetical protein CDS translation - hypothetical protein CDS
KP153451 (modified) - hypothetical protein CDS translation - hypothetical protein CDS
KU043413.1 - Hypothetical protein translation 2 - Hypothetical protein
KT149400 (modified) - hypothetical protein CDS translation 2 - Replication associated protein
KT149400 (modified) - hypothetical protein CDS translation 2 - hypothetical protein CDS
FJ665630 (modified) - Replication-associated protein translation - Replication-associated protein
FJ665634 (modified) - Replication-associated protein translation - Replication-associated protein
JX559642 (modified) - Replication-associated protein interval 1 translation - Replication-associated protein
KF147918 (modified) - Replication-associated protein interval 1 translation - Replication-associated protein
Y00514 (modified) - Replication-associated protein interval 1 translation - RepA
Y00514 (modified) - RepA translation - Replication-associated protein interval 1
Y00514 (modified) - Replication-associated protein interval 1 translation - Replication-associated protein
Y00514 (modified) - RepA translation - RepA
EF536860 (modified) - Replication-associated protein interval 1 translation - RepA
EF536860 (modified) - RepA translation - Replication-associated protein interval 1
EF536860 (modified) - Replication-associated protein interval 1 translation - Replication-associated protein
EF536860 (modified) - RepA translation - RepA
MH545523.1 - Replication associated protein translation - Replication associated protein
JQ920490 (modified) - Replication-associated protein interval 1 translation - Replication-associated protein
KP153405 (modified) - Replication associated protein translation - Replication associated protein
KP153406 (modified) - Replication associated protein translation - Replication associated protein
KT149394 (modified) - Replication associated protein translation - Replication associated protein
JQ085285 - Replication associated protein translation - Replication associated protein
KT945162 (modified) - Replication associated protein translation - Replication associated protein
KC248418 (modified) - Replication associated protein translation - Replication associated protein
KY884302.1 - Replication associated protein translation - Replication associated protein
KP153442 (modified) - Replication associated protein translation - Replication associated protein
X84735 (modified) - Movement protein translation - Movement protein
KJ547622 (modified) - Replication associated protein translation - Replication associated protein
KT149406 (modified) - hypothetical protein CDS translation - hypothetical protein CDS
KP410285 (modified) - Capsid protein translation - V2 CDS
KT388086 (modified) - Ren translation - TrAP
KT388086 (modified) - Ren translation - Ren
KT388088 (modified) - Ren translation - TrAP
KT388088 (modified) - Ren translation - Ren
MH545518.1 - Capsid protein translation - Replication associated protein
MH545519.1 - Capsid protein translation - Replication associated protein
JF755403 (modified) - ORF 4 (frame 3) translation - ORF 4 (frame 3)
KY487896.1 - Replication associated protein translation - Replication associated protein
KT149409 (modified) - hypothetical protein CDS translation - hypothetical protein CDS
KP153410 (modified) - Replication associated protein translation - Replication associated protein
KP153411 (modified) - Replication associated protein translation - Replication associated protein
KF133818 - hypothetical protein CDS translation - hypothetical protein CDS
KF133818 - hypothetical protein CDS translation - hypothetical protein CDS
KF133816 - hypothetical protein CDS translation 2 - hypothetical protein CDS
MF118167.1 - hypothetical protein CDS translation 4 - hypothetical protein CDS
MF118166.1 - hypothetical protein CDS translation 4 - hypothetical protein CDS
FJ959086 (modified) - hypothetical protein CDS translation - hypothetical protein CDS
JQ920490 (modified) - Movement protein translation - Movement protein
FJ959083 (modified) - hypothetical protein CDS translation - hypothetical protein CDS
MH545535.1 - Replication associated protein translation - Replication associated protein
KT732784 (modified) - hypothetical protein CDS translation - hypothetical protein CDS
KX388505.1 - Replication associated protein translation - Replication associated protein
MH545524.1 - Replication associated protein translation - Replication associated protein
HQ322117 (modified) - Replication associated protein translation - Replication associated protein
KX388497.1 - hypothetical protein CDS translation 2 - hypothetical protein CDS
KX618694 - Capsid protein translation - Pre-CP/V2 CDS
JQ920490 (modified) - putative BL1-like movement protein CDS translation - putative BL1-like movement protein CDS
KJ547651 (modified) - hypothetical protein CDS translation - hypothetical protein CDS
KX388497.1 - hypothetical protein CDS translation - hypothetical protein CDS
KJ955447 (modified) - truncated viral movement major protein CDS translation - truncated viral movement major protein CDS
KJ955448 (modified) - truncated viral movement major protein CDS translation - truncated viral movement major protein CDS
KJ955449 (modified) - truncated viral movement major protein CDS translation - truncated viral movement major protein CDS
KJ955450 (modified) - truncated viral movement major protein CDS translation - truncated viral movement major protein CDS
KJ955451 (modified) - truncated viral movement major protein CDS translation - truncated viral movement major protein CDS
KR134311 (modified) - truncated viral movement protein CDS translation - truncated viral movement protein CDS
KR134312 (modified) - truncated viral movement protein CDS translation - truncated viral movement protein CDS
KR134313 (modified) - truncated viral movement protein CDS translation - truncated viral movement protein CDS
KR134314 (modified) - truncated viral movement protein CDS translation - truncated viral movement protein CDS
KR134315 (modified) - truncated viral movement protein CDS translation - truncated viral movement protein CDS
KR134316 (modified) - truncated viral movement protein CDS translation - truncated viral movement protein CDS
KR134317 (modified) - truncated viral movement protein CDS translation - truncated viral movement protein CDS
KR134318 (modified) - truncated viral movement protein CDS translation - truncated viral movement protein CDS
KR134319 (modified) - truncated viral movement protein CDS translation - truncated viral movement protein CDS
KR134320 (modified) - truncated viral movement protein CDS translation - truncated viral movement protein CDS
KR134321 (modified) - truncated viral movement protein CDS translation - truncated viral movement protein CDS

KR134322 (modified) - truncated viral movement protein CDS translation - truncated viral movement protein CDS
KR134323 (modified) - truncated viral movement protein CDS translation - truncated viral movement protein CDS
KR134324 (modified) - truncated viral movement protein CDS translation - truncated viral movement protein CDS
KR134325 (modified) - truncated viral movement protein CDS translation - truncated viral movement protein CDS
KR134326 (modified) - truncated viral movement protein CDS translation - truncated viral movement protein CDS
KR134327 (modified) - truncated viral movement protein CDS translation - truncated viral movement protein CDS
KR134328 (modified) - truncated viral movement protein CDS translation - truncated viral movement protein CDS
KR134329 (modified) - truncated viral movement protein CDS translation - truncated viral movement protein CDS
KR134330 (modified) - truncated viral movement protein CDS translation - truncated viral movement protein CDS
KR134331 (modified) - truncated viral movement protein CDS translation - truncated viral movement protein CDS
KR134332 (modified) - truncated viral movement protein CDS translation - truncated viral movement protein CDS
KR134333 (modified) - truncated viral movement protein CDS translation - truncated viral movement protein CDS
KR134334 (modified) - truncated viral movement protein CDS translation - truncated viral movement protein CDS
KR134335 (modified) - truncated viral movement protein CDS translation - truncated viral movement protein CDS
KR134336 (modified) - truncated viral movement protein CDS translation - truncated viral movement protein CDS
KR134337 (modified) - truncated viral movement protein CDS translation - truncated viral movement protein CDS
KR134338 (modified) - truncated viral movement protein CDS translation - truncated viral movement protein CDS
KR134339 (modified) - truncated viral movement protein CDS translation - truncated viral movement protein CDS
KR134340 (modified) - truncated viral movement protein CDS translation - truncated viral movement protein CDS
KR134341 (modified) - truncated viral movement protein CDS translation - truncated viral movement protein CDS
KR134342 (modified) - truncated viral movement protein CDS translation - truncated viral movement protein CDS
KR134343 (modified) - truncated viral movement protein CDS translation - truncated viral movement protein CDS
KR134344 (modified) - truncated viral movement protein CDS translation - truncated viral movement protein CDS
KR134345 (modified) - truncated viral movement protein CDS translation - truncated viral movement protein CDS
KR134347 (modified) - truncated viral movement protein CDS translation - truncated viral movement protein CDS
KR134348 (modified) - truncated viral movement protein CDS translation - truncated viral movement protein CDS
KR134349 (modified) - truncated viral movement protein CDS translation - truncated viral movement protein CDS
KR134350 (modified) - truncated viral movement protein CDS translation - truncated viral movement protein CDS
KR134346 (modified) - truncated viral movement protein CDS translation - truncated viral movement protein CDS
KX388499.1 - hypothetical protein CDS translation - hypothetical protein CDS
KJ547624 (modified) - Replication associated protein translation - Replication associated protein
KF133817 - hypothetical protein CDS translation 2 - hypothetical protein CDS
KP153418 (modified) - Replication associated protein translation - Replication associated protein
KP153419 (modified) - Replication associated protein translation - Replication associated protein
KP153417 (modified) - Replication associated protein translation - Replication associated protein
KP153523 (modified) - Replication associated protein translation - Replication associated protein
KR528559 (modified) - Replication associated protein translation - Replication associated protein
KX388509.1 - hypothetical protein CDS translation - hypothetical protein CDS
KM386645 (modified) - Capsid protein translation - Movement protein
X84735 (modified) - Ren translation - TrAP
X84735 (modified) - Ren translation - Ren
KM386645 (modified) - Ren translation - TrAP
KM386645 (modified) - Ren translation - Ren
KF133818 - hypothetical protein CDS translation 4 - hypothetical protein CDS
KT732834 (modified) - Replication associated protein translation - Replication associated protein
KM821753 (modified) - Replication associated protein translation - Replication associated protein
KT732832 (modified) - hypothetical protein CDS translation - hypothetical protein CDS
FJ959078 (modified) - Replication associated protein translation - Replication associated protein
KP153437 (modified) - Replication associated protein translation - Replication associated protein
KP153440 (modified) - Replication associated protein translation - Replication associated protein
KP153439 (modified) - Replication associated protein translation - Replication associated protein
KP153438 (modified) - Replication associated protein translation - Replication associated protein
KX388494.1 - Replication associated protein translation - Replication associated protein
KX388494.1 - Replication associated protein translation - stem loop
KX388494.1 - Replication associated protein translation - stem loop
KX388494.1 - Replication associated protein translation - Replication associated protein
KM821763 (modified) - Replication associated protein translation - Replication associated protein
KM386645 (modified) - Movement protein translation - Movement protein
KT149401 (modified) - hypothetical protein CDS translation 2 - hypothetical protein CDS
KJ547629 (modified) - Replication associated protein translation - Replication associated protein
KM821770 (modified) - Replication associated protein interval 1 translation - Replication associated protein
KC248416 (modified) - hypothetical protein CDS translation - hypothetical protein CDS
KP153489 (modified) - Replication associated protein translation - Replication associated protein
KP153488 (modified) - Replication associated protein translation - Replication associated protein
KT862236 (modified) - Replication associated protein translation - Replication associated protein
KR528543 (modified) - Replication associated protein translation - Replication associated protein
KT862234 (modified) - Replication associated protein translation - Replication associated protein
KP153500 (modified) - Hypothetical 2 translation - Hypothetical 2
KP153500 (modified) - Replication associated protein translation - Replication associated protein
KT214386 (modified) - V3 CDS translation - V2 CDS
KT214386 (modified) - V3 CDS translation - V4 CDS
KT214386 (modified) - V3 CDS translation - V3 CDS
JX559642 (modified) - 14.7 kDa protein CDS translation - 14.7 kDa protein CDS
AF379637 (modified) - TrAP translation - Replication-associated protein
AF379637 (modified) - TrAP translation - TrAP
KR528546 (modified) - Replication associated protein translation - Replication associated protein
KP153423 (modified) - Replication associated protein translation - Replication associated protein
KP153403 (modified) - Replication associated protein translation - Replication associated protein
KJ547651 (modified) - Replication associated protein translation - Replication associated protein
KT149400 (modified) - Replication associated protein translation - Replication associated protein
EF536860 (modified) - Movement protein translation - Movement protein
DQ458791 (modified) - Movement protein translation - Movement protein
KC248424 (modified) - Replication associated protein translation - Replication associated protein
KM821762 (modified) - Replication associated protein translation - Replication associated protein
KP153477 (modified) - Replication associated protein interval 1 translation - Replication associated protein
KP153478 (modified) - Replication associated protein interval 1 translation - Replication associated protein
KP153479 (modified) - Replication associated protein interval 1 translation - Replication associated protein
KP153480 (modified) - Replication associated protein interval 1 translation - Replication associated protein
KP153481 (modified) - Replication associated protein interval 1 translation - Replication associated protein
KP153482 (modified) - Replication associated protein interval 1 translation - Replication associated protein
KP153476 (modified) - Replication associated protein interval 1 translation - Replication associated protein
KP153459 (modified) - Hypothetical 1 translation - Hypothetical 1
KP153461 (modified) - Hypothetical 1 translation - Hypothetical 1
KP153462 (modified) - Hypothetical 1 translation - Hypothetical 1
KP153460 (modified) - Hypothetical 1 translation - Hypothetical 1
KP153463 (modified) - Hypothetical 1 translation - Hypothetical 1
KT214373 (modified) - V3 CDS translation - V4 CDS
KT214373 (modified) - V3 CDS translation - V3 CDS
KJ547625 (modified) - Replication associated protein translation - Replication associated protein
KT149406 (modified) - hypothetical protein CDS translation 2 - hypothetical protein CDS
FJ959077 (modified) - hypothetical protein CDS translation 2 - hypothetical protein CDS
KT388088 (modified) - Movement protein translation - Movement protein
KT214373 (modified) - V4 CDS translation - V4 CDS
KT214386 (modified) - V4 CDS translation - V2 CDS
KT214386 (modified) - V4 CDS translation - V4 CDS
KT214389 (modified) - V4 CDS translation - V4 CDS
KX388501.1 - Replication associated protein translation - Replication associated protein
KX388498.1 - Replication associated protein translation - Replication associated protein
KX388500.1 - Replication associated protein translation - Replication associated protein
KF133817 - hypothetical protein CDS translation - hypothetical protein CDS
KT732784 (modified) - Replication associated protein translation - Replication associated protein
KP153404 (modified) - Replication associated protein translation - Replication associated protein
JF713717 (modified) - unknown CDS translation - unknown CDS
KR528566 (modified) - Replication associated protein translation - Replication associated protein
JF713718 (modified) - unknown CDS translation 2 - unknown CDS
KX388513.1 - hypothetical protein CDS translation - hypothetical protein CDS
KT149400 (modified) - hypothetical protein CDS translation 4 - hypothetical protein CDS
KU043406.1 - Replication associated protein translation - Replication associated protein
KM598390 (modified) - hypothetical protein CDS translation - hypothetical protein CDS
KM821764 (modified) - hypothetical protein CDS translation - hypothetical protein CDS
KT149395 (modified) - Replication associated protein translation - Replication associated protein
KT149396 (modified) - hypothetical protein CDS translation - hypothetical protein CDS
KC248425 (modified) - hypothetical protein CDS translation - hypothetical protein CDS
KT214386 (modified) - V2 CDS translation - V2 CDS
KR131749 (modified) - ORF4 CDS translation - ORF4 CDS
KT732824 (modified) - Replication associated protein translation - Replication associated protein
KM573768 (modified) - ORF2 CDS translation - ORF2 CDS
KM821750 (modified) - Replication associated protein translation - Replication associated protein
KM821760 (modified) - hypothetical protein CDS translation - hypothetical protein CDS
KJ547630 (modified) - Replication associated protein translation - Replication associated protein
KF133822 - hypothetical protein CDS translation - hypothetical protein CDS
KP153397 (modified) - Capsid protein translation - Replication associated protein
KP153398 (modified) - Capsid protein translation - Replication associated protein
KP153399 (modified) - Capsid protein translation - Replication associated protein
KP153400 (modified) - Capsid protein translation - Replication associated protein
KP153401 (modified) - Capsid protein translation - Replication associated protein
KM598394 (modified) - Replication associated protein interval 1 translation - Replication associated protein
KF738883 (modified) - Replication associated protein interval 1 translation - Replication associated protein
KF133814 - hypothetical protein CDS translation 2 - hypothetical protein CDS
KF133815 - ORF 5 (frame 2) translation - hypothetical protein CDS
KF133815 - ORF 5 (frame 2) translation - ORF 5 (frame 2)
KU043430.1 - Replication associated protein translation - Replication associated protein
FJ959077 (modified) - hypothetical protein CDS translation - hypothetical protein CDS
FJ959077 (modified) - hypothetical protein CDS translation - hypothetical protein CDS
KF738877 (modified) - Replication associated protein interval 1 translation - Replication associated protein
KM573767 (modified) - ORF3 CDS translation - ORF3 CDS
JX559642 (modified) - Movement protein translation - Movement protein
GU734126 (modified) - V2 CDS translation - V3 CDS
GU734126 (modified) - V2 CDS translation - V2 CDS
EU921828 (modified) - V2 CDS translation - V3 CDS
EU921828 (modified) - V2 CDS translation - V2 CDS
AF379637 (modified) - V2 translation - V3
AF379637 (modified) - V2 translation - V2
HQ443515 (modified) - V2 CDS translation - V3 CDS
HQ443515 (modified) - V2 CDS translation - V2 CDS
U49907 (modified) - V2 CDS translation - V3 CDS
U49907 (modified) - V2 CDS translation - V2 CDS
KF133819 - hypothetical protein CDS translation 2 - hypothetical protein CDS
JN857329 (modified) - Replication associated protein translation - Replication associated protein
KJ547653 (modified) - hypothetical protein CDS translation - hypothetical protein CDS
JQ085285 - Capsid protein translation - Replication associated protein
KT388086 (modified) - Replication-associated protein translation - Replication-associated protein
KF147918 (modified) - Movement protein translation - Movement protein
KM598396 (modified) - Replication associated protein translation - Replication associated protein
JX185418 (modified) - Replication associated protein translation - Replication associated protein
KT732819 (modified) - Replication associated protein translation - Replication associated protein
KP153371 (modified) - Replication associated protein translation - Replication associated protein
KP153373 (modified) - Replication associated protein translation - Replication associated protein
KP153372 (modified) - Replication associated protein translation - Replication associated protein
KT732822 (modified) - Replication associated protein translation - Replication associated protein
KT149406 (modified) - Replication associated protein translation - hypothetical protein CDS
KT149406 (modified) - Replication associated protein translation - Replication associated protein
KU043420.1 - Hypothetical protein translation - Replication associated protein
KU043420.1 - Hypothetical protein translation - Hypothetical protein
KR528558 (modified) - Replication associated protein interval 1 translation - Replication associated protein
JF755410 (modified) - ORF 4 (frame 3) translation - ORF 3 (frame 2)
JF755410 (modified) - ORF 4 (frame 3) translation - ORF 4 (frame 3)
JX305991 (modified) - hypothetical protein CDS translation 2 - hypothetical protein CDS
JX305992 (modified) - hypothetical protein CDS translation 2 - hypothetical protein CDS
JX305993 (modified) - hypothetical protein CDS translation 2 - hypothetical protein CDS
JX305994 (modified) - hypothetical protein CDS translation 2 - hypothetical protein CDS

JX305995 (modified) - hypothetical protein CDS translation 2 - hypothetical protein CDS
JX305996 (modified) - hypothetical protein CDS translation 2 - hypothetical protein CDS
JX305997 (modified) - hypothetical protein CDS translation 2 - hypothetical protein CDS
JQ023166 (modified) - hypothetical protein CDS translation 2 - hypothetical protein CDS
JX305998 (modified) - hypothetical protein CDS translation 2 - hypothetical protein CDS
KF133827 - hypothetical protein CDS translation 3 - hypothetical protein CDS
KF133827 - hypothetical protein CDS translation 3 - hypothetical protein CDS
KM874337 (modified) - Replication associated protein translation - Replication associated protein
KM874338 (modified) - Replication associated protein translation - Replication associated protein
KM874336 (modified) - Replication associated protein translation - Replication associated protein
KM874327 (modified) - Replication associated protein translation - Replication associated protein
KM874323 (modified) - Replication associated protein translation - Replication associated protein
KM874324 (modified) - Replication associated protein translation - Replication associated protein
KM874325 (modified) - Replication associated protein translation - Replication associated protein
KM874326 (modified) - Replication associated protein translation - Replication associated protein
MF351515.1 - Replication associated protein translation - Replication associated protein
NC_035475.1 - Replication associated protein translation - Replication associated protein
KM598391 (modified) - Replication associated protein translation - Replication associated protein
KU043449.1 - Hypothetical protein translation - Hypothetical protein
KP153504 (modified) - Replication associated protein translation - Replication associated protein
KM598392 (modified) - Replication associated protein translation - Replication associated protein
KY487835.1 - Replication associated protein translation - Replication associated protein
KU043396.1 - Replication associated protein translation - Replication associated protein
KX388507.1 - Replication associated protein translation - Replication associated protein
KF133815 - Replication associated protein translation - hypothetical protein CDS
KF133815 - Replication associated protein translation - Replication associated protein
KF133815 - hypothetical protein CDS translation - hypothetical protein CDS
KX388510.1 - Replication associated protein translation - Replication associated protein
KJ206566 (modified) - hypothetical protein CDS translation - hypothetical protein CDS
KT945163 (modified) - hypothetical protein CDS translation - hypothetical protein CDS
FJ959081 (modified) - Replication associated protein translation - Replication associated protein
KR528557 (modified) - Replication associated protein translation - Replication associated protein
JX904147 (modified) - Replication associated protein translation - Replication associated protein
GU456685 (modified) - Movement protein translation - Movement protein
KT388086 (modified) - Movement protein translation - Movement protein
KC108902 (modified) - Movement protein translation - Movement protein
KM821751 (modified) - Replication associated protein translation - Replication associated protein
KR528565 (modified) - Replication associated protein translation - Replication associated protein
KT149399 (modified) - hypothetical protein CDS translation 2 - hypothetical protein CDS
KF133808 - Replication associated protein translation - Replication associated protein
FJ959086 (modified) - Replication associated protein translation - Replication associated protein
KM874309 (modified) - Replication associated protein translation - Replication associated protein
KX388507.1 - hypothetical protein CDS translation - hypothetical protein CDS
KX388530.1 - hypothetical protein CDS translation 2 - hypothetical protein CDS
KM874319 (modified) - Capsid protein translation - Replication associated protein
KM874321 (modified) - Capsid protein translation - Replication associated protein
KM874322 (modified) - Capsid protein translation - Replication associated protein
AIF76255 - Q-loop/lid
AIF76255 - Q-loop/lid
KF133807 - Replication associated protein translation - Replication associated protein
KP153504 (modified) - hypothetical protein translation - hypothetical protein
KP153441 (modified) - Capsid protein translation - Replication associated protein
KT388088 (modified) - TrAP translation - Replication-associated protein
KT388088 (modified) - TrAP translation - TrAP
KT388086 (modified) - TrAP translation - Replication-associated protein
KT388086 (modified) - TrAP translation - TrAP
EU921828 (modified) - TrAP translation - Replication-associated protein
EU921828 (modified) - TrAP translation - TrAP
GU456685 (modified) - TrAP translation - Replication-associated protein
GU456685 (modified) - TrAP translation - TrAP
KC108902 (modified) - TrAP translation - Replication-associated protein
KC108902 (modified) - TrAP translation - TrAP
KU043446.1 - Replication associated protein translation - Replication associated protein
KM874294 (modified) - Replication associated protein translation - Replication associated protein
KM874292 (modified) - Replication associated protein translation - Replication associated protein
KM874293 (modified) - Replication associated protein translation - Replication associated protein
FJ665630 (modified) - Capsid protein translation - Movement protein
FJ665634 (modified) - Capsid protein translation - Movement protein
KT945162 (modified) - hypothetical protein CDS translation - Replication associated protein
KT945162 (modified) - hypothetical protein CDS translation - hypothetical protein CDS
KT149400 (modified) - hypothetical protein CDS translation - hypothetical protein CDS
KM874320 (modified) - Capsid protein translation - Replication associated protein
KT149405 (modified) - hypothetical protein CDS translation - Replication associated protein
KT149405 (modified) - hypothetical protein CDS translation - hypothetical protein CDS
KR186219 (modified) - Replication associated protein translation - Replication associated protein
KC248425 (modified) - Replication associated protein translation - Replication associated protein
KT149397 (modified) - hypothetical protein CDS translation 2 - Replication associated protein
KT149397 (modified) - hypothetical protein CDS translation 2 - hypothetical protein CDS
KF133807 - hypothetical protein CDS translation 2 - Replication associated protein
KF133807 - hypothetical protein CDS translation 2 - hypothetical protein CDS
KM874290 (modified) - Replication associated protein translation - Replication associated protein
KM874291 (modified) - Replication associated protein translation - Replication associated protein
KU203351 (modified) - Replication associated protein translation - Replication associated protein
KT149407 (modified) - Replication associated protein translation - Replication associated protein
KP153442 (modified) - Capsid protein translation - Replication associated protein
FJ959080 (modified) - Replication associated protein translation - Replication associated protein
KU203351 (modified) - ORF 6 (frame 3) translation - ORF 6 (frame 3)
KP153368 (modified) - Replication associated protein translation - Replication associated protein
KP153369 (modified) - Replication associated protein translation - Replication associated protein
MF118167.1 - hypothetical protein CDS translation 3 - hypothetical protein CDS
MF118166.1 - hypothetical protein CDS translation 3 - hypothetical protein CDS
KM598412 (modified) - Replication associated protein translation - Replication associated protein
MH545521.1 - Capsid protein translation - Replication associated protein
JF755404 (modified) - ORF 7 (frame 1) translation - ORF 7 (frame 1)
JF755405 (modified) - ORF 7 (frame 1) translation - ORF 7 (frame 1)
JF755406 (modified) - ORF 7 (frame 1) translation - ORF 7 (frame 1)
KT945154 (modified) - hypothetical protein CDS translation - Hypothetical 2
KT945154 (modified) - hypothetical protein CDS translation - hypothetical protein CDS
KP153466 (modified) - Replication associated protein translation - Replication associated protein
KP153464 (modified) - Replication associated protein translation - Replication associated protein
KP153465 (modified) - Replication associated protein translation - Replication associated protein
JF755404 (modified) - ORF 6 (frame 2) translation - ORF 6 (frame 2)
JF755405 (modified) - ORF 6 (frame 2) translation - ORF 6 (frame 2)
JF755406 (modified) - ORF 6 (frame 2) translation - ORF 6 (frame 2)
KP153468 (modified) - Replication associated protein translation - Replication associated protein
KP153467 (modified) - Replication associated protein translation - Replication associated protein
JF755404 (modified) - ORF 8 (frame 3) translation - ORF 8 (frame 3)
JF755405 (modified) - ORF 8 (frame 3) translation - ORF 8 (frame 3)
JF755406 (modified) - ORF 8 (frame 3) translation - ORF 8 (frame 3)
KP153461 (modified) - hypothetical 2 translation - hypothetical 2
KP153459 (modified) - hypothetical 2 translation - hypothetical 2
KP153463 (modified) - hypothetical 2 translation - hypothetical 2
KP153460 (modified) - hypothetical 2 translation - hypothetical 2
KJ955447 (modified) - viral protein RNase Z CDS translation - viral protein RNase Z CDS
KJ955448 (modified) - viral protein RNase Z CDS translation - viral protein RNase Z CDS
KJ955449 (modified) - viral protein RNase Z CDS translation - viral protein RNase Z CDS
KJ955450 (modified) - viral protein RNase Z CDS translation - viral protein RNase Z CDS
KJ955451 (modified) - viral protein RNase Z CDS translation - viral protein RNase Z CDS
KR134313 (modified) - viral protein RNase Z CDS translation - viral protein RNase Z CDS
KR134314 (modified) - viral protein RNase Z CDS translation - viral protein RNase Z CDS
KR134315 (modified) - viral protein RNase Z CDS translation - viral protein RNase Z CDS
KR134316 (modified) - viral protein RNase Z CDS translation - viral protein RNase Z CDS
KR134317 (modified) - viral protein RNase Z CDS translation - viral protein RNase Z CDS
KR134318 (modified) - viral protein RNase Z CDS translation - viral protein RNase Z CDS
KR134319 (modified) - viral protein RNase Z CDS translation - viral protein RNase Z CDS
KR134320 (modified) - viral protein RNase Z CDS translation - viral protein RNase Z CDS
KR134323 (modified) - viral protein RNase Z CDS translation - viral protein RNase Z CDS
KR134324 (modified) - viral protein RNase Z CDS translation - viral protein RNase Z CDS
KR134325 (modified) - viral protein RNase Z CDS translation - viral protein RNase Z CDS
KR134326 (modified) - viral protein RNase Z CDS translation - viral protein RNase Z CDS
KR134327 (modified) - viral protein RNase Z CDS translation - viral protein RNase Z CDS
KR134328 (modified) - viral protein RNase Z CDS translation - viral protein RNase Z CDS
KR134329 (modified) - viral protein RNase Z CDS translation - viral protein RNase Z CDS
KR134330 (modified) - viral protein RNase Z CDS translation - viral protein RNase Z CDS
KR134331 (modified) - viral protein RNase Z CDS translation - viral protein RNase Z CDS
KR134332 (modified) - viral protein RNase Z CDS translation - viral protein RNase Z CDS
KR134333 (modified) - viral protein RNase Z CDS translation - viral protein RNase Z CDS
KR134334 (modified) - viral protein RNase Z CDS translation - viral protein RNase Z CDS
KR134335 (modified) - viral protein RNase Z CDS translation - viral protein RNase Z CDS
KR134336 (modified) - viral protein RNase Z CDS translation - viral protein RNase Z CDS
KR134337 (modified) - viral protein RNase Z CDS translation - viral protein RNase Z CDS
KR134338 (modified) - viral protein RNase Z CDS translation - viral protein RNase Z CDS
KR134339 (modified) - viral protein RNase Z CDS translation - viral protein RNase Z CDS
KR134340 (modified) - viral protein RNase Z CDS translation - viral protein RNase Z CDS
KR134341 (modified) - viral protein RNase Z CDS translation - viral protein RNase Z CDS
KR134342 (modified) - viral protein RNase Z CDS translation - viral protein RNase Z CDS
KR134343 (modified) - viral protein RNase Z CDS translation - viral protein RNase Z CDS
KR134344 (modified) - viral protein RNase Z CDS translation - viral protein RNase Z CDS
KR134345 (modified) - viral protein RNase Z CDS translation - viral protein RNase Z CDS
KR134346 (modified) - viral protein RNase Z CDS translation - viral protein RNase Z CDS
KR134347 (modified) - viral protein RNase Z CDS translation - viral protein RNase Z CDS
KR134348 (modified) - viral protein RNase Z CDS translation - viral protein RNase Z CDS
KR134349 (modified) - viral protein RNase Z CDS translation - viral protein RNase Z CDS
KR134350 (modified) - viral protein RNase Z CDS translation - viral protein RNase Z CDS
KR134311 (modified) - viral protein RNase Z CDS translation - viral protein RNase Z CDS
KR134312 (modified) - viral protein RNase Z CDS translation - viral protein RNase Z CDS
KR134321 (modified) - viral protein RNase Z CDS translation - viral protein RNase Z CDS
KR134322 (modified) - viral protein RNase Z CDS translation - viral protein RNase Z CDS
KY052047.1 - Replication associated protein translation - Replication associated protein
KX388528.1 - hypothetical protein CDS translation - hypothetical protein CDS
KX388530.1 - hypothetical protein CDS translation - hypothetical protein CDS
KF133819 - Replication associated protein translation - Replication associated protein
KU043409.1 - Hypothetical protein translation - Hypothetical protein
KU043411.1 - Hypothetical protein translation - Hypothetical protein
KU043439.1 - Hypothetical protein translation - Hypothetical protein
KJ547618 (modified) - Replication associated protein translation - Replication associated protein
KU043413.1 - Hypothetical protein translation - Hypothetical protein
KP410285 (modified) - V3 CDS translation - V3 CDS
GU734126 (modified) - V3 CDS translation - V3 CDS
EU921828 (modified) - V3 CDS translation - V3 CDS
AF379637 (modified) - V3 translation - V3
HQ443515 (modified) - V3 CDS translation - V3 CDS
U49907 (modified) - V3 CDS translation - V3 CDS
KX388520.1 - hypothetical protein CDS translation - hypothetical protein CDS
MF351513.1 - Replication associated protein translation - Replication associated protein

MF351517.1 - Replication associated protein translation - Replication associated protein
MF351516.1 - Replication associated protein translation - Replication associated protein
MG522857.1 - Replication associated protein translation - Replication associated protein
MG522856.1 - Replication associated protein translation - Replication associated protein
MG522855.1 - Replication associated protein translation - Replication associated protein
MF351518.1 - Replication associated protein translation - Replication associated protein
MF351520.1 - Replication associated protein translation - Replication associated protein
MG522858.1 - Replication associated protein translation - Replication associated protein
MF351514.1 - Replication associated protein translation - Replication associated protein
KX388513.1 - Replication associated protein translation - Replication associated protein
JX908739 (modified) - Replication associated protein translation - Replication associated protein
KF133819 - hypothetical protein CDS translation - hypothetical protein CDS
JX908740 (modified) - Replication associated protein translation - Replication associated protein
KR528551 (modified) - Replication associated protein translation - Replication associated protein
FJ959079 (modified) - Replication associated protein interval 1 translation - Replication associated protein
KP153395 (modified) - hypothetical protein CDS translation - hypothetical protein CDS
KP153396 (modified) - hypothetical protein CDS translation - hypothetical protein CDS
KP153469 (modified) - Replication associated protein translation - Replication associated protein
KJ437671 (modified) - Replication-associated protein interval 1 translation - RepA CDS
KJ437671 (modified) - RepA CDS translation - Replication-associated protein interval 1
KJ437671 (modified) - Replication-associated protein interval 1 translation - Replication-associated protein
KJ437671 (modified) - RepA CDS translation - RepA CDS
KM874351 (modified) - Replication associated protein translation - Replication associated protein
KM874352 (modified) - Replication associated protein translation - Replication associated protein
KM874295 (modified) - Replication associated protein translation - Replication associated protein
KM874296 (modified) - Replication associated protein translation - Replication associated protein
KM874297 (modified) - Replication associated protein translation - Replication associated protein
KX388503.1 - Replication associated protein translation - Replication associated protein
KX388503.1 - Replication associated protein translation - stem loop
KX388503.1 - Replication associated protein translation - stem loop
KX388503.1 - Replication associated protein translation - Replication associated protein
KX388519.1 - hypothetical protein CDS translation - hypothetical protein CDS
KT149396 (modified) - Replication associated protein translation - Replication associated protein
KP153412 (modified) - Replication associated protein translation - Replication associated protein
KP153413 (modified) - Replication associated protein translation - Replication associated protein
MH545522.1 - Replication associated protein translation - Replication associated protein
KT732833 (modified) - Replication associated protein translation - Replication associated protein
AF379637 (modified) - Ren translation - TrAP
AF379637 (modified) - Ren translation - Ren
EU921828 (modified) - Ren translation - TrAP
EU921828 (modified) - Ren translation - Ren
KT149410 (modified) - Replication associated protein translation - Replication associated protein
KT149409 (modified) - Replication associated protein translation - Replication associated protein
KC108902 (modified) - Ren translation - TrAP
KC108902 (modified) - Ren translation - Ren
JF713718 (modified) - unknown CDS translation - unknown CDS
KF738880 (modified) - Capsid protein translation - Replication associated protein
KP153527 (modified) - Capsid protein translation - Replication associated protein
KT149408 (modified) - Replication associated protein translation - Replication associated protein
KM874360 (modified) - Replication associated protein translation - Replication associated protein
KM874361 (modified) - Replication associated protein translation - Replication associated protein
KR528548 (modified) - Replication associated protein translation - Replication associated protein
KM821767 (modified) - Replication associated protein interval 1 translation - Replication associated protein
KM821766 (modified) - hypothetical protein CDS translation 2 - hypothetical protein CDS
KJ417134 (modified) - Replication associated protein translation - Replication associated protein
KU043424.1 - Replication associated protein translation - Replication associated protein
KM598389 (modified) - Replication associated protein translation - Replication associated protein
KP153370 (modified) - Replication associated protein translation - Replication associated protein
JF938078 (modified) - Replication associated protein translation - Replication associated protein
KF133820 - hypothetical protein CDS translation - hypothetical protein CDS
KM874344 (modified) - Replication associated protein translation - Replication associated protein
KM874345 (modified) - Replication associated protein translation - Replication associated protein
KM874343 (modified) - Replication associated protein translation - Replication associated protein
KP153445 (modified) - Replication associated protein translation - Replication associated protein
JQ023166 (modified) - hypothetical protein CDS interval 1 translation - hypothetical protein CDS
JX305991 (modified) - hypothetical protein CDS interval 1 translation - hypothetical protein CDS
JX305992 (modified) - hypothetical protein CDS interval 1 translation - hypothetical protein CDS
JX305993 (modified) - hypothetical protein CDS interval 1 translation - hypothetical protein CDS
JX305994 (modified) - hypothetical protein CDS interval 1 translation - hypothetical protein CDS
JX305995 (modified) - hypothetical protein CDS interval 1 translation - hypothetical protein CDS
JX305996 (modified) - hypothetical protein CDS interval 1 translation - hypothetical protein CDS
JX305997 (modified) - hypothetical protein CDS interval 1 translation - hypothetical protein CDS
JQ023166 (modified) - hypothetical protein CDS interval 1 translation - Replication associated protein
JX305991 (modified) - hypothetical protein CDS interval 1 translation - Replication associated protein
JX305992 (modified) - hypothetical protein CDS interval 1 translation - Replication associated protein
JX305993 (modified) - hypothetical protein CDS interval 1 translation - Replication associated protein
JX305994 (modified) - hypothetical protein CDS interval 1 translation - Replication associated protein
JX305995 (modified) - hypothetical protein CDS interval 1 translation - Replication associated protein
JX305996 (modified) - hypothetical protein CDS interval 1 translation - Replication associated protein
JX305997 (modified) - hypothetical protein CDS interval 1 translation - Replication associated protein
JX305998 (modified) - hypothetical protein CDS interval 1 translation - hypothetical protein CDS
JX305998 (modified) - hypothetical protein CDS interval 1 translation - Replication associated protein
U49907 (modified) - C2 CDS translation - Replication-associated protein
U49907 (modified) - C2 CDS translation - C2 CDS
KM598390 (modified) - Replication associated protein translation - Replication associated protein
KP153395 (modified) - Replication associated protein translation - Replication associated protein
KP153396 (modified) - Replication associated protein translation - Replication associated protein
KM821749 (modified) - Replication associated protein interval 1 translation - Replication associated protein
KM598399 (modified) - Replication associated protein translation - Replication associated protein
MF118168.1 - hypothetical protein CDS translation - hypothetical protein CDS
KF133810 - Replication associated protein interval 2 translation - Replication associated protein
KF133821 - Replication associated protein translation - Replication associated protein
KM874320 (modified) - Replication associated protein translation - Replication associated protein
KF133811 - unk translation - unk
KT945164 (modified) - Replication associated protein translation - Replication associated protein
KF133825 - hypothetical protein CDS translation - hypothetical protein CDS
KU043405.1 - Hypothetical protein translation - Hypothetical protein
KX388516.1 - Replication associated protein interval 1 translation - Replication associated protein
KM573765 (modified) - ORF2 CDS translation - ORF2 CDS
MH545532.1 - Replication associated protein translation - Replication associated protein
KT732825 (modified) - Replication associated protein interval 1 translation - Replication associated protein
MH545517.1 - Replication associated protein translation - Replication associated protein
KM874350 (modified) - Replication associated protein interval 1 translation - Replication associated protein
KU043436.1 - Replication associated protein translation - Replication associated protein
KM821761 (modified) - Replication associated protein translation - Replication associated protein
JX185418 (modified) - Capsid protein translation - Replication associated protein
KM821750 (modified) - Capsid protein translation - Replication associated protein
MH552520.1 - Capsid protein translation - hypothetical protein CDS
KT732818 (modified) - Replication associated protein translation - Replication associated protein
HQ335087 (modified) - Replication associated protein interval 1 translation - Replication associated protein
KP153520 (modified) - Replication associated protein translation - Replication associated protein
KP153521 (modified) - Replication associated protein translation - Replication associated protein
KF738884 (modified) - Replication associated protein translation - Replication associated protein
KP153519 (modified) - Replication associated protein translation - Replication associated protein
KP153518 (modified) - Replication associated protein translation - Replication associated protein
KP153517 (modified) - Replication associated protein translation - Replication associated protein
KF738885 (modified) - Replication associated protein translation - Replication associated protein
KC248419 (modified) - Replication associated protein translation - Replication associated protein
KT945154 (modified) - Hypothetical 2 translation - Replication associated protein
KF133813 - hypothetical protein CDS translation - hypothetical protein CDS
KF481961 (modified) - Capsid protein translation - nonstructural protein 2 CDS
KT732816 (modified) - Replication associated protein translation - Replication associated protein
KF738873 (modified) - Replication associated protein translation - Replication associated protein
KP153437 (modified) - Capsid protein translation - Replication associated protein
KP153438 (modified) - Capsid protein translation - Replication associated protein
KP153439 (modified) - Capsid protein translation - Replication associated protein
KP153440 (modified) - Capsid protein translation - Replication associated protein
KR528552 (modified) - Replication associated protein translation - Replication associated protein
JQ837277 (modified) - hypothetical protein CDS translation - hypothetical protein CDS
KF246569 (modified) - Replication associated protein translation - Replication associated protein
KM573776 (modified) - ORF2 CDS translation - ORF2 CDS
AEL28791 - CDS
AEL28791 - replication-associated protein
KJ547620 (modified) - Replication associated protein interval 1 translation - Replication associated protein
MH616757.1 - Replication associated protein translation - hypothetical protein CDS
MH616757.1 - Replication associated protein translation - Replication associated protein
FJ959081 (modified) - hypothetical protein CDS translation - hypothetical protein CDS
KT149405 (modified) - Replication associated protein translation - Replication associated protein
KF133823 - Replication associated protein interval 1 translation - Replication associated protein
KX388512.1 - Replication associated protein translation - Replication associated protein
KT149404 (modified) - hypothetical protein CDS translation 3 - hypothetical protein CDS
KJ938716 (modified) - Replication associated protein translation - Replication associated protein
FJ959083 (modified) - Replication associated protein interval 1 translation - Replication associated protein
AF379637 (modified) - Replication-associated protein translation - L4 CDS
AF379637 (modified) - Replication-associated protein translation - Replication-associated protein
MH616747.1 - Capsid protein translation - hypothetical protein CDS
KJ547647 (modified) - hypothetical protein CDS translation 2 - hypothetical protein CDS
KJ547652 (modified) - Replication associated protein translation - Replication associated protein
KX388521.1 - Replication associated protein translation - Replication associated protein
KX388522.1 - Replication associated protein translation - Replication associated protein
KX388523.1 - Replication associated protein translation - Replication associated protein
KX388524.1 - Replication associated protein translation - Replication associated protein
KX388525.1 - Replication associated protein translation - Replication associated protein
KX388526.1 - Replication associated protein translation - Replication associated protein
KP410285 (modified) - Replication-associated protein interval 1 translation - Replication-associated protein
KR528555 (modified) - Replication associated protein translation - Replication associated protein
KR528556 (modified) - Replication associated protein translation - Replication associated protein
KF481961 (modified) - Replication associated protein translation - Replication associated protein
KJ547632 (modified) - hypothetical protein CDS translation - hypothetical protein CDS
KM386645 (modified) - Replication-associated protein translation - Replication-associated protein
MH378453.1 - Replication associated protein interval 1 translation - RepA CDS
MH378453.1 - RepA CDS translation - Replication associated protein
MH378453.1 - Replication associated protein interval 1 translation - Replication associated protein
MH378453.1 - RepA CDS translation - RepA CDS
KM598411 (modified) - Replication associated protein interval 1 translation - Replication associated protein
KF133815 - ORF 7 (frame 3) translation - ORF 7 (frame 3)
MH617390.1 - Capsid protein translation - hypothetical protein CDS
KR528543 (modified) - Capsid protein translation - Replication associated protein
MH545533.1 - Replication associated protein translation - Replication associated protein
MH425573.1 - Replication associated protein translation - Replication associated protein
GU734126 (modified) - Replication-associated protein translation - Replication-associated protein
KT214389 (modified) - Replication-associated protein interval 1 translation - RepA CDS
KT214389 (modified) - RepA CDS translation - Replication-associated protein interval 1
KT214389 (modified) - Replication-associated protein interval 1 translation - Replication-associated protein
KT214389 (modified) - RepA CDS translation - RepA CDS
U49907 (modified) - Replication-associated protein translation - Replication-associated protein

KT214386 (modified) - Replication-associated protein interval 1 translation - RepA CDS
KT214386 (modified) - RepA CDS translation - Replication-associated protein interval 1
KT214386 (modified) - Replication-associated protein interval 1 translation - Replication-associated protein
KT214386 (modified) - RepA CDS translation - RepA CDS
KT214373 (modified) - Replication-associated protein interval 1 translation - RepA CDS
KT214373 (modified) - RepA CDS translation - Replication-associated protein interval 1
KT214373 (modified) - Replication-associated protein interval 1 translation - Replication-associated protein
KT214373 (modified) - RepA CDS translation - RepA CDS
JX094280 (modified) - Replication-associated protein interval 1 translation - Replication-associated protein
KP153443 (modified) - Capsid protein translation - Replication associated protein
KP153444 (modified) - Capsid protein translation - Replication associated protein
JF713717 (modified) - unknown CDS translation 2 - unknown CDS
JF713716 (modified) - unknown CDS translation - unknown CDS
EU921828 (modified) - Replication-associated protein translation - Replication-associated protein
KC108902 (modified) - Replication-associated protein translation - Replication-associated protein
GU456685 (modified) - Replication-associated protein translation - Replication-associated protein
KF133828 - Replication associated protein translation - Replication associated protein
FJ959084 (modified) - Replication associated protein translation - Replication associated protein
KF133812 - Replication associated protein translation - Replication associated protein
KF133827 - Replication associated protein translation - Replication associated protein
KF133821 - hypothetical protein CDS translation - hypothetical protein CDS
KU043415.1 - Hypothetical protein translation - Hypothetical protein
DQ458791 (modified) - Replication-associated protein interval 1 translation - C1 CDS
DQ458791 (modified) - C1 CDS translation - Replication-associated protein interval 1
DQ458791 (modified) - Replication-associated protein interval 1 translation - Replication-associated protein
DQ458791 (modified) - C1 CDS translation - C1 CDS
KM510191 (modified) - Replication associated protein translation - Replication associated protein
KT862231 (modified) - Replication associated protein translation - Replication associated protein
KR528569 (modified) - Replication associated protein interval 1 translation - Replication associated protein
KP153501 (modified) - Replication associated protein translation - Replication associated protein
JF755410 (modified) - ORF 3 (frame 2) translation - ORF 3 (frame 2)
AF027217 (modified) - Replication associated protein translation - Replication associated protein
AF027217 (modified) - Replication associated protein translation - Replication associated protein
AF071879 (modified) - Replication associated protein translation - Replication associated protein
AF071879 (modified) - Replication associated protein translation - Replication associated protein
KM386645 (modified) - TrAP translation - Replication-associated protein
KM386645 (modified) - TrAP translation - TrAP
JQ837277 (modified) - Replication associated protein translation - Replication associated protein
X84735 (modified) - Replication-associated protein translation - Replication-associated protein
KT862235 (modified) - Replication associated protein translation - Replication associated protein
KT945163 (modified) - Replication associated protein translation - Replication associated protein
KT862256 (modified) - Replication associated protein translation - Replication associated protein
KF133810 - Replication associated protein interval 1 translation - Replication associated protein
KM874359 (modified) - Replication associated protein translation - Replication associated protein
KM821768 (modified) - Replication associated protein translation - Replication associated protein
KX388497.1 - Capsid protein translation - hypothetical protein CDS
KP153414 (modified) - Replication associated protein translation - Replication associated protein
KP153415 (modified) - Replication associated protein translation - Replication associated protein
KP153416 (modified) - Replication associated protein translation - Replication associated protein
KM598408 (modified) - Replication associated protein translation - Replication associated protein
KR528545 (modified) - Replication associated protein translation - Replication associated protein
MH545536.1 - Replication associated protein translation - Replication associated protein
KP153474 (modified) - Replication associated protein translation - Replication associated protein
KP153475 (modified) - Replication associated protein translation - Replication associated protein
KF133818 - Replication associated protein translation - Replication associated protein
JX904333 (modified) - Replication associated protein translation - Replication associated protein
LC133373 - Replication associated protein translation - Replication associated protein
KX618694 - Ren translation - TrAP
KX618694 - Ren translation - Ren
KC248414 (modified) - ORF 2 (frame 1) translation - Replication associated protein
KC248414 (modified) - ORF 2 (frame 1) translation - ORF 2 (frame 1)
KM874319 (modified) - Replication associated protein translation - Replication associated protein
KM874321 (modified) - Replication associated protein translation - Replication associated protein
KM874322 (modified) - Replication associated protein translation - Replication associated protein
KC248420 (modified) - hypothetical protein CDS translation - Replication associated protein
KC248420 (modified) - hypothetical protein CDS translation - hypothetical protein CDS
KF133823 - hypothetical protein CDS translation 2 - hypothetical protein CDS
X84735 (modified) - Capsid protein translation - Movement protein
GU734126 (modified) - C2 CDS translation - Replication-associated protein
GU734126 (modified) - C2 CDS translation - C2 CDS
KC248412.1 - Replication associated protein translation - Replication associated protein
KX618694 - TrAP translation - Replication associated protein
KX618694 - TrAP translation - TrAP
KP153394 (modified) - Replication associated protein interval 1 translation - Replication associated protein
KU043439.1 - Replication associated protein translation - Replication associated protein
KX388495.1 - Replication associated protein translation - Replication associated protein
KT732821 (modified) - Replication associated protein translation - Replication associated protein
KT732820 (modified) - Replication associated protein translation - Replication associated protein
KX388514.1 - hypothetical protein CDS translation - hypothetical protein CDS
KF133814 - Replication associated protein interval 1 translation - Replication associated protein
KP153522 (modified) - Replication associated protein translation - Replication associated protein
KM874339 (modified) - Replication associated protein translation - Replication associated protein
KM573763 (modified) - ORF2 CDS translation - ORF2 CDS
KX388494.1 - hypothetical protein CDS translation 3 - hypothetical protein CDS
KF133815 - hypothetical protein CDS interval 1 translation - Replication associated protein
KF133815 - hypothetical protein CDS interval 1 translation - hypothetical protein CDS
JF713716 (modified) - Replication associated protein translation - Replication associated protein
JF713717 (modified) - Replication associated protein translation - Replication associated protein
KP153499 (modified) - Replication associated protein translation - Replication associated protein
KX388494.1 - hypothetical protein CDS translation - hypothetical protein CDS
KT149401 (modified) - hypothetical protein CDS translation - hypothetical protein CDS
KT732830 (modified) - Replication associated protein translation - Replication associated protein
JX185415 (modified) - Capsid protein translation - Replication associated protein
KF133823 - hypothetical protein CDS translation - hypothetical protein CDS
KR528560 (modified) - Replication associated protein interval 1 translation - Replication associated protein
KC543331 (modified) - Replication associated protein translation - Replication associated protein
KC794539 (modified) - Replication associated protein translation - Replication associated protein
KC794540 (modified) - Replication associated protein translation - Replication associated protein
MH545526.1 - Replication associated protein translation - Replication associated protein
KF133814 - hypothetical protein CDS translation - hypothetical protein CDS
MF327576.1 - Capsid protein translation - Replication associated protein
AF379637 (modified) - Capsid protein translation - V2
EU921828 (modified) - Capsid protein translation - V2 CDS
KT149397 (modified) - hypothetical protein CDS translation - hypothetical protein CDS
KT149401 (modified) - Replication associated protein translation - hypothetical protein CDS
KT149401 (modified) - Replication associated protein translation - Replication associated protein
KU043429.1 - Replication associated protein translation - Replication associated protein
KF133825 - Replication associated protein translation - Replication associated protein
FJ959080 (modified) - hypothetical protein CDS translation - hypothetical protein CDS
U49907 (modified) - Capsid protein translation - V2 CDS
MG846356.1 - Replication associated protein translation - Replication associated protein
MG846355.1 - Replication associated protein translation - Replication associated protein
KT862232 (modified) - Replication associated protein translation - Replication associated protein
KT149395 (modified) - hypothetical protein CDS translation - hypothetical protein CDS
KT149397 (modified) - Replication associated protein translation - Replication associated protein
KT149398 (modified) - Replication associated protein translation - Replication associated protein
KF133818 - hypothetical protein CDS translation 3 - hypothetical protein CDS
KF133818 - hypothetical protein CDS translation 3 - hypothetical protein CDS
KM598396 (modified) - Capsid protein translation - Replication associated protein
KT945154 (modified) - Replication associated protein translation - Replication associated protein
KU043400.1 - Replication associated protein translation - Replication associated protein
MG522854.1 - Replication associated protein translation - Replication associated protein
MG522853.1 - Replication associated protein translation - Replication associated protein
KM598393 (modified) - Replication associated protein translation - Replication associated protein
MF351519.1 - Replication associated protein translation - Replication associated protein
KP153364 (modified) - Replication associated protein translation - Replication associated protein
KP153365 (modified) - Replication associated protein translation - Replication associated protein
KP153366 (modified) - Replication associated protein translation - Replication associated protein
KP153367 (modified) - Replication associated protein translation - Replication associated protein
MH545515.1 - Replication associated protein interval 1 translation - Replication associated protein
KU043425.1 - Replication associated protein translation - Replication associated protein
MH545534.1 - Replication associated protein translation - Replication associated protein
MF327574.1 - Replication associated protein translation - Replication associated protein
KX388508.1 - hypothetical protein CDS translation - hypothetical protein CDS
KM874315 (modified) - Replication associated protein translation - Replication associated protein
KM874316 (modified) - Replication associated protein translation - Replication associated protein
KM874317 (modified) - Replication associated protein translation - Replication associated protein
KM874318 (modified) - Replication associated protein translation - Replication associated protein
KT862230 (modified) - Replication associated protein translation - Replication associated protein
KF133813 - Replication associated protein interval 1 translation - Replication associated protein
KF133827 - hypothetical protein CDS translation 2 - hypothetical protein CDS
KP153451 (modified) - Replication associated protein translation - Replication associated protein
KP153452 (modified) - Replication associated protein translation - Replication associated protein
KP153453 (modified) - Replication associated protein translation - Replication associated protein
KU043420.1 - Hypothetical protein translation 2 - Replication associated protein
KU043420.1 - Hypothetical protein translation 2 - Hypothetical protein
KP153441 (modified) - Replication associated protein translation - Replication associated protein
KM598404 (modified) - Replication associated protein translation - Replication associated protein
KM598405 (modified) - Replication associated protein translation - Replication associated protein
MH545541.1 - Replication associated protein translation - Replication associated protein
MH425572.1 - Replication associated protein interval 1 translation - Replication associated protein
KU043411.1 - Hypothetical protein translation 2 - Hypothetical protein
KX388527.1 - Replication associated protein translation - Replication associated protein
FJ665634 (modified) - Movement protein translation - Movement protein
FJ665630 (modified) - Movement protein translation - Movement protein
KM874334 (modified) - Replication associated protein translation - Replication associated protein
KM874333 (modified) - Replication associated protein translation - Replication associated protein
KM874335 (modified) - Replication associated protein translation - Replication associated protein
KM874332 (modified) - Replication associated protein translation - Replication associated protein
KF133816 - Replication associated protein translation - Replication associated protein
KC441518 (modified) - Replication associated protein translation - Replication associated protein
GU456685 (modified) - Capsid protein translation - Movement protein
KT388086 (modified) - Capsid protein translation - Movement protein
KC108902 (modified) - Capsid protein translation - Movement protein
KF738883 (modified) - Capsid protein translation - Replication associated protein
KU043416.1 - Replication associated protein translation - Replication associated protein
KP202150 (modified) - Replication associated protein translation - Replication associated protein
MH545520.1 - Replication associated protein translation - Replication associated protein
KP153377 (modified) - Replication associated protein translation - Replication associated protein
KP153378 (modified) - Replication associated protein translation - Replication associated protein
KX388499.1 - Replication associated protein translation - Replication associated protein
KU043449.1 - Replication associated protein translation - Replication associated protein
KJ547648 (modified) - Replication associated protein translation - Replication associated protein

KM821769 (modified) - Replication associated protein translation - Replication associated protein
KT732828 (modified) - Replication associated protein translation - Replication associated protein
KF133807 - hypothetical protein CDS translation - hypothetical protein CDS
KX388514.1 - Replication associated protein translation - Replication associated protein
KX388515.1 - Replication associated protein translation - Replication associated protein
KR528564 (modified) - Replication associated protein interval 1 translation - Replication associated protein
KM598403 (modified) - Replication associated protein translation - Replication associated protein
KM598401 (modified) - Replication associated protein translation - Replication associated protein
KM598402 (modified) - Replication associated protein translation - Replication associated protein
KF133820 - Replication associated protein interval 1 translation - Replication associated protein
KU343138.1 - Hypothetical protein translation - Hypothetical protein
KM972725 (modified) - Replication associated protein interval 1 translation - Replication associated protein
KM972726 (modified) - Replication associated protein interval 1 translation - Replication associated protein
KC543331 (modified) - hypothetical protein CDS translation - hypothetical protein CDS
KC794539 (modified) - hypothetical protein CDS translation - hypothetical protein CDS
KC794540 (modified) - hypothetical protein CDS translation - hypothetical protein CDS
KX388496.1 - Replication associated protein translation - Replication associated protein
JX185416 (modified) - Capsid protein translation - Replication associated protein
JX185417 (modified) - Capsid protein translation - Replication associated protein
KU043436.1 - unk translation - unk
KR704912 - hypothetical protein CDS translation - hypothetical protein CDS
KR131749 (modified) - putative movement protein CDS translation - ORF1 CDS
KR131749 (modified) - putative movement protein CDS translation - putative movement protein CDS
KT600067 (modified) - Replication associated protein translation - Replication associated protein
KT600066 (modified) - Replication associated protein translation - Replication associated protein
KT600065 (modified) - Replication associated protein translation - Replication associated protein
KP153509 (modified) - Replication associated protein translation - Replication associated protein
KP153507 (modified) - Replication associated protein translation - Replication associated protein
KP153510 (modified) - Replication associated protein translation - Replication associated protein
KP153508 (modified) - Replication associated protein translation - Replication associated protein
KM821765 (modified) - Replication associated protein translation - Replication associated protein
KP153483 (modified) - Replication associated protein translation - Replication associated protein
KP153484 (modified) - Replication associated protein translation - Replication associated protein
KC248422 (modified) - Replication associated protein translation - Replication associated protein
KX388508.1 - Replication associated protein translation - Replication associated protein
KM874366 (modified) - Replication associated protein translation - Replication associated protein
KM874367 (modified) - Replication associated protein translation - Replication associated protein
KM874368 (modified) - Replication associated protein translation - Replication associated protein
KM598395 (modified) - Replication associated protein translation - Replication associated protein
KR528562 (modified) - Replication associated protein translation - Replication associated protein
KM821754 (modified) - Replication associated protein translation - Replication associated protein
JF713716 (modified) - unknown CDS translation 2 - unknown CDS
KJ547632 (modified) - Replication associated protein translation - Replication associated protein
KP005453 (modified) - Replication associated protein interval 1 translation - Replication associated protein
KP005454 (modified) - Replication associated protein interval 1 translation - Replication associated protein
KM510191 (modified) - hypothetical protein CDS translation - hypothetical protein CDS
KP153390 (modified) - Replication associated protein translation - Replication associated protein
KP153391 (modified) - Replication associated protein translation - Replication associated protein
KP153392 (modified) - Replication associated protein translation - Replication associated protein
KP153393 (modified) - Replication associated protein translation - Replication associated protein
KJ547646 (modified) - Replication associated protein translation - Replication associated protein
KP153456 (modified) - Replication associated protein translation - Replication associated protein
KP153457 (modified) - Replication associated protein translation - Replication associated protein
KP153458 (modified) - Replication associated protein translation - Replication associated protein
KM573767 (modified) - ORF2 CDS translation - ORF2 CDS
GU734126 (modified) - Capsid protein translation - V2 CDS
HQ443515 (modified) - Capsid protein translation - V2 CDS
KJ547621 (modified) - Replication associated protein translation - Replication associated protein
JN704610 (modified) - Replication associated protein translation - Replication associated protein
KM874328 (modified) - Replication associated protein translation - Replication associated protein
KM598397 (modified) - Replication associated protein translation - Replication associated protein
KP153460 (modified) - Replication associated protein translation - Replication associated protein
KP153462 (modified) - Replication associated protein translation - Replication associated protein
KT149403 (modified) - hypothetical protein CDS translation - hypothetical protein CDS
MH545529.1 - Replication associated protein interval 1 translation - Replication associated protein
KJ547650 (modified) - Replication associated protein translation - Replication associated protein
KR528547 (modified) - Replication associated protein interval 1 translation - Replication associated protein
MH545518.1 - Replication associated protein translation - Replication associated protein
MH545519.1 - Replication associated protein translation - Replication associated protein
JX185415 (modified) - Replication associated protein translation - Replication associated protein
KT732823 (modified) - Replication associated protein translation - Replication associated protein
KP153496 (modified) - Replication associated protein translation - Replication associated protein
KP153497 (modified) - Replication associated protein translation - Replication associated protein
KP153498 (modified) - Replication associated protein translation - Replication associated protein
MH545542.1 - Replication associated protein translation - Replication associated protein
KT149412 (modified) - Replication associated protein translation - Replication associated protein
JF713718 (modified) - Replication associated protein translation - Replication associated protein
KM874348 (modified) - Replication associated protein translation - Replication associated protein
KM874349 (modified) - Replication associated protein translation - Replication associated protein
KM874347 (modified) - Replication associated protein translation - Replication associated protein
MH617766.1 - Replication associated protein translation - hypothetical protein CDS
MH617766.1 - Replication associated protein translation - Replication associated protein
KU043420.1 - Replication associated protein translation - Hypothetical protein
KU043420.1 - Replication associated protein translation - Replication associated protein
KF133822 - Replication associated protein translation - Replication associated protein
JF912805 (modified) - Replication associated protein translation - Replication associated protein
KC248416 (modified) - Replication associated protein translation - Replication associated protein
MH545521.1 - Replication associated protein translation - Replication associated protein
KP153407 (modified) - Replication associated protein translation - Replication associated protein
KT149408 (modified) - hypothetical protein CDS translation - hypothetical protein CDS
KP153461 (modified) - Replication associated protein translation - Replication associated protein
KP153459 (modified) - Replication associated protein translation - Replication associated protein
KP153463 (modified) - Replication associated protein translation - Replication associated protein
KM821759 (modified) - Replication associated protein interval 1 translation - Replication associated protein
KF133811 - Replication associated protein translation - Replication associated protein
KF133824 - Replication associated protein translation - Replication associated protein
KM874305 (modified) - Replication associated protein translation - Replication associated protein
KM874306 (modified) - Replication associated protein translation - Replication associated protein
KM874308 (modified) - Replication associated protein translation - Replication associated protein
KM874304 (modified) - Replication associated protein translation - Replication associated protein
KM874307 (modified) - Replication associated protein translation - Replication associated protein
FJ959082 (modified) - Replication associated protein translation - Replication associated protein
MH545540.1 - Replication associated protein interval 1 translation - Replication associated protein
MF118167.1 - hypothetical protein CDS translation - hypothetical protein CDS
MF118166.1 - hypothetical protein CDS translation - hypothetical protein CDS
MH545539.1 - Replication associated protein translation - Replication associated protein
KT732827 (modified) - Replication associated protein translation - Replication associated protein
MH552520.1 - Replication associated protein translation - hypothetical protein CDS
MH552520.1 - Replication associated protein translation - Replication associated protein
MH425569.1 - Replication associated protein translation - Replication associated protein
KT149411 (modified) - Replication associated protein translation - Replication associated protein
JF713719 (modified) - Replication associated protein translation - Replication associated protein
KM598398 (modified) - Replication associated protein translation - Replication associated protein
KM821756 (modified) - Replication associated protein translation - Replication associated protein
KT149404 (modified) - Replication associated protein translation - Replication associated protein
KJ547649 (modified) - Replication associated protein translation - Replication associated protein
KF133811 - unk translation 2 - unk
KF133824 - unk translation - unk
KM821760 (modified) - Replication associated protein translation - Replication associated protein
KJ433989 (modified) - Replication associated protein translation - Replication associated protein
KR902498 (modified) - hypothetical protein CDS translation 2 - hypothetical protein CDS
KR902498 (modified) - hypothetical protein CDS translation 3 - hypothetical protein CDS
JX559622 (modified) - hypothetical protein CDS translation - hypothetical protein CDS
KP153402 (modified) - Replication associated protein translation - Replication associated protein
KP153360 (modified) - Replication associated protein translation - Replication associated protein
KT388088 (modified) - Capsid protein translation - Movement protein
KM821764 (modified) - Replication associated protein translation - Replication associated protein
JX559621 (modified) - hypothetical protein CDS translation - hypothetical protein CDS
KX388497.1 - Replication associated protein translation - Replication associated protein
KT732832 (modified) - Replication associated protein translation - Replication associated protein
KX388494.1 - hypothetical protein CDS translation 2 - hypothetical protein CDS
JQ898333 (modified) - hypothetical protein CDS translation - hypothetical protein CDS
KT862237 (modified) - Replication associated protein translation - Replication associated protein
KP153454 (modified) - Replication associated protein translation - Replication associated protein
KP153455 (modified) - Replication associated protein translation - Replication associated protein
KT149399 (modified) - hypothetical protein CDS translation - hypothetical protein CDS
KT149401 (modified) - hypothetical protein CDS translation 3 - hypothetical protein CDS
GU456685 (modified) - Ren translation - TrAP
GU456685 (modified) - Ren translation - Ren
KX388504.1 - Replication associated protein translation - Replication associated protein
KX388509.1 - Replication associated protein translation - Replication associated protein
KR131749 (modified) - ORF1 CDS translation - ORF1 CDS
JX559622 (modified) - Replication associated protein translation - Replication associated protein
JX559621 (modified) - Replication associated protein translation - Replication associated protein
KT149404 (modified) - hypothetical protein CDS translation - hypothetical protein CDS
KM874353 (modified) - Replication associated protein translation - Replication associated protein
KM874346 (modified) - Replication associated protein translation - Replication associated protein
KM874329 (modified) - Replication associated protein translation - Replication associated protein
KM874330 (modified) - Replication associated protein translation - Replication associated protein
KM874331 (modified) - Replication associated protein translation - Replication associated protein
KU343138.1 - Replication associated protein translation - Replication associated protein
KP153443 (modified) - Replication associated protein translation - Replication associated protein
KP153444 (modified) - Replication associated protein translation - Replication associated protein
KP153408 (modified) - Replication associated protein translation - Replication associated protein
KT945165 (modified) - Replication associated protein interval 1 translation - Replication associated protein
KY487829.1 - Replication associated protein translation - Replication associated protein
JQ898333 (modified) - Replication associated protein translation - Replication associated protein
KT149399 (modified) - Replication associated protein translation - hypothetical protein CDS
KT149399 (modified) - Replication associated protein translation - Replication associated protein
KM874298 (modified) - Replication associated protein translation - Replication associated protein
KM874299 (modified) - Replication associated protein translation - Replication associated protein
KM874300 (modified) - Replication associated protein translation - Replication associated protein
KJ547631 (modified) - hypothetical protein CDS translation - hypothetical protein CDS
KU058671.1 - Replication associated protein translation - Replication associated protein
KR528550 (modified) - Replication associated protein translation - Replication associated protein
KF133818 - hypothetical protein CDS translation 2 - hypothetical protein CDS
MH425571.1 - Replication associated protein translation - Replication associated protein
KP153503 (modified) - Replication associated protein translation - Replication associated protein
KX388527.1 - hypothetical protein CDS translation 2 - hypothetical protein CDS
KX388529.1 - hypothetical protein CDS translation - hypothetical protein CDS
KU043403.1 - Replication associated protein translation - Replication associated protein

JF713716 (modified) - unknown CDS translation 3 - unknown CDS
KT149411 (modified) - hypothetical protein CDS translation - hypothetical protein CDS
KU043411.1 - Replication associated protein translation - Replication associated protein
MF327575.1 - unk translation - unk
KU043422.1 - Replication associated protein translation - Replication associated protein
KU043421.1 - Replication associated protein translation - Replication associated protein
JF755402 (modified) - ORF 10 (frame 2) translation - ORF 10 (frame 2)
KT862257 (modified) - Replication associated protein translation - Replication associated protein
KM821752 (modified) - Replication associated protein translation - Replication associated protein
KP153397 (modified) - Replication associated protein translation - Replication associated protein
KP153398 (modified) - Replication associated protein translation - Replication associated protein
KP153399 (modified) - Replication associated protein translation - Replication associated protein
KP153400 (modified) - Replication associated protein translation - Replication associated protein
KP153401 (modified) - Replication associated protein translation - Replication associated protein
KF133814 - hypothetical protein CDS translation 3 - hypothetical protein CDS
KJ547626 (modified) - Replication associated protein interval 1 translation - Replication associated protein
KC248422 (modified) - hypothetical protein CDS translation - Replication associated protein
KC248422 (modified) - hypothetical protein CDS translation - hypothetical protein CDS
KR528561 (modified) - Replication associated protein translation - Replication associated protein
KM598400 (modified) - Replication associated protein interval 1 translation - Replication associated protein
KP153428 (modified) - Replication associated protein translation - Replication associated protein
KP153427 (modified) - Replication associated protein translation - Replication associated protein
FJ959077 (modified) - Replication associated protein translation - Replication associated protein
KX388509.1 - hypothetical protein CDS translation 2 - hypothetical protein CDS
KM821758 (modified) (reversed) - Replication associated protein translation - Replication associated protein
KM821758 (modified) - Replication associated protein translation - Replication associated protein
KM821757 (modified) - Replication associated protein translation - Replication associated protein
KC248414 (modified) - Replication associated protein translation - Replication associated protein
KP153379 (modified) - Replication associated protein translation - Replication associated protein
KP153380 (modified) - Replication associated protein translation - Replication associated protein
KP153381 (modified) - Replication associated protein translation - Replication associated protein
KP153382 (modified) - Replication associated protein translation - Replication associated protein
KP153383 (modified) - Replication associated protein translation - Replication associated protein
KT862233 (modified) - Replication associated protein translation - Replication associated protein
KT149403 (modified) - Replication associated protein translation - Replication associated protein
KU043405.1 - Replication associated protein translation - Replication associated protein
KX388519.1 - Replication associated protein translation - Replication associated protein
KX388520.1 - Replication associated protein translation - Replication associated protein
MH616973.1 - Capsid protein translation - hypothetical protein CDS
KM573766 (modified) - ORF2 CDS translation - ORF2 CDS
KX388496.1 - hypothetical protein CDS translation - hypothetical protein CDS
KP153446 (modified) - Replication associated protein translation - Replication associated protein
MH545522.1 - Capsid protein translation - Replication associated protein
KM510191 (modified) - Capsid protein translation - hypothetical protein CDS
KU043413.1 - Replication associated protein translation - Replication associated protein
KT214389 (modified) - Capsid protein translation - V3 CDS
KP153511 (modified) - Replication associated protein translation - Replication associated protein
KP153512 (modified) - Replication associated protein translation - Replication associated protein
KP153513 (modified) - Replication associated protein translation - Replication associated protein
KP153514 (modified) - Replication associated protein translation - Replication associated protein
KP153515 (modified) - Replication associated protein translation - Replication associated protein
MH545525.1 - Replication associated protein translation - Replication associated protein
X84735 (modified) - TrAP translation - Replication-associated protein
X84735 (modified) - TrAP translation - TrAP
MG748715.1 - Capsid protein translation - hypothetical protein CDS
MG748717.1 - Capsid protein translation - hypothetical protein CDS
MG748718.1 - Capsid protein translation - hypothetical protein CDS
MG748719.1 - Capsid protein translation - hypothetical protein CDS
MG748720.1 - Capsid protein translation - hypothetical protein CDS
MG748716.1 - Capsid protein translation - hypothetical protein CDS
JF755407 (modified) - Replication associated protein interval 2 translation - Replication associated protein
JF755416 (modified) - Replication associated protein interval 2 translation - Replication associated protein
JF755417 (modified) - Replication associated protein interval 2 translation - Replication associated protein
KF147918 (modified) - Capsid protein translation - Movement protein
JX559642 (modified) - Capsid protein translation - Movement protein
KJ547647 (modified) - Replication associated protein translation - Replication associated protein
KR528563 (modified) - Replication associated protein translation - Replication associated protein
KP410285 (modified) - V2 CDS translation - V3 CDS
KP410285 (modified) - V2 CDS translation - V2 CDS
KJ206566 (modified) - Capsid protein translation - hypothetical protein CDS
KT214373 (modified) - Capsid protein translation - V3 CDS
KP153462 (modified) - hypothetical 2 translation - hypothetical 2
JQ920490 (modified) - Capsid protein translation - Movement protein
KP202150 (modified) - Capsid protein translation - Replication associated protein
KT149402 (modified) - hypothetical protein CDS translation - hypothetical protein CDS
EF536860 (modified) - Capsid protein translation - Movement protein
KJ206566 (modified) - hypothetical protein CDS translation 2 - Replication associated protein
KJ206566 (modified) - hypothetical protein CDS translation 2 - hypothetical protein CDS
KT214386 (modified) - Capsid protein translation - V3 CDS
KF481961 (modified) - nonstructural protein 2 CDS translation - nonstructural protein 2 CDS
MH545520.1 - Capsid protein translation - Replication associated protein
KJ547627 (modified) - Replication associated protein interval 1 translation - Replication associated protein
KX618694 - Pre-CP/V2 CDS translation - Pre-CP/V2 CDS
KT732823 (modified) - Capsid protein translation - Replication associated protein
KX388529.1 - hypothetical protein CDS translation 2 - hypothetical protein CDS
KX388528.1 - hypothetical protein CDS translation 2 - hypothetical protein CDS
KU043433.1 - Replication associated protein translation - Replication associated protein
JF755409 (modified) - ORF 5 (frame 3) translation - ORF 5 (frame 3)
KX388527.1 - hypothetical protein CDS translation - hypothetical protein CDS
KM821766 (modified) - Replication associated protein translation - Replication associated protein
KC248423 (modified) - Replication associated protein translation - Replication associated protein
KM821766 (modified) - hypothetical protein CDS translation - hypothetical protein CDS
KM573773 (modified) - ORF2 CDS translation - ORF2 CDS
KU043416.1 - Hypothetical protein translation 2 - Hypothetical protein
MH545535.1 - Capsid protein translation - Replication associated protein
JF713719 (modified) - unknown CDS translation 2 - unknown CDS
KP153492 (modified) - Replication associated protein translation - Replication associated protein
KP153490 (modified) - Replication associated protein translation - Replication associated protein
KP153491 (modified) - Replication associated protein translation - Replication associated protein
KP153493 (modified) - Replication associated protein translation - Replication associated protein
JX185417 (modified) - Replication associated protein translation - Replication associated protein
JX185416 (modified) - Replication associated protein translation - Replication associated protein
KF133821 - hypothetical protein CDS translation 2 - hypothetical protein CDS
KP153486 (modified) - Replication associated protein translation - Replication associated protein
KP153485 (modified) - Replication associated protein translation - Replication associated protein
KM874310 (modified) - Replication associated protein translation - Replication associated protein
KM874311 (modified) - Replication associated protein translation - Replication associated protein
KM874312 (modified) - Replication associated protein translation - Replication associated protein
KM874313 (modified) - Replication associated protein translation - Replication associated protein
KM874314 (modified) - Replication associated protein translation - Replication associated protein
KP153528 (modified) - Replication associated protein translation - Replication associated protein
KR902498 (modified) - hypothetical protein CDS translation 4 - hypothetical protein CDS
JF713719 (modified) - unknown CDS translation - unknown CDS
KU043416.1 - Hypothetical protein translation - Hypothetical protein
KM573773 (modified) - ORF3 CDS translation - ORF3 CDS
KU203351 (modified) - ORF 7 (frame 2) translation - ORF 6 (frame 3)
KU203351 (modified) - ORF 7 (frame 2) translation - ORF 7 (frame 2)
MH425570.1 - Replication associated protein translation - Replication associated protein
KJ547649 (modified) - hypothetical protein CDS translation - hypothetical protein CDS
KU043446.1 - Hypothetical protein translation - Hypothetical protein
FJ959082 (modified) - hypothetical protein CDS translation - hypothetical protein CDS
KC248417 (modified) - hypothetical protein CDS translation - hypothetical protein CDS
KP153361 (modified) - Replication associated protein translation - Replication associated protein
KP153362 (modified) - Replication associated protein translation - Replication associated protein
KY487779.1 - Replication associated protein translation - Replication associated protein
KR902498 (modified) - hypothetical protein CDS translation - hypothetical protein CDS
KF133812 - hypothetical protein CDS translation 2 - hypothetical protein CDS
WP_027090230 - other interval 3
WP_027090230 - other interval 3
WP_027090230 - other interval 3
WP_027090230 - other interval 3
YP_009237578 - other interval 3
YP_009237578 - other interval 3
YP_009237578 - other interval 3
KM874329 (modified) - Capsid protein translation - Replication associated protein
KM874330 (modified) - Capsid protein translation - Replication associated protein
KM874331 (modified) - Capsid protein translation - Replication associated protein
KX388498.1 - Capsid protein translation - misc
KX388498.1 - Capsid protein translation - misc
KX388500.1 - Capsid protein translation - Ori
KX388500.1 - Capsid protein translation - Ori
KX388501.1 - Capsid protein translation - Ori
KX388501.1 - Capsid protein translation - Ori
KF738883 (modified) - Replication associated protein interval 2 translation - Replication associated protein
KT149396 (modified) - Replication associated protein translation - hypothetical protein CDS
KU043409.1 - Replication associated protein translation - Hypothetical protein
KU043409.1 - Replication associated protein translation - Replication associated protein
KT149410 (modified) - Replication associated protein translation - hypothetical protein CDS
KT149409 (modified) - Replication associated protein translation - hypothetical protein CDS
AIF76278 - CDS
AIF76278 - Rep Protein
KM874332 (modified) - Capsid protein translation - Replication associated protein
KM874333 (modified) - Capsid protein translation - Replication associated protein
KM874334 (modified) - Capsid protein translation - Replication associated protein
KM874335 (modified) - Capsid protein translation - Replication associated protein
KT945165 (modified) - Replication associated protein interval 1 translation - Replication associated protein
KT945165 (modified) - Replication associated protein interval 1 translation - stem loop
KT945165 (modified) - Replication associated protein interval 1 translation - stem loop
APG55798 - CDS
APG55798 - Rep Protein
APG55798 - Viral Rep
APG55798 - Viral Rep
APG55798 - Viral Rep
MF118167.1 - hypothetical protein CDS translation - hypothetical protein CDS
KX388505.1 - Replication associated protein translation - misc
KX388505.1 - Replication associated protein translation - misc
KX388495.1 - Replication associated protein translation - Ori
KX388495.1 - Replication associated protein translation - Ori
KX388495.1 - Replication associated protein translation - Ori
KX388504.1 - Replication associated protein translation - Ori
KX388504.1 - Replication associated protein translation - Ori
KX388507.1 - Replication associated protein translation - Ori
KX388507.1 - Replication associated protein translation - Ori
KX388507.1 - Replication associated protein translation - Ori
MF118167.1 - hypothetical protein CDS translation - Ori
MF118167.1 - hypothetical protein CDS translation - Ori
KX388495.1 - Replication associated protein translation - Replication associated protein
KX388504.1 - Replication associated protein translation - Replication associated protein
KX388505.1 - Replication associated protein translation - Replication associated protein
KX388507.1 - Replication associated protein translation - Replication associated protein
KX388495.1 - Replication associated protein translation - stem loop
KX388507.1 - Replication associated protein translation - stem loop
KF738880 (modified) - Replication associated protein interval 2 translation - Replication associated protein
KT214386 (modified) - RepA CDS translation - Replication-associated protein interval 2

KT214373 (modified) - RepA CDS translation - Replication-associated protein interval 2
MH545530.1 - Replication associated protein interval 1 translation - Replication associated protein
MH545530.1 - Replication associated protein interval 1 translation - stem loop
MH545530.1 - Replication associated protein interval 1 translation - stem loop
KF738878 (modified) - Replication associated protein interval 2 translation - Replication associated protein
KF738879 (modified) - Replication associated protein interval 2 translation - Replication associated protein
FJ959086 (modified) - Replication associated protein translation - Replication associated protein
FJ959086 (modified) - Replication associated protein translation - stem loop
FJ959086 (modified) - Replication associated protein translation - stem loop
KM598411 (modified) - Replication associated protein interval 2 translation - Replication associated protein
KR528558 (modified) - Replication associated protein interval 2 translation - Replication associated protein
KT214389 (modified) - RepA CDS translation - Replication-associated protein interval 2
KM821748 (modified) - Replication associated protein interval 2 translation - Replication associated protein
KR528558 (modified) - Replication associated protein interval 2 translation - contains nonanucleotide motif stem loop
KR528558 (modified) - Replication associated protein interval 2 translation - Replication associated protein
KR528558 (modified) - Replication associated protein interval 2 translation - contains nonanucleotide motif stem loop
YP_009237578 - other interval 1
YP_009237578 - other interval 1
YP_009237578 - other interval 1
YP_009237578 - other interval 1
WP_027090230 - other interval 1
WP_027090230 - other interval 1
WP_027090230 - other interval 1
WP_027090230 - other interval 1
WP_027090230 - other interval 1
WP_027090230 - other interval 1
JX094280 (modified) - Replication-associated protein interval 2 translation - Replication-associated protein
KM598400 (modified) - Replication associated protein interval 2 translation - Replication associated protein
KT388086 (modified) - Capsid protein translation - Ren
KM821757 (modified) - Capsid protein translation - Replication associated protein
AXH73061 - CDS
AXH73061 - putative viral replication protein
AXH73061 - TIP49
AXH73061 - TIP49
AXH73061 - TIP49
KJ437671 (modified) - RepA CDS translation - Replication-associated protein interval 2
AQU11746 - CDS
AQU11746 - replication protein
AQU11746 - Viral Rep
AQU11746 - Viral Rep
AQU11746 - Viral Rep
KF133823 - Replication associated protein interval 2 translation - Replication associated protein
MH545540.1 - Replication associated protein interval 2 translation - Replication associated protein
YP_009237554 - Gemini AL1
YP_009237554 - Gemini AL1
YP_009109660 - replication-associated protein
YP_009109660 - start codon not determined CDS
YP_009109660 - Viral Rep
YP_009109660 - Viral Rep
YP_009109660 - Viral Rep
WP_027090230 - arginine finger
WP_027090230 - arginine finger
WP_027090230 - arginine finger
WP_027090230 - arginine finger
MH617615.1 - Replication associated protein translation - hypothetical protein CDS
MH617615.1 - Replication associated protein translation - Replication associated protein
AIF34798 - Viral Rep
AIF34798 - Viral Rep
AGA18393 - Viral Rep
AGA18393 - Viral Rep
AXH76451 - Viral Rep
AXH76451 - Viral Rep
YP_009226567 - Viral Rep
YP_009226567 - Viral Rep
AQU11736 - Viral Rep
AQU11736 - Viral Rep
KX388516.1 - Replication associated protein interval 2 translation - Replication associated protein
AGA18448 - Viral Rep
AGA18448 - Viral Rep
AQU11717 - Viral Rep
AQU11717 - Viral Rep
AQU11724 - Viral Rep
AQU11724 - Viral Rep
AXH76508 - Viral Rep
AXH76508 - Viral Rep
KM874350 (modified) - Replication associated protein interval 2 translation - Replication associated protein
AIF34812 - Gemini AL1
AIF34812 - Gemini AL1
AJP36430 - Viral Rep
AJP36430 - Viral Rep
AQU11726 - Viral Rep
AQU11726 - Viral Rep
YP_009237541 - AAA
YP_009237541 - AAA
YP_009448204 - Viral Rep
YP_009448204 - Viral Rep
YP_009126925 - Viral Rep
YP_009126925 - Viral Rep
YP_009115538 - Viral Rep
YP_009115538 - Viral Rep
AGA18286 - Viral Rep
AGA18286 - Viral Rep
KJ547626 (modified) - Replication associated protein interval 2 translation - Replication associated protein
KP153394 (modified) - Replication associated protein interval 2 translation - Replication associated protein
YP_009237541 - Viral Rep
YP_009237541 - Viral Rep
KY487837.1 - Replication associated protein translation - Replication associated protein
AKO84203 - Viral Rep
AKO84203 - Viral Rep
YP_009170674 - Viral Rep
YP_009170674 - Viral Rep
ADY62649 - Viral Rep
ADY62649 - Viral Rep
AHB63242 - Viral Rep
AHB63242 - Viral Rep
YP_009126938 - Viral Rep
YP_009126938 - Viral Rep
AXH76879 - Viral Rep
AXH76879 - Viral Rep
AIF76269 - Viral Rep
AIF76269 - Viral Rep
KF133815 - hypothetical protein CDS interval 2 translation - hypothetical protein CDS
KF133815 - hypothetical protein CDS interval 2 translation - stem loop
KF133815 - hypothetical protein CDS interval 2 translation - stem loop
KF133815 - hypothetical protein CDS interval 2 translation - hypothetical protein CDS
AXH77740 - Viral Rep
AXH77740 - Viral Rep
AVA16977 - Viral Rep
AVA16977 - Viral Rep
AVV68420 - Viral Rep
AVV68420 - Viral Rep
AVA17000 - Viral Rep
AVA17000 - Viral Rep
AVA16996 - Viral Rep
AVA16996 - Viral Rep
ATD53351 - Viral Rep
ATD53351 - Viral Rep
AVA16998 - Viral Rep
AVA16998 - Viral Rep
AWB80902 - Viral Rep
AWB80902 - Viral Rep
YP_008052687 - Viral Rep
YP_008052687 - Viral Rep
ADC79191 - Viral Rep
ADC79191 - Viral Rep
YP_009021888 - Viral Rep
YP_009021888 - Viral Rep
AUT13975 - Viral Rep
AUT13975 - Viral Rep
ACE62799 - Viral Rep
ACE62799 - Viral Rep
AIW81537 - Viral Rep
AIW81537 - Viral Rep
KR528560 (modified) - Replication associated protein interval 2 translation - Replication associated protein
MH378453.1 - Capsid protein translation - Replication associated protein
KT732784 (modified) - Capsid protein translation - hypothetical protein CDS
KJ955447 (modified) - Replication associated protein translation - viral protein RNase Z CDS
KJ955448 (modified) - Replication associated protein translation - viral protein RNase Z CDS
KJ955449 (modified) - Replication associated protein translation - viral protein RNase Z CDS
KJ955450 (modified) - Replication associated protein translation - viral protein RNase Z CDS
KJ955451 (modified) - Replication associated protein translation - viral protein RNase Z CDS
KR134313 (modified) - Replication associated protein translation - viral protein RNase Z CDS
KR134314 (modified) - Replication associated protein translation - viral protein RNase Z CDS
KR134315 (modified) - Replication associated protein translation - viral protein RNase Z CDS
KR134316 (modified) - Replication associated protein translation - viral protein RNase Z CDS
KR134317 (modified) - Replication associated protein translation - viral protein RNase Z CDS
KR134318 (modified) - Replication associated protein translation - viral protein RNase Z CDS
KR134319 (modified) - Replication associated protein translation - viral protein RNase Z CDS
KR134320 (modified) - Replication associated protein translation - viral protein RNase Z CDS
KR134323 (modified) - Replication associated protein translation - viral protein RNase Z CDS
KR134324 (modified) - Replication associated protein translation - viral protein RNase Z CDS
KR134325 (modified) - Replication associated protein translation - viral protein RNase Z CDS
KR134326 (modified) - Replication associated protein translation - viral protein RNase Z CDS
KR134327 (modified) - Replication associated protein translation - viral protein RNase Z CDS
KR134328 (modified) - Replication associated protein translation - viral protein RNase Z CDS
KR134329 (modified) - Replication associated protein translation - viral protein RNase Z CDS
KR134330 (modified) - Replication associated protein translation - viral protein RNase Z CDS
KR134331 (modified) - Replication associated protein translation - viral protein RNase Z CDS
KR134332 (modified) - Replication associated protein translation - viral protein RNase Z CDS
KR134333 (modified) - Replication associated protein translation - viral protein RNase Z CDS
KR134334 (modified) - Replication associated protein translation - viral protein RNase Z CDS
KR134335 (modified) - Replication associated protein translation - viral protein RNase Z CDS
KR134336 (modified) - Replication associated protein translation - viral protein RNase Z CDS
KR134337 (modified) - Replication associated protein translation - viral protein RNase Z CDS
KR134338 (modified) - Replication associated protein translation - viral protein RNase Z CDS
KR134339 (modified) - Replication associated protein translation - viral protein RNase Z CDS
KR134340 (modified) - Replication associated protein translation - viral protein RNase Z CDS
KR134341 (modified) - Replication associated protein translation - viral protein RNase Z CDS
KR134342 (modified) - Replication associated protein translation - viral protein RNase Z CDS
KR134343 (modified) - Replication associated protein translation - viral protein RNase Z CDS
KR134344 (modified) - Replication associated protein translation - viral protein RNase Z CDS
KR134345 (modified) - Replication associated protein translation - viral protein RNase Z CDS
KR134346 (modified) - Replication associated protein translation - viral protein RNase Z CDS
KR134347 (modified) - Replication associated protein translation - viral protein RNase Z CDS
KR134348 (modified) - Replication associated protein translation - viral protein RNase Z CDS
KR134349 (modified) - Replication associated protein translation - viral protein RNase Z CDS
KR134350 (modified) - Replication associated protein translation - viral protein RNase Z CDS
KR528549 (modified) - Capsid protein translation - Replication associated protein
EF536860 (modified) - RepA translation - Replication-associated protein interval 2
KJ547653 (modified) - hypothetical protein CDS translation - hypothetical protein CDS
AEL22996 - Viral Rep
AEL22996 - Viral Rep
YP_004778177 - Viral Rep
YP_004778177 - Viral Rep
AXH73508 - Viral Rep
AXH73508 - Viral Rep
JQ920490 (modified) - Replication-associated protein interval 2 translation - Replication-associated protein
AXH75487 - Viral Rep
AXH75487 - Viral Rep
AXH76667 - Viral Rep
AXH76667 - Viral Rep
AQU11728 - Viral Rep
AQU11728 - Viral Rep
KM821749 (modified) - Replication associated protein interval 2 translation - Replication associated protein
HQ443515 (modified) - Replication-associated protein interval 2 translation - Replication-associated protein
KR131749 (modified) - putative RepA-like protein CDS translation - Replication-associated protein interval 2
KU043398.1 - Replication associated protein translation - Replication associated protein
AXH74936 - Viral Rep
AXH74936 - Viral Rep
ARO38300 - Viral Rep
ARO38300 - Viral Rep
YP_009116906 - Viral Rep
YP_009116906 - Viral Rep
AXH73393 - Viral Rep
AXH73393 - Viral Rep
AXH76040 - Viral Rep
AXH76040 - Viral Rep
AIF76255 - Viral Rep
AIF76255 - Viral Rep
YP_009237516 - Viral Rep
YP_009237516 - Viral Rep
ALE29847 - Viral Rep
ALE29847 - Viral Rep
YP_009001747 - Viral Rep
YP_009001747 - Viral Rep
YP_009237564 - Viral Rep
YP_009237564 - Viral Rep
AJD07478 - Viral Rep
AJD07478 - Viral Rep
ALE29688 - Viral Rep
ALE29688 - Viral Rep
YP_009001742 - Viral Rep
YP_009001742 - Viral Rep
YP_009237586 - Viral Rep
YP_009237586 - Viral Rep
AMH87735 - Viral Rep
AMH87735 - Viral Rep
AXH73382 - Viral Rep
AXH73382 - Viral Rep
KF133823 - hypothetical protein CDS translation 2 - hypothetical protein CDS
ARE67375 - Viral Rep
ARE67375 - Viral Rep
NP_955176 - Viral Rep
NP_955176 - Viral Rep
AKR53286 - Viral Rep
AKR53286 - Viral Rep
AGA18388 - Viral Rep
AGA18388 - Viral Rep
YP_009126890 - Viral Rep
YP_009126890 - Viral Rep
AGA18473 - Viral Rep

AGA18473 - Viral Rep
AGS47835 - Viral Rep
AGS47835 - Viral Rep
YP_009109670 - Viral Rep
YP_009109670 - Viral Rep
AQR57898 - Viral Rep
AQR57898 - Viral Rep
AGA18245 - Viral Rep
AGA18245 - Viral Rep
AQR57902 - Viral Rep
AQR57902 - Viral Rep
AXH73061 - Viral Rep
AXH73061 - Viral Rep
Y00514 (modified) - Replication-associated protein interval 2 translation - RepA
Y00514 (modified) - Replication-associated protein interval 2 translation - Replication-associated protein
NP_619761 - Viral Rep
NP_619761 - Viral Rep
YP_009237578 - Viral Rep
YP_009237578 - Viral Rep
KF133827 - hypothetical protein CDS translation 3 - hypothetical protein CDS
AQU11729 - Viral Rep
AQU11729 - Viral Rep
KJ547621 (modified) - Capsid protein translation - Replication associated protein
KM821759 (modified) - Replication associated protein interval 2 translation - Replication associated protein
KM510189 (modified) - Replication associated protein interval 2 translation - Replication associated protein
KM510190 (modified) - Replication associated protein interval 2 translation - Replication associated protein
MH545531.1 - Replication associated protein interval 2 translation - Replication associated protein
MH545531.1 - Replication associated protein interval 2 translation - stem loop
MH545531.1 - Replication associated protein interval 2 translation - stem loop
MH545531.1 - Replication associated protein interval 2 translation - Replication associated protein
KR528569 (modified) - Replication associated protein interval 2 translation - Replication associated protein
DQ458791 (modified) - C1 CDS translation - Replication-associated protein interval 2
KU043409.1 - Hypothetical protein translation - Replication associated protein
MH545530.1 - Replication associated protein interval 2 translation - Replication associated protein
MH545530.1 - Replication associated protein interval 2 translation - stem loop
MH545530.1 - Replication associated protein interval 2 translation - stem loop
MH545530.1 - Replication associated protein interval 2 translation - Replication associated protein
FJ959083 (modified) - Replication associated protein interval 2 translation - hypothetical protein CDS
KR528560 (modified) - Capsid protein translation - Replication associated protein
KJ547627 (modified) - Replication associated protein interval 2 translation - Replication associated protein
AJD07493 - Viral Rep
AJD07493 - Viral Rep
KR528544 (modified) - Replication associated protein translation - Replication associated protein
AQU11749 - CDS
AQU11749 - replication protein
AQU11749 - Viral Rep
AQU11749 - Viral Rep
AQU11749 - Viral Rep
HQ335087 (modified) - Replication associated protein interval 2 translation - Replication associated protein
KX388495.1 - Replication associated protein translation - Replication associated protein
KX388495.1 - Replication associated protein translation - stem loop
KX388495.1 - Replication associated protein translation - stem loop
KM510191 (modified) - hypothetical protein CDS translation 2 - hypothetical protein CDS
KM510191 (modified) - Replication associated protein translation - hypothetical protein CDS
KM510191 (modified) - hypothetical protein CDS translation 2 - Replication associated protein
MH539648.1 - Capsid protein translation - hypothetical protein CDS
AQU11726 - P-loop NTPase
AQU11726 - P-loop NTPase
AVA16977 - RNA helicase
AVA16977 - RNA helicase
AVV68420 - RNA helicase
AVV68420 - RNA helicase
AVH76405 - Viral Rep
AVH76405 - Viral Rep
KR131749 (modified) - ORF1 CDS translation - putative movement protein CDS
YP_009163920 - P-loop NTPase
YP_009163920 - P-loop NTPase
ARD71303 - P-loop NTPase
ARD71303 - P-loop NTPase
AIF34818 - P-loop NTPase
AIF34818 - P-loop NTPase
AXH73508 - RNA helicase
AXH73508 - RNA helicase
AXG50856 - Viral Rep
AXG50856 - Viral Rep
AHC72271 - Viral Rep
AHC72271 - Viral Rep
YP_008997794 - Viral Rep
YP_008997794 - Viral Rep
AHC72167 - Viral Rep
AHC72167 - Viral Rep
AHC72177 - Viral Rep
AHC72177 - Viral Rep
WP_027090230 - AAA
WP_027090230 - P-loop NTPase
WP_027090230 - AAA
WP_027090230 - AAA
WP_027090230 - P-loop NTPase
WP_027090230 - P-loop NTPase
AXH76632 - P-loop NTPase
AXH76632 - P-loop NTPase
MG846357.1 - Replication associated protein interval 1 translation - Replication associated protein
ACE62799 - RNA helicase
ACE62799 - RNA helicase
AHB63242 - RNA helicase
AHB63242 - RNA helicase
ADY62649 - RNA helicase
ADY62649 - RNA helicase
ARE68406 - RNA helicase
ARE68406 - RNA helicase
YP_009109643 - CDS
YP_009109643 - HTH
YP_009109643 - replication-associated protein
YP_009109643 - HTH
YP_009109643 - HTH
MH545529.1 - Replication associated protein interval 2 translation - Replication associated protein
YP_004376332 - Viral Rep
YP_004376332 - Viral Rep
YP_009163936 - Viral Rep
YP_009163936 - Viral Rep
MH378453.1 - Replication associated protein interval 2 translation - Replication associated protein
KX388527.1 - Replication associated protein translation - Ori
KX388527.1 - Replication associated protein translation - Ori
KX388527.1 - Replication associated protein translation - Ori
KX388527.1 - Replication associated protein translation - Replication associated protein
KX388527.1 - Replication associated protein translation - stem loop
ATY42470 - RNA helicase
ATY42470 - RNA helicase
AIL50149 - RNA helicase
AIL50149 - RNA helicase
AVT56110 - RNA helicase
AVT56110 - RNA helicase
ADC79191 - P-loop NTPase
ADC79191 - P-loop NTPase
JF755404 (modified) - ORF 9 (frame 1) translation - ORF 8 (frame 3)
JF755405 (modified) - ORF 9 (frame 1) translation - ORF 8 (frame 3)
JF755406 (modified) - ORF 9 (frame 1) translation - ORF 8 (frame 3)
JF755404 (modified) - ORF 8 (frame 3) translation - ORF 9 (frame 1)
JF755404 (modified) - ORF 9 (frame 1) translation - ORF 9 (frame 1)
JF755405 (modified) - ORF 8 (frame 3) translation - ORF 9 (frame 1)
JF755405 (modified) - ORF 9 (frame 1) translation - ORF 9 (frame 1)
JF755406 (modified) - ORF 8 (frame 3) translation - ORF 9 (frame 1)
JF755406 (modified) - ORF 9 (frame 1) translation - ORF 9 (frame 1)
KP153476 (modified) - Replication associated protein interval 2 translation - Replication associated protein
KP153477 (modified) - Replication associated protein interval 2 translation - Replication associated protein
KP153478 (modified) - Replication associated protein interval 2 translation - Replication associated protein
KP153479 (modified) - Replication associated protein interval 2 translation - Replication associated protein
KP153480 (modified) - Replication associated protein interval 2 translation - Replication associated protein
KP153481 (modified) - Replication associated protein interval 2 translation - Replication associated protein
KP153482 (modified) - Replication associated protein interval 2 translation - Replication associated protein
AKO84203 - RNA helicase
AKO84203 - RNA helicase
YP_009237578 - Walker B motif
YP_009237578 - Walker B motif
YP_009237578 - Walker B motif
YP_009051960 - P-loop NTPase
YP_009051960 - P-loop NTPase
APG55798 - P-loop NTPase
APG55798 - P-loop NTPase
APZ87906 - RNA helicase
APZ87906 - RNA helicase
KT149412 (modified) - hypothetical protein CDS translation 2 - hypothetical protein CDS
KT149412 (modified) - Replication associated protein translation - hypothetical protein CDS
KT149412 (modified) - hypothetical protein CDS translation 2 - Replication associated protein
MG846357.1 - Replication associated protein interval 2 translation - Replication associated protein
KT149404 (modified) - hypothetical protein CDS translation 3 - hypothetical protein CDS
KT149404 (modified) - hypothetical protein CDS translation 4 - hypothetical protein CDS
KT149404 (modified) - hypothetical protein CDS translation 4 - hypothetical protein CDS
AFH02742 - Viral Rep
AFH02742 - Viral Rep
AEL28813 - Viral Rep
AEL28813 - Viral Rep
AXH74140 - Viral Rep
AXH74140 - Viral Rep
AVA16977 - replicase-associated protein
AVA16977 - start codon not determined CDS
AVV68420 - CDS
AVV68420 - replication-associated protein
AVA17000 - replicase-associated protein
AVA17000 - start codon not determined CDS
AVA16996 - replicase-associated protein
AVA16996 - start codon not determined CDS
ATD53351 - ORF1 CDS
ATD53351 - Rep Protein
KT869077.1 - Replication associated protein translation - Replication associated protein
KT869077.1 - Replication associated protein translation - Replication associated protein
AVA16998 - replicase-associated protein
AVA16998 - start codon not determined CDS
AWB80902 - Rep CDS
AWB80902 - replicase protein
YP_009458619 - Viral Rep
YP_009458619 - Viral Rep
BAP81877 - Viral Rep
BAP81877 - Viral Rep
APZ87906 - Viral Rep
APZ87906 - Viral Rep
AVT56110 - Viral Rep
AVT56110 - Viral Rep
AIL50149 - Viral Rep
AIL50149 - Viral Rep
ATY42470 - Viral Rep
ATY42470 - Viral Rep
MH545531.1 - Replication associated protein interval 1 translation - Replication associated protein
MH545531.1 - Replication associated protein interval 1 translation - stem loop
MH545531.1 - Replication associated protein interval 1 translation - stem loop
AKO71368 - Viral Rep
AKO71368 - Viral Rep
ALA65733 - Viral Rep
ALA65733 - Viral Rep
AIF76278 - RNA helicase
AIF76278 - RNA helicase
YP_009126925 - RNA helicase
YP_009126925 - RNA helicase
YP_009109643 - P-loop NTPase
YP_009109643 - P-loop NTPase
AXH74443 - RNA helicase
AXH74443 - RNA helicase
AQU11749 - RNA helicase
AQU11749 - RNA helicase
YP_009126932 - P-loop NTPase
YP_009126932 - P-loop NTPase
AXH73382 - P-loop NTPase
AXH73382 - P-loop NTPase
YP_009126898 - P-loop NTPase
YP_009126898 - P-loop NTPase
AWB80902 - P-loop NTPase
AWB80902 - P-loop NTPase
ATD53351 - RNA helicase
ATD53351 - RNA helicase
AVA16996 - RNA helicase
AVA16996 - RNA helicase
AVA16998 - RNA helicase
AVA16998 - RNA helicase
AVA17000 - RNA helicase
AVA17000 - RNA helicase
YP_004376332 - RNA helicase
YP_004376332 - RNA helicase
ATY70087 - Viral Rep
ATY70087 - Viral Rep
KF133815 - hypothetical protein CDS interval 2 translation - ORF 5 (frame 2)
YP_003084282 - Viral Rep
YP_003084282 - Viral Rep
AXH77564 - Viral Rep
AXH77564 - Viral Rep
CBK25810 - Viral Rep
CBK25810 - Viral Rep
AQU11776 - Viral Rep
AQU11776 - Viral Rep
AQU11773 - Viral Rep
AQU11773 - Viral Rep
AEI54346 - Viral Rep
AEI54346 - Viral Rep
AGA18409 - Viral Rep
AGA18409 - Viral Rep
AXH75991 - Viral Rep
AXH75991 - Viral Rep
YP_009163927 - Viral Rep
YP_009163927 - Viral Rep
AEM05804 - Viral Rep
AEM05804 - Viral Rep
AXH74683 - Viral Rep
AXH74683 - Viral Rep
KT149406 (modified) - hypothetical protein CDS translation 2 - Replication associated protein
FJ959077 (modified) - hypothetical protein CDS translation 2 - hypothetical protein CDS
AXH76451 - RNA helicase
AXH76451 - RNA helicase
AGA18245 - RNA helicase
AGA18245 - RNA helicase
YP_009021888 - P-loop NTPase
YP_009021888 - P-loop NTPase
ALE29635 - RNA helicase
ALE29635 - RNA helicase
YP_009237530 - RNA helicase
YP_009237530 - RNA helicase
YP_009115538 - P-loop NTPase
YP_009115538 - P-loop NTPase
AWW06057 - RNA helicase
AWW06057 - RNA helicase
AGA18441 - P-loop NTPase

AGA18441 - P-loop NTPase
YP_009116906 - P-loop NTPase
YP_009116906 - P-loop NTPase
AGA18448 - RNA helicase
AGA18448 - RNA helicase
AMH87735 - P-loop NTPase
AMH87735 - P-loop NTPase
YP_009126890 - RNA helicase
YP_009126890 - RNA helicase
AXH76508 - RNA helicase
AXH76508 - RNA helicase
AXH73061 - RNA helicase
AXH73061 - RNA helicase
AXH73061 - RNA helicase
YP_009163927 - P-loop NTPase
YP_009163927 - P-loop NTPase
AXH75991 - P-loop NTPase
AXH75991 - P-loop NTPase
AQR57898 - P-loop NTPase
AQR57898 - P-loop NTPase
ARO38300 - P-loop NTPase
ARO38300 - P-loop NTPase
AIF76269 - P-loop NTPase
AIF76269 - P-loop NTPase
YP_009126879 - P-loop NTPase
YP_009126879 - P-loop NTPase
YP_009116902 - P-loop NTPase
YP_009116902 - P-loop NTPase
YP_009109675 - P-loop NTPase
YP_009109675 - P-loop NTPase
AIF34802 - P-loop NTPase
AIF34802 - P-loop NTPase
AJD20393 - Gemini AL1 M
AJD20393 - Gemini AL1 M
KY487844.1 - Replication associated protein translation - Replication associated protein
YP_006281010 - P-loop NTPase
YP_006281010 - P-loop NTPase
KR528559 (modified) - Capsid protein translation - Replication associated protein
AVH76405 - RNA helicase
AVH76405 - RNA helicase
AVH76405 - RNA helicase
AXH76879 - P-loop NTPase
AXH76879 - P-loop NTPase
ALE29847 - RNA helicase
ALE29847 - RNA helicase
YP_009001747 - RNA helicase
YP_009001747 - RNA helicase
AQU11773 - P-loop NTPase
AQU11773 - P-loop NTPase
YP_009021245 - RNA helicase
YP_009021245 - RNA helicase
AXH78100 - RNA helicase
AXH78100 - RNA helicase
AQU11776 - P-loop NTPase
AQU11776 - P-loop NTPase
KT214386 (modified) - V2 CDS translation - V3 CDS
AXH75780 - Viral Rep
AXH75780 - Viral Rep
KP005454 (modified) - Capsid protein translation - hypothetical protein CDS
KP005454 (modified) - hypothetical protein CDS translation - hypothetical protein CDS
KP005453 (modified) - Capsid protein translation - hypothetical protein CDS
KP005453 (modified) - hypothetical protein CDS translation - hypothetical protein CDS
AIF34818 - Viral Rep
AIF34818 - Viral Rep
KM510189 (modified) - hypothetical protein CDS translation - hypothetical protein CDS
KM510190 (modified) - hypothetical protein CDS translation - hypothetical protein CDS
KM510189 (modified) - Capsid protein translation - hypothetical protein CDS
KM510189 (modified) - hypothetical protein CDS translation - hypothetical protein CDS
KM510190 (modified) - Capsid protein translation - hypothetical protein CDS
KM510190 (modified) - hypothetical protein CDS translation - hypothetical protein CDS
AIX11626 - RNA helicase
AIX11626 - RNA helicase
AOV86234 - RNA helicase
AOV86234 - RNA helicase
AOV86234 - RNA helicase
KJ547647 (modified) - hypothetical protein CDS translation - hypothetical protein CDS
KJ547647 (modified) - Replication associated protein translation - hypothetical protein CDS
KJ547647 (modified) - hypothetical protein CDS translation - Replication associated protein
KT149400 (modified) - Replication associated protein translation - hypothetical protein CDS
KJ547631 (modified) - hypothetical protein CDS translation - Replication associated protein
APG55798 - CDS
APG55798 - GluZincin
APG55798 - Rep Protein
APG55798 - GluZincin
APG55798 - GluZincin
YP_009237586 - P-loop NTPase
YP_009237586 - P-loop NTPase
AXH73393 - RNA helicase
AXH73393 - RNA helicase
AXH76040 - RNA helicase
AXH76040 - RNA helicase
KT862236 (modified) - Capsid protein translation - Replication associated protein
AVX29443 - RNA helicase
AVX29443 - RNA helicase
AXH73290 - Viral Rep
AXH73290 - Viral Rep
AXH75585 - Viral Rep
AXH75585 - Viral Rep
AGS47835 - P-loop NTPase
AGS47835 - P-loop NTPase
KU043420.1 - Replication associated protein translation - Hypothetical protein
AXH73792 - Viral Rep
AXH73792 - Viral Rep
AIF34798 - P-loop NTPase
AIF34798 - P-loop NTPase
AEL22996 - P-loop NTPase
AEL22996 - P-loop NTPase
YP_004778177 - P-loop NTPase
YP_004778177 - P-loop NTPase
AJD07493 - RNA helicase
AJD07493 - RNA helicase
AGA18388 - P-loop NTPase
AGA18388 - P-loop NTPase
AUW34331 - RNA helicase
AUW34331 - RNA helicase
YP_009163936 - P-loop NTPase
YP_009163936 - P-loop NTPase
AGA18473 - P-loop NTPase
AGA18473 - P-loop NTPase
AJM89742 - P-loop NTPase
AJM89742 - P-loop NTPase
YP_009237516 - RNA helicase
YP_009237516 - RNA helicase
AXH77740 - P-loop NTPase
AXH77740 - P-loop NTPase
KF133827 - hypothetical protein CDS translation - hypothetical protein CDS
KM874309 (modified) - Capsid protein translation - Replication associated protein
KJ547632 (modified) - hypothetical protein CDS translation - hypothetical protein CDS
ARE68406 - CDS
ARE68406 - replication associated protein
ARE68406 - Viral Rep
ARE68406 - Viral Rep
ARE68406 - Viral Rep
MH617615.1 - Capsid protein translation - CHAP domain protein CDS
KU043420.1 - Hypothetical protein translation 2 - Hypothetical protein
MH617135.1 - Replication associated protein translation - hypothetical protein CDS
MH617135.1 - Replication associated protein translation - Replication associated protein
JF713716 (modified) - Replication associated protein translation - unknown CDS
AXH74140 - RNA helicase
AXH74140 - RNA helicase
AIF76278 - CDS
AIF76278 - Rep Protein
AIF76278 - Viral Rep
AIF76278 - Viral Rep
AIF76278 - Viral Rep
BAP81877 - RNA helicase
BAP81877 - RNA helicase
YP_009109660 - P-loop NTPase
YP_009109660 - P-loop NTPase
JX908740 (modified) - Capsid protein translation - Replication associated protein
AJD07478 - RNA helicase
AJD07478 - RNA helicase
AXH75674 - P-loop NTPase
AXH75674 - P-loop NTPase
YP_009109670 - P-loop NTPase
YP_009109670 - P-loop NTPase
KF133814 - Replication associated protein interval 2 translation - Replication associated protein
AIF76259 - RNA helicase
AIF76259 - RNA helicase
KF133828 - Capsid protein translation - hypothetical protein CDS
AMD39533 - AAA 16
AMD39533 - AAA 16
YP_764455 - AAA 16
YP_764455 - AAA 16
KT732825 (modified) - Replication associated protein interval 2 translation - Replication associated protein
YP_009237530 - CDS
YP_009237530 - replication associated protein
YP_009237530 - Viral Rep
YP_009237530 - Viral Rep
YP_009237530 - Viral Rep
ALE29635 - CDS
ALE29635 - replication associated protein
ALE29635 - Viral Rep
ALE29635 - Viral Rep
ALE29635 - Viral Rep
AGA19549 - Gemini AL1
AGA19549 - Gemini AL1
FJ959083 (modified) - Replication associated protein interval 2 translation - Replication associated protein
AXH76632 - CDS
AXH76632 - helicase Protein
AXH76632 - Viral Rep
AXH76632 - Viral Rep
AXH76632 - Viral Rep
YP_009458619 - RNA helicase
YP_009458619 - RNA helicase
YP_009021245 - CDS
YP_009021245 - replication-associated protein
YP_009021245 - Viral Rep
AOV86234 - CDS
AOV86234 - putative rep protein
AOV86234 - Viral Rep
AXH73056 - CDS
AXH73056 - putative viral replication protein
AXH73056 - Viral Rep
YP_009109643 - CDS
YP_009109643 - replication-associated protein
YP_009109643 - Viral Rep
ARI44308 - Rep CDS
ARI44308 - replication-associated protein
ARI44308 - Viral Rep
AHH31400 - CDS
AHH31400 - replication-associated protein
AHH31400 - Viral Rep
AEL28791 - CDS
AEL28791 - replication-associated protein
AEL28791 - Viral Rep
YP_764455 - ORFV1; putative replicase CDS
YP_764455 - rep protein
YP_764455 - Viral Rep
AIF34812 - CDS
AIF34812 - Gemini AL1 M
AIF34812 - replication-associated protein
YP_009116902 - CDS
YP_009116902 - replication-associated protein
YP_009116902 - Viral Rep
AEL28793 - CDS
AEL28793 - replication-associated protein
AEL28793 - Viral Rep
AWW06057 - CDS
AWW06057 - helicase Protein
AWW06057 - Viral Rep
ALE29827 - CDS
ALE29827 - replication associated protein
ALE29827 - Viral Rep
AMD39533 - replication-associated protein
AMD39533 - V1 CDS
AMD39533 - Viral Rep
AVX29443 - CDS
AVX29443 - replication initiator protein
AVX29443 - Viral Rep
AXH74443 - CDS
AXH74443 - putative viral replication protein
AXH74443 - Viral Rep
AGA18441 - CDS
AGA18441 - hypothetical protein
AGA18441 - Viral Rep
AIF76259 - CDS
AIF76259 - Rep Protein
AIF76259 - Viral Rep
YP_009163920 - CDS
YP_009163920 - putative spliced replication initiation protein
YP_009163920 - Viral Rep
AIF34802 - CDS
AIF34802 - replication-associated protein
AIF34802 - Viral Rep
AIF76277 - CDS
AIF76277 - Rep Protein
AIF76277 - Viral Rep
KF738877 (modified) - Replication associated protein interval 2 translation - Replication associated protein
YP_009126898 - CDS
YP_009126898 - replication-associated protein
YP_009126898 - Viral Rep
AGA18265 - CDS
AGA18265 - hypothetical protein
AGA18265 - Viral Rep
AIX11626 - Rep CDS
AIX11626 - replicase Protein
AIX11626 - Viral Rep
AUW34331 - Rep CDS
AUW34331 - replication-associated protein
AUW34331 - Viral Rep
YP_009109675 - CDS
YP_009109675 - replication-associated protein
YP_009109675 - Viral Rep
AGG39817 - CDS
AGG39817 - replication-associated protein
AGG39817 - Viral Rep
AJM89742 - CDS
AJM89742 - replication associated protein
AJM89742 - Viral Rep
WP_027090230 - other interval 2
YP_009237578 - other interval 2
WP_027090230 - Walker B motif
YP_009237578 - Walker B motif
AAK73450 - C1/C2 CDS
AAK73450 - Gemini AL1 M
AAK73450 - Rep Protein
YP_009126879 - CDS
YP_009126879 - replication-associated protein
YP_009126879 - Viral Rep
YP_009142778 - Gemini AL1
YP_009142778 - putative replication protein
YP_009142778 - rep CDS
YP_009237504 - CDS
YP_009237504 - Gemini AL1
YP_009237504 - replication associated protein
AIF34803 - CDS
AIF34803 - Gemini AL1
AIF34803 - replication-associated protein
YP_009237555 - CDS
YP_009237555 - Gemini AL1
YP_009237555 - replication associated protein
AXH77952 - Gemini AL1
AXH77952 - Geminivirus Rep catalytic domain CDS
AXH77952 - Rep catalytic domain protein
AXH74707 - CDS
AXH74707 - Gemini AL1
AXH74707 - Rep Protein
AAK73450 - C1/C2 CDS
AAK73450 - Gemini AL1
AAK73450 - Rep Protein
AXH78100 - CDS
AXH78100 - helicase Protein
AXH78100 - Viral Rep
AJD07498 - CDS

AJD07498 - replication-associated protein
AJD07498 - Viral Rep
AXH73056 - CDS
AXH73056 - P-loop NTPase
AXH73056 - putative viral replication protein
ARE68406 - CDS
ARE68406 - replication associated protein
YP_009237554 - Gemini AL1
YP_009237554 - CDS
YP_009237554 - Gemini AL1 M
YP_009237554 - replication associated protein
AMD39533 - AAA 16
AMD39533 - AAA 16
AMD39533 - replication-associated protein
AMD39533 - replication-associated protein
AMD39533 - RNA helicase
AMD39533 - RNA helicase
AMD39533 - V1 CDS
AMD39533 - V1 CDS
AMD39533 - Walker A/P-loop
AMD39533 - Walker A/P-loop
AMD39533 - Walker A/P-loop
AMD39533 - Walker A/P-loop
AIF76255 - CDS
AIF76255 - P-loop NTPase
AIF76255 - Rep Protein
AIF76255 - Walker A/P-loop
YP_009237578 - CDS
YP_009237578 - replication associated protein
YP_009237578 - RNA helicase
YP_009237578 - Walker A motif
WP_027090230 - AAA
WP_027090230 - ATP-dependent Clp protease ATP-binding subunit Protein
WP_027090230 - P-loop NTPase
WP_027090230 - P-loop NTPase
WP_027090230 - Walker A motif
AGA19549 - C1 CDS
AGA19549 - Gemini AL1 M
AGA19549 - Rep Protein
AKR53286 - CDS
AKR53286 - P-loop NTPase
AKR53286 - viral replicase protein
KX388507.1 - Replication associated protein translation - Replication associated protein
KX388507.1 - Replication associated protein translation - stem loop
JX559621 (modified) - hypothetical protein CDS translation 2 - hypothetical protein CDS
JX559622 (modified) - hypothetical protein CDS translation 2 - hypothetical protein CDS
JX559621 (modified) - hypothetical protein CDS translation 2 - stem loop
JX559622 (modified) - hypothetical protein CDS translation 2 - stem loop
AJD20393 - CDS
AJD20393 - Gemini AL1
AJD20393 - replication associated protein
AGG39817 - CDS
AGG39817 - P-loop NTPase
AGG39817 - replication-associated protein
WP_027090230 - ATP-dependent Clp protease ATP-binding subunit Protein
WP_027090230 - P-loop NTPase
WP_027090230 - AAA
WP_027090230 - ATP-dependent Clp protease ATP-binding subunit Protein
WP_027090230 - P-loop NTPase
WP_027090230 - Walker B motif
AQR57902 - TIP49
AQR57902 - Rep CDS
AQR57902 - replicase Protein
AQR57902 - RNA helicase
AXH73290 - CDS
AXH73290 - P-loop NTPase
AXH73290 - putative viral replication protein
YP_009237541 - AAA
YP_009237541 - CDS
YP_009237541 - P-loop NTPase
YP_009237541 - replication associated protein
AUF34964 - CDS
AUF34964 - P-loop NTPase
AUF34964 - putative replication-associated protein
YP_009126892 - CDS
YP_009126892 - replication-associated protein
YP_009126892 - RNA helicase
AQU11729 - CDS
AQU11729 - P-loop NTPase
AQU11729 - replication protein
AHH31400 - CDS
ARI44308 - Rep CDS
AHH31400 - replication-associated protein
ARI44308 - replication-associated protein
AHH31400 - RNA helicase
ARI44308 - RNA helicase
AMD39533 - AAA 16
AMD39533 - replication-associated protein
AMD39533 - RNA helicase
AMD39533 - V1 CDS
AUT13975 - CDS
AUT13975 - replication protein
AUT13975 - RNA helicase
YP_009170674 - rep CDS
YP_009170674 - replicase Protein
YP_009170674 - RNA helicase
YP_009237578 - CDS
YP_009237578 - replication associated protein
YP_009237578 - RNA helicase
YP_764455 - AAA 16
YP_764455 - ORFV1; putative replicase CDS
YP_764455 - rep protein
YP_764455 - RNA helicase
AGA18409 - CDS
AGA18409 - hypothetical protein
AGA18409 - P-loop NTPase
NP_955176 - CDS
NP_955176 - Rep-like protein
ARE67375 - RNA helicase
NP_955176 - RNA helicase
ARE67375 - SWPV2-147 CDS
ARE67375 - SWPV2-ORF147 Protein
AEL28813 - CDS
AEL28813 - replication-associated protein
AEL28813 - RNA helicase
AFH02742 - CDS
AFH02742 - P-loop NTPase
AFH02742 - putative Rep Protein
AQU11733 - CDS
AQU11733 - P-loop NTPase
AQU11733 - replication protein
AIF76255 - CDS
AIF76255 - P-loop NTPase
AIF76255 - Rep Protein
AXH77121 - CDS
AXH77121 - helicase Protein
AXH77121 - RNA helicase
AJP36430 - CDS
AJP36430 - replication-associated protein
AJP36430 - RNA helicase
YP_009126938 - CDS
YP_009126938 - P-loop NTPase
YP_009126938 - replication-associated protein
AXG50856 - M-Rep CDS
AXG50856 - master replication initiator protein
AXG50856 - P-loop NTPase
NP_619761 - putative CDS
NP_619761 - RNA helicase
NP_619761 - virus replication-associated protein
AKO71368 - CDS
ALA65733 - CDS
AKO71368 - Replication associated protein
ALA65733 - replication initiation protein
AKO71368 - RNA helicase
ALA65733 - RNA helicase
AHC72271 - M-Rep CDS
AHC72271 - master replication initiator protein
AHC72271 - RNA helicase
YP_008997794 - M-Rep CDS
YP_008997794 - master replication initiator protein
YP_008997794 - RNA helicase
AHC72177 - M-Rep CDS
AHC72177 - master replication initiator protein
AHC72177 - RNA helicase
AHC72167 - M-Rep CDS
AHC72167 - master replication initiator protein
AHC72167 - P-loop NTPase
CBK25810 - rep CDS
CBK25810 - replication association protein
CBK25810 - RNA helicase
ATY70087 - Rep CDS
ATY70087 - replication initiator protein
ATY70087 - RNA helicase
ADC79191 - P-loop NTPase
ADC79191 - CDS
ADC79191 - RNA helicase
ADC79191 - V1 Protein
AGA18391 - CDS
AGA18391 - hypothetical protein
AGA18391 - P-loop NTPase
AWR89667 - CDS
AWR89667 - P-loop NTPase
AWR89667 - replication initiation protein
AQU11717 - CDS
AQU11724 - CDS
AQU11717 - P-loop NTPase
AQU11724 - P-loop NTPase
AQU11717 - replication protein
AQU11724 - replication protein
AEL87784 - CDS
AEL87784 - P-loop NTPase
AEL87784 - putative replication-associated protein
YP_009109660 - replication-associated protein
YP_009109660 - start codon not determined CDS
KX388528.1 - hypothetical protein CDS translation - hypothetical protein CDS
KX388528.1 - hypothetical protein CDS translation - stem loop
KY487868.1 - Replication associated protein translation - Replication associated protein
YP_009237571 - CDS
YP_009237571 - P-loop NTPase
YP_009237571 - replication associated protein
AWR89667 - CDS
AWR89667 - replication initiation protein
KY487956.1 - Replication associated protein translation - Replication associated protein
YP_003084282 - CDS
YP_003084282 - putative Rep protein
YP_003084282 - RNA helicase
AOV86234 - CDS
AOV86234 - DNA pol3 delta2
AOV86234 - putative rep protein
AIW81537 - ORF1 CDS
AIW81537 - replicase protein
AIW81537 - RNA helicase
AQU11726 - P-loop NTPase
AQU11726 - CDS
AQU11726 - P-loop NTPase
AQU11726 - replication protein
AGA18265 - CDS
AGA18265 - hypothetical protein
AGA18265 - P-loop NTPase
AJD07498 - CDS
AJD07498 - replication-associated protein
AJD07498 - RNA helicase
AXH73792 - CDS
AXH73792 - P-loop NTPase
AXH73792 - putative viral replication protein
YP_009237564 - CDS
YP_009237564 - P-loop NTPase
YP_009237564 - replication associated protein
ALE29688 - CDS
ALE29688 - replication associated protein
ALE29688 - RNA helicase
YP_009001742 - CDS
YP_009001742 - P-loop NTPase
YP_009001742 - replication-associated protein
YP_009448204 - rep CDS
YP_009448204 - RNA helicase
YP_009448204 - rolling-circle replication protein
AGA18286 - CDS
AGA18286 - hypothetical protein
AGA18286 - P-loop NTPase
AQR57898 - Rep CDS
AQR57898 - Rep CDS
AQR57898 - replicase Protein
AQR57898 - replicase Protein
AEL22996 - CDS
AEL22996 - CDS
YP_004778177 - CDS
YP_004778177 - CDS
AEL22996 - Rep protein
AEL22996 - Rep protein
YP_004778177 - Rep protein
YP_004778177 - Rep protein
AGA18286 - CDS
AGA18286 - CDS
AGA18286 - hypothetical protein
AGA18286 - hypothetical protein
JX904185 (modified) - Replication associated protein translation - Replication associated protein
YP_009163936 - CDS
YP_009163936 - CDS
YP_009163936 - putative replication initiation protein
YP_009163936 - putative replication initiation protein
JX904077 (modified) - Replication associated protein translation - Replication associated protein
APA62649 - CDS
APA62649 - CDS
APA62649 - putative replication protein
APA62649 - putative replication protein
AGA18409 - CDS
AGA18409 - CDS
AGA18409 - hypothetical protein
AGA18409 - hypothetical protein
JX904473 (modified) - Replication associated protein translation - Replication associated protein
KT149398 (modified) - hypothetical protein CDS translation - hypothetical protein CDS
AIF76259 - CDS
AIF76259 - CDS
AIF76259 - Rep Protein
AIF76259 - Rep Protein
ALE29847 - CDS
ALE29847 - replication associated protein
ALE29847 - CDS
ALE29847 - replication associated protein
YP_009001747 - CDS
YP_009001747 - CDS
YP_009001747 - replication-associated protein
YP_009001747 - replication-associated protein
KY348843.1 - Replication associated protein translation - Replication associated protein
KP153359 (modified) - Replication associated protein translation - Replication associated protein
AVX29443 - CDS
AVX29443 - CDS
AVX29443 - replication initiator protein
AVX29443 - replication initiator protein
ALE29635 - CDS
ALE29635 - CDS
ALE29635 - replication associated protein
ALE29635 - replication associated protein
AUT13975 - CDS
AUT13975 - CDS
AUT13975 - replication protein
AUT13975 - replication protein
AIF34802 - CDS
AIF34802 - CDS
AIF34802 - replication-associated protein
AIF34802 - replication-associated protein
AXL65935 - CDS
AXL65935 - CDS
AXL65935 - replication-associated protein
AXL65935 - replication-associated protein
AUM61711 - Rep CDS
AUM61711 - Rep CDS
AUM61711 - Rep Protein
AUM61711 - Rep Protein
AXH76667 - CDS
AXH76667 - CDS
AXH76667 - putative viral replication protein
AXH76667 - putative viral replication protein
JX904469 (modified) - Replication associated protein translation - Replication associated protein
AQU11726 - CDS
AQU11726 - CDS
AQU11726 - replication protein
AQU11726 - replication protein
BAP81877 - Rep CDS
BAP81877 - Rep CDS
BAP81877 - rolling circle replication initiator protein
BAP81877 - rolling circle replication initiator protein
ALE29688 - CDS
ALE29688 - CDS
ALE29688 - replication associated protein
ALE29688 - replication associated protein
YP_009001742 - CDS
YP_009001742 - CDS

YP_009001742 - replication-associated protein
YP_009001742 - replication-associated protein
YP_009458619 - rep CDS
YP_009458619 - rep CDS
YP_009458619 - replication-association protein
YP_009458619 - replication-association protein
YP_009237530 - CDS
YP_009237530 - CDS
YP_009237530 - replication associated protein
YP_009237530 - replication associated protein
ARI44308 - Rep CDS
ARI44308 - Rep CDS
ARI44308 - replication-associated protein
ARI44308 - replication-associated protein
AHH31400 - CDS
AHH31400 - CDS
AHH31400 - replication-associated protein
AHH31400 - replication-associated protein
YP_009506303 - CDS
YP_009506303 - CDS
YP_009506303 - Rep protein
YP_009506303 - Rep protein
AKR53286 - CDS
AKR53286 - CDS
AKR53286 - viral replicase protein
AKR53286 - viral replicase protein
ADC79191 - CDS
ADC79191 - CDS
ADC79191 - V1 Protein
ADC79191 - V1 Protein
AOV86234 - CDS
AOV86234 - CDS
AOV86234 - putative rep protein
AOV86234 - putative rep protein
AKO84203 - CDS
AKO84203 - CDS
AKO84203 - replicase Protein
AKO84203 - replicase Protein
AVT56110 - CDS
AVT56110 - CDS
AVT56110 - replicase Protein
AVT56110 - replicase Protein
AIL50149 - CDS
AIL50149 - CDS
AIL50149 - replicase Protein
AIL50149 - replicase Protein
ATY42470 - CDS
ATY42470 - CDS
ATY42470 - putative replication associated protein
ATY42470 - putative replication associated protein
KC241982 (modified) - Replication associated protein translation - Replication associated protein
KC241982 (modified) - Replication associated protein translation - Replication associated protein
AXR86424 - CDS
AXR86424 - CDS
AXR86424 - replicase Protein
AXR86424 - replicase Protein
AIF34812 - CDS
AIF34812 - CDS
AIF34812 - replication-associated protein
AIF34812 - replication-associated protein
AXG50856 - M-Rep CDS
AXG50856 - M-Rep CDS
AXG50856 - master replication initiator protein
AXG50856 - master replication initiator protein
AHC72167 - M-Rep CDS
AHC72167 - M-Rep CDS
AHC72167 - master replication initiator protein
AHC72167 - master replication initiator protein
AHC72177 - M-Rep CDS
AHC72177 - M-Rep CDS
AHC72177 - master replication initiator protein
AHC72177 - master replication initiator protein
AIF34818 - CDS
AIF34818 - CDS
AIF34818 - replication-associated protein
AIF34818 - replication-associated protein
JX904605 (modified) - Replication associated protein translation - Replication associated protein
AKO71368 - CDS
AKO71368 - CDS
AKO71368 - Replication associated protein
AKO71368 - Replication associated protein
ALA65733 - CDS
ALA65733 - CDS
ALA65733 - replication initiation protein
ALA65733 - replication initiation protein
AGA18393 - CDS
AGA18393 - CDS
AGA18393 - hypothetical protein
AGA18393 - hypothetical protein
JX904431 (modified) - Replication associated protein translation - Replication associated protein
JF755404 (modified) - Replication associated protein translation - Replication associated protein
JF755405 (modified) - Replication associated protein translation - Replication associated protein
JF755406 (modified) - Replication associated protein translation - Replication associated protein
AGA18441 - CDS
AGA18441 - CDS
AGA18441 - hypothetical protein
AGA18441 - hypothetical protein
JX904562 (modified) - Replication associated protein translation - Replication associated protein
AAK73450 - C1/C2 CDS
AAK73450 - C1/C2 CDS
AAK73450 - Rep Protein
AAK73450 - Rep Protein
AXL65907 - CDS
AXL65907 - CDS
AXL65907 - replication-associated protein
AXL65907 - replication-associated protein
JX904518 (modified) - Replication associated protein translation - Replication associated protein
AXH74936 - CDS
AXH74936 - CDS
AXH74936 - putative viral replication protein
AXH74936 - putative viral replication protein
AUM62033 - Rep CDS
AUM62033 - Rep CDS
AUM62033 - Rep Protein
AUM62033 - Rep Protein
AXH74683 - CDS
AXH74683 - CDS
AXH74683 - putative viral replication protein
AXH74683 - putative viral replication protein
YP_009506293 - CDS
YP_009506293 - CDS
YP_009506293 - Rep Protein
YP_009506293 - Rep Protein
YP_009047144 - CDS
YP_009047144 - CDS
YP_009047144 - replication-associated protein
YP_009047144 - replication-associated protein
KJ547653 (modified) - Replication associated protein translation - Replication associated protein
AOV86303 - CDS
AOV86303 - CDS
AOV86303 - putative rep protein
AOV86303 - putative rep protein
ARE67375 - SWPV2-147 CDS
ARE67375 - SWPV2-147 CDS
ARE67375 - SWPV2-ORF147 Protein
ARE67375 - SWPV2-ORF147 Protein
NP_955176 - CDS
NP_955176 - CDS
NP_955176 - Rep-like protein
NP_955176 - Rep-like protein
JF755411 (modified) - Replication associated protein translation - Replication associated protein
JF755412 (modified) - Replication associated protein translation - Replication associated protein
JF755413 (modified) - Replication associated protein translation - Replication associated protein
JF755414 (modified) - Replication associated protein translation - Replication associated protein
AQU11733 - CDS
AQU11733 - CDS
AQU11733 - replication protein
AQU11733 - replication protein
AXH76451 - CDS
AXH76451 - CDS
AXH76451 - helicase Protein
AXH76451 - helicase Protein
JX904076 (modified) - Replication associated protein translation - Replication associated protein
AEI54346 - CDS
AEI54346 - CDS
AEI54346 - rep protein
AEI54346 - rep protein
KP153430 (modified) - Replication associated protein translation - Replication associated protein
KP153431 (modified) - Replication associated protein translation - Replication associated protein
KP153432 (modified) - Replication associated protein translation - Replication associated protein
KP153433 (modified) - Replication associated protein translation - Replication associated protein
KP153429 (modified) - Replication associated protein translation - Replication associated protein
AGA18265 - CDS
AGA18265 - CDS
AGA18265 - hypothetical protein
AGA18265 - hypothetical protein
JX904144 (modified) - Replication associated protein translation - Replication associated protein
AUM61703 - Rep CDS
AUM61703 - Rep CDS
AUM61703 - Rep Protein
AUM61703 - Rep Protein
AVH76405 - CDS
AVH76405 - CDS
AVH76405 - putative Rep Protein
AVH76405 - putative Rep Protein
AIF34803 - CDS
AIF34803 - CDS
AIF34803 - replication-associated protein
AIF34803 - replication-associated protein
YP_009448204 - rep CDS
YP_009448204 - rep CDS
YP_009448204 - rolling-circle replication protein
YP_009448204 - rolling-circle replication protein
AIF76255 - CDS
AIF76255 - CDS
AIF76255 - Rep Protein
AIF76255 - Rep Protein
AXH74707 - CDS
AXH74707 - CDS
AXH74707 - Rep Protein
AXH74707 - Rep Protein
YP_009315918 - CDS
YP_009315918 - CDS
YP_009315918 - Rep Protein
YP_009315918 - Rep Protein
AXQ66426 - CDS
AXQ66426 - CDS
AXQ66426 - putative viral replication protein
AXQ66426 - putative viral replication protein
AXH75585 - CDS
AXH75585 - CDS
AXH75585 - putative viral replication protein
AXH75585 - putative viral replication protein
AUM61904 - Rep CDS
AUM61904 - Rep CDS
AUM61904 - Rep Protein
AUM61904 - Rep Protein
KM573768 (modified) - Replication associated protein translation - Replication associated protein
ETO15557 - CDS
ETO15557 - CDS
ETO15557 - rep Protein
ETO15557 - rep Protein
YP_009237504 - CDS
YP_009237504 - CDS
YP_009237504 - replication associated protein
YP_009237504 - replication associated protein
JX904523 (modified) - Replication associated protein translation - Replication associated protein
AUM61656 - Rep CDS
AUM61656 - Rep CDS
AUM61656 - Rep Protein
AUM61656 - Rep Protein
AGA18391 - CDS
AGA18391 - CDS
AGA18391 - hypothetical protein
AGA18391 - hypothetical protein
JX904427 (modified) - Replication associated protein translation - Replication associated protein
ORX42374 - CDS
ORX42374 - CDS
ORX42374 - hypothetical protein
ORX42374 - hypothetical protein
AUM61793 - Rep CDS
AUM61793 - Rep CDS
AUM61793 - Rep Protein
AUM61793 - Rep Protein
KU203355.1 - Replication associated protein translation - Replication associated protein
KU203354.1 - Replication associated protein translation - Replication associated protein
JX904394 (modified) - Replication associated protein translation - Replication associated protein
AQU11724 - CDS
AQU11724 - CDS
AQU11724 - replication protein
AQU11724 - replication protein
AQU11717 - CDS
AQU11717 - CDS
AQU11717 - replication protein
AQU11717 - replication protein
JF755407 (modified) - Replication associated protein interval 1 translation - Replication associated protein
KJ437671 (modified) - Movement protein translation - Movement protein
AJD07478 - CDS
AJD07478 - CDS
AJD07478 - replication-associated protein
AJD07478 - replication-associated protein
AFH02742 - CDS
AFH02742 - CDS
AFH02742 - putative Rep Protein
AFH02742 - putative Rep Protein
AXH74140 - CDS
AXH74140 - CDS
AXH74140 - helicase Protein
AXH74140 - helicase Protein
YP_009021245 - CDS
YP_009021245 - CDS
YP_009021245 - replication-associated protein
YP_009021245 - replication-associated protein
ARO38300 - Rep CDS
ARO38300 - Rep CDS
ARO38300 - replicase Protein
ARO38300 - replicase Protein
AGA18473 - CDS
AGA18473 - CDS
AGA18473 - hypothetical protein
AGA18473 - hypothetical protein
AUM61742 - Rep CDS
AUM61742 - Rep CDS
AUM61742 - Rep Protein
AUM61742 - Rep Protein
AQU11736 - CDS
AQU11736 - CDS
AQU11736 - replication protein
AQU11736 - replication protein
AQU11746 - CDS
AQU11746 - CDS
AQU11746 - replication protein
AQU11746 - replication protein
AUM61928 - Rep CDS
AUM61928 - Rep CDS
AUM61928 - Rep Protein
AUM61928 - Rep Protein
JX904548 (modified) - Replication associated protein translation - Replication associated protein
MF327576.1 - Replication associated protein translation - Replication associated protein
AXH74443 - CDS
AXH74443 - CDS
AXH74443 - putative viral replication protein
AXH74443 - putative viral replication protein
YP_009126892 - CDS
YP_009126892 - CDS
YP_009126892 - replication-associated protein
YP_009126892 - replication-associated protein
AUW34331 - Rep CDS
AUW34331 - Rep CDS
AUW34331 - replication-associated protein
AUW34331 - replication-associated protein
AGA18388 - CDS
AGA18388 - CDS
AGA18388 - hypothetical protein
AGA18388 - hypothetical protein
JX904420 (modified) - Replication associated protein translation - Replication associated protein

JX305998 (modified) - Replication associated protein
translation - hypothetical protein CDS
JX305998 (modified) - Replication associated protein
translation - Replication associated protein
JQ023166 (modified) - Replication associated protein
translation - hypothetical protein CDS
JX305991 (modified) - Replication associated protein
translation - hypothetical protein CDS
JX305992 (modified) - Replication associated protein
translation - hypothetical protein CDS
JX305993 (modified) - Replication associated protein
translation - hypothetical protein CDS
JX305994 (modified) - Replication associated protein
translation - hypothetical protein CDS
JX305995 (modified) - Replication associated protein
translation - hypothetical protein CDS
JX305996 (modified) - Replication associated protein
translation - hypothetical protein CDS
JX305997 (modified) - Replication associated protein
translation - hypothetical protein CDS
JX305992 (modified) - Replication associated protein
translation - Replication associated protein
JQ023166 (modified) - Replication associated protein
translation - Replication associated protein
JX305991 (modified) - Replication associated protein
translation - Replication associated protein
JX305993 (modified) - Replication associated protein
translation - Replication associated protein
JX305994 (modified) - Replication associated protein
translation - Replication associated protein
JX305995 (modified) - Replication associated protein
translation - Replication associated protein
JX305996 (modified) - Replication associated protein
translation - Replication associated protein
JX305997 (modified) - Replication associated protein
translation - Replication associated protein
AIF76255 - CDS
AIF76255 - Q-loop/lid
AIF76255 - Rep Protein
WP_095545523 - hypothetical protein
WP_095545523 - hypothetical protein
YP_009126938 - CDS
YP_009126938 - replication-associated protein
YP_009126938 - CDS
YP_009126938 - replication-associated protein
AUM61876 - Rep CDS
AUM61876 - Rep CDS
AUM61876 - Rep Protein
AUM61876 - Rep Protein
PKM67799 - CDS
PKM67799 - CDS
PKM67799 - replication protein
PKM67799 - replication protein
JX904439 (modified) - Replication associated protein
translation - Replication associated protein
AXQ66129 - CDS
AXQ66129 - CDS
AXQ66129 - putative viral replication protein
AXQ66129 - putative viral replication protein
YP_009126932 - CDS
YP_009126932 - CDS
YP_009126932 - replication-associated protein
YP_009126932 - replication-associated protein
OMJ25023 - CDS
OMJ25023 - CDS
OMJ25023 - hypothetical protein
OMJ25023 - hypothetical protein
AXH77121 - CDS
AXH77121 - CDS
AXH77121 - helicase Protein
AXH77121 - helicase Protein
JX904407 (modified) - Replication associated protein
translation - Replication associated protein
YP_009237564 - CDS
YP_009237564 - CDS
YP_009237564 - replication associated protein
YP_009237564 - replication associated protein
GAC77803 - CDS
GAC77803 - replication protein
GAC77803 - CDS
GAC77803 - replication protein
AUM62002 - Rep CDS
AUM62002 - Rep CDS
AUM62002 - Rep Protein
AUM62002 - Rep Protein
AUM61946 - Rep CDS
AUM61946 - Rep CDS
AUM61946 - Rep Protein
AUM61946 - Rep Protein
YP_009021888 - CDS
YP_009021888 - CDS
YP_009021888 - replication associated protein
YP_009021888 - replication associated protein
GAC77860 - CDS
GAC77860 - CDS
GAC77860 - replication protein
GAC77860 - replication protein
YP_008052687 - CDS
YP_008052687 - CDS
YP_008052687 - Rep domain protein
YP_008052687 - Rep domain protein
WP_027090230 - ATP-dependent Clp protease ATP-binding subunit Protein
WP_027090230 - ATP-dependent Clp protease ATP-binding subunit Protein
JX904139 (modified) - Replication associated protein
translation - Replication associated protein
AEL28813 - CDS
AEL28813 - CDS
AEL28813 - replication-associated protein
AEL28813 - replication-associated protein
YP_009126930 - CDS
YP_009126930 - CDS
YP_009126930 - replication-associated protein
YP_009126930 - replication-associated protein
JX904368 (modified) - Replication associated protein
translation - Replication associated protein
AQU11728 - CDS
AQU11728 - CDS
AQU11728 - replication protein
AQU11728 - replication protein
AXH76906 - CDS
AXH76906 - CDS
AXH76906 - putative replicase Protein
AXH76906 - putative replicase Protein
KU203356.1 - Replication associated protein translation - Replication associated protein
KR902498 (modified) - Replication associated protein
translation - Replication associated protein
YP_009170674 - rep CDS
YP_009170674 - rep CDS
YP_009170674 - replicase Protein
YP_009170674 - replicase Protein
AMB43004 - CDS
AMB43004 - CDS
AMB43004 - putative Rep protein
AMB43004 - putative Rep protein
YP_009126881 - CDS
YP_009126881 - CDS
YP_009126881 - replication-associated protein
YP_009126881 - replication-associated protein
MF327575.1 - Replication associated protein translation - Replication associated protein
KM573766 (modified) - Replication associated protein
translation - Replication associated protein
GAC77844 - CDS
GAC77844 - CDS
GAC77844 - replication protein
GAC77844 - replication protein
AEL87784 - CDS
AEL87784 - CDS
AEL87784 - putative replication-associated protein
AEL87784 - putative replication-associated protein
CBK25810 - rep CDS
CBK25810 - rep CDS
CBK25810 - replication association protein
CBK25810 - replication association protein
YP_009237555 - CDS
YP_009237555 - CDS
YP_009237555 - replication associated protein
YP_009237555 - replication associated protein
AUM62047 - Rep CDS
AUM62047 - Rep CDS
AUM62047 - Rep Protein
AUM62047 - Rep Protein
AEL28793 - CDS
AEL28793 - replication-associated protein
AEL28793 - CDS
AEL28793 - replication-associated protein
YP_009109670 - CDS
YP_009109670 - CDS
YP_009109670 - replication-associated protein
YP_009109670 - replication-associated protein
YP_004376332 - CDS
YP_004376332 - CDS
YP_004376332 - putative replication protein
YP_004376332 - putative replication protein
JX904250 (modified) - Replication associated protein
translation - Replication associated protein
JX904245 (modified) - Replication associated protein
translation - Replication associated protein
AXH73508 - CDS
AXH73508 - CDS
AXH73508 - putative viral replication protein
AXH73508 - putative viral replication protein
ALE29827 - CDS
ALE29827 - CDS
ALE29827 - replication associated protein
ALE29827 - replication associated protein
AGG39817 - CDS
AGG39817 - CDS
AGG39817 - replication-associated protein
AGG39817 - replication-associated protein
AXH76879 - CDS
AXH76879 - CDS
AXH76879 - putative viral replication protein
AXH76879 - putative viral replication protein
AMH87735 - CDS
AMH87735 - CDS
AMH87735 - replication-associated protein
AMH87735 - replication-associated protein
AUF34964 - CDS
AUF34964 - CDS
AUF34964 - putative replication-associated protein
AUF34964 - putative replication-associated protein
PVV02139 - CDS
PVV02139 - CDS
PVV02139 - hypothetical protein
PVV02139 - hypothetical protein
AEL28791 - CDS
AEL28791 - replication-associated protein
AQU11749 - CDS
AQU11749 - CDS
AQU11749 - replication protein
AQU11749 - replication protein
YP_764455 - ORFV1; putative replicase CDS
YP_764455 - ORFV1; putative replicase CDS
YP_764455 - rep protein
YP_764455 - rep protein
AUM61805 - Rep CDS
AUM61805 - Rep CDS
AUM61805 - Rep Protein
AUM61805 - Rep Protein
AQU11779 - CDS
AQU11779 - CDS
AQU11779 - replication protein
AQU11779 - replication protein
JX904377 (modified) - Replication associated protein
translation - Replication associated protein
OWF45429 - CDS
OWF45429 - CDS
OWF45429 - Master replication protein
OWF45429 - Master replication protein
AXH75487 - CDS
AXH75487 - CDS
AXH75487 - putative viral replication protein
AXH75487 - putative viral replication protein
AXL65927 - CDS
AXL65927 - CDS
AXL65927 - replication-associated protein
AXL65927 - replication-associated protein
AGA19549 - C1 CDS
AGA19549 - C1 CDS
AGA19549 - Rep Protein
AGA19549 - Rep Protein
YP_009109643 - CDS
YP_009109643 - CDS
YP_009109643 - replication-associated protein
YP_009109643 - replication-associated protein
ATY70087 - Rep CDS
ATY70087 - Rep CDS
ATY70087 - replication initiator protein
ATY70087 - replication initiator protein
AXL65946 - CDS
AXL65946 - CDS
AXL65946 - replication-associated protein
AXL65946 - replication-associated protein
ADY62649 - CDS
ADY62649 - CDS
ADY62649 - Rep Protein
ADY62649 - Rep Protein
AKO63006 - CDS
AKO63006 - CDS
AKO63006 - replication-associated protein
AKO63006 - replication-associated protein
ACE62799 - rep CDS
ACE62799 - rep CDS
ACE62799 - Rep Protein
ACE62799 - Rep Protein
ANH56850 - CDS
ANH56850 - CDS
ANH56850 - rep protein
ANH56850 - rep protein
YP_009508846 - rep CDS
YP_009508846 - rep CDS
YP_009508846 - rep protein
YP_009508846 - rep protein
AJE25853 - Rep CDS
AJE25853 - Rep CDS
AJE25853 - Rep Protein
AJE25853 - Rep Protein
AJF23058 - rep CDS
AJF23058 - rep CDS
AJF23058 - rep protein
AJF23058 - rep protein
AIF76277 - CDS
AIF76277 - CDS
AIF76277 - Rep Protein
AIF76277 - Rep Protein
AIW81537 - ORF1 CDS
AIW81537 - ORF1 CDS
AIW81537 - replicase protein
AIW81537 - replicase protein
YP_009126879 - CDS
YP_009126879 - CDS
YP_009126879 - replication-associated protein
YP_009126879 - replication-associated protein
JX904541 (modified) - Replication associated protein
translation - Replication associated protein
KR704912 - Replication associated protein translation - Replication associated protein
AXH77057 - CDS
AXH77057 - CDS
AXH77057 - Rep Protein
AXH77057 - Rep Protein
NP_619761 - putative CDS
NP_619761 - putative CDS
NP_619761 - virus replication-associated protein
NP_619761 - virus replication-associated protein
JX904559 (modified) - Replication associated protein
translation - Replication associated protein
YP_009126898 - CDS
YP_009126898 - CDS
YP_009126898 - replication-associated protein
YP_009126898 - replication-associated protein
AUM61616 - Rep CDS
AUM61616 - Rep CDS
AUM61616 - Rep Protein
AUM61616 - Rep Protein
MF118167.1 - Replication associated protein translation - Replication associated protein
YP_009237554 - CDS
YP_009237554 - CDS
YP_009237554 - replication associated protein
YP_009237554 - replication associated protein
AJP36430 - CDS
AJP36430 - CDS
AJP36430 - replication-associated protein
AJP36430 - replication-associated protein
AOV86285 - CDS
AOV86285 - CDS
AOV86285 - putative rep protein
AOV86285 - putative rep protein
HM228875 (modified) - Replication associated protein
translation - Replication associated protein
YP_009163920 - CDS
YP_009163920 - CDS
YP_009163920 - putative spliced replication initiation protein

YP_009163920 - putative spliced replication initiation protein
AMD39533 - replication-associated protein
AMD39533 - replication-associated protein
AMD39533 - V1 CDS
AMD39533 - V1 CDS
JX904107 (modified) - Replication associated protein translation - Replication associated protein
YP_009259728 - CDS
YP_009259728 - CDS
YP_009259728 - putative Rep protein
YP_009259728 - putative Rep protein
AUM62041 - Rep CDS
AUM62041 - Rep CDS
AUM62041 - Rep Protein
AUM62041 - Rep Protein
YP_009142778 - putative replication protein
YP_009142778 - putative replication protein
YP_009142778 - rep CDS
YP_009142778 - rep CDS
KM105952 (modified) - Replication associated protein translation - Replication associated protein
AXH75991 - CDS
AXH75991 - CDS
AXH75991 - putative viral replication protein
AXH75991 - putative viral replication protein
AXQ66530 - CDS
AXQ66530 - CDS
AXQ66530 - replication protein
AXQ66530 - replication protein
EES99438 - CDS
EES99438 - CDS
EES99438 - Replicase-associated protein, putative
EES99438 - Replicase-associated protein, putative
AUM61662 - Rep CDS
AUM61662 - Rep CDS
AUM61662 - Rep Protein
AUM61662 - Rep Protein
AGA18245 - CDS
AGA18245 - CDS
AGA18245 - hypothetical protein
AGA18245 - hypothetical protein
JX904075 (modified) - Replication associated protein translation - Replication associated protein
AUM61736 - Rep CDS
AUM61736 - Rep CDS
AUM61736 - Rep Protein
AUM61736 - Rep Protein
AXH77952 - Geminivirus Rep catalytic domain CDS
AXH77952 - Geminivirus Rep catalytic domain CDS
AXH77952 - Rep catalytic domain protein
AXH77952 - Rep catalytic domain protein
AUM61958 - Rep CDS
AUM61958 - Rep CDS
AUM61958 - Rep Protein
AUM61958 - Rep Protein
MF118166.1 - Replication associated protein translation - Replication associated protein
AJD07498 - CDS
AJD07498 - CDS
AJD07498 - replication-associated protein
AJD07498 - replication-associated protein
AGS47835 - CDS
AGS47835 - CDS
AGS47835 - replication-associated protein
AGS47835 - replication-associated protein
SCN47931 - Rep CDS
SCN47931 - Rep CDS
SCN47931 - Replication initiator protein
SCN47931 - Replication initiator protein
AIF76269 - CDS
AIF76269 - CDS
AIF76269 - Rep Protein
AIF76269 - Rep Protein
AUM61856 - Rep CDS
AUM61856 - Rep CDS
AUM61856 - Rep Protein
AUM61856 - Rep Protein
KM573767 (modified) - Replication associated protein translation - Replication associated protein
JF755401 (modified) - Replication associated protein translation - Replication associated protein
AXH76896 - CDS
AXH76896 - CDS
AXH76896 - replication protein
AXH76896 - replication protein
AXH73382 - CDS
AXH73382 - CDS
AXH73382 - putative viral replication protein
AXH73382 - putative viral replication protein
AUM61874 - Rep CDS
AUM61874 - Rep CDS
AUM61874 - Rep Protein
AUM61874 - Rep Protein
AXH75674 - CDS
AXH75674 - CDS
AXH75674 - Rep Protein
AXH75674 - Rep Protein
AJD07493 - CDS
AJD07493 - CDS
AJD07493 - replication-associated protein
AJD07493 - replication-associated protein
APZ87906 - CDS
APZ87906 - CDS
APZ87906 - replication-associated protein
APZ87906 - replication-associated protein
AJM89742 - CDS
AJM89742 - CDS
AJM89742 - replication associated protein
AJM89742 - replication associated protein
AQR57902 - Rep CDS
AQR57902 - Rep CDS
AQR57902 - replicase Protein
AQR57902 - replicase Protein
AXH73061 - CDS
AXH73061 - CDS
AXH73061 - putative viral replication protein
AXH73061 - putative viral replication protein
AUM61773 - Rep CDS
AUM61773 - Rep CDS
AUM61773 - Rep Protein
AUM61773 - Rep Protein
AXH73290 - CDS
AXH73290 - CDS
AXH73290 - putative viral replication protein
AXH73290 - putative viral replication protein
AXH75836 - CDS
AXH75836 - CDS
AXH75836 - replication-associated protein
AXH75836 - replication-associated protein
KM573773 (modified) - Replication associated protein translation - Replication associated protein
APA62657 - CDS
APA62657 - CDS
APA62657 - putative replication protein
APA62657 - putative replication protein
YP_009116902 - CDS
YP_009116902 - CDS
YP_009116902 - replication-associated protein
YP_009116902 - replication-associated protein
JF713717 (modified) - unknown CDS translation 3 - unknown CDS
AIF34798 - CDS
AIF34798 - CDS
AIF34798 - replication-associated protein
AIF34798 - replication-associated protein
MF327573.1 - Replication associated protein translation - Replication associated protein
JX904344 (modified) - Replication associated protein translation - Replication associated protein
APA62647 - CDS
APA62647 - CDS
APA62647 - putative replication protein
APA62647 - putative replication protein
AUM61894 - Rep CDS
AUM61894 - Rep CDS
AUM61894 - Rep Protein
AUM61894 - Rep Protein
AUM61960 - Rep CDS
AUM61960 - Rep CDS
AUM61960 - Rep Protein
AUM61960 - Rep Protein
KU203352.1 - Replication associated protein translation - Replication associated protein
JF755410 (modified) - Replication associated protein translation - Replication associated protein
APC94137 - putative replication-associated protein
APC94137 - putative replication-associated protein
APC94137 - similar to YP 009126890.1 CDS
APC94137 - similar to YP 009126890.1 CDS
KM573776 (modified) - Replication associated protein translation - Replication associated protein
YP_009237578 - CDS
YP_009237578 - CDS
YP_009237578 - replication associated protein
YP_009237578 - replication associated protein
AEM05804 - CDS
AEM05804 - CDS
AEM05804 - REP Protein
AEM05804 - REP Protein
JF755409 (modified) - Replication associated protein translation - Replication associated protein
AXL65944 - CDS
AXL65944 - CDS
AXL65944 - replication-associated protein
AXL65944 - replication-associated protein
YP_009259737 - CDS
YP_009259737 - CDS
YP_009259737 - putative Rep protein
YP_009259737 - putative Rep protein
GAC77783 - CDS
GAC77783 - CDS
GAC77783 - replication protein
GAC77783 - replication protein
AGA18448 - CDS
AGA18448 - CDS
AGA18448 - hypothetical protein
AGA18448 - hypothetical protein
JX904581 (modified) - Replication associated protein translation - Replication associated protein
YP_009126925 - CDS
YP_009126925 - CDS
YP_009126925 - replication-associated protein
YP_009126925 - replication-associated protein
YP_009109675 - CDS
YP_009109675 - CDS
YP_009109675 - replication-associated protein
YP_009109675 - replication-associated protein
YP_006281010 - CDS
YP_006281010 - CDS
YP_006281010 - putative viral replication protein
YP_006281010 - putative viral replication protein
AXH75780 - CDS
AXH75780 - CDS
AXH75780 - replication-associated protein
AXH75780 - replication-associated protein
AXH73792 - CDS
AXH73792 - CDS
AXH73792 - putative viral replication protein
AXH73792 - putative viral replication protein
AHC72271 - M-Rep CDS
AHC72271 - M-Rep CDS
AHC72271 - master replication initiator protein
AHC72271 - master replication initiator protein
YP_008997794 - M-Rep CDS
YP_008997794 - M-Rep CDS
YP_008997794 - master replication initiator protein
YP_008997794 - master replication initiator protein
YP_009126890 - CDS
YP_009126890 - CDS
YP_009126890 - replication-associated protein
YP_009126890 - replication-associated protein
AOV86255 - CDS
AOV86255 - CDS
AOV86255 - putative rep protein
AOV86255 - putative rep protein
AUM61787 - Rep CDS
AUM61787 - Rep CDS
AUM61787 - Rep Protein
AUM61787 - Rep Protein
YP_009506291 - CDS
YP_009506291 - CDS
YP_009506291 - Rep Protein
YP_009506291 - Rep Protein
AUM61801 - Rep CDS
AUM61801 - Rep CDS
AUM61801 - Rep Protein
AUM61801 - Rep Protein
MF118169.1 - Replication associated protein translation - Replication associated protein
JF755408 (modified) - Replication associated protein translation - Replication associated protein
AXQ65661 - CDS
AXQ65661 - CDS
AXQ65661 - putative viral replication protein
AXQ65661 - putative viral replication protein
YP_009508165 - V2 CDS
YP_009508165 - V2 CDS
YP_009508165 - V2 Protein
YP_009508165 - V2 Protein
KM573765 (modified) - Replication associated protein translation - Replication associated protein
YP_009116906 - CDS
YP_009116906 - CDS
YP_009116906 - replication-associated protein
YP_009116906 - replication-associated protein
AXH73393 - CDS
AXH73393 - CDS
AXH73393 - putative viral replication protein
AXH73393 - putative viral replication protein
AXH76040 - CDS
AXH76040 - CDS
AXH76040 - putative viral replication protein
AXH76040 - putative viral replication protein
JX904401 (modified) - Replication associated protein translation - Replication associated protein
JX904192 (modified) - Replication associated protein translation - Replication associated protein
MF118168.1 - Replication associated protein translation - Replication associated protein
JX904412 (modified) - Replication associated protein translation - Replication associated protein
AXH77580 - CDS
AXH77580 - CDS
AXH77580 - replication protein
AXH77580 - replication protein
AXH77287 - CDS
AXH77287 - CDS
AXH77287 - Rep Protein
AXH77287 - Rep Protein
AXH78100 - CDS
AXH78100 - CDS
AXH78100 - helicase Protein
AXH78100 - helicase Protein
JF755416 (modified) - Replication associated protein interval 1 translation - Replication associated protein
JF755417 (modified) - Replication associated protein interval 1 translation - Replication associated protein
YP_009237571 - CDS
YP_009237571 - CDS
YP_009237571 - replication associated protein
YP_009237571 - replication associated protein
AXH75489 - CDS
AXH75489 - CDS
AXH75489 - replication-associated protein
AXH75489 - replication-associated protein
JX904395 (modified) - Replication associated protein translation - Replication associated protein
AQU11729 - CDS
AQU11729 - replication protein
AQU11729 - CDS
AQU11729 - replication protein
AQU11743 - CDS
AQU11743 - CDS
AQU11743 - replication protein
AQU11743 - replication protein
AXH74669 - CDS
AXH74669 - CDS
AXH74669 - replication protein
AXH74669 - replication protein
YP_009115538 - CDS
YP_009115538 - CDS
YP_009115538 - replication-associated protein
YP_009115538 - replication-associated protein
AXH77740 - CDS
AXH77740 - CDS
AXH77740 - putative viral replication protein
AXH77740 - putative viral replication protein
AUM61811 - Rep CDS
AUM61811 - Rep CDS
AUM61811 - Rep Protein
AUM61811 - Rep Protein
YP_009237516 - CDS
YP_009237516 - CDS
YP_009237516 - replication associated protein
YP_009237516 - replication associated protein
AUM61707 - Rep CDS
AUM61707 - Rep CDS
AUM61707 - Rep Protein

AUM61707 - Rep Protein
AUM61940 - Rep CDS
AUM61940 - Rep CDS
AUM61940 - Rep Protein
AUM61940 - Rep Protein
AJF23074 - rep CDS
AJF23074 - rep CDS
AJF23074 - rep protein
AJF23074 - rep protein
AJF23080 - rep CDS
AJF23080 - rep CDS
AJF23080 - rep protein
AJF23080 - rep protein
YP_009226567 - CDS
YP_009226567 - CDS
YP_009226567 - replication-associated protein
YP_009226567 - replication-associated protein
AUM61730 - Rep CDS
AUM61730 - Rep CDS
AUM61730 - Rep Protein
AUM61730 - Rep Protein
AJD20393 - CDS
AJD20393 - CDS
AJD20393 - replication associated protein
AJD20393 - replication associated protein
JF755415 (modified) - Replication associated protein interval 1 translation - Replication associated protein
AXQ65784 - CDS
AXQ65784 - CDS
AXQ65784 - replication associated protein
AXQ65784 - replication associated protein
AUM61795 - Rep CDS
AUM61795 - Rep CDS
AUM61795 - Rep Protein
AUM61795 - Rep Protein
ARD71303 - putative replication initiator protein
ARD71303 - putative replication initiator protein
ARD71303 - Rep CDS
ARD71303 - Rep CDS
AXH73056 - CDS
AXH73056 - CDS
AXH73056 - putative viral replication protein
AXH73056 - putative viral replication protein
AIX11626 - Rep CDS
AIX11626 - Rep CDS
AIX11626 - replicase Protein
AIX11626 - replicase Protein
APG55798 - CDS
APG55798 - CDS
APG55798 - Rep Protein
APG55798 - Rep Protein
AHB63242 - CDS
AHB63242 - CDS
AHB63242 - replication associated protein
AHB63242 - replication associated protein
JX904561 (modified) - Replication associated protein translation - Replication associated protein
AXH76632 - CDS
AXH76632 - CDS
AXH76632 - helicase Protein
AXH76632 - helicase Protein
JF755402 (modified) - Replication associated protein translation - Replication associated protein
JF755403 (modified) - Replication associated protein translation - Replication associated protein
KM573763 (modified) - Replication associated protein translation - Replication associated protein
KM573764 (modified) - Replication associated protein translation - Replication associated protein
AOV86329 - CDS
AOV86329 - CDS
AOV86329 - putative rep protein
AOV86329 - putative rep protein
YP_003084282 - CDS
YP_003084282 - CDS
YP_003084282 - putative Rep protein
YP_003084282 - putative Rep protein
JX904312 (modified) - Replication associated protein translation - Replication associated protein
AXH77906 - CDS
AXH77906 - CDS
AXH77906 - replication protein
AXH77906 - replication protein
AXH74257 - CDS
AXH74257 - CDS
AXH74257 - replication associated protein
AXH74257 - replication associated protein
AWW06057 - CDS
AWW06057 - CDS
AWW06057 - helicase Protein
AWW06057 - helicase Protein
AQU11773 - CDS
AQU11773 - CDS
AQU11773 - replication protein
AQU11773 - replication protein
AQU11776 - CDS
AQU11776 - CDS
AQU11776 - replication protein
AQU11776 - replication protein
AXH74332 - CDS
AXH74332 - CDS
AXH74332 - replication-associated protein
AXH74332 - replication-associated protein
AUM61906 - Rep CDS
AUM61906 - Rep CDS
AUM61906 - Rep Protein
AUM61906 - Rep Protein
YP_009237586 - CDS
YP_009237586 - CDS
YP_009237586 - replication associated protein
YP_009237586 - replication associated protein
JX559621 (modified) - hypothetical protein CDS translation 2 - hypothetical protein CDS
JX559622 (modified) - hypothetical protein CDS translation 2 - hypothetical protein CDS
YP_009163927 - CDS
YP_009163927 - CDS
YP_009163927 - putative replication initiation protein
YP_009163927 - putative replication initiation protein
JX904478 (modified) - Replication associated protein translation - Replication associated protein
AXH76508 - CDS
AXH76508 - CDS
AXH76508 - helicase Protein
AXH76508 - helicase Protein
AXH77564 - CDS
AXH77564 - CDS
AXH77564 - putative viral replication protein
AXH77564 - putative viral replication protein
JX904416 (modified) - Replication associated protein translation - Replication associated protein
YP_009051960 - CDS
YP_009051960 - CDS
YP_009051960 - replication-associated protein
YP_009051960 - replication-associated protein
KJ206566 (modified) - Replication associated protein translation - Replication associated protein
YP_009237541 - CDS
YP_009237541 - CDS
YP_009237541 - replication associated protein
YP_009237541 - replication associated protein
WP_027090230 - other interval 3
YP_009237578 - other interval 3
AIF76278 - CDS
AIF76278 - Rep Protein
YP_009237578 - Walker A motif
YP_009237578 - other interval 1
YP_009237578 - CDS
YP_009237578 - replication associated protein
YP_009237578 - RNA helicase
WP_027090230 - Walker A motif
WP_027090230 - other interval 1
WP_027090230 - P-loop NTPase
WP_027090230 - AAA
WP_027090230 - ATP-dependent Clp protease ATP-binding subunit Protein
WP_027090230 - P-loop NTPase
YP_009237554 - CDS
YP_009237554 - Gemini AL1
YP_009237554 - replication associated protein
WP_027090230 - AAA
WP_027090230 - arginine finger
WP_027090230 - ATP-dependent Clp protease ATP-binding subunit Protein
WP_027090230 - P-loop NTPase
AIF34798 - CDS
AIF34798 - replication-associated protein
AIF34798 - Viral Rep
AGA18393 - CDS
AGA18393 - hypothetical protein
AGA18393 - Viral Rep
AXH76451 - CDS
AXH76451 - helicase Protein
AXH76451 - Viral Rep
YP_009226567 - CDS
YP_009226567 - replication-associated protein
YP_009226567 - Viral Rep
AQU11736 - CDS
AQU11736 - replication protein
AQU11736 - Viral Rep
AGA18448 - CDS
AGA18448 - hypothetical protein
AGA18448 - Viral Rep
AQU11717 - CDS
AQU11724 - CDS
AQU11717 - replication protein
AQU11724 - replication protein
AQU11717 - Viral Rep
AQU11724 - Viral Rep
AXH76508 - CDS
AXH76508 - helicase Protein
AXH76508 - Viral Rep
AIF34812 - CDS
AIF34812 - Gemini AL1
AIF34812 - replication-associated protein
AJP36430 - CDS
AJP36430 - replication-associated protein
AJP36430 - Viral Rep
AQU11726 - CDS
AQU11726 - replication protein
AQU11726 - Viral Rep
YP_009237541 - AAA
YP_009237541 - CDS
YP_009237541 - replication associated protein
YP_009448204 - rep CDS
YP_009448204 - rolling-circle replication protein
YP_009448204 - Viral Rep
YP_009126925 - CDS
YP_009126925 - replication-associated protein
YP_009126925 - Viral Rep
YP_009115538 - CDS
YP_009115538 - replication-associated protein
YP_009115538 - Viral Rep
AGA18286 - CDS
AGA18286 - hypothetical protein
AGA18286 - Viral Rep
YP_009237541 - CDS
YP_009237541 - replication associated protein
YP_009237541 - Viral Rep
AKO84203 - CDS
AKO84203 - replicase Protein
AKO84203 - Viral Rep
YP_009170674 - rep CDS
YP_009170674 - replicase Protein
YP_009170674 - Viral Rep
ADY62649 - CDS
ADY62649 - Rep Protein
ADY62649 - Viral Rep
AHB63242 - CDS
AHB63242 - replication associated protein
AHB63242 - Viral Rep
YP_009126938 - CDS
YP_009126938 - replication-associated protein
YP_009126938 - Viral Rep
AXH76879 - CDS
AXH76879 - putative viral replication protein
AXH76879 - Viral Rep
AIF76269 - CDS
AIF76269 - Rep Protein
AIF76269 - Viral Rep
AXH77740 - CDS
AXH77740 - putative viral replication protein
AXH77740 - Viral Rep
AVA16977 - replicase-associated protein
AVA16977 - start codon not determined CDS
AVA16977 - Viral Rep
AVV68420 - CDS
AVV68420 - replication-associated protein
AVV68420 - Viral Rep
AVA17000 - replicase-associated protein
AVA17000 - start codon not determined CDS
AVA17000 - Viral Rep
AVA16996 - replicase-associated protein
AVA16996 - start codon not determined CDS
AVA16996 - Viral Rep
ATD53351 - ORF1 CDS
ATD53351 - Rep Protein
AVA16998 - replicase-associated protein
AVA16998 - start codon not determined CDS
ATD53351 - Viral Rep
AVA16998 - Viral Rep
AWB80902 - Rep CDS
AWB80902 - replicase protein
AWB80902 - Viral Rep
YP_008052687 - CDS
YP_008052687 - Rep domain protein
YP_008052687 - Viral Rep
ADC79191 - CDS
ADC79191 - V1 Protein
ADC79191 - Viral Rep
YP_009021888 - CDS
YP_009021888 - replication associated protein
YP_009021888 - Viral Rep
AUT13975 - CDS
AUT13975 - replication protein
AUT13975 - Viral Rep
AIW81537 - ORF1 CDS
ACE62799 - rep CDS
ACE62799 - Rep Protein
AIW81537 - replicase protein
ACE62799 - Viral Rep
AIW81537 - Viral Rep
JX305991 (modified) - hypothetical protein CDS interval 2 translation - hypothetical protein CDS
JX305993 (modified) - hypothetical protein CDS interval 2 translation - hypothetical protein CDS
JX305994 (modified) - hypothetical protein CDS interval 2 translation - hypothetical protein CDS
JX305995 (modified) - hypothetical protein CDS interval 2 translation - hypothetical protein CDS
JX305996 (modified) - hypothetical protein CDS interval 2 translation - hypothetical protein CDS
JX305997 (modified) - hypothetical protein CDS interval 2 translation - hypothetical protein CDS
JX305998 (modified) - hypothetical protein CDS interval 2 translation - hypothetical protein CDS
JX305992 (modified) - hypothetical protein CDS interval 2 translation - hypothetical protein CDS
JQ023166 (modified) - hypothetical protein CDS interval 2 translation - hypothetical protein CDS
AEL22996 - CDS
YP_004778177 - CDS
AEL22996 - Rep protein
YP_004778177 - Rep protein
AEL22996 - Viral Rep
YP_004778177 - Viral Rep
AXH73508 - CDS
AXH73508 - putative viral replication protein
AXH73508 - Viral Rep
AXH75487 - CDS
AXH75487 - putative viral replication protein
AXH75487 - Viral Rep
AXH76667 - CDS
AXH76667 - putative viral replication protein
AXH76667 - Viral Rep
AQU11728 - CDS
AQU11728 - replication protein
AQU11728 - Viral Rep
AXH74936 - CDS
AXH74936 - putative viral replication protein
AXH74936 - Viral Rep
ARO38300 - Rep CDS
ARO38300 - replicase Protein
ARO38300 - Viral Rep
YP_009116906 - CDS
YP_009116906 - replication-associated protein
YP_009116906 - Viral Rep
AXH73393 - CDS
AXH76040 - CDS
AXH73393 - putative viral replication protein
AXH76040 - putative viral replication protein
AXH73393 - Viral Rep
AXH76040 - Viral Rep
AIF76255 - CDS
AIF76255 - Rep Protein
AIF76255 - Viral Rep
YP_009237516 - CDS
YP_009237516 - replication associated protein
YP_009237516 - Viral Rep

ALE29847 - CDS
YP_009001747 - CDS
ALE29847 - replication associated protein
YP_009001747 - replication-associated protein
ALE29847 - Viral Rep
YP_009001747 - Viral Rep
YP_009237564 - CDS
YP_009237564 - replication associated protein
YP_009237564 - Viral Rep
AJD07478 - CDS
AJD07478 - replication-associated protein
AJD07478 - Viral Rep
ALE29688 - CDS
YP_009001742 - CDS
ALE29688 - replication associated protein
YP_009001742 - replication-associated protein
ALE29688 - Viral Rep
YP_009001742 - Viral Rep
YP_009237586 - CDS
YP_009237586 - replication associated protein
YP_009237586 - Viral Rep
AMH87735 - CDS
AMH87735 - replication-associated protein
AMH87735 - Viral Rep
AXH73382 - CDS
AXH73382 - putative viral replication protein
AXH73382 - Viral Rep
ARE67375 - SWPV2-147 CDS
ARE67375 - SWPV2-ORF147 Protein
ARE67375 - Viral Rep
NP_955176 - CDS
NP_955176 - Rep-like protein
NP_955176 - Viral Rep
AKR53286 - CDS
AKR53286 - Viral Rep
AKR53286 - viral replicase protein
AGA18388 - CDS
AGA18388 - hypothetical protein
AGA18388 - Viral Rep
YP_009126890 - CDS
YP_009126890 - replication-associated protein
YP_009126890 - Viral Rep
AGA18473 - CDS
AGA18473 - hypothetical protein
AGA18473 - Viral Rep
AGS47835 - CDS
AGS47835 - replication-associated protein
AGS47835 - Viral Rep
YP_009109670 - CDS
YP_009109670 - replication-associated protein
YP_009109670 - Viral Rep
AQR57898 - Rep CDS
AQR57898 - replicase Protein
AQR57898 - Viral Rep
AGA18245 - CDS
AGA18245 - hypothetical protein
AGA18245 - Viral Rep
AQR57902 - Rep CDS
AQR57902 - replicase Protein
AQR57902 - Viral Rep
AXH73061 - CDS
AXH73061 - putative viral replication protein
AXH73061 - Viral Rep
NP_619761 - putative CDS
NP_619761 - Viral Rep
NP_619761 - virus replication-associated protein
YP_009237578 - CDS
YP_009237578 - replication associated protein
YP_009237578 - Viral Rep
AQU11729 - CDS
AQU11729 - replication protein
AQU11729 - Viral Rep
AJD07493 - CDS
AJD07493 - replication-associated protein
AJD07493 - Viral Rep
KM972725 (modified) - Replication associated protein interval 2 translation - Replication associated protein
KM972726 (modified) - Replication associated protein interval 2 translation - Replication associated protein
AQU11726 - CDS
AQU11726 - P-loop NTPase
AQU11726 - replication protein
AVV68420 - CDS
AVA16977 - replicase-associated protein
AVV68420 - replication-associated protein
AVA16977 - RNA helicase
AVV68420 - RNA helicase
AVA16977 - start codon not determined CDS
AVH76405 - CDS
AVH76405 - putative Rep Protein
AVH76405 - Viral Rep
YP_009163920 - CDS
YP_009163920 - P-loop NTPase
YP_009163920 - putative spliced replication initiation protein
ARD71303 - P-loop NTPase
ARD71303 - putative replication initiator protein
ARD71303 - Rep CDS
AIF34818 - CDS
AIF34818 - P-loop NTPase
AIF34818 - replication-associated protein
AXH73508 - CDS
AXH73508 - putative viral replication protein
AXH73508 - RNA helicase
AXG50856 - M-Rep CDS
AXG50856 - master replication initiator protein
AXG50856 - Viral Rep
AHC72271 - M-Rep CDS
AHC72271 - master replication initiator protein
AHC72271 - Viral Rep
YP_008997794 - M-Rep CDS
YP_008997794 - master replication initiator protein
YP_008997794 - Viral Rep
AHC72167 - M-Rep CDS
AHC72167 - master replication initiator protein
AHC72167 - Viral Rep
AHC72177 - M-Rep CDS
AHC72177 - master replication initiator protein
AHC72177 - Viral Rep
WP_027090230 - P-loop NTPase
WP_027090230 - P-loop NTPase
WP_027090230 - AAA
WP_027090230 - ATP-dependent Clp protease ATP-binding subunit Protein
WP_027090230 - P-loop NTPase
WP_027090230 - AAA
WP_027090230 - ATP-dependent Clp protease ATP-binding subunit Protein
WP_027090230 - P-loop NTPase
AXH76632 - CDS
AXH76632 - helicase Protein
AXH76632 - P-loop NTPase
ACE62799 - rep CDS
ACE62799 - Rep Protein
ACE62799 - RNA helicase
AHB63242 - CDS
AHB63242 - replication associated protein
AHB63242 - RNA helicase
ADY62649 - CDS
ADY62649 - Rep Protein
ADY62649 - RNA helicase
ARE68406 - CDS
ARE68406 - replication associated protein
ARE68406 - RNA helicase
YP_004376332 - CDS
YP_004376332 - putative replication protein
YP_004376332 - Viral Rep
YP_009163936 - CDS
YP_009163936 - putative replication initiation protein
YP_009163936 - Viral Rep
ATY42470 - CDS
ATY42470 - putative replication associated protein
ATY42470 - RNA helicase
AIL50149 - CDS
AIL50149 - replicase Protein
AIL50149 - RNA helicase
AVT56110 - CDS
AVT56110 - replicase Protein
AVT56110 - RNA helicase
ADC79191 - CDS
ADC79191 - P-loop NTPase
ADC79191 - V1 Protein
AKO84203 - CDS
AKO84203 - replicase Protein
AKO84203 - RNA helicase
YP_009237578 - CDS
YP_009237578 - replication associated protein
YP_009237578 - RNA helicase
YP_009237578 - Walker B motif
YP_009051960 - CDS
YP_009051960 - P-loop NTPase
YP_009051960 - replication-associated protein
APG55798 - CDS
APG55798 - P-loop NTPase
APG55798 - Rep Protein
APZ87906 - CDS
APZ87906 - replication-associated protein
APZ87906 - RNA helicase
AFH02742 - CDS
AFH02742 - putative Rep Protein
AFH02742 - Viral Rep
AEL28813 - CDS
AEL28813 - replication-associated protein
AEL28813 - Viral Rep
AXH74140 - CDS
AXH74140 - helicase Protein
AXH74140 - Viral Rep
AVA16977 - replicase-associated protein
AVA16977 - start codon not determined CDS
AVV68420 - CDS
AVV68420 - replication-associated protein
AVA17000 - replicase-associated protein
AVA17000 - start codon not determined CDS
AVA16996 - replicase-associated protein
AVA16996 - start codon not determined CDS
ATD53351 - ORF1 CDS
ATD53351 - Rep Protein
AVA16998 - replicase-associated protein
AVA16998 - start codon not determined CDS
AWB80902 - Rep CDS
AWB80902 - replicase protein
YP_009458619 - rep CDS
YP_009458619 - replication-association protein
YP_009458619 - Viral Rep
BAP81877 - Rep CDS
BAP81877 - rolling circle replication initiator protein
BAP81877 - Viral Rep
APZ87906 - CDS
APZ87906 - replication-associated protein
APZ87906 - Viral Rep
AVT56110 - CDS
AVT56110 - replicase Protein
AVT56110 - Viral Rep
AIL50149 - CDS
AIL50149 - replicase Protein
AIL50149 - Viral Rep
ATY42470 - CDS
ATY42470 - putative replication associated protein
ATY42470 - Viral Rep
AKO71368 - CDS
AKO71368 - Replication associated protein
AKO71368 - Viral Rep
ALA65733 - CDS
ALA65733 - replication initiation protein
ALA65733 - Viral Rep
AIF76278 - CDS
AIF76278 - Rep Protein
AIF76278 - RNA helicase
YP_009126925 - CDS
YP_009126925 - replication-associated protein
YP_009126925 - RNA helicase
YP_009109643 - CDS
YP_009109643 - P-loop NTPase
YP_009109643 - replication-associated protein
AXH74443 - CDS
AXH74443 - putative viral replication protein
AXH74443 - RNA helicase
AQU11749 - CDS
AQU11749 - replication protein
AQU11749 - RNA helicase
YP_009126932 - CDS
YP_009126932 - P-loop NTPase
YP_009126932 - replication-associated protein
AXH73382 - CDS
AXH73382 - P-loop NTPase
AXH73382 - putative viral replication protein
YP_009126898 - CDS
YP_009126898 - P-loop NTPase
YP_009126898 - replication-associated protein
AWB80902 - P-loop NTPase
AWB80902 - Rep CDS
AWB80902 - replicase protein
ATD53351 - ORF1 CDS
ATD53351 - Rep Protein
AVA16996 - replicase-associated protein
AVA16998 - replicase-associated protein
AVA17000 - replicase-associated protein
ATD53351 - RNA helicase
AVA16996 - RNA helicase
AVA16998 - RNA helicase
AVA17000 - RNA helicase
AVA16996 - start codon not determined CDS
AVA16998 - start codon not determined CDS
AVA17000 - start codon not determined CDS
YP_004376332 - CDS
YP_004376332 - putative replication protein
YP_004376332 - RNA helicase
ATY70087 - Rep CDS
ATY70087 - replication initiator protein
ATY70087 - Viral Rep
YP_003084282 - CDS
YP_003084282 - putative Rep protein
YP_003084282 - Viral Rep
AXH77564 - CDS
AXH77564 - putative viral replication protein
AXH77564 - Viral Rep
CBK25810 - rep CDS
CBK25810 - replication association protein
CBK25810 - Viral Rep
AQU11776 - CDS
AQU11776 - replication protein
AQU11776 - Viral Rep
AQU11773 - CDS
AQU11773 - replication protein
AQU11773 - Viral Rep
AEI54346 - CDS
AEI54346 - rep protein
AEI54346 - Viral Rep
AGA18409 - CDS
AGA18409 - hypothetical protein
AGA18409 - Viral Rep
AXH75991 - CDS
AXH75991 - putative viral replication protein
AXH75991 - Viral Rep
YP_009163927 - CDS
YP_009163927 - putative replication initiation protein
YP_009163927 - Viral Rep
AEM05804 - CDS
AEM05804 - REP Protein
AEM05804 - Viral Rep
AXH74683 - CDS
AXH74683 - putative viral replication protein
AXH74683 - Viral Rep
AXH76451 - CDS
AXH76451 - helicase Protein
AXH76451 - RNA helicase
AGA18245 - CDS
AGA18245 - hypothetical protein
AGA18245 - RNA helicase
YP_009021888 - CDS
YP_009021888 - P-loop NTPase
YP_009021888 - replication associated protein
ALE29635 - CDS
ALE29635 - replication associated protein
ALE29635 - RNA helicase
YP_009237530 - CDS
YP_009237530 - replication associated protein
YP_009237530 - RNA helicase
YP_009115538 - CDS
YP_009115538 - P-loop NTPase
YP_009115538 - replication-associated protein
AWW06057 - CDS
AWW06057 - helicase Protein
AWW06057 - RNA helicase
AGA18441 - CDS
AGA18441 - hypothetical protein
AGA18441 - P-loop NTPase
YP_009116906 - CDS
YP_009116906 - P-loop NTPase
YP_009116906 - replication-associated protein
AGA18448 - CDS
AGA18448 - hypothetical protein
AGA18448 - RNA helicase
AMH87735 - CDS
AMH87735 - P-loop NTPase
AMH87735 - replication-associated protein
YP_009126890 - CDS
YP_009126890 - replication-associated protein
YP_009126890 - RNA helicase

AXH76508 - CDS
AXH76508 - helicase Protein
AXH76508 - RNA helicase
AXH73061 - TIP49
AXH73061 - CDS
AXH73061 - putative viral replication protein
AXH73061 - RNA helicase
YP_009163927 - CDS
YP_009163927 - P-loop NTPase
YP_009163927 - putative replication initiation protein
AXH75991 - CDS
AXH75991 - P-loop NTPase
AXH75991 - putative viral rplication protein
AQR57898 - P-loop NTPase
AQR57898 - Rep CDS
AQR57898 - replicase Protein
ARO38300 - P-loop NTPase
ARO38300 - Rep CDS
ARO38300 - replicase Protein
AIF76269 - CDS
AIF76269 - P-loop NTPase
AIF76269 - Rep Protein
YP_009126879 - CDS
YP_009126879 - P-loop NTPase
YP_009126879 - replication-associated protein
YP_009116902 - CDS
YP_009116902 - P-loop NTPase
YP_009116902 - replication-associated protein
YP_009109675 - CDS
YP_009109675 - P-loop NTPase
YP_009109675 - replication-associated protein
AIF34802 - CDS
AIF34802 - P-loop NTPase
AIF34802 - replication-associated protein
AJD20393 - CDS
AJD20393 - Gemini AL1 M
AJD20393 - replication associated protein
YP_006281010 - CDS
YP_006281010 - P-loop NTPase
YP_006281010 - putative viral replication protein
AVH76405 - CDS
AVH76405 - Parvo NS1
AVH76405 - putative Rep Protein
AVH76405 - RNA helicase
KT945154 (modified) - Hypothetical 2 translation - Hypothetical 2
AXH76879 - CDS
AXH76879 - P-loop NTPase
AXH76879 - putative viral replication protein
ALE29847 - CDS
ALE29847 - replication associated protein
ALE29847 - RNA helicase
YP_009001747 - CDS
YP_009001747 - replication-associated protein
YP_009001747 - RNA helicase
AQU11773 - CDS
AQU11773 - P-loop NTPase
AQU11773 - replication protein
YP_009021245 - CDS
YP_009021245 - replication-associated protein
YP_009021245 - RNA helicase
AXH78100 - CDS
AXH78100 - helicase Protein
AXH78100 - RNA helicase
AQU11776 - CDS
AQU11776 - P-loop NTPase
AQU11776 - replication protein
AXH75780 - CDS
AXH75780 - replication-associated protein
AXH75780 - Viral Rep
AIF34818 - CDS
AIF34818 - replication-associated protein
AIF34818 - Viral Rep
AIX11626 - Rep CDS
AIX11626 - replicase Protein
AIX11626 - RNA helicase
AOV86234 - DNA pol3 delta2
AOV86234 - CDS
AOV86234 - putative rep protein
AOV86234 - RNA helicase
YP_009237586 - CDS
YP_009237586 - P-loop NTPase
YP_009237586 - replication associated protein
AXH73393 - CDS
AXH73393 - putative viral replication protein
AXH73393 - RNA helicase
AXH76040 - CDS
AXH76040 - putative viral replication protein
AXH76040 - RNA helicase
AVX29443 - CDS
AVX29443 - replication initiator protein
AVX29443 - RNA helicase
AXH73290 - CDS
AXH73290 - putative viral replication protein
AXH73290 - Viral Rep
AXH75585 - CDS
AXH75585 - putative viral replication protein
AXH75585 - Viral Rep
AGS47835 - CDS
AGS47835 - P-loop NTPase
AGS47835 - replication-associated protein
AXH73792 - CDS
AXH73792 - putative viral replication protein
AXH73792 - Viral Rep
AIF34798 - CDS
AIF34798 - P-loop NTPase
AIF34798 - replication-associated protein
AEL22996 - CDS
YP_004778177 - CDS
AEL22996 - P-loop NTPase
YP_004778177 - P-loop NTPase
AEL22996 - Rep protein
YP_004778177 - Rep protein
AJD07493 - CDS
AJD07493 - replication-associated protein
AJD07493 - RNA helicase
AGA18388 - CDS
AGA18388 - hypothetical protein
AGA18388 - P-loop NTPase
AUW34331 - Rep CDS
AUW34331 - replication-associated protein
AUW34331 - RNA helicase
YP_009163936 - CDS
YP_009163936 - P-loop NTPase
YP_009163936 - putative replication initiation protein
AGA18473 - CDS
AGA18473 - hypothetical protein
AGA18473 - P-loop NTPase
AJM89742 - CDS
AJM89742 - P-loop NTPase
AJM89742 - replication associated protein
YP_009237516 - CDS
YP_009237516 - replication associated protein
YP_009237516 - RNA helicase
AXH77740 - CDS
AXH77740 - P-loop NTPase
AXH77740 - putative viral replication protein
AXH74140 - CDS
AXH74140 - helicase Protein
AXH74140 - RNA helicase
BAP81877 - Rep CDS
BAP81877 - RNA helicase
BAP81877 - rolling circle replication initiator protein
YP_009109660 - P-loop NTPase
YP_009109660 - replication-associated protein
YP_009109660 - start codon not determined CDS
AJD07478 - CDS
AJD07478 - replication-associated protein
AJD07478 - RNA helicase
AXH75674 - CDS
AXH75674 - P-loop NTPase
AXH75674 - Rep Protein
YP_009109670 - CDS
YP_009109670 - P-loop NTPase
YP_009109670 - replication-associated protein
AIF76259 - CDS
AIF76259 - Rep Protein
AIF76259 - RNA helicase
AMD39533 - AAA 16
AMD39533 - replication-associated protein
AMD39533 - V1 CDS
YP_764455 - AAA 16
YP_764455 - ORFV1; putative replicase CDS
YP_764455 - rep protein
AGA19549 - C1 CDS
AGA19549 - Gemini AL1
AGA19549 - Rep Protein
YP_009458619 - rep CDS
YP_009458619 - replication-association protein
YP_009458619 - RNA helicase
