## Supplementary material for "Unveiling Crucivirus Diversity by Mining Metagenomic Data": Supp. Table 4

### Supplementary Table 4: Crucivirus Rep motifs

| | Endonuclease domain | | | Helicase domain | | | |
|---|---|---|---|---|---|---|---|
| | Motif I | Motif II | Motif III | Walker A | Walker B | Motif C | Arg finger |
| **CruV-081** | FFTWNN | KHLQG | KANAYCCK | EMGENGKTVFCK | IFHFP | VFSN | LSQDRW |
| **CruV-083** | QPRISK | | | GRTGTGKSRRAW | VIDEF | ITSN | ALRRRL |
| **CruV-084** | QPRISK | | | GRTGTGKSRRAW | VIDEF | ITSN | ALRRRL |
| **CruV-086** | TFTINN | PHIQG | NNANYCSK | | | | ETVKKY |
| **CruV-087** | DFTSSC | THFQG | LNMFYVLK | PKGGKGKTHVCR | LVDIP | VFTN | LSSDRW |
| **CruV-088** | IFTINN | PHIQG | QNYNYCSK | EDGNKGKSYLCK | LLDIP | AFSN | MSSDRW |
| **CruV-089** | IATIPK | QHWQI | AAKDYVWK | GPTGTGKSHNAW | IIDEF | ILSN | ALARRL |
| **CruV-090** | LLTYPG | DHTHC | NQIAYIAK | PKGGAGKTWLAR | ILDLP | VFAN | MSADRW |
| **CruV-091** | LLTYPG | DHTHC | NQIAYIAK | PKGGAGKTWLAR | ILDLP | VFAN | MSADRW |
| **CruV-092** | CFTVFD | HHWQC | DNIAYCTK | PNGNAGKTTMSK | FNFV | CFAN | MGSYKW |
| **CruV-093** | TGTWNN | PHIQF | ACMKYCSK | HEGNSGKTTFAK | MFDLT | IFAN | LSLDRW |
| **CruV-094** | DLTIPC | LHYQC | GNEFYVTK | TIGQRGKTFLTM | FIDLP | VFTN | LSIDRW |
| **CruV-095** | VFTWNN | PHIQG | KDQDYCKK | GPTATGKTSRAR | IIDDL | ITSC | QLYRRI |
| **CruV-096** | DFTAPA | IHYQG | GNDWYVTK | EKGNEGKSILCQ | TIDMP | VFTN | LSMDRW |
| **CruV-097** | CWTSYE | NHWQG | ENRIYCG | GPTGTGKSLKAR | IIDDF | ITSP | QFERRF |
| **CruV-098** | SWTSYE | KHWQG | ENRIYCGA | GPTGCGKSFKAY | LINDF | ITSS | QFQRRF |
| **CruV-099** | FFTWNN | PHIQG | AAENYCSK | PLTCQGKTQLTK | IFNFA | IFSN | LSKDRW |
| **CruV-100** | VFTAWD | VHWQG | QARDYTMK | GPSGTGKSRLAE | IVNEF | FTTP | QLLRRI |
| **CruV-101** | MLTYKT | KHTHV | NFVLYIAK | KKGCVGKTVLCK | LINLS | VFAN | LSLDRW |
| **CruV-102** | AFTLFY | KHLQG | QNIKYCSK | PTGDIGKSELCK | VIDIP | VFSN | LSKDRW |
| **CruV-103** | FLTYAQ | RHFHA | ATATYIQK | | SRDEF | | |
| **CruV-104** | FLTYPK | PHFHV | ACAAYLQK | | TQDEF | | |
| **CruV-105** | FLTYPK | PHFHV | ACAAYLQK | | TQDEF | | |
| **CruV-106** | RFTWNN | PHLQG | ANIAYAQK | EKGNQGGKSWIAQ | IIDLT | | |
| **CruV-107** | | | | | DSDDF | | |
| **CruV-108** | FFTWNN | PHIQG | RAQEYCCK | PIGNNGKTFFMK | IFDLA | VFSN | MSADRW |
| **CruV-109** | FFTLNN | PHVQG | ACIKYCKK | GNAGTLKTRTAF | IIDDF | ITCE | QVNRRI |
| **CruV-111** | DFTLYD | LHYQG | DNNFYVCK | KNGNIGKSSICR | LIDMP | VFTN | LSKDMW |
| **CruV-112** | FFTFNN | PHLQG | KADEYCEK | KSGNNGKTEFCK | LFDFP | VMSN | MSIDRW |
| **CruV-113** | FITYPQ | PHIHA | NSKAYVCK | GAPDFGKTYAIE | IYDDI | VVTN | AFESRF |
| **CruV-115** | FITIKG | | QSICYGTK | KIGGNGKSFFSR | FIDLP | IFSN | LSKDRW |
| **CruV-116** | FLTWH | RHFQG | ASKKYCTK | KIGGRGKSFLAK | LFDVA | VFAN | LSADRW |
| **CruV-117** | DFRANE | VHWQG | ESFFYAMK | IDGNNGKSFISA | FCDMP | VFTN | LSRDRW |
| **CruV-119** | DMRLSQ | VHFQI | KNFNYVLK | FEGNNGKSTIAS | LLDMP | VFTN | LSKDRW |
| **CruV-120** | GCTWNN | PHLQF | VASNYCKK | GPAGYGKSHEAK | IVDDF | VTSN | PIEQRF |
| **CruV-121** | LLTWPL | EHTHT | NAVKYLSK | EEGGKGKSRLAR | IFDLP | IFAN | LSADRW |
| **CruV-122** | CFTYNN | PHLQG | KAIAYCKK | GEPGSGKSQLAR | VLDDI | ITTL | QLLRRI |
| **CruV-123** | CITKHL | PHIHI | QADDYVKK | GKAGLGKSRLAH | LIEDL | VTSD | QLTRRI |
| **CruV-124** | LLTYST | EHTHV | DAKQYIAK | TVGLCQKTDLTR | IIDLP | VLAN | LSFDKW |
| **CruV-125** | CFTQNF | PHWQG | AAREYCMK | GPTGCGKSHDAF | VFDEF | ISSR | EVLRRI |
| **CruV-126** | MVTIRE | RHWQV | AAKDYVWK | GDTGTGKSRRAW | VFDEF | ITSN | ALMRRL |
| **CruV-127** | QITVFD | EHWHI | ENIDYVKK | GPSGVGKTERAK | IYDEW | ITSV | QWERRI |
| **CruV-128** | LFTWNN | AHLQG | QNKTYCSK | SAGGVGKSAMYK | TFDFA | CFAN | LTLSRW |
| **CruV-129** | CFTWNN | PHVQG | ANLAYCSK | GTTGVGKSYDAL | IIDDF | ATTC | QLYRRI |

| | Endonuclease domain | | | Helicase domain | | | |
|---|---|---|---|---|---|---|---|
| | Motif I | Motif II | Motif III | Walker A | Walker B | Motif C | Arg finger |
| CruV-130 | CWTLNN | PHLQG | DAVNYCKK | GSSGTGKSHKAR | IIDDF | VTSI | QLLRRI |
| CruV-131 | LFTLNN | PHLQA | INWLYCSK | GESGLGKTQYAC | VFDDC | FTIN | AIDSRV |
| CruV-132 | AFTAWY | KHIQG | GNIKYCKK | GKTGAGKSYSAF | IIDEF | ITSN | QLLRRI |
| CruV-133 | CFTLNN | PHIQG | DASNYCKK | GATSTGKSETSH | MIDEF | ITSC | QLVRRC |
| CruV-134 | CFTINN | RHIQG | RYLYCMK | | | | |
| CruV-135 | VLTIWM | IHIQA | DAREYCMV | GETDLGKTRFVF | LLDEF | ITSN | ALMRRI |
| CruV-136 | AFTLNN | PHHQG | QNYNYCSK | GKTNVGKSHLAM | VLDEL | ITSA | QLGRRI |
| CruV-137 | CFTLNN | PHLQG | QASEYCKK | GLAGVGKSRGAF | VIDDF | ITCE | QVMRRI |
| CruV-138 | CFTYWG | LHHQG | QNIKYCSK | GPTGCSKTHSAY | IIDDF | VTSC | QLLRRI |
| CruV-139 | CFTSYK | THIQG | QNINYCKK | GEPGTGKTRAVW | LIDDF | FTSN | AFFRRV |
| CruV-140 | FFTWNN | PHIQG | KAEIYCSK | EKTNMGKTQLCK | IFNIS | IFAN | MSIDRW |
| CruV-141 | LLTYKT | LHTHV | GALKYLSK | SLGRKGKTQFGK | IINLT | LTCN | LSQDRW |
| CruV-142 | TFVSWE | LHWQG | QNRMYCCK | GDAGSGKTEWCR | LFDDF | ITCS | QLYRRI |
| CruV-143 | FFTYNN | KHLQG | SANMYCCK | KSGNIGKTVFTK | IFHYS | VFSN | LTLDRW |
| CruV-144 | VFTFNN | PHYQG | KAAAYCMK | GPTGTGKTHRAR | VIDEY | ITCP | EITRRL |
| CruV-145 | CFTINN | PHIQG | QNIDYCSK | GPTGTGKSALAG | VIDDF | VTSP | QLLRRI |
| CruV-146 | CFTVNN | PHLQG | HNLAYCTK | GPSGAGKSYWCH | YLDDL | VTSQ | ALLRRF |
| CruV-147 | CFTVNN | PHLQG | HNLAYCTK | GPSGAGKSYWCH | YLDDL | VTSQ | ALLRRF |
| CruV-148 | CFTVNN | PHLQG | HNLAYCTK | GPSGAGKSYWCH | YLDDL | VTSQ | ALLRRF |
| CruV-149 | CYTYWG | RHIQG | QAATYCMK | GGPGTGKSAAAD | IFDDF | FTAN | PFMRRV |
| CruV-150 | SLTINN | EHYQL | ALEKYVKK | | | | |
| CruV-151 | CFTINH | EHLQC | QASDYCKK | GESGVGKSKSVR | LIDDL | ITSQ | AITRRF |
| CruV-152 | CFTVNN | PHLQG | HNLAYCTK | GPSGAGKSYWCH | YLDDL | VTSQ | ALLRRF |
| CruV-153 | TFTWNN | PHLQG | ANRLYCMK | TVGGMGKSWMTD | VFDFS | CFAN | MSQDRW |
| CruV-154 | CFTLNN | PHLQG | SAAAYCKK | GPAGAGKTAGPV | IIDDF | ITCD | QVIRRI |
| CruV-156 | CYTYWG | RHIQG | QAADYCMK | GGPGTGKSAAAA | IFDDF | FTAN | PFMRRV |
| CruV-157 | DFRHTA | VHFQG | DDFFYTMK | DSGNNGKSTIAS | FVDMP | VFSN | LSRDRW |
| CruV-158 | VFTLNN | PHLQG | SNKTYCSK | EKGSKGKTFLAK | IYDVP | IFTN | FSVDRL |
| CruV-159 | CFTLNN | PHLQG | QASNYCKK | GATGVGKSRGAR | IIDDF | ITCE | QVLRRI |
| CruV-160 | EFRLSE | RHFQG | NYDCYASK | NIGNNGKSTIAH | FVDLP | VFTN | VSRDRW |
| CruV-161 | DFTYHS | LHFQG | TSFDYVMK | PKGNLGKTIFRK | ILDFA | IFTN | MSDDRW |
| CruV-162 | CWTLNN | PHLQG | LAIRYCVK | ENGNTGKSELTK | IIDIP | VFSN | LSLDRW |
| CruV-163 | DFTAHG | IHLQG | KNSYYVSK | KSGNIGKSTLCT | LFDLP | IFTN | LSADRW |
| CruV-164 | CWTLNN | PHLQG | LAIRYCVK | ENGNTGKSELTK | IIDIP | VFSN | LSLDRW |
| CruV-165 | DFRANQ | LHWQG | DLFFYVMK | PTGGVGKSTIAA | FLDMP | VFTN | LSRDRW |
| CruV-166 | CFTINN | PHIQG | RAAAYCQ | | | | |
| CruV-167 | CYTYWG | RHIQG | QAADYCMK | GGPGTGKSAAAD | IFDDF | FTAN | PFMRRV |
| CruV-168 | TFTLYD | LHYQG | HNYNYCTK | KESETGKTIFKQ | | VTTN | LLPKRI |
| CruV-169 | DFRANQ | LHWQG | DLFFYVMK | PTGGVGKSTIAA | FLDMP | VFTN | LSRDRW |
| CruV-170 | | VHFHV | | GPPDIGKTHWIN | VYDDV | IFLN | LFVARF |
| CruV-171 | VFTWNN | PHLQG | ANLKYCSK | KDGNTGKSDMLK | VFDYS | CFSN | LSQDRW |
| CruV-172 | DFTLFD | IHYQG | DNDFYVLK | ENGNVGKTTLVR | FVDMP | IFTN | LSKDMW |
| CruV-173 | | PHYHI | DNIHYIKK | GPSGVGKTTKAM | LYDEW | ITSV | QWERRI |
| CruV-174 | VIVTNN | PHLQG | SQRAYCTK | GEAGVGKSVKAR | | | |
| CruV-175 | VIVTNN | PHLQG | SQRAYCTK | GEAGVGKSVKAR | | | |
| CruV-176 | VFTINN | KHLQG | QNVKYCTK | KNGNSGKSKIVK | LLDIS | IFSN | MSLDRW |

| | Endonuclease domain | | | Helicase domain | | | |
|---|---|---|---|---|---|---|---|
| | Motif I | Motif II | Motif III | Walker A | Walker B | Motif C | Arg finger |
| CruV-177 | DFTYVK | EHYQG | KNFNYVLK | INGNNGKSYLSE | IMDMP | LFCN | LSKDRM |
| CruV-178 | DMTWHQ | LHYQC | GNLNYVTK | SKGVVGKSTLIN | LIDIP | VFTN | LSNGRW |
| CruV-179 | CFTAYH | PHLQG | EAIDYCKK | GGPGSGKSHHCK | ILDEF | ITSN | PLIRRI |
| CruV-180 | CFTAYH | PHLQG | EAIDYCKK | GGPGSGKSHHCK | ILDEF | ITSN | PLIRRI |
| CruV-181 | DFTLRR | EHFQG | SNFNYVLK | PAGNVGKSTLSG | FLDLP | VFTN | LSYDRW |
| CruV-182 | VFTSFN | KHLQG | QAIDYCKK | GDTQTGKTKYVY | LLDDF | ITSN | ALKRRI |
| CruV-183 | RFTINN | FHFQG | SNVVYASK | GPSGTGKTHRAT | IIDDL | ITAP | QLLRRI |
| CruV-184 | CFTMFF | LHYQG | QNVKYCSK | GPTGVGKSKLAF | IFDDF | ITSY | QLGRRI |
| CruV-185 | QLTVFD | EHWHI | DNIHYIKK | GPSGIGKTQRAK | VYDEW | ITSV | QWQRRI |
| CruV-186 | CFTAWD | FHWQG | QASEYCKK | HNSGTGKSTLID | TFNIP | VFAN | LMPKRC |
| CruV-187 | CFTIHD | LHVQG | QNKEYCSK | GPPRTGKSLLVR | VFDEF | FTSN | IRGPRF |
| CruV-188 | DFRISK | IHYQG | RTFNYQLK | PVGNNGKSTIKD | LVDIP | IFTN | LSPDRM |
| CruV-189 | MIRISR | DHIQG | ALSKYSMK | KIGNTGKSKFCK | LFNLT | VFAN | MSLDRW |
| CruV-190 | VFTYNN | KHLQG | QAITYCKE | DNGRSGKSELVA | LFDLP | IFAN | LTMDKW |
| CruV-191 | TWTIFD | IHLQG | QNKVYCSK | GPPNIGKSGLAR | VFGEF | FLSN | IRGPRF |
| CruV-192 | VFVLPN | KHLQG | ANIAYCKK | PEGNQGKSWMCN | VFDFS | CFSN | MSQDRW |
| CruV-193 | CFTNFA | LHHQG | QNVDYCSK | LTGNLGKSFFST | IFDFE | AMSN | FSIDRW |
| CruV-194 | VFTSWC | KHWQG | QASSYCMK | LKSATGKSTMRD | IFDLP | VFAN | EMPERC |
| CruV-195 | VFTLNN | PHLQG | QAVDYCKK | GPTGTGKSRTAR | LIEDV | ITSN | PILRRF |
| CruV-196 | CFTWNN | PHLQG | SNMTYCTK | KEGKTGKTEFAK | IYPLS | IFAN | MSKDRW |
| CruV-197 | CFTNFA | LHHQG | QNVEYCSK | ITGNLGKSFFSD | VFDFE | CLSN | FSLDRW |
| CruV-198 | PWTLNN | PHLQG | QNIDYIVL | GPSGIGKTYAAE | FLDEF | ITTP | QLSRRI |
| CruV-199 | FITYFA | IHFHC | QAIAYILK | | | | |
| CruV-200 | CITINN | LHIQG | QAIDYCKK | GESGCSKSHTAK | IIDEL | ITTS | QLIRRI |
| CruV-201 | LLTYKT | EHTHV | DTFFYLTK | DVGGSGKTNLSC | MIDIP | ILAN | IKWDRW |
| CruV-202 | CFTLNN | PHLQG | ERYLYCMK | | | | |
| CruV-203 | VFTFNN | PHLQG | QCRTYCSK | GPTGSGKSRGVR | LLDDV | VTSQ | ALERRF |
| CruV-204 | CFTFNN | PHLQG | QNTAYCSK | GETGTGKSHCVE | YLEDF | VTSN | PLERRF |
| CruV-205 | CFTFNN | PHLQG | QNTAYCSK | GETGTGKSHCVE | YLEDF | VTSN | PLERRF |
| CruV-206 | VFTWNN | PHLQG | ASLKYCSK | KEGNTGKSDMLK | VFDYS | CFSN | LSQDRW |
| CruV-207 | CFTFNN | KHLQG | QAIAYCKK | GEPEAGKSFYAR | ILDDI | ITTL | QFTRRI |
| CruV-208 | LGTIPR | EHWQI | AAEDYVWK | GKTGTGKSRRAW | IIDEF | ITSN | ALMRRL |
| CruV-209 | MFTYAT | LHTHA | QKVLYLCK | IKGCCGKTIFAQ | LEDVG | VLAN | LTWDKW |
| CruV-210 | LLTYRT | LHTHV | DASKYIAK | GNGNDGKSLGLC | IIDLP | VMAN | LSLDRW |
| CruV-211 | CYTINN | PHLQG | EASDYCKK | GKAGVGKTKLAH | IIDDY | ITCE | QITRRI |
| CruV-212 | FFTFNN | THLQG | ASINYCSK | GNGNDGKSLGLC | ILDLP | IFTN | LTLDRW |
| CruV-213 | TFTLNN | KHVQG | QNYEYCSK | GKSGSGKSREAR | IIDDV | VTSQ | ALKRRF |
| CruV-214 | CCTDYV | EHYQV | QNKRYCSK | GPPGTCKSRTAI | LFDDF | FTSN | AWIRRL |
| CruV-215 | CFTHNN | PHLQG | SNKTYCSK | PEGNVGKSDMAK | VIDLA | CFAN | LSADRW |
| CruV-216 | DLSYNK | EHYQI | KNFNYVMK | KNGNNGKSYLSE | IMDMP | VFCN | LSKDRM |
| CruV-217 | CFTLNN | PHHQG | DNYDYCSK | GPTGVGKTHLAH | IFDDI | VTTC | QLGRRI |
| CruV-218 | CFTINN | PHIQG | QNRTYCSK | GGARLGKSKLAH | LWDEF | ITSI | QIMGRI |
| CruV-219 | VFTSWC | KHWQG | QAAQYCKK | QKSGTGKSTMRD | VFDLP | VFAN | DMPMRC |
| CruV-220 | CFTINN | PHIQG | QASDYCKE | GPTGTGKTRTAM | IIDDF | ITCP | QLTRRI |
| CruV-221 | DFRCNE | IHWQG | DLFFYCMK | PTGGKGKSTIAA | FFDMP | VFSN | LSRDRW |
| CruV-222 | IFVYNN | PHLQG | EAAAYCKK | GETGVGKTRGAI | IIDDF | ITCD | QLTRRI |

| | Endonuclease domain | | | Helicase domain | | | |
|---|---|---|---|---|---|---|---|
| | Motif I | Motif II | Motif III | Walker A | Walker B | Motif C | Arg finger |
| CruV-223 | FLTSYA | PHSHT | QAIEYVVK | ALRHMGMIERAQ | | | |
| CruV-224 | CWTLNN | PHLQG | QARDYCLK | TKGGRGKSYLAK | VFDYS | CFSN | |
| CruV-225 | DVTIWE | RHFQI | RNFNYVLK | PTGNYGKGFAAD | ILDMP | VFTN | YSKDRF |
| CruV-226 | DFTLFD | IHFQG | TNDFYVTK | PEGCKGKTALTR | MIDMP | VFTN | MSKDMW |
| CruV-227 | CFTFNN | RHLQG | QNDDYINK | GNTGTGKSHAVE | YLEDI | VTSN | PLQRRF |
| CruV-228 | ILTINN | VHIQA | AAAAYCIK | GDGGVGKTKYVI | CIDDF | ITCE | QVMRRI |
| CruV-229 | VFTINN | LHIQG | DSINYTKK | GKANTGKTYKAR | VMEDF | FSTN | CENGRI |
| CruV-230 | VFTWNN | PHHQG | QNFNYSNK | GPSGAGKTRWCY | IVDDF | FTSC | QLLRRI |
| CruV-231 | CFTMFW | EHYQG | QNVKYCAK | GPTGVFKTRYAS | IFDDF | ITSC | QLGRRI |
| CruV-232 | IPTIWI | LHAHL | | | | ICFS | |
| CruV-233 | VFTWNN | PHLQG | ASLKYCSK | KRGNTGKTAMFK | IFDLA | CFSN | LSIDRW |
| CruV-234 | CYTINN | PHLQG | EASDYCKK | GKAGVGKTKLAH | IIDDY | ITCE | QITRRI |
| CruV-235 | CFTMYW | EHYQG | QNVHYCAK | GPTGTGKSKMAF | IFDDF | ITSC | QLGRRI |
| CruV-236 | CFTYNN | PHLQG | ENIKYCSK | LVGNKGKTAMTK | IFDFS | IFAN | LSKDRW |
| CruV-237 | CFTHHI | PHYQG | EASEYCKE | GVMDAGKTTIIR | FMDEV | VASN | AIKKRF |
| CruV-238 | DFTVWP | LHFQG | KNFFYVMK | GVGNNGKTICCQ | IFDFP | LFMN | LSADRW |
| CruV-239 | DFTVWP | LHFQG | KNFFYVMK | GVGNNGKTICCQ | IFDFP | LFMN | LSADRW |
| CruV-240 | DFTLDV | LHYQG | NSFIYVMK | AQPNLGKTHATQ | FIDMP | VFTN | LGDDRW |
| CruV-241 | DFTYFG | LHYQG | DNNFYVCK | KLGNKGKSYLSF | IFDMP | IFTN | LSEDRW |
| CruV-242 | AITISR | PHFQG | ALKKYCMK | QEGNQGKSEFVK | LVDLT | VMAN | LSNDRW |
| CruV-243 | LYTWFN | PHVQG | NNFEYCSK | GPSNTGKTHYAA | IFDDM | FTHN | AIERRF |
| CruV-244 | CFTSFN | EHFQG | QARVYCMK | DAGNNGKTFLSK | ICDYA | VMAN | LSVDRI |
| CruV-245 | LYTWFN | PHVQG | NNFEYCSK | GPSNTGKTHYAA | IFDDM | FTHN | AIERRF |
| CruV-246 | FFTYKT | EHTHA | GSIAYCIK | KAGGKGKSTFLD | INLPR | IFAN | MSLRKW |
| CruV-247 | CFTINN | HHFQC | SNIVYCTK | GPTGTGKTYHVM | LIDDL | VTSN | TLWRRF |
| CruV-248 | CFTYNN | PHLQG | ENIKYCSK | LVGNKGKTAMTK | IFDFS | IFAN | LSKDRW |
| CruV-249 | VFTHHC | RHFQG | GSLDDCLN | GDGDLGKSDLCK | IFDFP | CLAN | YTSNRW |
| CruV-250 | CFTLNN | PHVQG | QNYEYCSK | GESGAGKSHKSR | ILEDV | VTSQ | ALHRRF |
| CruV-251 | CFTLNN | PHVQG | QNYEYCSK | GESGAGKSHKSR | ILEDV | VTSQ | ALHRRF |
| CruV-252 | CFTMFW | EHYQG | QNVRYCAK | GPTGTGKSKIAF | IFDDF | ITSC | QLGRRL |
| CruV-253 | CFTLNN | PHLQG | QASEYCKK | GLAGVGKTRHIY | IIDDF | ITCE | QIMRRI |
| CruV-254 | CFTMYW | EHYQG | QNVRYCAK | GPTGTGKSKLAF | IFDDF | ITSC | QLGRRL |
| CruV-255 | FLTINN | KHLHC | DCIVYCQK | GASGTGKTRCIF | LWDEI | ITSS | EIMRRL |
| CruV-256 | CFTAYP | QHHQG | DAAQYCWK | GKAGTGKTRLAL | IIDDF | ITSC | QLLRRI |
| CruV-257 | VYTYFD | LHLQG | QASTYCKE | GLAGTGKSRTAK | VIDDF | ITCE | QIMRRL |
| CruV-258 | CFTVNN | QHLQC | QAADYCKK | GDSGVGKSMSVR | LIDDL | ITSQ | AISRRF |
| CruV-259 | TLTIWD | QHFQA | HNNIYTSK | GKSGCGKTHQAQ | IFDEF | FTSN | AFQRRI |
| CruV-260 | FYTFNN | PHLQG | QAANYCLK | GTTGTHKSYLSR | ILEDF | INTP | QLLRRI |
| CruV-261 | WLTYKT | EHTHC | RIMRYMAK | ETGNVGKSWMAL | IFDFP | VLAN | MSLDRW |
| CruV-262 | CFTMFW | EHYQG | QNVKYCSK | GPAGVGKTQSAL | IIDDF | ITCD | QIGRRI |
| CruV-263 | CFVINN | PHIQG | QNRAYCSK | GDTGAGKSRLGH | LWDEF | ITSI | QILRRI |
| CruV-264 | LYTWFN | PHVQG | NNFEYCSK | GPSNTGKTHYAA | IFDDM | FTHN | AIERRF |
| CruV-265 | CFTLNN | PHVQG | QNYEYCSK | GESGVGKSHKAR | ILEDV | VTSQ | ALGRRF |
| CruV-266 | CFTLNN | PHVQG | QNYEYCSK | GESGVGKSHKAR | ILEDV | VTSQ | ALGRRF |
| CruV-267 | CFTIFG | PHLQG | NNVIYCSK | AQGDTGKSFFVK | CIDYA | CFAN | LSMDRW |
| CruV-268 | VFTAWD | GHWQG | QAREYALK | GPTSTGKTRLVM | LVDDF | ITSP | QLLRRI |

| | Endonuclease domain | | | Helicase domain | | | |
|---|---|---|---|---|---|---|---|
| | Motif I | Motif II | Motif III | Walker A | Walker B | Motif C | Arg finger |
| CruV-269 | IFVKHY | EHLQG | QASDYCRG | VVGSAGKSELAK | AYDLP | VFAN | MSADRW |
| CruV-270 | FFTYKT | EHTHA | GSIAYCIK | KAGGKGKSTFLD | INLPR | IFAN | MSLRKW |
| CruV-271 | EIEINN | PHIHC | RYVDYCKG | | | | VLHREI |
| CruV-272 | CFTDFV | PHIQG | QAADYCKE | GEPGCGKSHRAV | VLDDF | ITTN | QWLRRV |
| CruV-273 | TFTLFY | PHLQG | ENNTYCSK | PEGGVGKSDMFK | VFDLA | CFAN | LSRDRW |
| CruV-274 | DLTGWT | LHFQC | KNFDYVMK | KIGGIGKTTLTD | IVDLP | VFTN | ISKFRW |
| CruV-275 | LLTYGS | LHTHI | NSLKYLAK | SIGGSGKSWLID | MNLT | VMAN | LTTDRW |
| CruV-276 | LYTWFN | PHVQG | NNFEYCSK | GPSNTGKTHYAA | IFDDM | FTHN | AIERRF |
| CruV-277 | CFTDFV | PHIQG | QAADYCKE | GEPGCGKSHRAV | VLDDF | ITTN | QWLRRV |
| CruV-278 | LYTWFN | PHVQG | NNFEYCSK | GPSNTGKTHYAA | IFDDM | FTHN | AIERRF |
| CruV-279 | | EHMQC | QASDYCKK | GPPGVGKSHIAR | | VTSN | AIVRRF |
| CruV-280 | AFTWNN | PHLQG | ANLKYCSK | KRGNTGKTAMFK | IFDLA | CFSN | LSIDRW |
| CruV-281 | CFTDFV | PHIQG | QAADYCKE | GEPGCGKSHRAV | VLDDF | ITTN | QWLRRV |
| CruV-282 | CFTDFV | PHIQG | QAADYCKE | GEPGCGKSHRAV | VLDDF | ITTN | QWLRRV |
| CruV-283 | EITIQP | LHFQG | DNNFYVMK | QKGGIGKSKLVE | LIDYP | IFTN | LTKNRW |
| CruV-284 | DFRISQ | LHFQG | QNFNYVLK | PVGNNGKSTVAS | LIDMP | VFTN | LSADRW |
| CruV-285 | VFTNFK | LHHQG | SNNKYTSK | GSAGRGKSHIAK | WFDEF | VTSP | QMLRRI |
| CruV-286 | DLTIYE | RHFQI | KNFNYVMK | PDGNRGKGFIAD | ILDMP | LFTN | YSKDRY |
| CruV-287 | DFTYFI | WHLQG | AFYRYPMK | SHGNTGKTTFKS | VIDIP | VMSN | LSIDRW |
| CruV-288 | VFTAWQ | LHWQG | EAAAYCEK | EASATGKSTMYD | FNVPR | VFAN | DMPDRC |
| CruV-289 | CFTLNN | PHLQG | QASEYCKK | GATGVGKTRTAR | IIDDF | ITCE | QVIRRI |
| CruV-290 | RWTHNN | PHLQG | ANIVYVRK | PLGKCGKTVFAA | IIDLM | VLSN | MAQERF |
| CruV-291 | CFTDFV | PHIQG | QAADYCKE | GEPGCGKSHRAV | VLDDF | ITTN | QWLRRV |
| CruV-292 | CFTLNN | PHLQG | QASMYCKK | GATGVGKSRTAR | IIDDF | ITCE | QVLRRI |
| CruV-293 | MFTYYI | PHLQF | QNYKYCSK | GPPGVGKSRRAR | LIDDL | VTSN | SFDRML |
| CruV-294 | CFTINN | PHMQC | HNQRYCKK | GTTGSAKTREYT | LSRLR | ITSC | QLHRRY |
| CruV-295 | CFTMFW | DHYQG | DNVKYCSK | GPTGTGKSKFAF | IFDDF | ITSC | QLGRRI |
| CruV-297 | CFTINN | PHIQG | QASDYCKK | GPTGTGKTRTAI | LIDDF | ITCP | QVTRRI |
| CruV-298 | CWTLNN | PHLQG | QAAEYCKK | GPTGTGKSRRAH | IIDDF | VTSN | ALRRRI |
| CruV-299 | VFTNFK | PHHQG | SNEKYTQK | GESGRGKTYIAK | WFDEF | FTSP | QLKRRI |
| CruV-300 | DATIGE | RHYQC | DNHFYVCK | PDGGIGKSTFCT | LMDMP | IFTN | LTRNRW |
| CruV-301 | CFTVNN | PHLQG | QNRVYCMK | GPTGIGKSRYCF | ILEDV | ITSN | PLLRRF |
| CruV-302 | KFTWNN | PHIQG | NHVRYIQG | LSSTTGKTTFMQ | HIDIP | VTSN | KLPNRF |
| CruV-303 | CFTLNN | PHLQG | EAAMYCKK | GLAGVGKTRKPI | IVDDF | ITCE | QVLRRI |
| CruV-304 | LFTYYI | PHLQF | QNYKYCSK | GPPGVGKSRRAR | LIDDL | VTSN | SILRRF |
| CruV-305 | CFTVNN | PHLQG | QNRVYCMK | GPTGIGKSRYCF | ILEDV | ITSN | PLLRRF |
| CruV-306 | CFTHNN | PHLQG | ANKTYCSK | PEGNVGKSDMAK | VIDLA | CFAN | LSADRW |
| CruV-307 | LTTYWP | LHGHM | QAKDYVTK | | | | |
| CruV-308 | CYTSRE | PHIHV | KWTNYVLK | | | | |
| CruV-309 | CFTVNN | PHLQG | QNRVYCMK | GPTGIGKSRYCF | ILEDV | ITSN | PLLRRF |
| CruV-310 | CFTVNN | PHLQG | QNRVYCMK | GPTGIGKSRYCF | ILEDV | ITSN | PLLRRF |
| CruV-311 | CFTVNN | PHLQG | QNRVYCMK | GPTGIGKSRYCF | ILEDV | ITSN | PLLRRF |
| CruV-312 | DFTSYF | LHLQG | TDDAYVMK | KKGNIGKTTLVG | FIDLP | VFTN | MSSDRW |
| CruV-313 | TWTLNN | PHLQG | QNRNYCGK | GKTGQAKTRLAG | IIDEV | YTSP | QLHRRI |
| CruV-314 | VFTWNN | LHHQG | QNFNYSNK | GPAGIGKTRWCA | IVDDF | FTSC | QLLRRI |
| CruV-315 | IFTWNN | PHHQG | QNFNYSNK | GPAGIGKTRWCA | IVDDF | FTSC | QLLRRI |

| | Endonuclease domain | | | Helicase domain | | | |
|---|---|---|---|---|---|---|---|
| | Motif I | Motif II | Motif III | Walker A | Walker B | Motif C | Arg finger |
| CruV-316 | CFTINN | PHMQC | QNQIYCKK | GTTGSAKTREFT | MNEVR | ITSC | QFYRRY |
| CruV-317 | FITYPK | HLHSI | YAIDYCDK | | | | |
| CruV-318 | CFTAFD | PHYQG | SNEAYTQK | GGAGKGKSTVAR | WFDEF | ITSS | QLLRRI |
| CruV-319 | VFTYNN | PHLQG | ENEDYCSK | GDTGTGKSHAVE | YLEDF | VTSN | PLMRRF |
| CruV-320 | VFTYNN | PHLQG | ENEDYCSK | GDTGTGKSHAVE | YLEDF | VTSN | PLMRRF |
| CruV-321 | VFTINN | PHLQG | NNAIYCSK | KDGNAGKSDMCK | CMDYA | CFSN | LSTDRW |
| CruV-322 | CFTLNN | PHLQG | DNIDYCSK | GVTGVGKSTKYR | ILEDI | VTSQ | ALKRRF |
| CruV-323 | CFTLNN | PHLQG | DNIDYCSK | GVTGVGKSTKYR | ILEDI | VTSQ | ALKRRF |
| CruV-324 | CFTLNN | PHLQG | DNIDYCSK | GVTGVGKSTKYR | ILEDI | VTSQ | ALKRRF |
| CruV-325 | MIRISK | HHIQG | ALEKYSMK | QIGNTGKSKFCK | FNLSR | VMAN | LSSDRW |
| CruV-326 | IFTWNN | PHLQG | TNTSYCSK | GQSGSGKSQFAE | VIEEF | ITSV | QIVRRM |
| CruV-327 | LLTYKT | EHTHT | DMKRYISK | PMGNSGKSTFAK | IFDFP | VFAN | LSFDRW |
| CruV-328 | CFTLNN | PHLQG | QASNYCKK | GPTGSGKTRHAL | VIDDF | ITSC | QLLRRI |
| CruV-329 | DLTISL | DHIQA | SGDFYCCK | EKGNIGKSILVA | MFDMP | IFTN | LSRDRW |
| CruV-330 | FLTYWP | IHAHM | QAIDYCTK | | | | |
| CruV-331 | CFTLNN | PHLQG | SNIAYCTK | GPTGTGKSRAAN | IIDDY | ITTP | QLMRRI |
| CruV-332 | CFTYYS | LHVQG | EAAAYCEK | GPSRCGKSVLAR | FSEFA | FTSN | PWLKRI |
| CruV-333 | VFTINN | PHLQG | NNAIYCSK | HKGNAGKSDMCK | CFDYA | CFAN | LSADRW |
| CruV-334 | CFTINN | FHLQG | QAREYCMK | GEPGCGKTRLAK | LFDDF | ITSN | ALERRI |
| CruV-335 | MLTYKT | EHTHV | KCINYLCK | GGSGIGKTQLAL | VFDDI | FTSN | AIKRRF |
| CruV-336 | CFTIFD | PHLQG | EASEYCKK | GPTGTGKTSMSM | IIDEY | FTSN | PIARRV |
| CruV-337 | VLTVSQ | WHYQG | NLKEYCIK | FDGGKGKSSLCK | FNLSR | VFTN | FSPDRW |
| CruV-338 | DITAPC | LHWQC | GDEFYVIK | PTGQVGKSTLAL | FIDLP | VFTN | LSRDRW |
| CruV-339 | TFTWNN | PHLQG | DSIRYCSK | KDGCAGKSSLID | IVDLS | VMAN | FSLDRW |
| CruV-340 | CITLNN | PHLQG | QNKTYCSK | HWGPTGKFLQEC | | IMSN | |
| CruV-341 | CFTEFD | KHMQG | ENQKYCTK | GKSGSGKTKYVY | LLDDF | ITSN | AVERRI |
| CruV-342 | LLTYKT | EHTHT | DMKRYISK | PMGNSGKSTFAK | IFDFP | VFAN | LSFDRW |
| CruV-343 | CFTLNN | PHLQC | QASEYCKK | GLAGVGKSRGAY | VIDDF | ITCE | QVMRRL |
| CruV-344 | CFTVNL | HHYQG | QAIDYVSK | GVSDSGKTRKVY | LIDDY | YTSN | AFKRRI |
| CruV-345 | CYTWNN | PHIQG | QSLTYCSK | GKTAVGKSHLAF | | ITSS | QLLRRL |
| CruV-346 | CFTLNN | PHLQG | SNIKYCSK | GKSRAAREHNGY | ILDDF | IVTSN | |
| CruV-347 | CFTLNN | PHLQG | QNIDYCIK | GSTGTGKSMKAR | IIDDF | VTSN | PLLRRF |
| CruV-348 | AFTHHT | LHWQT | DNARYCKK | GKGGTGKSRLAR | MIEEF | FTSN | QTERRF |
| CruV-349 | IITSYL | MHHQG | QGAEYCGY | GESGTGKTELIY | LIDDI | LTSN | AIIRRI |
| CruV-350 | SWTLFN | KHLQG | ENRAYCTK | GDTGTGKTLFAR | ILDDI | VTGP | QLLRRC |
| CruV-351 | MLTYPD | EHSHV | NSVRYLGK | AIGGKGKSCFAD | IFDWP | IFAN | LSLDRW |
| CruV-352 | CFTINK | LHIQG | QARDYCMG | GKSGTGKSKLAW | IIDDY | FTTN | TFERRI |
| CruV-353 | IITSYL | MHHQG | QGAEYCGY | GESGTGKTELIY | LIDDI | LTSN | AIMRRI |
| CruV-354 | VFTNWK | LHHQG | SNDKYTSK | GKSGRGKTHIAK | WLDEF | VTSS | QLRRRI |
| CruV-355 | AVTINN | RHLQC | QAAEYCKK | GPTRIGKSTRAR | LIEDI | VTSQ | AIVARC |
| CruV-356 | CFTINQ | LHLQG | QARDYCMK | GPPGTGKTTLIA | IFDDF | FTSN | AIRRRC |
| CruV-357 | AFTWFN | KHLQG | ANIAYCSK | QIGNVGKTHMAM | VFDFT | IFAN | MSLDRW |
| CruV-358 | SFNFFI | EHYQC | NFDYVMK | VGGVHMKSSFEE | IIDIE | VFAN | LTGGRF |
| CruV-359 | QFTLNN | PHLQG | QNITYCKK | NLGNIGKSDTVM | IFDLP | IYAN | LSLDRW |
| CruV-360 | LFVLNN | PHLQG | VNIKYCQK | GKTSTGKSHYIH | FIDDF | ITSP | QLLRRV |
| CruV-361 | VGTLNN | PHLQF | QNKDYCSK | GAAGTGKSMTAR | LIDDF | VTSQ | AIKRRF |

| | Endonuclease domain | | | Helicase domain | | | |
|---|---|---|---|---|---|---|---|
| | Motif I | Motif II | Motif III | Walker A | Walker B | Motif C | Arg finger |
| CruV-362 | DFTLDM | RHWQG | NVFTYIEK | NKGNNGKTWFAG | IIDIP | VFVN | LSADRM |
| CruV-363 | CFTNFD | PHHQG | DNVKYCSE | YDGQLGKSYFQK | VNLTR | IFAN | LSIHRW |
| CruV-364 | CFTDFV | PHIQG | QAADYCKE | GEPGCGKSHRAV | VLDDF | ITTN | QWLRRV |
| CruV-365 | CFTLNN | PHLQG | QNDAYCKK | GLSGCGKSRYAR | IIEDL | ITSQ | ALTRRC |
| CruV-366 | EIRANE | RHYQG | GKAFYSMK | FAGCDGKSTCAS | FFDMP | VFTN | MSKDRW |
| CruV-367 | VFTLNN | PHLQG | KAIDYCKK | GNTGLGKTRFVH | LFDDY | ITSN | ALERRI |
| CruV-368 | | | QARDYTMK | GPTGINKSRNAY | IIDDY | LTSH | ELARRV |
| CruV-369 | CFTDFV | KHLQG | QNIEYCSK | GESGAGKSYPIN | LFDDF | FTAN | AWNRRI |
| CruV-370 | | | | | VFDEF | ILSN | AFLSRV |
| CruV-371 | SSTTIP | PHAMI | SISRYAEV | | | | CVARRV |
| CruV-372 | CYTLNS | LHIQG | QAADYCRD | GESGTGKSQAAY | LFDEF | FTSN | PLTRRI |
| CruV-373 | LVTCNV | KHLQG | EMMMYCKK | GPAGSGKSHYAR | LLDDF | VTSQ | ALNRRF |
| CruV-374 | VFTLHN | PHLQG | HSIIYCKK | GESGTGKSRHAR | IIEDI | ITSN | AIERRY |
| CruV-375 | FLTYKT | KHSHM | KALNYIGK | PPGQGGKTSLLR | VNLTR | VFAN | MSRGRI |
| CruV-376 | HFVYKH | LHTHV | TLVRYLAK | ETGACGKSTFIR | LFDLP | VFAN | LSLDRW |
| CruV-377 | SFTWNK | LHYQG | NAEFYCCK | AEGCTGKSNFAT | IFDVP | VFAN | LSVDRW |
| CruV-378 | VFTLNN | PHLQG | QNKEYTQK | GPAGAGKSRGAR | IMDDF | ITSQ | AIKRRF |
| CruV-379 | CYTLNN | LHCQG | QAADYCKR | GATGVGKSAAAY | IFDEM | FTSN | PFMRRI |
| CruV-380 | VFTCNN | PHLQG | ASIEYCEK | GPTGSGKSRAAR | ILDDV | VTSN | ALHRRF |
| CruV-381 | PLTRFN | FHWQI | QNYKYCTK | GPSGIGKTSWAL | VFDEI | FCSN | AIERRV |
| CruV-382 | CFTIHD | PHLQG | LARHYCMK | GKSRSGKSSSAI | VNEMS | MTTN | IFNRIT |
| CruV-383 | MVTYND | PHLQM | DAAGYCTL | GDGATLKSSCIR | AIDDP | ITAN | PMIRRF |
| CruV-384 | VYTLNN | PHLQG | EAVTYCQK | GNPRTGKSRSAR | LIEDI | ITSN | PLQERF |
| CruV-385 | CVTYNL | RHIQG | QNVRYCSK | GRTGLAKTHGTI | LIDDL | VTSN | ALKRRM |
| CruV-386 | | WHYQC | AAWRYCYN | AEGQSGTTQMFK | CIDIT | IKWN | VSAGRM |
| CruV-387 | CYTLNS | LHIQG | QAADYCRD | GESGTGKSQAAY | LFDEF | FTSN | PLTRRI |
| CruV-388 | CWTHYD | KHMQG | KNEEYVGK | KGGGAGKSACAD | CFDFS | VFAN | LSEDKL |
| CruV-389 | LLTYKT | PHTHA | RVLKYITK | RVGNMGKTVLAK | TIDLP | VLAN | LSLDRW |
| CruV-390 | VFTLNN | LHLQG | KNFTYCTK | GPSGTGKSRYAR | LVEDV | VTSQ | ALKRRF |
| CruV-391 | FLTYSQ | IHIHA | GLVRYLRD | APSGWGKSEWIQ | IFDDI | VLSN | QIKQRF |
| CruV-392 | FLTYAQ | VHRHA | GSLKYVLE | GGTNIGKTTLMN | VFDEF | FLSN | TLMNRL |
| CruV-393 | CYTLNN | PHLQG | QNRAYCTK | GPTGSGKSHAAR | IFDDF | FTAP | QFTRRI |
| CruV-394 | IFTLNN | HHLQG | QAEAYATK | GPTGTGKTKAAA | IIDDY | ITAP | QLGRRL |
| CruV-395 | AFTWFG | PHLQG | ANFRYASK | IAGGQGKTLFSK | IVDLT | VFCN | LSADRV |
| CruV-396 | CITLNN | PHIQG | QNREYCSK | GPTGTGKTFRAI | IFDDF | ITSN | PLYRRI |
| CruV-397 | NFTDPN | PHLQC | ACIEYCGK | GPTGSGKSRWAM | IIDDF | FTTP | QFSRRL |
| CruV-398 | FFTWNN | KHIQG | EALAYCTK | PNGGAGKTVLAT | IWDIP | IFSN | LSKDRW |
| CruV-399 | SFTWNK | LHYQG | EAEFYCMK | ETGRTGKSEFAT | IFDVP | VFSN | LSHDRW |
| CruV-400 | | LHWQV | AVLSYCMK | TTGQAGFTSLMS | IIDIT | IKTN | LSEGRM |
| CruV-401 | AFTMNN | RHLQG | KSIDYCKK | GPTGTGKSSTAR | IMDDI | ITSN | ALNRRF |
| CruV-402 | CFTLNN | AHVQG | ENQAYCSK | GSTGTGKSRFAN | IIDDY | ITTP | QLERRV |
| CruV-403 | AFTWNS | DHYQG | GLDFYVVK | QSGGVGKSKLQK | CNIP | VFSN | ASPDRW |
| CruV-404 | CFTLNN | PHLQG | SCIDYCKK | GKTGVGKTRYLY | IFDDI | FTSS | QLLRRV |
| CruV-405 | CGTLNN | PHLQF | QATDYCRK | GPPGTGKTRAAY | LLDEF | ISAN | ALRRRF |
| CruV-406 | LVTWFI | KHYQG | QAIAYCTK | GETGTGKSRKVY | LIDDF | ITSN | AFFRRV |
| CruV-407 | LVTWFI | KHYQG | QAIAYCTK | GETGTGKSRKVY | LIDDF | ITSN | AFFRRV |

| | Endonuclease domain | | | Helicase domain | | | |
|---|---|---|---|---|---|---|---|
| | Motif I | Motif II | Motif III | Walker A | Walker B | Motif C | Arg finger |
| CruV-408 | CFTVFG | VHIQG | KNTKYCTK | GGSGMGKTFSTY | LIDDF | ITSE | NLRRRA |
| CruV-409 | CWTWNN | PHLQA | QNYTYCSK | GTAGAGKTRAAF | ILDDF | ITSN | ALERRL |
| CruV-410 | | PHLQA | QNYTYCSK | GTAGAGKTRAAF | ILDDF | ITSN | ALERRL |
| CruV-411 | CFTWNN | PHLQG | QNYAYCSK | GTAGAGKTRAAF | ILDDF | ITSN | ALERRL |
| CruV-412 | LLTTKL | IHWQW | QACEYVIK | EEGGKGKSMLAR | IIDIA | IMTN | LSSDRW |
| CruV-413 | CFTLNN | PHLQG | DNYLYCSK | RTGGVGKSWLAT | CFDIS | VFAN | FSADRL |
| CruV-414 | VFTLNN | PHLQG | QARDYCRK | GPTGSGKSRAAF | VIDDY | ITSP | QLVRRV |
| CruV-415 | CFTSFN | EHAQG | QAIYYCEK | GSSRVRKTSVAY | IIDEI | FTSN | ALKNRF |
| CruV-416 | MLTIYN | PHLQG | SNTAYCKK | GTTGTGKSRWTK | VIDDL | ITNP | QLTRRI |
| CruV-417 | FGTVNG | PHLHF | KCLDYMKK | GESGSGKSLSAR | IIDDL | VTSN | ALLRRF |
| CruV-418 | IFTWWC | RHLQG | GSRDYCLS | GSAGKNKTRAAY | LIDEF | ITTN | AILRRI |
| CruV-419 | CFTMNN | PHLQG | ASIEYCRK | GPTGTGKSTRAR | ILDDV | VTSQ | ALRRRF |
| CruV-420 | VFRHSN | PHLQG | DSQNYCMK | GPTGTGKTRTIY | VLDEF | ITSP | QLIRRI |
| CruV-421 | SWTWHL | LHYQG | EAEFYCLK | EEGCTGKSHFAT | IFDIP | VFSN | LSKDRW |
| CruV-422 | SFTLNN | PHLQG | ENIVYCSK | GRTGSGKSKFAF | IVDEF | ITSQ | ALLRRI |
| CruV-423 | SWTWNK | LHYQG | DAEWYCLK | NEGCNGKSHFAT | IFDIP | VLAN | LSVDRW |
| CruV-424 | CFTHNN | RHLQG | ASIEYCKK | GPTGTGKSTLAR | IIDDL | VTSQ | ALERRF |
| CruV-425 | CFTINH | PHLQG | QAADYCKK | GASGTGKSKTAR | IIEEW | ITSN | PLKRRI |
| CruV-426 | VVTVNN | PHLQC | QNFIYCSK | | IIEDM | | |
| CruV-427 | MFTWFD | PHLQG | DNYNYCRK | GDTGCGKSVAIK | VMDDI | FISN | QLTRRI |
| CruV-428 | CYTLNN | LHCQG | QAADYCRD | GETGTGKSNAAY | LMEEF | FTSN | PFLRRI |
| CruV-429 | CFTLNN | PHLQG | QNIDYCFK | GKTGFGKTRCSM | ICDDF | ITTP | QLTRRV |
| CruV-430 | SFTWNK | LHYQG | DAEFYCTK | NIGCVGKTNFAT | IFDVP | ILAN | MSADRW |
| CruV-431 | MLTYMC | LHTHV | NRVKYLSK | RSGSVGKTDLVK | LIDIP | IFSN | LKIDRW |
| CruV-432 | CFTLNN | PHLQG | SNKVYCSK | GATGSGKSFAAH | VLDDY | VTSN | AACRRV |
| CruV-433 | CPVCAD | PHLQM | KSITYCTK | GPPGTGKSLLAA | ILDDF | ITTN | AILRRM |
| CruV-434 | FITSSN | PHLQF | SNILYVLK | ATGNVGKSWMAT | CFDIS | VFAN | FSQDRL |
| CruV-435 | SWTWNK | LHYQG | DAEFYCMK | ETGCTGKSYLAT | LFDIP | VLAN | LSLDRW |
| CruV-437 | CFTMWD | PHLQG | ACRAYCLK | GPPGTGKTHLAK | FIDEL | ITSN | PLKRRL |
| CruV-438 | AFTLNN | PHLQG | SSIAYCKK | GPTGTGKSRFAR | LIEDV | ITSN | PLLRRC |
| CruV-439 | CFTDYD | PHLQG | DCISYCEK | GKTGVGKTYAAI | IIDDF | VTSN | PLRRRF |
| CruV-440 | KLTTPA | QHSKA | LSLLCQEV | GESTGSTGRTVG | LSDEL | ALVP | TPNKRC |
| CruV-441 | VFTSFP | RHVHV | GWIDYCQK | | | IELI | |
| CruV-442 | AFTWNN | PHLQG | ASIEYCKK | GPTGSGKSRSAR | IVDDV | VTSN | PLMRRF |
| CruV-443 | CFTFNN | PHLQA | QASDYCKK | GESGSGKTELAK | IIDDF | VTCD | QIKSRF |
| CruV-444 | MFTIFQ | EHLQG | QCKTYCSK | QDGNSGKTLLAK | VMDYT | VFAN | LSIDRF |
| CruV-445 | CLTVYN | VHQHW | EAVMYFLK | GPTGTGKSLLCT | VLDEV | ITAP | QLRRRL |
| CruV-446 | CFTDYD | PHLQG | DCISYCEK | GKTGVGKTYAAI | IIDDF | VTSN | PLRRRF |
| CruV-447 | CFTLNN | RHLQG | ASIAYCKK | GETGSGKSRCAR | VIEDV | VTSN | ALFRRF |
| CruV-448 | AFTQWN | RHLQG | DNIDYCSK | GKSRAGKSHDAK | FENL | MCGN | HTERKE |
| CruV-449 | CYTVNN | EHIQG | QAKHYCMK | GPSGSGKSRIAE | ILDDF | FTSN | ALHRRI |
| CruV-450 | CLTGHF | PHLQG | QNIAYCSK | GPTGSGKSFSAR | ILDDF | ITSN | PILRRF |
| CruV-451 | SWTFNK | LHYQG | QAEFYCLK | TEGCTGKSSFAT | IFDVP | VFAN | LSMDRW |
| CruV-452 | SWTWNR | LHYQG | DAEWYCMK | DQGCKGKSHFAT | IFDIP | VLAN | MSVDRW |
| CruV-453 | LVYNFN | VHLQC | HATEYCKK | GPTGTGKSRDAN | FWDEF | FTAP | QFLRRL |
| CruV-454 | LVYNFN | VHLQC | HATEYCKK | GPTGTGKSRDAN | FWDEF | FTAP | QFLRRL |

| | Endonuclease domain | | | Helicase domain | | | |
|---|---|---|---|---|---|---|---|
| | Motif I | Motif II | Motif III | Walker A | Walker B | Motif C | Arg finger |
| CruV-455 | LVYNFN | VHLQC | HATEYCKK | GPTGTGKSRDAN | FWDEF | FTAP | QFLRRL |
| CruV-456 | LVYNFN | VHLQC | HATEYCKK | GPTGTGKSRDAN | FWDEF | FTAP | QFLRRL |
| CruV-457 | CFTIYD | IHLQC | QNRDYCKK | RTPGRGKTDCLS | VIDDF | VTSN | PIKARF |
| CruV-458 | CFTQKF | KHYQG | QNFEYVTK | GPSGCGKSHAAM | IMDDF | ITAI | QFRRRF |
| CruV-459 | CFTLND | QHLQG | QAKQYCEK | GPTGSGKSRRAL | VLDDF | ITSN | ALVRRL |
| CruV-460 | CLTGHY | PHLQA | DNIAYCSK | GKTGTGKSHAMR | IIDDF | VTSN | PILRRF |
| CruV-461 | MFTWNN | RHYQG | QADAYCSK | GGPGVGKTRDAL | VLDEI | IFLC | QVERRI |
| CruV-462 | VFTLNN | PHLQG | QASDYCKK | GAPGTGKSLCAR | LIEEW | VTSN | ALKRRF |
| CruV-463 | CVTINN | KHFQM | QNEDYCAG | RTSGVGKTSAVR | LLDDF | VTSN | PINNRF |
| CruV-464 | SWTFHM | LHYQG | NAEFYCMK | DTGCTGKTDFSI | IFDIP | VFSN | LSKDRW |
| CruV-465 | PFTYNY | EHVQG | QNRDYCAK | GASGTGKSYYAR | IDDFD | VTSN | PILRRF |
| CruV-466 | SITFNK | LHYQL | NAEFYCMK | ENGCKGKSHFAT | IFDIP | VLSN | LSADRW |
| CruV-467 | CFTLNN | PHLQG | ENRVYCSK | GPTGCGKSKSVR | LIEDI | VTSQ | ALERRF |
| CruV-468 | LLTYKS | RHTHV | DAKAYIGK | KRGNTGKSWLSR | IMDLA | VFAN | MSLDRW |
| CruV-469 | VFTCFQ | VHWQG | RARAYCKK | GPTGTGKSLWAH | IMEEF | LTSN | AAQRRL |
| CruV-470 | CFTVFF | PHFQG | QAIDYCHT | TDGRCGKTAFTR | LFDIE | VFSN | ATPDVW |
| CruV-472 | VFTLNG | SHLQG | HVRSYCVK | GPSNSGKTYQAN | INEF | VTCV | QFLRRI |
| CruV-473 | MSLAKN | | | GPSGTGKSHAAR | IIEDF | VTSN | PLLRRF |
| CruV-474 | CFTLNN | PHLQG | KAWDYCDK | VEGGTGKSTFAR | LFDFA | IFTN | LSRDRW |
| CruV-475 | FYTARE | LSTTG | RASRYSAR | GGTGTGKTRSVY | LLDEL | ITAP | QLLRRI |
| CruV-476 | FCTYSD | LHFHV | RTYEYCTK | GPPDCGKSFWVN | IYDDV | IVCN | TFTTRF |
| CruV-477 | CFTQHQ | LHLQG | QAINYCKK | GPSGVGKSREAA | IIDDY | ITSN | ALNRRI |
| CruV-478 | CFVHWI | DHYQG | DNYDYITC | YNFGTGKSHSAK | LIDEI | ITSN | CLKRRM |
| CruV-479 | LLTYKS | RHTHV | DAKAYIGK | KRGNTGKSWFSR | IMDLA | VFAN | MSLDRW |
| CruV-480 | CLTAYN | AHLQG | ANYKYCSK | GTTGVGKTFSVD | VIDDM | ITNP | QLMRRI |
| CruV-481 | VFTLNH | PDIQG | QSLDYCTK | GPTGTQKTRSSF | IFDDY | VTCP | QLSRRI |
| CruV-482 | CFTKFL | LHIQG | SAREYCRK | GASGAGKTRRAI | IIDDF | ITSN | ALRRRI |
| CruV-483 | CWTLNN | PHLQG | IAIEYCKK | | | | |
| CruV-484 | TITYNK | PHLQM | FNSDYCEG | KASGVGKTTAIR | LLDDF | VTSN | PICNRF |
| CruV-485 | DFRANE | RHYQG | GQAFYALK | HTGNLGKSFLVA | LIDMP | VFTN | LSADRW |
| CruV-486 | HLTYRS | DHTHV | KNVVYTTY | GETGLAKTQWAV | IFDDM | FTSN | AVLRRV |
| CruV-487 | CGTLHN | PHLQI | QAREYCMK | GKPNAGKTYKFY | LIDDF | VTTN | ALLRRI |
| CruV-488 | MFTIFQ | EHLQG | QCKAYCSK | QDGNSGKTVLAK | VMDYT | VLAN | LSQDRW |
| CruV-489 | | PHLQG | QNERYCSK | GRPGCGKTRQCY | LLDEF | LTSN | ALRRRI |
| CruV-490 | CFTVNN | PHLQG | EAWDYCLK | GPTGSGKSRAAF | ILDDF | ITSC | QLLRRI |
| CruV-491 | | KHLQG | ASIRYCKK | GPTGTGKSHFAR | IIEEL | VTSN | PLRRRF |
| CruV-492 | CGTLHN | PHLQI | QAREYCMK | GKPGCGKTYKFY | LIDDF | VTTN | ALLRRF |
| CruV-493 | | | ANFEYCSK | GVPGCGKTEYAL | LIDDF | ITTN | ALFRRI |
| CruV-494 | CFTINN | VHLQG | QNFLYCAK | GPSGTGKTQFAL | VFDDM | FTHN | AIERRY |
| CruV-495 | | | DSNRYCRK | GKTGRGKTRAVY | LFDDF | FTTP | QLSRRV |
| CruV-496 | | AHTHV | RLWEYHQK | GPTGIGKTQWAL | VFDDT | ITSN | AIKRRC |
| CruV-497 | PWTLNN | PHLQGSHI TG | QAINYCKK | GKPGTGKTRKAY | LLDEF | VTSN | ALRRRF |
| CruV-498 | VFTINN | LHLQG | LARNYVLK | GDTNIGKTSFAK | VFDDM | FTCN | AILRRV |
| CruV-499 | SLTVNN | PHLQC | SLHKYCEK | | | | |
| CruV-500 | VFTLNN | PHLQS | NTTQYYPT | SCSNGRCSQIHL | | | PMSNRK |

| | Endonuclease domain | | | Helicase domain | | | |
|---|---|---|---|---|---|---|---|
| | Motif I | Motif II | Motif III | Walker A | Walker B | Motif C | Arg finger |
| CruV-501 | LYTKFN | PHIQG | QNFVYCSK | GSSGIGKTHYAA | IFDDM | FTHN | AIERRL |
| CruV-502 | CFTIHR | VHLQG | SNWDYCTK | GATGTGKTYYAS | IIDDY | ITSA | QLTDRC |
| CruV-503 | FGTCNN | RHLQF | QNITYCSK | GESGSGKSTLAD | CQEFR | FTQC | QLLRRV |
| CruV-504 | CFTLNN | PHLQG | QARDYCRK | GETYSGKTHQAF | LLDDF | ITTN | ALKRRI |
| CruV-505 | CFRANA | LHHQG | QAWAYSTK | GDSNCGKTHFVL | VFDDL | FTHN | AIDRRV |
| CruV-506 | CFRANA | LHHQG | QAWAYSTK | GDSNCGKTHFVL | VFDDL | FTHN | AIDRRV |
| CruV-507 | CFTINN | PHIQG | QNMLYCKK | GPSGTGKTQWAL | VFDDM | FTTN | AIERRY |
| CruV-508 | VFTLNN | PHLQG | QASDYCKK | GAPGTGKSLCAR | LIEEW | VTSN | QALRRR |
| CruV-509 | LYTKFN | PHIQG | QNFVYCSK | GSSGIGKTHYAA | IFDDM | FTHN | AIERRL |
| CruV-510 | TFRKSA | LHFQG | QNWAYATK | GPSNTGKTSFVK | IFDDM | FTHN | AIDRRV |
| CruV-511 | CFSHWN | HHYQG | ENHVYCTK | GVKGAGKTQDAM | IIDEA | ITMN | YPVRHP |
| CruV-512 | HLTYPS | PHTHA | ACWNYHEK | GRTGIGKTQWAL | VFDDL | FTSN | AIARRC |
| CruV-513 | | LHYQG | QVWAYSTK | GEPGSGKTEFAR | IIEDF | | |
| CruV-514 | TFVLNN | KHFQG | TNAKYCSK | GPPGTGKSFMAW | IIDDF | ITTN | QLRRRV |
| CruV-515 | VFTLNN | PHLQG | QASDYCKK | | | | |
| CruV-516 | SLVVER | GHSWM | RHRAYFSQ | GGSGIGKSMFAQ | IFDDC | ITSN | AIERRM |
| CruV-517 | VFTLNN | PHLQG | QASDYCKK | | | | |
| CruV-518 | IFRMSN | PHLQG | QNEKYCSK | GPPGCGKTELAK | LIDDF | ITSN | ALFRRF |
| CruV-519 | | THWWA | | GAAGLGKTHAAL | FNEA | LTTN | AFRRRG |
| CruV-520 | CFTVNN | PHLQG | QNRDYCLK | GSPGTGKSVWAR | LIEDV | VTSN | ALRRRF |
| CruV-521 | CFTLND | KHYQG | QARDYCMK | GKPGSGKTQLFW | LIDDF | ITTN | ALKRRV |
| CruV-522 | | LHYQG | QNWAYCTK | GPPGVGKSEFAR | ILEDF | ITTN | |
| CruV-523 | CVTYFG | LHCQA | ENIVYCTK | GETGGGKTRAAF | IFDDF | ITCP | QLLRRI |
| CruV-524 | VFTWND | KHYQG | QAREYAMK | GPPGTGKSRCVR | LIDDF | ITTN | ALKRRI |
| CruV-525 | CFTLND | KHYQG | QARDYCMK | GTPGTGKTQNFW | LIDDF | VTSN | ALKRRF |
| CruV-526 | LLTQQG | AHAHI | RAVEYVKK | GPSRTGKSYHAR | LIEDL | VTSN | PICNRF |
| CruV-527 | CFTWFK | EHLQC | RARLYCMK | GKPGTGKTRRAY | LLDDF | ITTN | ALKRRF |
| CruV-528 | CFTWFK | EHLQC | RARLYCMK | GKPGTGKTRRAY | LLDDF | ITTN | ALKRRF |
| CruV-529 | VFTTNN | PHLQG | EAATYCKK | QVAVRNGGFPKC | ILDDW | ITTT | QLLRRV |
| CruV-530 | ISTPTL | | AAIQYYRV | GASRTGKSTHMK | VHDCA | IDSQ | AFCKRF |
| CruV-531 | VLTHNN | PHVQA | GGVDYILR | GVSGSGKSHTAR | INDFY | ITSN | AVERRI |
| CruV-532 | | LHWQV | QSLDYCNK | GPSGAGKSTRCR | IIEDL | ITTN | RWARRF |
| CruV-534 | VFRSNA | LHHQG | QAWAYSVK | GPPGTGKTDFAY | ILDEF | ITTN | ALFRRI |
| CruV-535 | VFTARA | LHYQG | SNWTYCTK | GPPGTGKTDFAY | IIDEF | ITTN | ALFRRI |
| CruV-536 | VFTLNN | PHFQG | QAIEYCKK | GKTGTGKTYKAF | LLDEV | ITTN | ALLRRF |
| CruV-537 | CWTLNN | PHLQG | | | | | |
| CruV-538 | IVVINN | PHFQC | NNQDYCKK | GPTGLGKSKLAR | LIEDF | VTSN | PLLRRF |
| CruV-539 | FLTKNN | PHLHA | SCITYCKK | GETGFGKTSFAF | IIDEI | ITAP | HLLRRI |
| CruV-540 | FFTLFV | LHIQG | FGMNYCEK | GPTSTGKTTYAE | IDDLT | LTSN | |
| CruV-541 | VLTCNN | PHLQC | | GKPGTGKTHAAT | IIDDF | VTSN | AIMRRC |
